# Opposing copy number variation dynamics accompany adaptation to glucose and galactose in diploid yeast

**DOI:** 10.1101/2025.10.01.679718

**Authors:** Prachitha Nagendra, Saket Choudhary, Supreet Saini

**Affiliations:** Department of Chemical Engineering, Indian Institute of Technology Bombay, Mumbai, India 400 076; Koita Centre for Digital Health, Indian Institute of Technology Bombay, Mumbai, India 400 076

**Keywords:** S. cerevisiae, adaptation, glucose, galactose, duplication, deletion, SNPs

## Abstract

Adaptation to new environments arises from multiple forms of genetic variation, including single-nucleotide polymorphisms (SNPs) and structural variants such as copy number variations (CNVs). Although both mutation classes have been implicated in evolutionary change, their relative contributions during adaptation remain incompletely understood and may depend strongly on environmental context. Here we investigated genome evolution during experimental adaptation of diploid *Saccharomyces cerevisiae* populations propagated for 1200 generations in two alternative carbon sources, glucose or galactose. Whole-genome sequencing of replicate populations at multiple evolutionary time points revealed relatively few SNPs but widespread copy number variation across the genome. Strikingly, CNV dynamics differed markedly between environments. Populations evolved in glucose exhibited extensive early deletions followed by later genomic remodeling that reduced the overall deletion burden. In contrast, populations evolved in galactose accumulated persistent subtelomeric duplications throughout the experiment. Many loci displayed opposing copy number changes in the two environments, with genes deleted during evolution in glucose frequently duplicated during evolution in galactose. These contrasting trajectories indicate that structural genomic changes repeatedly arise during adaptation and that their direction and persistence depend strongly on environmental conditions. Together, our results illustrate how distinct genomic routes can emerge during adaptation of genetically identical populations and highlight the importance of considering multiple classes of mutations when studying evolutionary responses to environmental change.

**Impact statement:** This study demonstrates that copy number variations (CNVs), rather than single-nucleotide polymorphisms, dominate genome evolution during adaptation of diploid *Saccharomyces cerevisiae* to alternative carbon sources. By evolving genetically identical populations in glucose and galactose, the work reveals that environmental context drives strikingly opposing CNV trajectories, widespread deletions in glucose and persistent duplications in galactose, often targeting the same genomic loci in opposite directions. These findings provide direct evidence that structurally mediated gene dosage changes are a primary and environment-dependent mechanism of adaptation, reshaping metabolic networks and specialization strategies. More broadly, the study challenges the prevailing SNP-centric view of adaptive evolution and highlights CNVs as a predictable and major axis of genome remodeling in microbial systems.

## Introduction

Adaptation arises from genetic variation that alters cellular physiology and organismal fitness (1-3). Historically, single-nucleotide polymorphisms (SNPs) have been viewed as the primary substrate of adaptive evolution because they are abundant, arise continuously, and can fine-tune gene function through changes in coding or regulatory sequences (1, 4-6). Consequently, many microbial experimental evolution studies have focused on identifying beneficial point mutations and tracing their dynamics during adaptation (5, 7, 8). However, organisms and genomes can also evolve through structural changes, including gene duplications, deletions, which alter gene dosage and can affect large genomic regions simultaneously (9, 10). These distinct classes of mutations differ not only in their molecular properties but also in their rates of occurrence, phenotypic consequences, and evolutionary dynamics (9, 11-14).

Structural variants can play important roles in evolutionary adaptation across diverse systems. Gene amplifications have been associated with resistance to metals, toxins, and drugs in microbes, while deletions and other large-scale rearrangements can rapidly reconfigure metabolic pathways or regulatory networks (12, 15-18). Because copy number changes can simultaneously affect multiple genes and regulatory elements, they may enable rapid physiological shifts that would require multiple point mutations to achieve (12, 13, 19, 20). At the same time, point mutations can provide more precise adjustments of gene function, refining adaptive phenotypes following larger genomic changes. (1, 2, 6, 19, 21). Consequently, adaptation may proceed through different combinations of SNPs and structural variants depending on the organism, genetic background, and ecological context (2, 22-24).

Despite extensive documentation of both mutation classes, relatively few studies have directly compared the contributions of SNPs and CNVs to adaptation within the same experimental system (5, 10, 13, 17, 19). Moreover, it remains unclear whether the genomic routes taken during adaptation depend strongly on environmental context, particularly when environments differ only subtly in their metabolic demands (2, 21, 23, 25, 26). Addressing this question requires experimental designs in which genetically identical populations evolve independently under distinct but comparable conditions, allowing evolutionary trajectories to be contrasted across environments while controlling for initial genetic background (27-29).

Experimental evolution with microbes provides a powerful framework for addressing these questions because populations can be propagated for hundreds to thousands of generations under controlled conditions, and genomic changes can be monitored over time (30-32). In the budding yeast *Saccharomyces cerevisiae*, carbon source availability strongly influences cellular physiology and gene regulation, particularly through the regulatory networks governing glucose repression and galactose utilization (33-36). Glucose and galactose therefore provide closely related yet distinct metabolic environments that engage overlapping but different transcriptional and metabolic programs, making them a useful system for examining how environmental context shapes evolutionary trajectories (37-41).

Here, we evolved replicate diploid populations of *S. cerevisiae* for 1200 generations in either glucose or galactose and tracked genomic changes across four evolutionary time points. By combining whole-genome sequencing with phenotypic assays, we compared the accumulation of SNPs and CNVs and examined how genomic changes correspond to patterns of specialization and cross-environment performance. We find that while SNPs occur rarely across populations, extensive copy number variation emerges repeatedly during evolution in both environments. Strikingly, the direction and persistence of these copy number changes differ between environments: glucose-evolved populations exhibit extensive early deletions whereas galactose-evolved populations accumulate persistent duplications across many genomic regions. Many of the same loci experience opposing copy number changes in the two environments, suggesting that gene dosage changes in shared metabolic modules are resolved differently under distinct selective pressures. Together, these results highlight the prominence of copy number variation in shaping genome evolution in this system and illustrate how environmental context can lead to divergent structural trajectories even among genetically identical populations.

## Results

### Adaptive dynamics of glucose- and galactose-evolved populations

We evolved six replicate diploid *Saccharomyces cerevisiae* populations in either 0.5% glucose or 0.5% galactose for 1200 generations. Fitness was assayed every 300 generations, while whole-genome sequencing of all six glucose- and six galactose-adapted populations, each at four time points was performed to identify the underlying genetic basis of adaptation **(Figure 1A)**.

**Figure 1.**
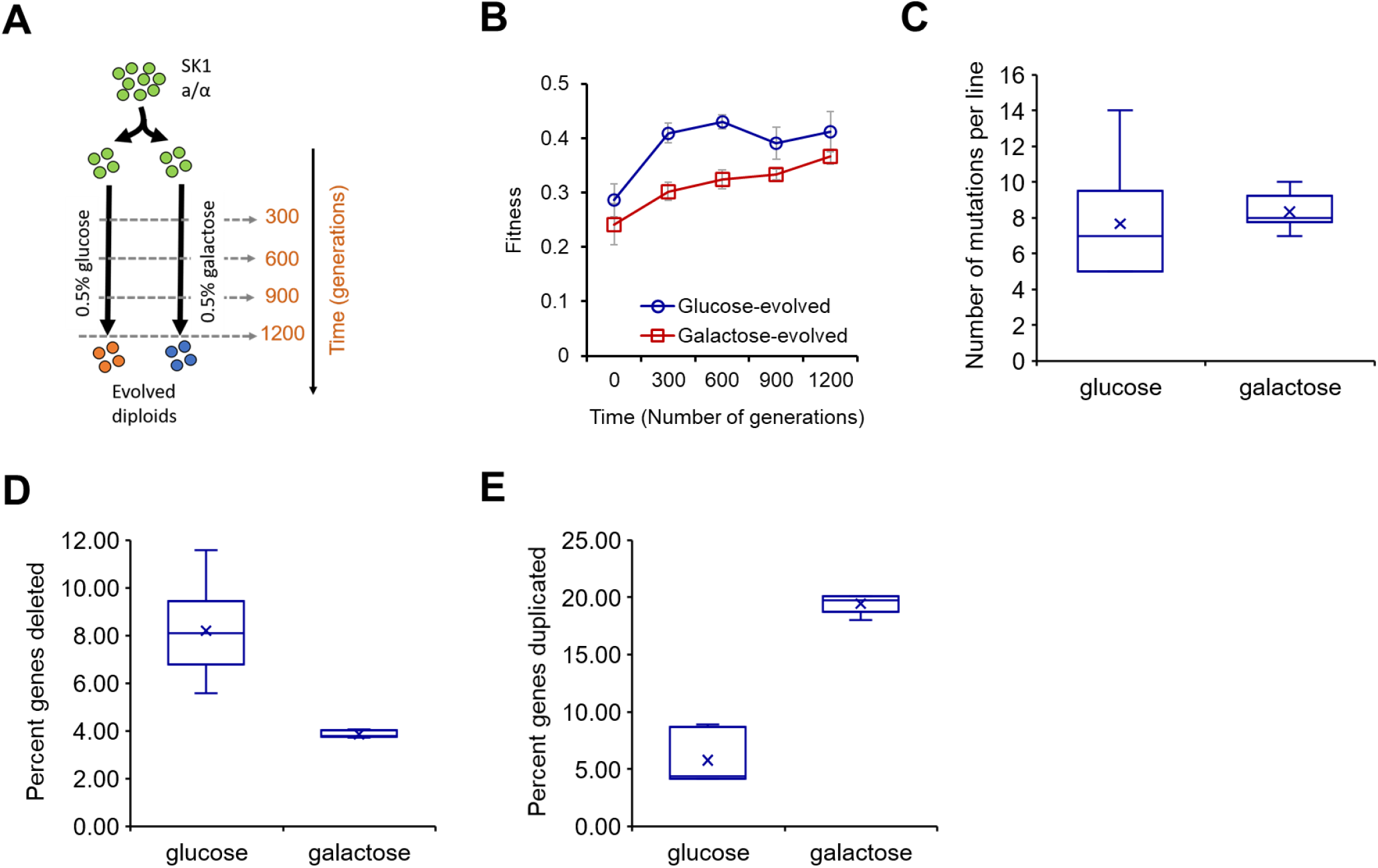
Experimental design, fitness gains, and a genomic overview of adaptation. **(A)** Schematic of the experimental evolution. Six independent diploid *S. cerevisiae* populations were propagated for 1200 generations in either glucose or galactose. Populations were sampled at 0, 300, 600, 900, and 1200 generations for phenotyping and whole-genome sequencing. **(B)** Home-environment fitness trajectories (mean relative growth rate ± s.e.m., *n* = 6 lines per environment) show steady improvement in both media; glucose-evolved lines (blue) rise rapidly and then plateau, while galactose-evolved lines (red) increase more gradually. **(C)** SNP accumulation is rare. The box plots indicate the total number of SNPs including variants that are fixed (allele frequency is either 0 or 1 depending on reference or alternate allele being fixed) and variants that are present but not fixed (allele frequency greater than 0.2 in case of de novo variants) in glucose and galactose environments. (joint calling across time points; stringent depth and allele-frequency filters; visual confirmation in IGV). **(D)** and **(E)** Copy-number variation (CNV) dominates genome change and differs by environment. Box-and-whisker plots summarize CNV across 1200 generations. **(D)** deletions only and **(E)** duplications only. Glucose-evolved lines carry abundant deletions, whereas galactose-evolved lines show persistent, higher duplication loads, exhibiting environment-specific CNV strategies. In figures C-E, boxes show interquartile range with median; whiskers span full range; ✕ indicates mean.

In both environments, populations initially experienced a fitness increase in their home environment, which later fluctuated mildly. Upon assaying fitness by preculturing in the same medium, we see that by 1200 generations, glucose-adapted populations grew ~20–30% faster than their ancestor in glucose, while galactose-adapted populations improved ~15– 25% in galactose **(Figure 1B and Supplement Figure S1)**.

### Fitness changes correlate with environment-specific copy number variation dynamics

Despite the different selective environments, we observed single-nucleotide polymorphisms (SNPs) to be rare across all lines and time points (**Figure 1C and TableS1**). Contrastingly, we found the dominant mode of genomic change was copy-number variation (CNV), which occurred repeatedly and independently in multiple replicate lines **(Figure 1D-E and TableS2 and S3)**.

Adaptation was accompanied by widespread CNVs whose signatures differed sharply between the two environments. Overall, glucose lines carried roughly equal numbers of deletions and duplications, while galactose lines accumulated several-fold more duplications than deletions, a bias that remained consistent across all four time points **(TableS4 and S5)**.

However, duplications and deletions exhibited strong contrasts, depending on the environment in which the populations evolved. In glucose-evolved populations, deletions were frequent, particularly in telomeric and subtelomeric regions of all chromosomes, except chromosomes I and II **(Figure 2 and Supplement Figure S2)**. The extent of these deletions was most pronounced in the early intervals (first 300 generations) but declined with further evolution. This suggests that early adaptive gains involved a large number of deletions that were later refined or compensated by subsequent changes or that the populations carrying the deletion were lost with time.

**Figure 2.**
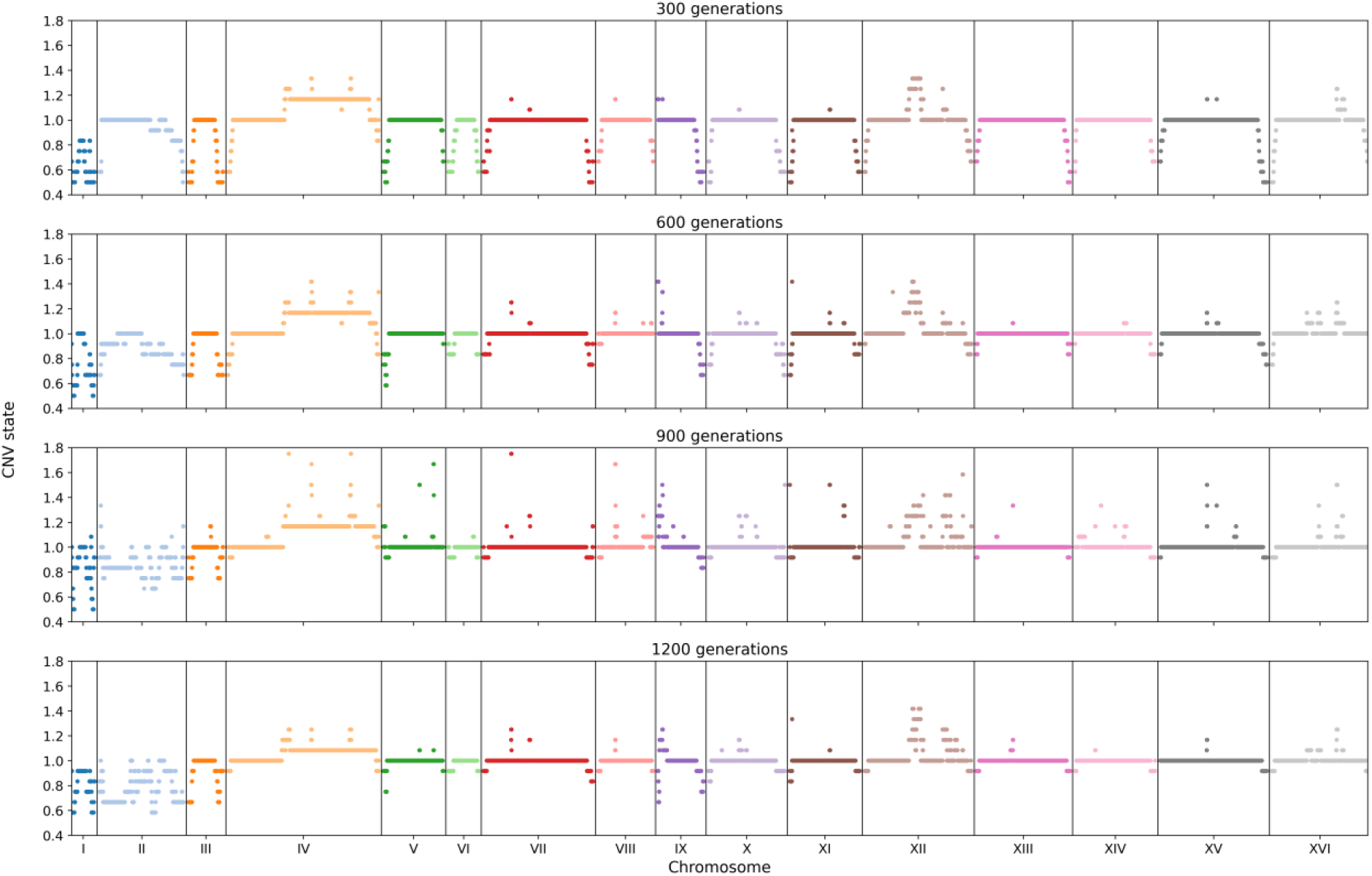
CNV dynamics in glucose-evolved populations across 1200 generations. CNV states for the six glucose-evolved lines at 300, 600, 900, and 1200 generations. The *x*-axis lists the 16 *S. cerevisiae* chromosomes. Each point is a 500-bp genomic window; values are normalized to the ancestor such that state = 1 indicates no change, <1 indicates deletion, and >1 indicates duplication. The colored horizontal bars denote the mean CNV state for that chromosome at that time point. At 300 generations, windows with deletion states (≈0.6–0.95) are widespread and concentrated near chromosome ends, revealing early, extensive telomeric loss. Through 600–900 generations, the deletion burden decreases (mean states trend toward 1). By 1200 generations, most chromosomes sit close to state 1 with fewer and weaker deletions, consistent with refinement/compensation of early structural changes. Overall, the trajectory indicates that early telomeric deletions dominate adaptation in glucose, followed by later remodeling that reduces deletion extent and introduces targeted duplications.

In contrast, galactose-evolved populations exhibited widespread telomeric and sub-telomeric duplications **(Figure 3 and Supplement Figure S3)** except in chromosome II. These duplications were evident from as early as 300 generations and were maintained or expanded throughout 1200 generations, indicating persistent selective benefit in the galactose environment. The opposing trend in glucose- and galactose-evolved populations remain even after removal of telomeric regions from each chromosome **(Supplement figure S4a and b)**.

**Figure 3.**
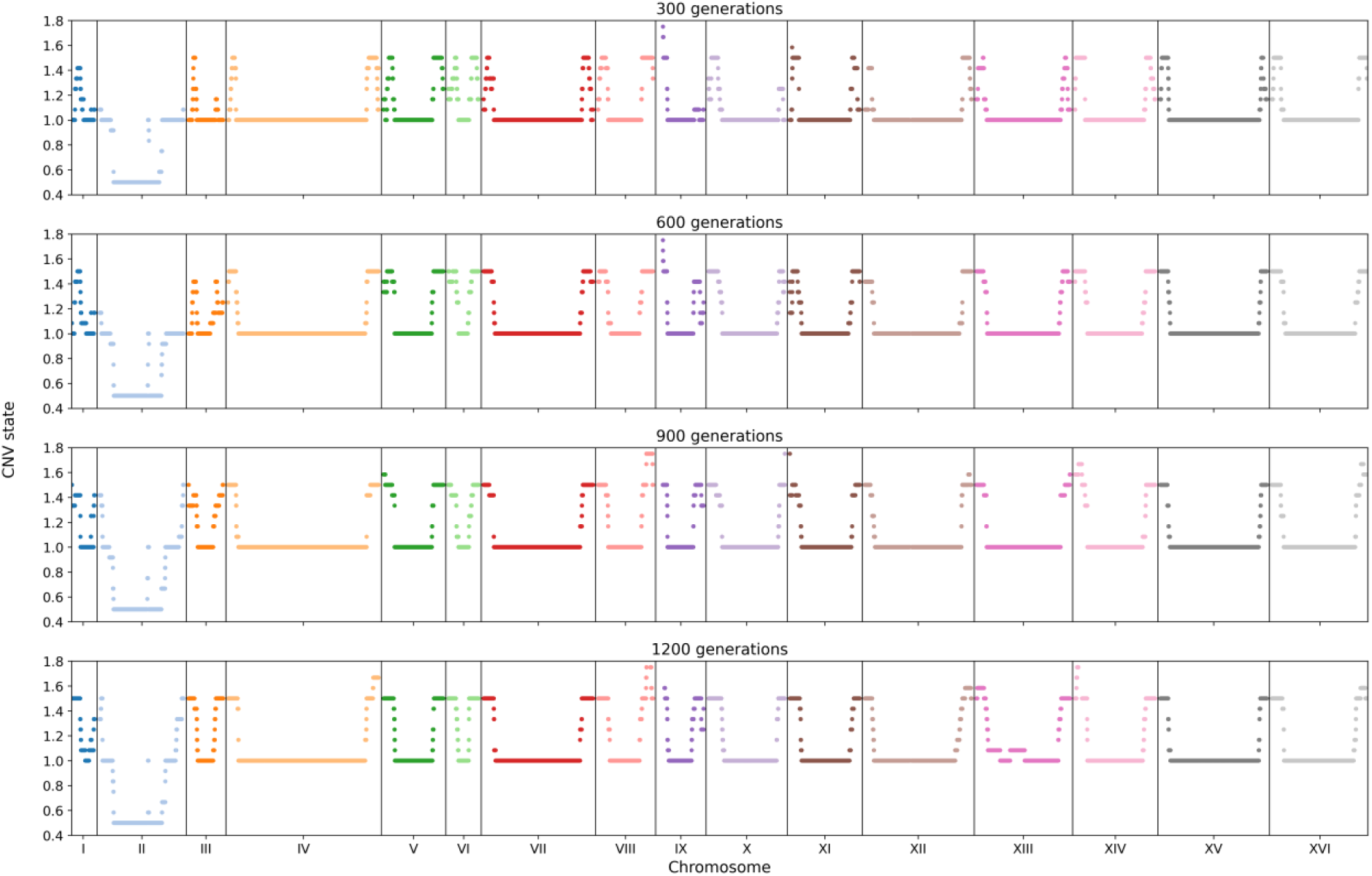
Persistent subtelomeric duplications in galactose-evolved populations. CNV states for the six galactose-evolved lines at 300, 600, 900, and 1200 generations. The *x*-axis lists the 16 *S. cerevisiae* chromosomes. Each point represents a 500-bp window; state = 1 indicates no change, <1 deletion, >1 duplication. The colored horizontal bar marks the mean CNV state for that chromosome at that time point. Across all time points, windows with duplication states (>1.0–1.7) are abundant and cluster near chromosome ends, revealing early-onset, persistent subtelomeric duplications under galactose selection except in chromosome II. Unlike the glucose-evolved trajectories (Figure 2), deletions are rare and mean chromosome states remain consistently >1 from 300 to 1200 generations, indicating sustained duplication loads rather than deletion-first remodeling. Together, these patterns demonstrate a duplication-biased CNV program in galactose that is stable over evolutionary time and concentrated in telomeric neighborhoods enriched for transport/metabolic modules.

Strikingly, 474 genes that were deleted in at least one of the replicate in glucose-evolved lines were found duplicated in at least one of the replicate in galactose-evolved lines during some stage of the evolution experiment **(Figure 4, Supplement Figure S5a and S5b)**.

**Figure 4.**
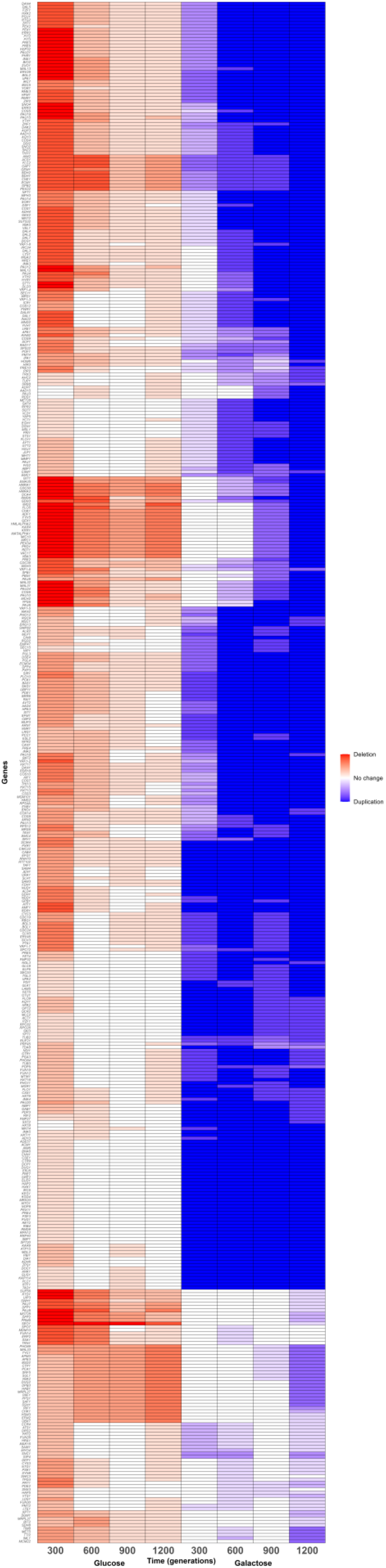
Divergent copy number evolution in glucose versus galactose. 474 genes deleted in at least one of the replicates during adaptation in glucose appear duplicated during adaptation in galactose in at least one of the replicates. This highlights environment specific resolution of selection on the same loci over evolutionary time. Each row is a gene. Columns show four time points in glucose (300, 600, 900, 1200 generations) followed by four time points in galactose (300, 600, 900, 1200 generations). Color encodes the direction of copy number change relative to the ancestor: red indicates deletions, blue indicates duplications, white indicates no detected change, the intensity of color indicates the number of replicates showing deletion/duplication for a given gene (scale bar at right).

### Specialization trajectories are mirrored in genomic remodeling

Adaptation to glucose increases fitness. However, we asked if there were particular costs associated with this adaptation. Specifically, we investigated the cost of adaptation in the ability of the cells to switch environments and exhibit growth. To quantify this, we compared the glucose-evolved lines’ growth to the glucose ancestor, when the cells were first pre-cultured in galactose. Glucose-evolved populations showed a strong penalty or growth defect at 300 generations because of prior exposure to galactose, consistent with the cost of specialization on glucose **(Figure 5)**. However, this penalty declined sharply by generation 900, reaching nearly zero, before increasing slightly again later in the experiment. This dynamic trajectory suggested a transient phase of specialization followed by the acquisition of mutations conferring generalist performance.

**Figure 5.**
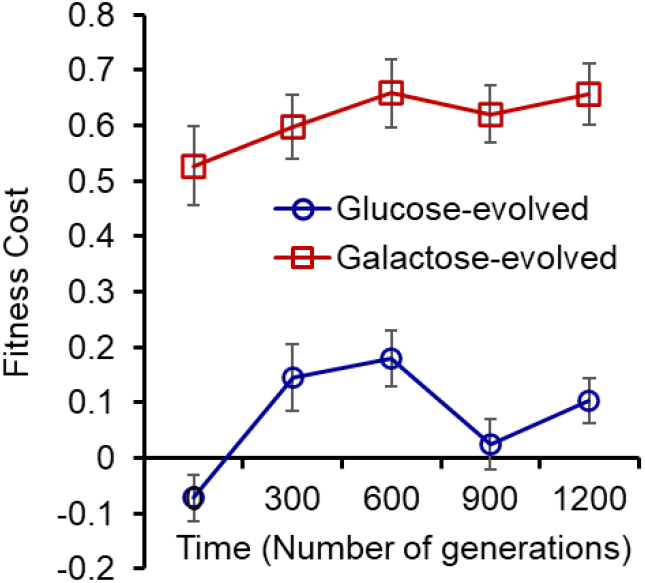
History-dependent fitness costs after cross-environment preculture. For each evolved background, fitness cost was computed as the relative decrease in composite growth rate in the home environment when cells were precultured in the non-home carbon source. Composite growth rate integrates lag and exponential growth from OD time courses (Methods). Lines show the mean cost across six replicate populations at each evolutionary time point (300, 600, 900, 1200 generations); blue, glucose-evolved populations precultured in galactose and assayed in glucose; red, galactose-evolved populations precultured in glucose and assayed in galactose. Glucose-evolved populations display a transient specialization: a pronounced cost early (peaking by ~600 generations) that collapses by 900 generations and remains low thereafter, indicating reduced sensitivity to preculture history. In contrast, galactose-evolved populations show a persistent, increasing cost across the experiment, consistent with deepening specialization to galactose and strong history dependence following glucose exposure.

The genomic data shows that early adaptation coincided with extensive telomeric deletions, including in regions encompassing genes involved in galactose metabolism and alternative carbon source utilization. Pathway-level analyses revealed frequent targeting of galactose metabolism, pyruvate metabolism, and fructose and mannose metabolism **(Supplement Figure S6)**. These deletions likely conferred rapid specialization to glucose but incurred fitness costs when preculture history involved galactose. Later in evolution, however, the extent of deletions decreased, suggesting the appearance of compensatory CNVs (including duplications) that may have restored broader metabolic flexibility and enabled the observed decline in fitness penalties.

In contrast, galactose-evolved populations showed steadily increasing penalties across the 1200 generations, consistent with a trajectory of deepening specialization **(Figure 5)**. The underlying CNV patterns reinforced this phenotype: galactose lines accumulated persistent duplications in telomeric and subtelomeric regions, often in loci involved in fructose and mannose metabolism, starch and sucrose metabolism, and galactose metabolism **(Supplement Figure S7)**. Rather than being compensated, these duplications were maintained and amplified over time, reinforcing a genetic architecture that promoted strong specialization and increasing costs of cross-environment exposure.

### Physiological history dependence and its genomic correlates

We further tested the influence of short-term growth history on fitness. We grew galactose-evolved populations briefly in glucose (~15 generations) and then returned them to galactose media. The transferred population displayed reduced growth rates in galactose, but this penalty was mitigated when, following growth in glucose, cells were re-exposed to galactose for 7 or 15 generations **(Figure 6A)**. This plasticity suggests that the specialized physiology of galactose-adapted lines is sensitive to immediate history but can be restored with sufficient exposure to galactose. Genomic signatures provide a plausible explanation: the duplications enriched in galactose-evolved lines were heavily biased toward genes encoding metabolic enzymes and transporters, creating a cellular state optimized for galactose but disrupted by prior glucose exposure.

**Figure 6.**
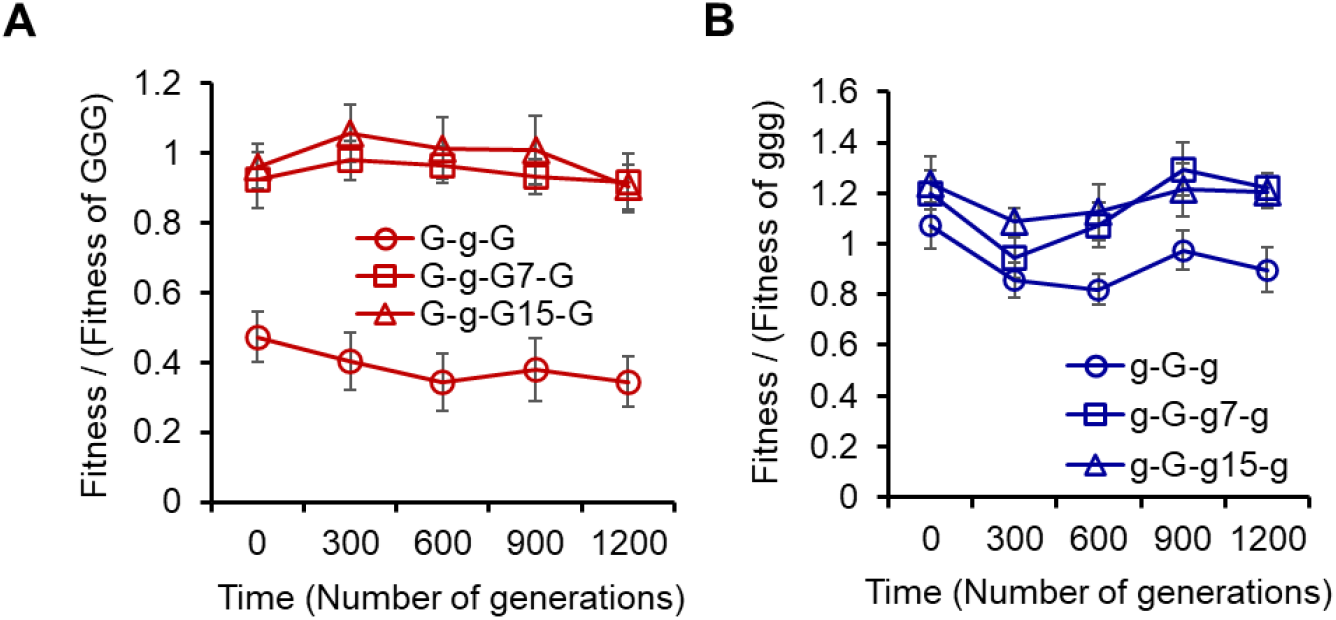
Short-term history effects and recovery in home environment. For each evolutionary background, we measured growth in the home environment after a brief exposure to the non-home carbon source, with or without a recovery phase back in the home medium. Fitness is shown as a ratio to the corresponding home-only control (for (A): divided by fitness of ggg; for (B): divided by fitness of GGG), where “g/G” denote glucose/galactose and the string indicates evolution → preculture → (optional recovery duration) → assay. Thus, g-G-g = glucose-evolved, precultured in galactose, assayed in glucose; g-G-g7-g / g-G-g15-g include ~7 or ~15 generations of recovery in glucose before assay. Analogously, G-g-G, G-g-G7-G, G-g-G15-G are for galactose-evolved lines. Lines show the mean across six replicate populations at each evolutionary time point (0, 300, 600, 900, 1200 generations). Composite fitness integrates lag and exponential growth from OD time courses (Methods). **(A)** Galactose-evolved populations (right) show a strong penalty when assayed immediately after glucose preculture (G-g-G), whereas 7–15 generations of re-exposure to galactose (G-g-G7-G, G-g-G15-G) restore fitness toward the home-only baseline across all evolutionary time points. **(B)** Glucose-evolved populations exhibit history dependence early: performance in glucose after galactose preculture depends on recovery duration (g-G-g < g-G-g7-g ≈ g-G-g15-g) through ~600 generations, after which the curves converge, indicating a loss of history dependence.

Glucose-evolved populations behaved differently. During the first 600 generations, the duration of glucose preculture (7 vs. 15 generations) influenced measured growth rates, consistent with continued sensitivity to short-term history **(Figure 6B)**. However, beyond generation 600, this effect disappeared: fitness became independent of preculture duration. This history-independence parallels the genomic trend in which extensive deletions present early in evolution diminished later, with glucose lines acquiring duplications that likely stabilized metabolism and reduced sensitivity to environmental history.

### Predictability of evolutionary outcomes

To investigate the contributors to fitness of the evolved populations in different conditions, we analyzed our dataset using a linear model incorporating ancestor fitness, preconditioning environment, recovery duration, evolutionary time, and their interactions. We ran an ANOVA analysis which showed fitness of the ancestral strain as the strongest predictor of evolved fitness (η^2^ ≈ 0.71, p < 10^−62^), demonstrating that the initial genotype largely sets the baseline for subsequent adaptation. Evolutionary time also contributed significantly to fitness changes (η^2^ ≈ 0.22, p < 10^−28^), consistent with the gradual accumulation of adaptive changes observed in both glucose- and galactose-evolved lines.

Other main effects, including preconditioning environment and recovery duration, were individually non-significant. However, modestly significant interactions with evolutionary time were detected (preconditioning × evolution: p ≈ 0.0014; recovery × evolution: p ≈ 0.027), indicating that environmental context can subtly modulate the pace or trajectory of adaptation. Higher-order interactions contributed minimally to overall variance in fitness.

Linear regression analysis further supported these conclusions. The coefficient for ancestral fitness was large and highly significant (β = 1.324, p < 0.001), whereas the effect of evolutionary time per generation was small but detectable (β = 8.7 × 10^−5^, p < 0.001).

Interaction terms were largely negligible, emphasizing that while environmental history and short-term physiological conditions can influence fitness to a limited extent, the initial genetic background dominates subsequent evolutionary outcomes.

These findings suggest that the evolutionary response in our experiment is highly predictable at the level of gross fitness changes: the starting genotype establishes a strong constraint on the magnitude and direction of adaptation, while environmental factors such as preconditioning and recovery modulate the response only subtly. When interpreted alongside our genomic analyses, these results imply that predictable structural changes, particularly environment-specific CNVs in key metabolic loci, act within the bounds set by the ancestral genotype to produce incremental but repeatable fitness gains.

## Discussion

Adaptation can proceed through multiple forms of genetic variation, including point mutations and structural variants. In this study, we compared the genomic trajectories of replicate diploid *Saccharomyces cerevisiae* populations evolving in two related carbon environments. Across 1200 generations, we observed relatively few SNPs but extensive copy number variation in both environments. Rather than focusing on the adaptive value of individual mutations, our results highlight how different classes of mutations contribute unevenly to genome evolution and how environmental context shapes the structural trajectories that emerge during adaptation.

A striking feature of our experiment is that copy number changes follow distinct and reproducible patterns in the two environments. Populations evolving in glucose exhibit widespread deletions early in evolution that diminish over time, whereas populations evolving in galactose accumulate persistent duplications across many genomic regions. These contrasting trajectories suggest that structural genome remodeling is not random but reflects environment-dependent selective pressures acting on shared metabolic and regulatory modules. Notably, many of the loci affected by deletions in glucose are duplicated in galactose-evolved populations, indicating that the same genomic regions experience opposing dosage changes depending on the metabolic context. Such opposing trajectories illustrate how related environments can impose divergent evolutionary pressures on common genetic targets.

The predominance of CNVs in our dataset is consistent with growing evidence that structural variants can play important roles during microbial adaptation (13, 19, 42). Gene duplications and deletions can arise at higher rates than point mutations and can alter gene dosage across multiple loci simultaneously, enabling rapid physiological shifts (11, 12, 18, 43). In diploid organisms, structural variation may be particularly influential because changes in copy number can modify gene dosage immediately (44-48). In this context, CNVs may provide a flexible genomic substrate through which populations explore alternative physiological strategies during adaptation (9, 24).

Our results also reveal that genome evolution in this system is shaped not only by mutation supply but also by environmental context and the architecture of cellular regulation. Glucose and galactose engage overlapping metabolic pathways but differ substantially in regulatory control and carbon utilization strategies (34, 35, 40, 41). The opposing CNV patterns observed here suggest that evolutionary responses may preferentially target genomic regions whose dosage influences these metabolic programs (10, 16, 17, 49). It has been reported that sub-telomeric regions in *S. cerevisiae* contain carbohydrate uptake genes and are known to evolve faster than other regions of the chromosome (50). Thus, rather than converging on identical genetic solutions, populations evolving in related environments can remodel similar genomic regions in opposite directions (25, 51).

More broadly, these findings illustrate how examining multiple classes of genetic variation can provide a more complete view of evolutionary dynamics (19, 24, 52, 53). Studies of experimental evolution have traditionally emphasized point mutations, but structural variants may represent an additional and sometimes prominent axis of genome evolution (6, 18, 30, 54). By comparing adaptation across two closely related environments, our work highlights how environmental context can influence which genomic changes accumulate and how those changes reshape genome architecture over evolutionary time.

Together, our results show that genetically identical populations evolving under distinct nutrient conditions can follow divergent structural genomic trajectories, even when adapting over the same timescale. Understanding how different classes of mutations contribute to these trajectories will be important for developing a broader view of the mechanisms underlying evolutionary adaptation in microbial systems (23, 24, 53, 55).

## Methods

### Strains used and experimental evolution

A diploid SK1 (*a/α*) strain was generated by mating two isogenic haploids:

- *SK1 MATa ARS314::TRP1 ho::LYS2 lys2 ura3 leu2::hisG his3::hisG trp1::hisG*
- *SK1 MATα ARS314::LEU2 ho::LYS2 lys2 ura3 leu2::hisG his3::hisG trp1::hisG*

For each evolution experiment, the diploid strain was streaked from frozen stock onto YPD agar plates. A single colony was inoculated into glycerol–lactate medium (3 mL 100% glycerol, 2 mL 40% lactate, 50 mg complete amino acid mixture, 660 mg yeast nitrogen base per 100 mL) and cultured for 48 h.

From this starter culture, 1% (v/v) inoculum was transferred daily into fresh medium to establish six independent replicate populations in each of two carbon environments:

- Glucose (0.5% w/v) supplemented with 50 mg complete amino acid mixture and 0.66 mg yeast nitrogen base
- Galactose (0.5% w/v) supplemented with the same additives

All populations were propagated by serial 1:100 dilutions every 24 h and maintained at 30°C, 250 rpm shaking conditions for a total of ~1200 generations.

### Growth Rate Measurements as a Proxy for Adaptation

Growth rates were quantified at four evolutionary time points: 300, 600, 900, and 1200 generations. At each point, populations were revived from frozen stocks and inoculated into the same carbon source in which they had been evolved. Optical density (OD) measurements were then recorded to estimate growth rate, which served as a proxy for fitness.

We denote these assays as follows:

- ggg: populations evolved in glucose, passaged in glucose, and assayed in glucose.
- GGG: populations evolved in galactose, passaged in galactose, and assayed in galactose.
- gGg: populations evolved in glucose, passaged in galactose, and assayed in glucose.
- GgG: populations evolved in galactose, passaged in glucose, and assayed in galactose.

This design ensured that the measured growth rates reflected adaptation to the *native selection environment* of each evolving population.

### Reciprocal Environment Assays

To assess the stability and plasticity of adaptation, evolved populations were exposed to the non-home carbon source for a short duration (~15 generations) and subsequently returned to their original environment. For glucose-evolved populations, the non-home environment was 0.5% galactose medium, while for galactose-evolved populations, the non-home environment was 0.5% glucose medium.

Fitness was assayed in the home environment at three distinct stages:

- Immediately after growth in the non-home environment
- Following ~7 generations of regrowth in the home environment
- Following ~15 generations of regrowth in the home environment

At each assay point, 100–200 µL of culture (volume adjusted according to OD) was inoculated into 10 mL of the relevant assay medium. Cultures were incubated at 30 °C with shaking at 250 rpm, and optical density (OD) measurements were collected at regular intervals. Growth dynamics were quantified by calculating a composite growth rate, incorporating both the lag phase duration and the exponential growth rate derived from the OD growth curves.

Each kinetic assay was performed in triplicate for every population at every time point. In parallel, ancestral strains were revived and assayed under identical conditions, providing a consistent baseline for estimating relative fitness gains during evolution.

### Whole-Genome Sequencing

Evolved populations at four evolutionary time points (300, 600, 900, and 1200 generations) were revived in 5 mL of their respective evolution environments. Cells were harvested, and genomic DNA was extracted using the phenol–chloroform method (56). Sequencing libraries were prepared, and 150 bp paired-end sequencing was performed on the DNBSEQ platform at ~100x coverage by Haystack Analytics. The ancestral diploid SK1 strain was also sequenced at the same depth for comparison.

### Variant Calling and Filtering

Complete assembled SK1 sequence was used as reference genome for alignment of all evolved populations (57). Liftoff was used to extract gene annotations from S288C and annotate the reference SK1 genome (58).

For each evolved population, sequencing reads were aligned to the SK1 reference genome using BWA-MEM. Duplicate reads were identified and marked with Picard Tools. Variant calling was performed with GATK4, following the recommended best practices pipeline: HaplotypeCaller, GenomicsDBImport, and GenotypeGVCFs. Variants were jointly called across all time points for each replicate line to maintain consistency in variant representation. Variant annotation was carried out using SnpEff (59).

From the resulting VCF files, total read depth and allele-specific read depths (reference and alternate alleles) were extracted for each sample. Variants were filtered out based on the following conditions:

- the average coverage across all time points was <30x; and
- all populations across the four time points had <20x coverage; and
- variant calls with more than two alleles; and
- the allele frequency remained unchanged across time.

Mutations were considered fixed when the alternate allele frequency reached thresholds of:

- 0.4–0.6 for heterozygous alternate alleles,
- 0.8–1.0 for homozygous alternate alleles, or
- 0–0.2 for homozygous reference alleles, relative to the reference genome.

All variants were visually validated using the Integrated Genome Viewer (IGV).

Mitochondrial genome variant analysis was carried out independently following the same pipeline using SK1 mitochondrial genome as reference.

The raw sequencing data of all evolved populations is available at http://www.ncbi.nlm.nih.gov/bioproject/1298216.

### Copy Number Variation (CNV) Analysis

To ensure comparable sequencing coverage across samples, the median coverage across the genome was calculated for each evolved replicate at every time point and in both evolution environments **(Supplementary Figure S8)**.

A custom coverage-based pipeline was used to identify copy number variants (CNVs) in all evolved populations. First, per-site read depth across the genome was calculated using samtools depth. To detect contiguous genomic regions exhibiting CNVs, the genome was divided into 500 bp windows, and the median coverage per window was determined. Each window’s coverage was then normalized by the genome-wide median coverage for that sample. To account for standing variation present in the ancestor, the ratio of normalized coverage in evolved populations to that of the ancestral reference was computed for each window, yielding a measure of relative coverage. All the windows that had a normalized coverage less than 0.25 in ancestor were excluded from further analysis.

To classify CNV states, we implemented an unsupervised Hidden Markov Model (HMM) allowing discrete states of 0, 0.5, 1, 1.5, 2, 3, and 4. A CNV state of 1 indicated equal copy number in evolved and ancestral genomes, values <1 indicated deletions, and values >1 indicated duplications in the evolved populations (52).

### CNV Annotation and Pathway Enrichment Analysis

Using the CNV states obtained from coverage-based analysis, all windows with a state of 1 (no change relative to ancestor) were excluded. The remaining windows were categorized as duplications (>1) or deletions (<1). Each set of windows was annotated to identify the genes contained within the windows for every replicate at each time point. Genes of unknown function were filtered out, and annotations across time points were merged within each replicate to obtain the total number of unique genes affected by CNVs. To visualize genomic variation, the average CNV state per window was calculated across replicates and plotted for all time points in both environments.

The ends of chromosomes are prone to coverage variation in *S. cerevisiae*. Based on the mean telomeric length (60) across isolates of *S. cerevisiae*, 500bp was eliminated from both the ends of each chromosome and the analysis was repeated. Average CNV state across genome was plotted for each environment.

For Pathway Enrichment Analysis (PEA), only CNVs shared across all replicates were considered. A t-test was performed for each genomic window based on the relative coverage ratio across replicates. Windows with p < 0.05 and a CNV state ≠1 in at least four replicates were identified as high-confidence CNVs. These were further separated into duplications and deletions, annotated, and filtered to exclude genes of unknown function. For each time point and environment, a nonredundant list of unique genes showing CNVs across replicates was compiled.

Because the resulting gene lists were large (300–1200 genes), they were randomly partitioned into bins of 100 genes, and KEGG pathway enrichment analysis was performed on each bin using the clusterProfiler package in R (23). This random segregation and enrichment analysis was repeated 3–12 times, depending on gene list size. The adjusted p-values from all runs were combined using Fisher’s method, and pathways were ranked by gene count and adjusted p-value to identify the most significantly affected pathways. This analysis was performed independently for each time point in both evolution environments.

## Supporting information

Supplement Figures, Tables, and Codes.

## Acknowledgements

This work was supported by the DBT/Wellcome Trust India Alliance grant IA/S/19/2/504632 awarded to SS.

## Supplement Material

Supplement Figures

Supplement Tables.

Codes for CNV analysis, SNP analysis and linear model of adaptation.

## Data Availability Statement

All data supporting the findings of this study are provided in the Supplementary Information files. Raw sequencing data have been deposited with NCBI and is available at: http://www.ncbi.nlm.nih.gov/bioproject/1298216. All analysis scripts and custom codes used in this study are included in the Supplementary Materials.

## Conflict of interest statement

All authors declare that they have no conflict of interest to report.

## Notes

### Competing Interest Statement

The authors have declared no competing interest.

### Summary of Updates

- Extended genomic analysis. - Rewritten sections of the manuscript.

