## Supplement Figures, Tables, and Codes. for "Opposing copy number variation dynamics accompany adaptation to glucose and galactose in diploid yeast"

Running Title: *CNVs and adaptation in glucose, galactose*

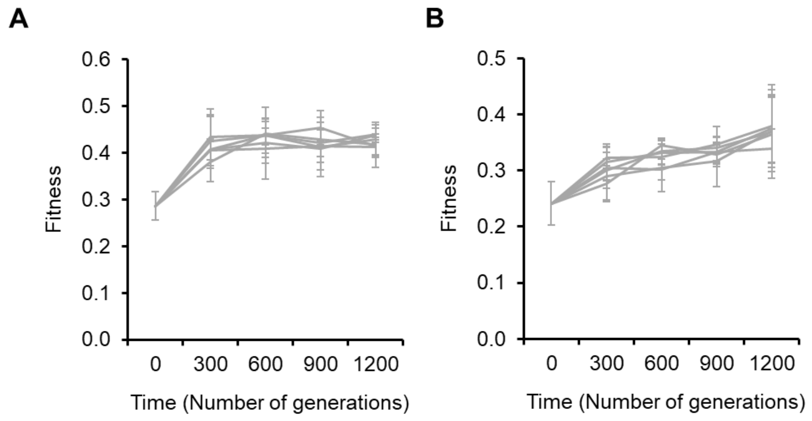


**Figure S1. Adaptation of yeast populations over 1,200 generations in glucose and galactose.** **(A)** Fitness trajectories of six independent yeast lines evolved in glucose. **(B)** Fitness trajectories of six independent yeast lines evolved in galactose. Fitness was measured every 300 generations. Data points represent mean values from three replicate experiments, with error bars indicating standard deviation.

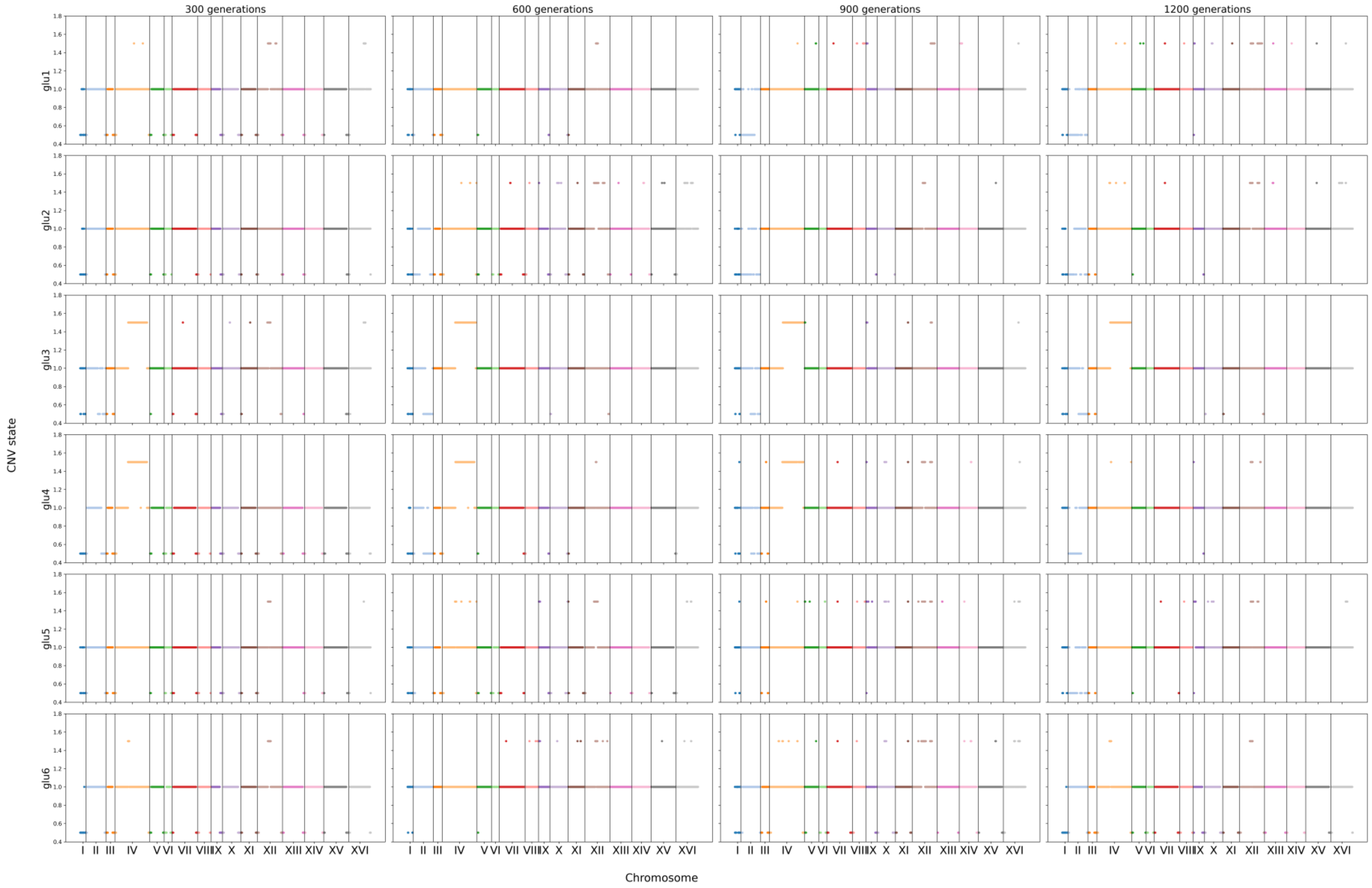
300 generations
600 generations
900 generations
1200 generations
glu1
glu2
glu3
glu4
glu5
glu6
CNV state
0.4 0.6 0.8 1.0 1.2 1.4 1.6 1.8
I II III IV V VI VII VIII IX X XI XII XIII XIV XV XVI
Chromosome

**Figure S2. Replicate-wise CNV dynamics in glucose-evolved populations across 1,200 generations.** CNV states in each of the six glucose-evolved lines (rows) at 300, 600, 900, and 1,200 generations (columns). The *x*-axis lists the 16 *S. cerevisiae* chromosomes and y-axis is CNV state. Each point is 500bp window. The CNV states are either <1 (deletion), 1 (no change) or >1 (duplication). All six replicates show similar CNV dynamics – early onset of telomeric deletions followed by compensatory duplications in the same regions. Only chromosome I and II exhibit deletions throughout the chromosome. Duplications are sporadic but persistent. Replicate lines 3 and 4 however show a significant duplication of the right arm of chromosome IV which is not seen in the rest of replicate lines.

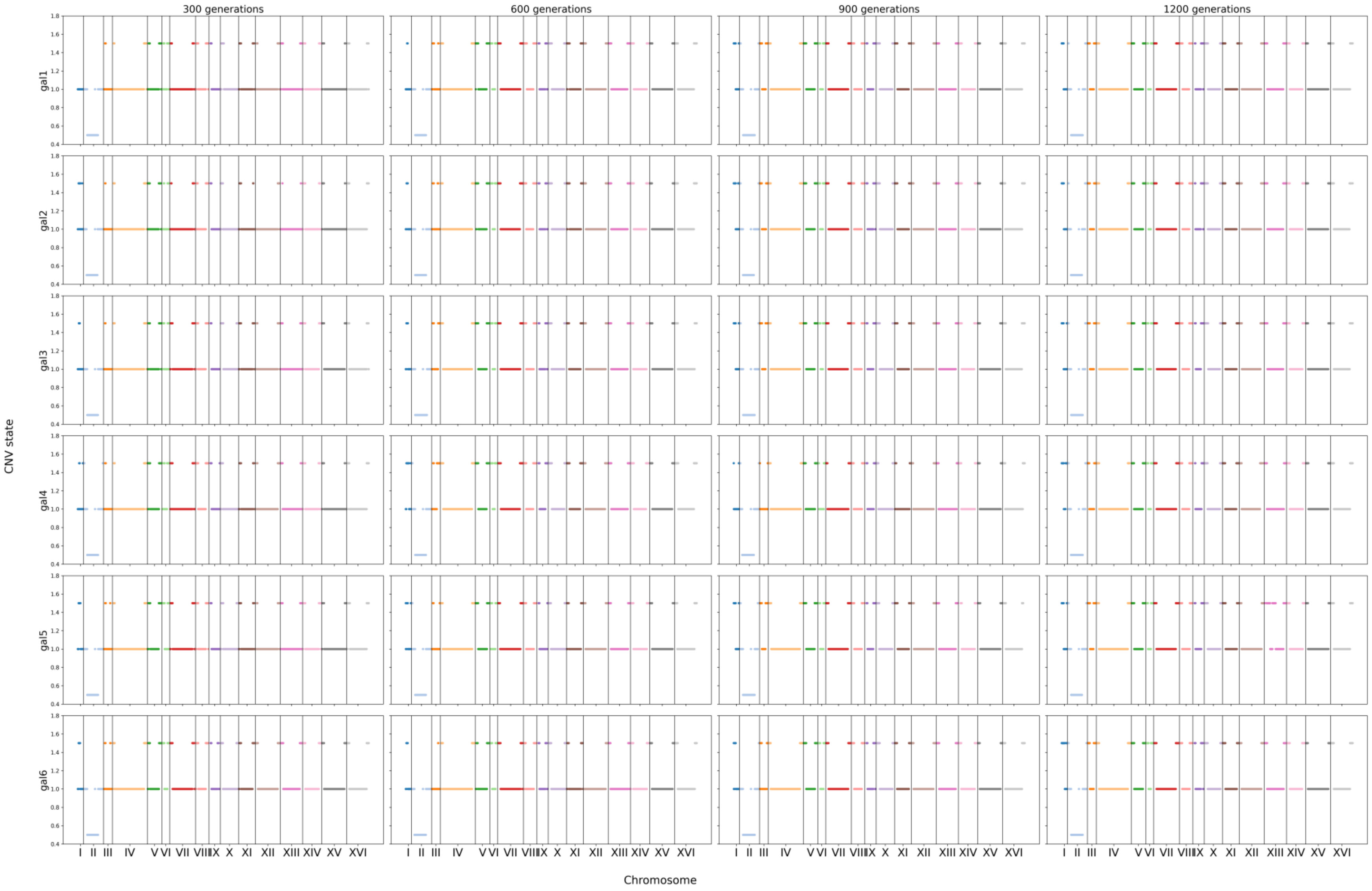
300 generations
600 generations
900 generations
1200 generations
CNV state
gal1
gal2
gal3
gal4
gal5
gal6
1.8
1.6
1.4
1.2
1.0
0.8
0.6
0.4
I II III IV V VI VII VIII IX X XI XII XIII XIV XV XVI
Chromosome

**Figure S3. Replicate-wise CNV dynamics in galactose-evolved populations across 1,200 generations.** CNV states in each of the six galactose-evolved lines (rows) at 300, 600, 900, and 1,200 generations (columns). The *x*-axis lists the 16 *S. cerevisiae* chromosomes while Y-axis represent the CNV state. Each coloured dot is a 500bp window. A CNV state of <1 is deletion, 1 is no change with respect to ancestor, >1 is duplication. All the replicate lines exhibit a similar trend of persistent duplications which increase in number with time. Deletions are observed only on chromosome II.

**(a)**

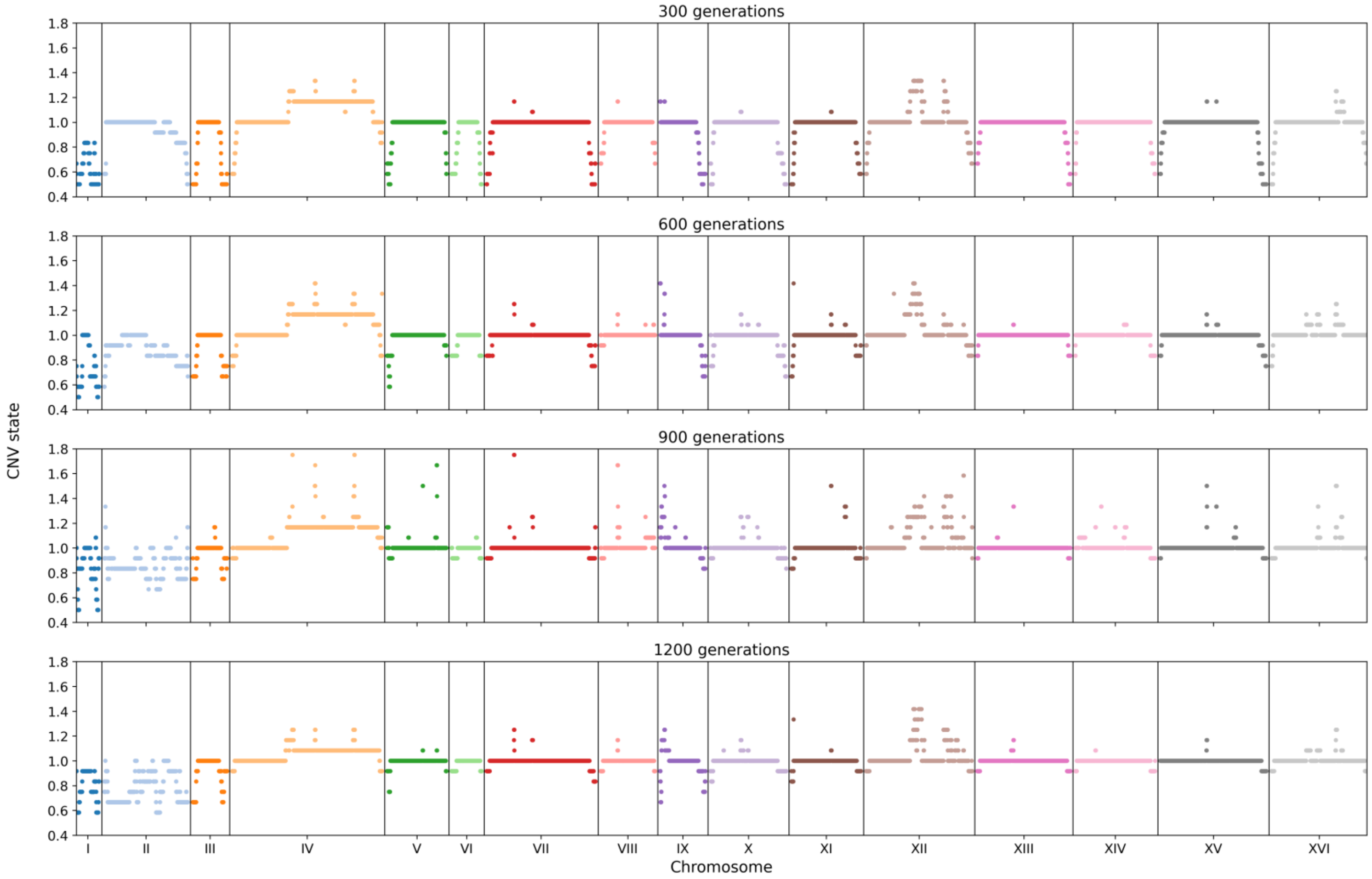

300 generations
600 generations
900 generations
1200 generations
CNV state
1.8
1.6
1.4
1.2
1.0
0.8
0.6
0.4
I
II
III
IV
V
VI
VII
VIII
IX
X
XI
XII
XIII
XIV
XV
XVI
Chromosome

**(b)**

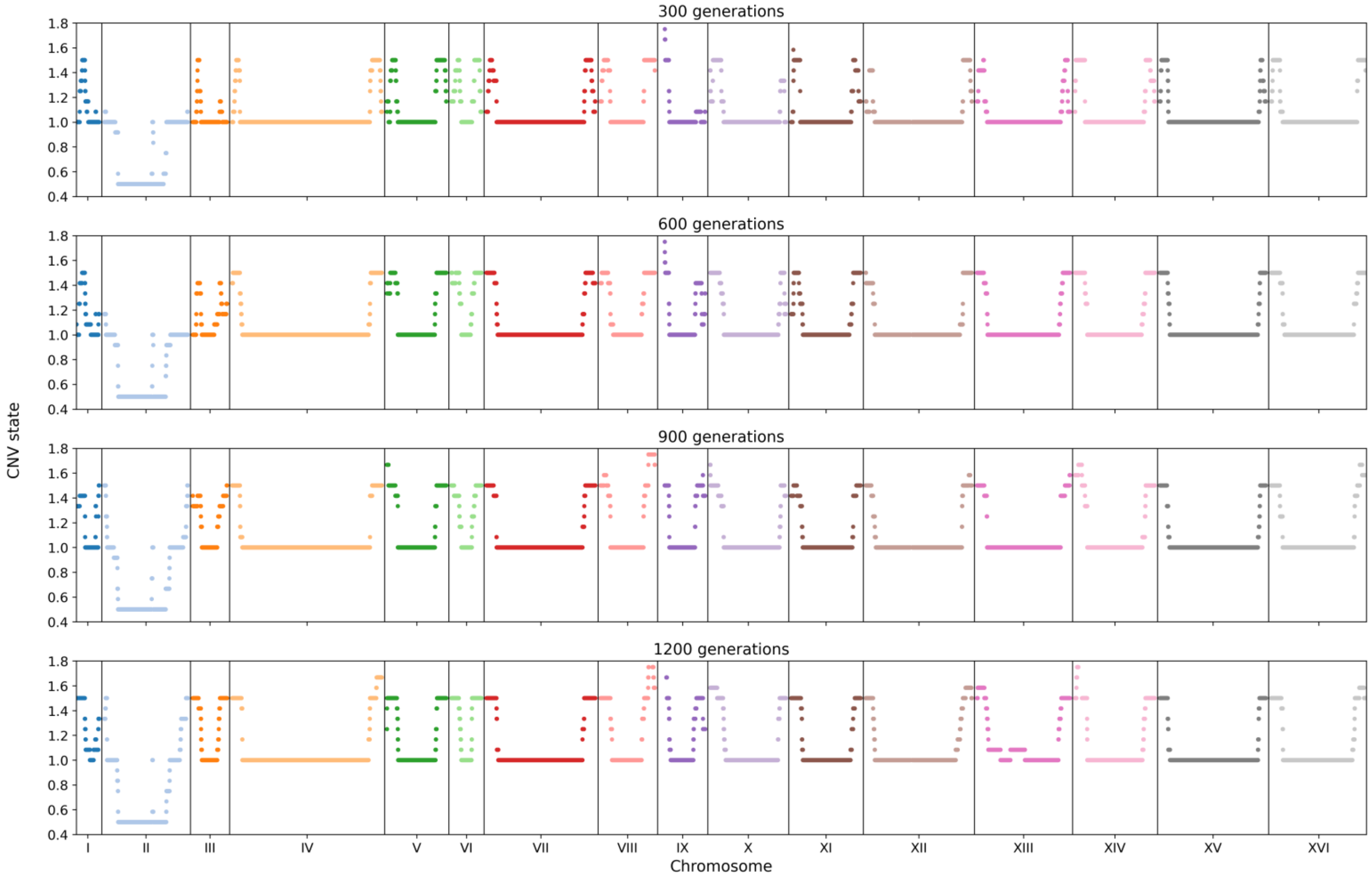


**Figure S4**. **Average genome-wide CNV state plots across 1200 generations in (a) glucose- and (b) galactose-evolved populations.** 500bp from both the ends of each chromosome was eliminated and average CNV state across the genome was plotted to see if the variations extended beyond telomeric regions. The opposing trend of deletions in glucose and duplications in galactose remain even after elimination of telomeric regions.

**(a)**

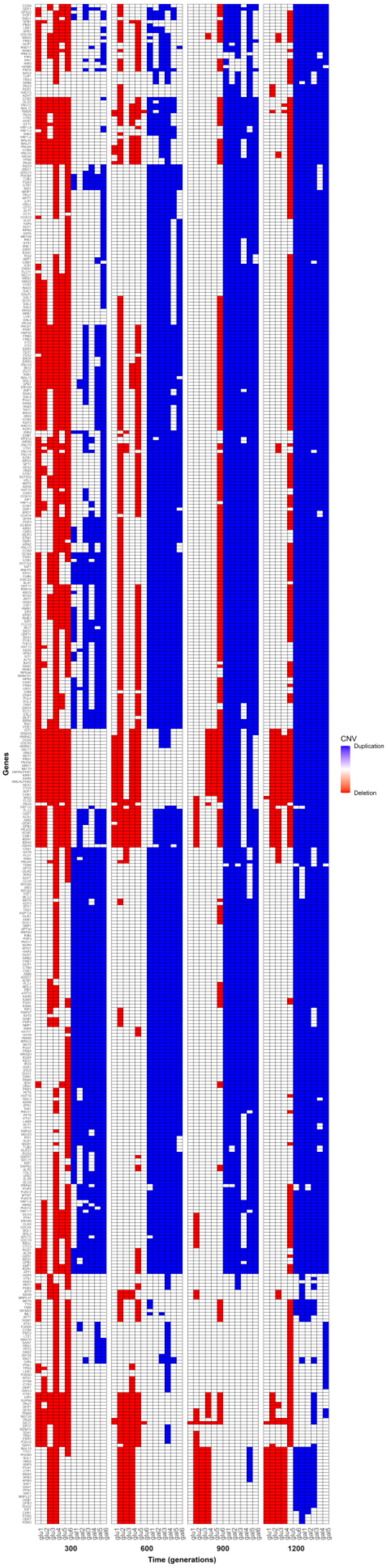
CNV
Duplication
Deletion
Genes
300
600
900
1200
Time (generations)

**(b)**

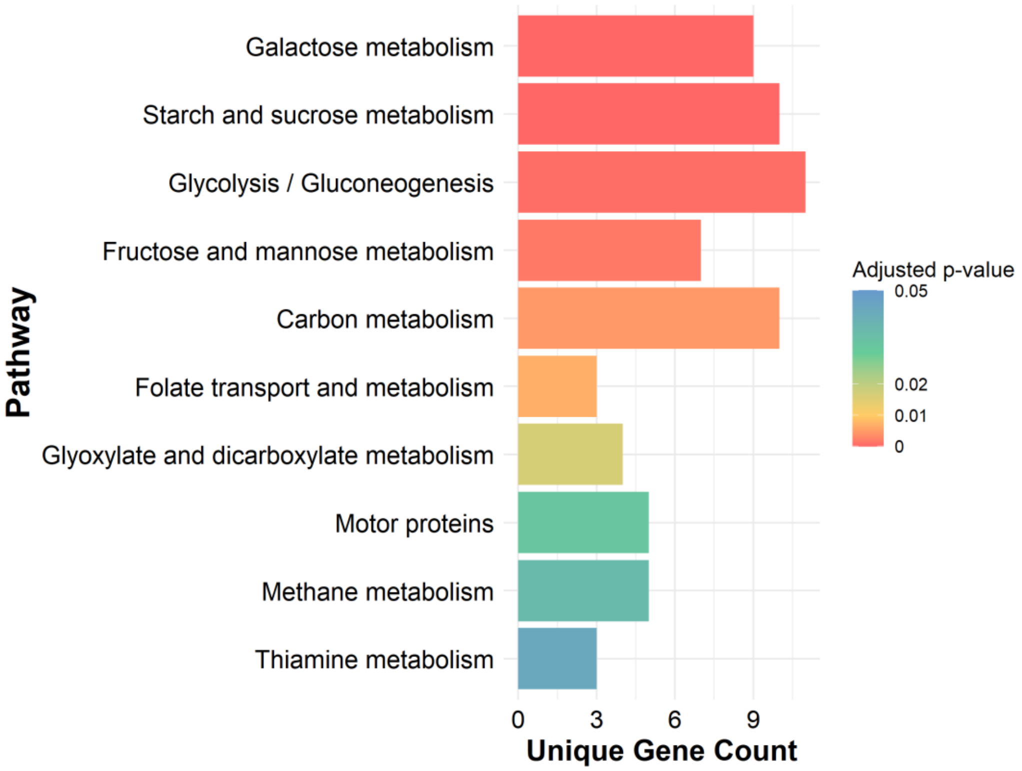


**Figure S5**. **Replicate-wise antagonistic copy number evolution in glucose versus galactose. (a)** The pattern of opposing copy number variation is observed even at a replicate level. Each row is a gene, and column is a replicate line evolved in either glucose or galactose. Each block represents a single evolutionary time point. Deletions are represented by red, duplications by blue, and no change with respect to ancestor by white. With increasing period of evolution, the deletions in glucose decrease (disappearance of red boxes) while duplications in galactose increase (appearance of blue). **(b)** KEGG pathway enrichment analysis (using clusterProfiler) of the genes undergoing opposing selection in glucose and galactose in 1200 generations. The ten most significantly enriched pathways are listed on y-axis, x-axis represents the number of genes, and the colour gradient is based on the adjusted p-value.

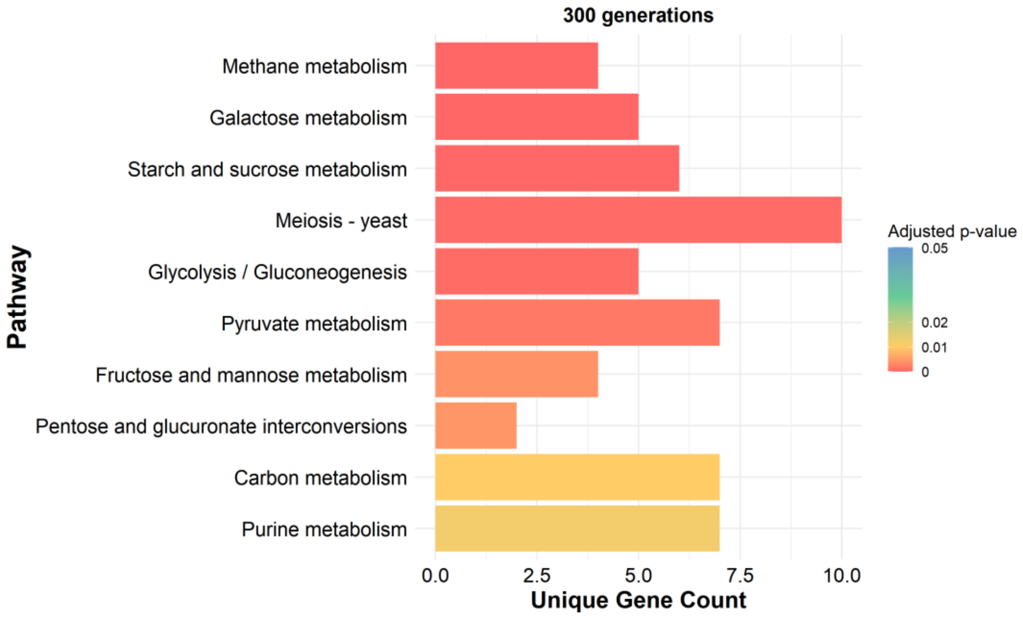


**Figure S6. KEGG Pathway Enrichment Analysis of genes that have undergone deletion in glucose.** The genes that were commonly deleted in all the replicate lines evolved in glucose were subjected to KEGG pathway enrichment analysis using the cluster Profiler package in R (see Methods). X-axis represents the number of genes belonging to each of the pathway and the pathways are ordered based on significance (represented by the colour gradient). As the deletion burden decreased with evolutionary time, no significant enrichment for pathways was observed beyond 300 generations. Also, genes that have undergone duplications do not show enrichment for any specific pathway.

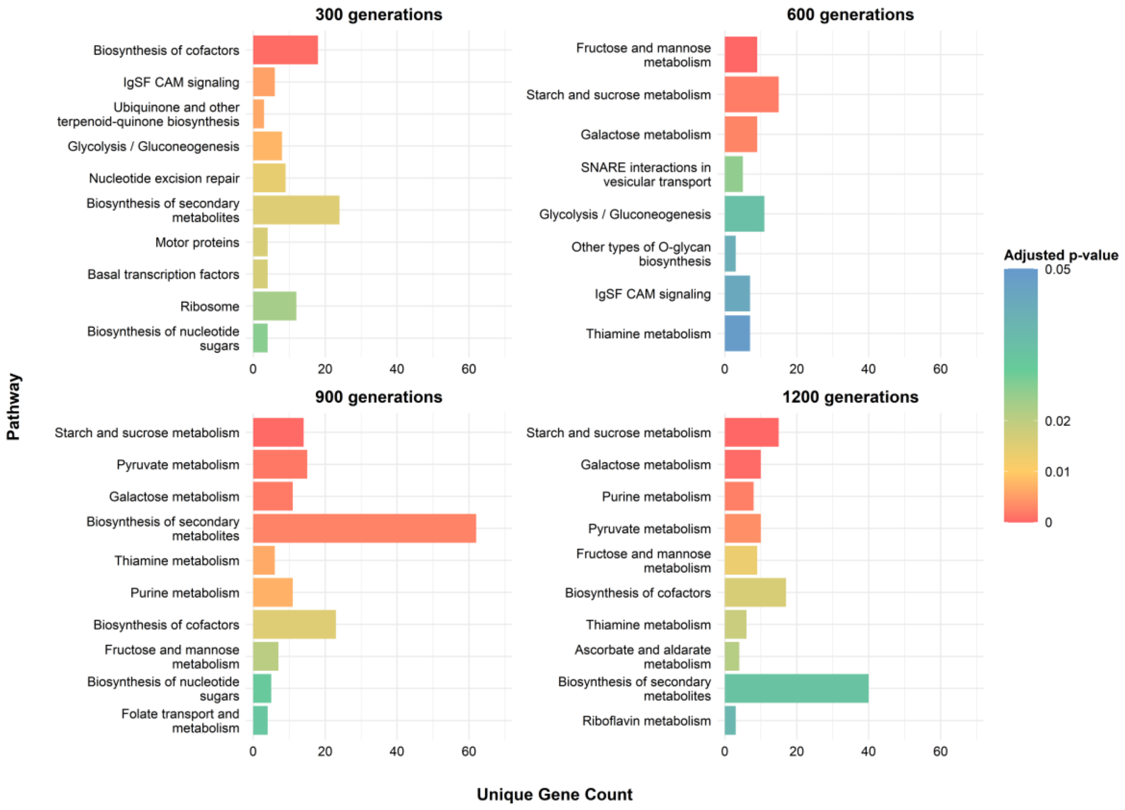


**Figure S7. KEGG Pathway Enrichment Analysis of genes that have undergone duplication in galactose.** The genes that were commonly duplicated in all the replicate lines evolved in galactose were subjected to KEGG pathway enrichment analysis using the cluster Profiler package in R (see Methods). X-axis represents the number of genes belonging to each of the pathway and the pathways are ordered based on the significance (represented by the colour gradient). The number of genes as well as pathways enriched due to duplications increase with time. Only ten most significant pathways are listed here. The genes that have undergone deletion in galactose do not show enrichment in any specific pathway.

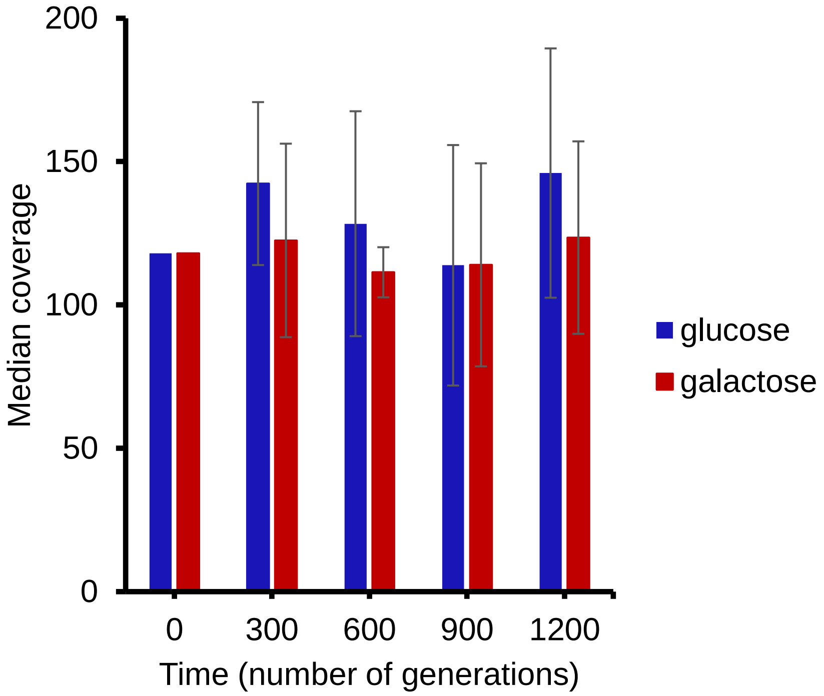


**Figure S8. Average of the median of genome coverage of all populations sequenced.** Coverage at each position and hence the median of the coverage across genome was calculated for all the evolved populations and ancestor. Each bar represents the mean of six glucose- or galactose-evolved replicates sequenced at the given time point. The error bar is the standard deviation. Only exception is gal5 line at 600 generations. The median coverage of this line is 393 and is excluded from this plot. Time point 0 on y-axis corresponds to the coverage of ancestor which is common to both glucose- and galactose-evolved populations.

**Table S1:** List of SNP/Indel along with the allele frequency at each time point sequenced of each glucose and galactose replicate.

**Glucose-evolved lines:**

**glu1**

| CHROM | POS | REF | ALT | Consequence | Impact | GeneName | 0gen | 300gen | 600gen | 900gen | 1200gen |
|---|---|---|---|---|---|---|---|---|---|---|---|
| chrII | 456577 | C | T | synonymous_variant | LOW | SIF2 | 0.3817 | 0.3103 | 0.3233 | 0.0000 | 0.0000 |
| chrII | 477457 | C | T | synonymous_variant | LOW | RAD16 | 0.3324 | 0.7657 | 0.7720 | 0.9580 | 1.0000 |
| chrII | 684747 | G | A | downstream_gene_variant | MODIFIER | SLX1 | 0.0000 | 0.0000 | 0.0000 | 0.0000 | 0.5000 |
| chrIV | 174965 | T | A | stop_gained | HIGH | NRP1 | 0.0000 | 1.0000 | 1.0000 | 1.0000 | 1.0000 |
| chrVIII | 149476 | A | T | synonymous_variant | LOW | MYO1 | 0.0000 | 0.0000 | 0.0000 | 0.1589 | 0.4000 |
| chrXI | 194390 | T | A | stop_gained | HIGH | APL2 | 0.0000 | 0.0000 | 0.0000 | 0.0000 | 0.4375 |
| chrXII | 173452 | G | GT | frameshift_variant | HIGH | PPR1 | 0.0000 | 0.0000 | 0.0000 | 0.0000 | 0.3871 |
| chrXII | 280475 | C | T | downstream_gene_variant | MODIFIER | FMP25 | 0.5143 | 0.0000 | 0.0000 | 0.0000 | 0.0000 |
| chrXII | 563956 | T | G | upstream_gene_variant | MODIFIER | YNCL0032C | 0.0000 | 0.0000 | 0.1455 | 0.5208 | 0.5574 |
| chrXII | 563961 | A | AC | upstream_gene_variant | MODIFIER | YNCL0032C | 0.0000 | 0.0000 | 0.1481 | 0.5349 | 0.5517 |
| chrXIII | 561137 | T | A | stop_gained | HIGH | MSS11 | 0.0000 | 0.0000 | 0.1405 | 0.9437 | 1.0000 |
| chrXIV | 122233 | A | G | upstream_gene_variant | MODIFIER | BOR1 | 0.0000 | 0.0000 | 0.2048 | 0.1978 | 0.6056 |
| chrXV | 831184 | C | CT | upstream_gene_variant | MODIFIER | SNF2 | 0.5397 | 0.0000 | 0.0000 | 0.0000 | 0.0000 |
| chrXVI | 148816 | T | C | missense_variant | MODERATE | THI6 | 0.0000 | 0.0000 | 0.0000 | 0.0000 | 0.3430 |

**glu2**

| CHROM | POS | REF | ALT | Consequence | Impact | GeneName | 0gen | 300gen | 600gen | 900gen | 1200gen |
|---|---|---|---|---|---|---|---|---|---|---|---|
| chrIV | 50225 | AG | A | frameshift_variant | HIGH | MFG1 | 0.0000 | 0.0000 | 0.9529 | 1.0000 | 1.0000 |
| chrIV | 174965 | T | A | stop_gained | HIGH | NRP1 | 0.0000 | 0.8242 | 1.0000 | 1.0000 | 1.0000 |
| chrXII | 280475 | C | T | downstream_gene_variant | MODIFIER | FMP25 | 0.5143 | 0.1546 | 0.0000 | 0.0000 | 0.0000 |
| chrXII | 277333 | C | A | upstream_gene_variant | MODIFIER | FMP25 | 0.5179 | 0.0000 | 0.0000 | 0.0000 | 0.0000 |
| chrXV | 831184 | C | CT | upstream_gene_variant | MODIFIER | SNF2 | 0.5397 | 0.0964 | 0.0000 | 0.0000 | 0.0000 |
| chrXV | 300039 | C | A | stop_gained | HIGH | SIN3 | 0.0000 | 0.0000 | 0.0000 | 0.3182 | 0.3400 |
| chrVII | 566793 | A | C | upstream_gene_variant | MODIFIER | YNCG0025C | 0.4128 | 0.1296 | 0.1395 | 0.0000 | 0.0000 |
| chrXIV | 122233 | A | G | upstream_gene_variant | MODIFIER | BOR1 | 0.0000 | 0.0000 | 0.4091 | 0.3654 | 0.4048 |

**glu3**

| CHROM | POS | REF | ALT | Consequence | Impact | GeneName | 0gen | 300gen | 600gen | 900gen | 1200gen |
|---|---|---|---|---|---|---|---|---|---|---|---|

| | | | | | | | | | | | |
|---|---|---|---|---|---|---|---|---|---|---|---|
| chrV | 396704 | GT | G | downstream_gene_variant | MODIFIER | KAP123 | 0.0000 | 0.0000 | 0.3333 | 0.6000 | 0.6789 |
| chrMT | 81543 | TA | T | intron_variant | MODIFIER | COX1 | 0.0000 | 1.0000 | 1.0000 | 1.0000 | 1.0000 |
| chrXII | 280475 | C | T | downstream_gene_variant | MODIFIER | FMP25 | 0.5143 | 0.0000 | 0.0000 | 0.0000 | 0.0000 |
| chrXV | 391585 | G | A | upstream_gene_variant | MODIFIER | WHI2 | 0.0000 | 0.0000 | 0.0000 | 0.1463 | 0.2276 |
| chrXIII | 585172 | T | A | synonymous_variant | LOW | ECM5 | 0.0000 | 0.0000 | 0.0000 | 0.3529 | 0.2579 |

**glu4**

| CHROM | POS | REF | ALT | Consequence | Impact | GeneName | 0gen | 300gen | 600gen | 900gen | 1200gen |
|---|---|---|---|---|---|---|---|---|---|---|---|
| chrI | 194352 | T | C | missense_variant | MODERATE | YAT1 | 0.0000 | 0.0000 | 0.0000 | 0.0000 | 0.5325 |
| chrIV | 50225 | AG | A | frameshift_variant | HIGH | MFG1 | 0.0000 | 0.0000 | 0.0000 | 0.0000 | 0.9904 |
| chrIV | 174965 | T | A | stop_gained | HIGH | NRP1 | 0.0000 | 0.0000 | 0.0000 | 0.0000 | 1.0000 |
| chrXII | 280475 | C | T | downstream_gene_variant | MODIFIER | FMP25 | 0.5143 | 0.4757 | 0.5436 | 0.4588 | 0.0000 |
| chrXV | 831184 | C | CT | upstream_gene_variant | MODIFIER | SNF2 | 0.5397 | 0.5060 | 0.5283 | 0.5429 | 0.0000 |
| chrVII | 642759 | G | GT | downstream_gene_variant | MODIFIER | PIL1 | 0.0000 | 0.0000 | 0.0000 | 0.0930 | 0.5238 |
| chrXVI | 831790 | A | G | downstream_gene_variant | MODIFIER | SUE1 | 0.0000 | 0.0000 | 0.0000 | 0.0000 | 0.4783 |
| chrXV | 300039 | C | A | stop_gained | HIGH | SIN3 | 0.0000 | 0.0000 | 0.0000 | 0.0000 | 0.3696 |

**glu5**

| CHROM | POS | REF | ALT | Consequence | Impact | GeneName | 0gen | 300gen | 600gen | 900gen | 1200gen |
|---|---|---|---|---|---|---|---|---|---|---|---|
| chrII | 468840 | G | T | downstream_gene_variant | MODIFIER | YSA1 | 0.0000 | 0.2849 | 0.2941 | 0.4271 | 0.4868 |
| chrXIII | 562220 | G | A | stop_gained | HIGH | MSS11 | 0.0000 | 0.0000 | 0.0000 | 0.0000 | 0.8925 |
| chrXIV | 444577 | GT | G | frameshift_variant | HIGH | PHO23 | 0.0000 | 0.8476 | 0.9732 | 1.0000 | 1.0000 |
| chrXII | 1028098 | G | A | missense_variant | MODERATE | CSS1 | 0.3750 | 0.4338 | 0.5058 | 0.0000 | 0.0000 |
| chrXII | 967165 | G | C | upstream_gene_variant | MODIFIER | CAR2 | 0.0000 | 0.0000 | 0.0000 | 0.0000 | 0.2067 |
| chrXV | 16996 | C | G | upstream_gene_variant | MODIFIER | ZPS1 | 0.3693 | 0.4472 | 0.4699 | 0.0000 | 0.0000 |

**glu6**

| CHROM | POS | REF | ALT | Consequence | Impact | GeneName | 0gen | 300gen | 600gen | 900gen | 1200gen |
|---|---|---|---|---|---|---|---|---|---|---|---|
| chrII | 468840 | G | T | downstream_gene_variant | MODIFIER | YSA1 | 0.0000 | 0.0000 | 0.4053 | 0.3168 | 0.3298 |
| chrIII | 164960 | TGTG | T | conservative_inframe_deletion | MODERATE | NPP1 | 0.0000 | 0.0000 | 0.0000 | 0.4350 | 0.4811 |
| chrV | 285041 | G | C | missense_variant | MODERATE | HIS1 | 0.0000 | 0.0000 | 0.0000 | 0.4417 | 0.4903 |
| chrXIV | 444577 | GT | G | frameshift_variant | HIGH | PHO23 | 0.0000 | 0.0000 | 0.9630 | 0.9862 | 0.9935 |
| chrII | 468840 | G | T | downstream_gene_variant | MODIFIER | ALG1 | 0.0000 | 0.0000 | 0.4053 | 0.3168 | 0.3298 |

**Galactose-evolved lines:**

**gal1**

| CHROM | POS | REF | ALT | Consequence | Impact | GeneName | 0gen | 300gen | 600gen | 900gen | 1200gen |
|---|---|---|---|---|---|---|---|---|---|---|---|
| chrVII | 380205 | C | G | missense_variant | MODERATE | SGF73 | 0.5174 | 0.5000 | 0.0000 | 0.0000 | 0.0000 |
| chrVII | 500460 | C | A | missense_variant | MODERATE | ERG26 | 0.5195 | 1.0000 | 1.0000 | 1.0000 | 1.0000 |
| chrX | 84263 | CT | C | downstream_gene_variant | MODIFIER | RPS14B | 0.4336 | 0.3939 | 0.0000 | 0.0000 | 0.0000 |
| chrXIII | 561824 | T | TTC | frameshift_variant | HIGH | MSS11 | 0.0000 | 0.0000 | 1.0000 | 1.0000 | 1.0000 |
| chrXV | 120611 | A | C | synonymous_variant | LOW | HMI1 | 0.0000 | 0.0000 | 0.0000 | 0.0000 | 0.5000 |
| chrXV | 354354 | G | T | stop_gained | HIGH | SFM1 | 0.0000 | 0.0000 | 0.4271 | 0.3701 | 0.4336 |
| chrXI | 689230 | T | C | missense_variant | MODERATE | YEL077C | 0.0000 | 0.1741 | 0.0000 | 0.0000 | 0.2203 |
| chrXVI | 232025 | C | T | synonymous_variant | LOW | MRX4 | 0.0000 | 0.0000 | 0.2154 | 0.3333 | 0.2947 |

**gal2**

| CHROM | POS | REF | ALT | Consequence | Impact | GeneName | 0gen | 300gen | 600gen | 900gen | 1200gen |
|---|---|---|---|---|---|---|---|---|---|---|---|
| chrII | 535403 | A | G | missense_variant | MODERATE | BMT2 | 0.0000 | 0.5036 | 0.5140 | 0.5091 | 0.4175 |
| chrVII | 500460 | C | A | missense_variant | MODERATE | ERG26 | 0.5195 | 0.0000 | 0.0000 | 0.0000 | 0.0000 |
| chrXIII | 562283 | C | A | stop_gained | HIGH | MSS11 | 0.0000 | 0.0000 | 0.0000 | 0.2149 | 0.6593 |
| chrII | 456577 | C | T | synonymous_variant | LOW | SIF2 | 0.3817 | 0.0000 | 0.0000 | 0.0000 | 0.0000 |
| chrX | 180278 | G | C | missense_variant | MODERATE | URA2 | 0.0000 | 0.0000 | 0.0000 | 0.1154 | 0.3375 |
| chrII | 216155 | A | G | upstream_gene_variant | MODIFIER | YNCB0004W | 0.0000 | 0.0000 | 0.1935 | 0.0000 | 0.2083 |
| chrII | 216156 | C | T | upstream_gene_variant | MODIFIER | YNCB0004W | 0.0000 | 0.0000 | 0.1935 | 0.0000 | 0.2083 |

**gal3**

| CHROM | POS | REF | ALT | Consequence | Impact | GeneName | 0gen | 300gen | 600gen | 900gen | 1200gen |
|---|---|---|---|---|---|---|---|---|---|---|---|
| chrV | 396400 | G | A | downstream_gene_variant | MODIFIER | KAP123 | 0.0000 | 0.0000 | 0.0000 | 0.0000 | 0.5660 |
| chrVII | 500460 | C | A | missense_variant | MODERATE | ERG26 | 0.5195 | 0.9463 | 0.9623 | 1.0000 | 1.0000 |
| chrX | 84263 | CT | C | downstream_gene_variant | MODIFIER | RPS14B | 0.4336 | 0.9317 | 1.0000 | 1.0000 | 1.0000 |
| chrVII | 523574 | C | G | stop_gained | HIGH | MSB2 | 0.0000 | 0.0000 | 0.0000 | 0.0000 | 0.3772 |
| chrXIV | 699421 | G | T | missense_variant | MODERATE | SOL1 | 0.0000 | 0.0000 | 0.0000 | 0.0000 | 0.3498 |
| chrI | 222232 | A | G | downstream_gene_variant | MODIFIER | PAU8 | 0.0000 | 0.2187 | 0.1446 | 0.2476 | 0.2166 |
| chrIV | 6517 | A | G | upstream_gene_variant | MODIFIER | YRF1-4 | 0.0000 | 0.1458 | 0.4651 | 0.2353 | 0.3036 |
| chrXIV | 122233 | A | G | upstream_gene_variant | MODIFIER | BOR1 | 0.0000 | 0.0000 | 0.0000 | 0.5714 | 0.4200 |
| chrXV | 55042 | G | A | downstream_gene_variant | MODIFIER | MED7 | 0.0000 | 0.0000 | 0.0000 | 0.0000 | 0.3158 |

| | | | | | | | | | | | |
|---|---|---|---|---|---|---|---|---|---|---|---|
| chrVII | 523574 | C | G | stop_gained | HIGH | MSB2 | 0.0000 | 0.0000 | 0.0000 | 0.0000 | 0.3772 |

**gal4**

| CHROM | POS | REF | ALT | Consequence | Impact | GeneName | 0gen | 300gen | 600gen | 900gen | 1200gen |
|---|---|---|---|---|---|---|---|---|---|---|---|
| chrV | 396420 | CT | C | downstream_gene_variant | MODIFIER | KAP123 | 0.0000 | 0.0000 | 0.0000 | 0.6872 | 1.0000 |
| chrX | 84263 | CT | C | downstream_gene_variant | MODIFIER | RPS14B | 0.4336 | 0.5769 | 0.9931 | 1.0000 | 1.0000 |
| chrXII | 494744 | C | T | upstream_gene_variant | MODIFIER | EMG1 | 0.0000 | 0.1111 | 0.5205 | 0.7326 | 0.7416 |
| chrVII | 380205 | C | G | missense_variant | MODERATE | SGF73 | 0.5174 | 0.1028 | 0.0000 | 0.0000 | 0.0000 |
| chrIII | 202611 | A | C | upstream_gene_variant | MODIFIER | PHO87 | 0.0000 | 0.0000 | 0.0000 | 0.0000 | 1.0000 |
| chrIII | 202636 | C | T | upstream_gene_variant | MODIFIER | PHO87 | 0.0000 | 0.0000 | 0.0000 | 0.0000 | 0.6466 |
| chrII | 456577 | C | T | synonymous_variant | LOW | SIF2 | 0.3817 | 0.4603 | 0.9217 | 1.0000 | 1.0000 |
| chrX | 181359 | G | A | stop_gained | HIGH | URA2 | 0.0000 | 0.0000 | 0.9859 | 0.9907 | 1.0000 |

**gal5**

| CHROM | POS | REF | ALT | Consequence | Impact | GeneName | 0gen | 300gen | 600gen | 900gen | 1200gen |
|---|---|---|---|---|---|---|---|---|---|---|---|
| chrVII | 380205 | C | G | missense_variant | MODERATE | SGF73 | 0.5174 | 0.9496 | 0.9755 | 0.9929 | 0.9913 |
| chrVII | 500460 | C | A | missense_variant | MODERATE | ERG26 | 0.5195 | 0.8586 | 0.9769 | 1.0000 | 1.0000 |
| chrXII | 30802 | T | C | missense_variant | MODERATE | YCT1 | 0.0000 | 0.0000 | 0.0000 | 0.2011 | 0.4178 |
| chrXIII | 798607 | G | T | stop_gained | HIGH | ATR2 | 0.0000 | 0.0000 | 0.0000 | 0.5172 | 0.4950 |
| chrII | 456577 | C | T | synonymous_variant | LOW | SIF2 | 0.3817 | 0.5181 | 0.8675 | 1.0000 | 1.0000 |
| chrXIII | 440270 | C | A | missense_variant | MODERATE | MUB1 | 0.0000 | 0.0000 | 0.0000 | 0.3258 | 0.2214 |
| chrV | 396484 | G | A | downstream_gene_variant | MODIFIER | KAP123 | 0.0000 | 0.0000 | 0.0000 | 0.9290 | 0.9760 |
| chrVIII | 527986 | G | A | downstream_gene_variant | MODIFIER | TDA8 | 0.0000 | 0.0000 | 0.0000 | 0.4594 | 0.3913 |

**gal6**

| CHROM | POS | REF | ALT | Consequence | Impact | GeneName | 0gen | 300gen | 600gen | 900gen | 1200gen |
|---|---|---|---|---|---|---|---|---|---|---|---|
| chrX | 84263 | CT | C | downstream_gene_variant | MODIFIER | RPS14B | 0.4336 | 0.0000 | 0.0000 | 0.0000 | 0.0000 |
| chrXII | 280475 | C | T | downstream_gene_variant | MODIFIER | FMP25 | 0.5143 | 0.0000 | 0.0000 | 0.0000 | 0.0000 |
| chrXIII | 561419 | G | A | stop_gained | HIGH | MSS11 | 0.0000 | 0.0000 | 0.0000 | 1.0000 | 0.9737 |
| chrX | 709079 | C | A | missense_variant | MODERATE | DAN1 | 0.0000 | 0.0000 | 0.0000 | 0.3068 | 0.2509 |
| chrXII | 103452 | C | A | stop_gained | HIGH | SPA2 | 0.0000 | 0.0000 | 0.0000 | 0.2000 | 0.2222 |
| chrII | 456577 | C | T | synonymous_variant | LOW | SIF2 | 0.3817 | 0.0000 | 0.0000 | 0.0000 | 0.0000 |
| chrII | 216155 | A | G | upstream_gene_variant | MODIFIER | YNCB0004W | 0.0000 | 0.1667 | 0.0000 | 0.1579 | 0.3438 |

| | | | | | | | | | | | |
|---|---|---|---|---|---|---|---|---|---|---|---|
| chrII | 216156 | C | T | upstream_gene_variant | MODIFIER | YNCB0004W | 0.0000 | 0.1667 | 0.0000 | 0.1579 | 0.3438 |
| chrIV | 526089 | A | T | upstream_gene_variant | MODIFIER | LYS14 | 0.0000 | 0.0000 | 0.0000 | 0.0000 | 0.2121 |

**Table S2:** List of genes that have undergone deletion at each time point in each of the glucose and galactose replicate.

**300 generations**

| Replicate | Gene list |
|---|---|
| **glu1** | AAD10, AAD4, ACS1, ADE1, ADF1, ADH4, AGP3, AIF1, AIM2, ALD4, AMF1, APA1, AQY3, ATF1, AVT2, BAS1, BAT2, BDH1, BDH2, BGL2, BIO2, BRR6, BSC5, BUD14, CAB4, CAN1, CDC15, CDC39, CDC50, CHA1, CIN8, CNE1, COS1, COS10, COS12, COS4, COS5, COS6, COS7, COX14, CSS3, CWC22, DAK2, DAL1, DAL2, DAL3, DAL4, DAL5, DAL7, DAL81, DAN1, DAN4, DCG1, DDI2, DFP1, DFP2, DLD3, DSE4, DSF1, ECM1, EFB1, EFG1, EMA35, EMC4, ERO1, ERP1, ERP2, ERR2, ERR3, ERV29, ESL2, FDC1, FEX1, FIT2, FIT3, FLC2, FLO10, FLO5, FRE2, FRE3, FRE4, FRE5, FUN14, FYV5, FZF1, GDH1, GDH3, GEM1, GEX1, GEX2, GIT1, GPB1, GPB2, GTT1, HFM1, HIR3, HMLALPHA2, HMRA1, HMRA2, HMS2, HOM6, HPA3, HSP32, HXK2, HXT13, HXT15, HXT17, HYR1, ICR1, IMA1, IMA3, IMD2, IML1, INA22, IPA1, IRC24, IRC4, IRC7, JEN1, KAR4, KIN3, KIN82, KRR1, KTD1, LDS1, LRE1, LRG1, LYS1, MAL12, MAL13, MAL31, MAL32, MATALPHA1, MCH2, MCM22, MDM10, MET5, MGA2, MGM101, MIC10, MLP1, MND2, MNS1, MNT2, MPH3, MRC1, MRS1, MRS6, MSH3, MST28, NDD1, NFT1, NMD5, NPR2, NRE1, NUD1, NUP60, OAF1, OCA4, PAD1, PAU10, PAU12, PAU14, PAU15, PAU18, PAU19, PAU21, PAU24, PAU4, PAU6, PAU7, PAU8, PBN1, PCC1, PCK1, PDE1, PDE2, PDR18, PEX22, PEX34, PGU1, PHR1, PIP2, PMT4, POF1, PRD1, PRE10, PRM9, PRT1, PSK1, PUL3, PUL4, PWR1, PXR1, RAD17, RAI1, RDR1, RDT1, RFA1, RMD6, RME3, RMR1, RNH70, RPS12, RPS4A, RTG2, RTT102, SCP1, SCW4, SEC11, SEN34, SEO1, SGM1, SIR1, SIT1, SKG1, SLH1, SNO2, SNO4, SNR18, SNZ2, SOR1, SPB1, SPO7, SPS22, SRY1, SSA1, SUP56, SUT532, SWD1, SWH1, TAF1, TAN1, TFC3, TGA1, THI11, THI13, TIM8, TPD3, TRN1, TTI2, UBP11, UIP3, UPA1, URA1, VAC17, VBA3, VBA5, VEL1, VPS8, VTH1, VTH2, XPT1, YAT1, YOR1, YPS6, YRF1-2, YRF1-4, YRF1-6, YRF1-8, YVH1, ZIP2, ZNF1, ZRT1, ZUO1 |
| **glu2** | AAD10, ACS1, ADE1, ADF1, AGP3, AIM2, ALD4, AMF1, AQY3, ATF1, BAT2, BDH1, BDH2, BGL2, BIO2, BOL1, BOL3, BUD14, CDC15, CDC19, CDC24, CDC50, CHA1, CLN3, CNE1, COS1, COS4, COS5, COS6, COS7, CYC3, CYS3, DAK2, DAL1, DAL2, DAL3, DAL4, DAL5, DAL7, DAL81, DAN1, DAN4, DCG1, DDI2, DEP1, DFP2, DLD3, DSF1, ECM1, EFB1, EMA35, ERP1, ERP2, ERR2, ERR3, ERV29, ERV46, FEX1, FEX2, FIT2, FIT3, FLC2, FLO5, FRE3, FRE5, FUN14, FYV5, GCV3, GDH1, GDH3, GEM1, GEX1, GEX2, GIT1, GPB1, GPB2, GTT1, HMLALPHA2, HMRA1, HMRA2, HSP32, HXT13, HYR1, IMA1, IMA3, IMD2, INA22, IRC24, KAR4, KIN3, KRR1, KTD1, LYS1, MAL12, MAL13, MAL31, MAL32, MAN2, MATALPHA1, MCH2, MDM10, MGA2, MIC10, MND2, MRC1, MST28, NDD1, NRE1, NTG1, NUD1, NUP60, OAF1, OCA4, PAU10, PAU12, PAU15, PAU18, PAU19, PAU21, PAU24, PAU4, PAU6, PAU8, PEX22, PEX34, PGU1, PHO12, PHR1, PRD1, PRM9, PTA1, RBG1, RDR1, RDT1, RFA1, RMD6, SEN34, SEO1, SNO2, SNO4, SNR18, SNZ2, SPB1, SPC72, SPO7, SSA1, SUP56, SWC3, SWD1, SWH1, SYN8, TFC3, TGA1, THI11, TPD3, TRN1, UIP3, UPA1, VAC17, VBA3, VBA5, VPS8, VTH2, YAT1, YPS6, YRF1-2, YRF1-3, YRF1-4, YRF1-5, YRF1-6, YRF1-7, YRF1-8, YVH1, ZNF1, ZUO1 |
| **glu3** | ACM1, ADE1, ADF1, ADH4, ADH5, AGP2, ALG7, AMN1, APD1, APE3, APM3, ARA1, ARL1, ARO4, ATG42, ATP15, BGL2, BIO2, BIT2, BMT2, BRR6, BSD2, BUD14, CAB4, CCZ1, CDC15, CDC28, CDC39, CDC50, CHA1, CHK1, CIR2, CKS1, CNS1, COS5, COS6, COS9, CSH1, CTP1, CWC22, DFP1, DFP2, DLD3, DPB3, DUG2, DUT1, EFG1, EFM2, EMA35, ENP1, ERP1, ERR2, ERR3, ERV29, EXO5, FDH1, FEX1, FEX2, FIT2, FIT3, FLO5, FMP27, FRE2, FRE3, FRE5, FYV5, FZF1, GAB1, GDS1, GEX1, GIT1, GPX2, GTT1, HAB1, HAP5, HFM1, HIS7, HMLALPHA2, HMRA1, HMRA2, HSL7, HSM3, HSP32, HXK2, ICS2, IFA38, IMA1, IMD2, IRA1, ISW1, KAR4, KAR9, KIN3, KIN82, KRR1, KTD1, MAK5, MAL12, MAL13, MAL31, MAL32, MAL33, MATALPHA1, MCH2, MDL2, MEC1, MIC10, MIC12, MNT2, MRC1, MRPL27, MRPL37, MRPS5, MRPS9, MRS6, MSC6, MSH3, MST28, MTC4, NBP1, NPL4, NUP60, OCA4, OPY1, PAF1, PAU10, PAU12, PAU15, PAU19, PAU20, PAU21, PAU24, PAU6, PAU7, PAU8, PAU9, PBI1, PCA1, PDE1, PDE2, PDP3, PEX32, PEX34, PHO89, PHR1, PIP2, PLC1, POP4, POP7, PPS1, PRD1, PRE10, PRM9, PRT1, PXR1, RAD17, RAI1, RCF3, RDT1, REI1, RFA1, RGD1, RIB5, RIB7, RIF1, RIF2, RMD6, RME3, RMR1, RNH70, RPB5, RPS12, RRT2, RTC2, RTG2, RTT102, SAF1, SAM3, SAM4, SCP1, SCW4, SDH8, SEC66, SEN34, SEO1, SHE3, SHG1, SHM1, SLH1, SLI15, SLM6, SNF5, SNO4, SNX3, SPO23, SPP381, SRB6, SRY1, SSE2, SSH1, SUL1, SUP45, SUP56, SUT532, SWD1, TAE1, TAF1, TBS1, TFC3, TGA1, TRS20, TSC10, TYC1, TYR1, UBS1, UBX7, UIP3, UPA1, VAC17, VBA2, VBA3, VEL1, VPS8, VTH1, VTS1, YOR1, YPS6, YPT10, YSW1, YSY6, ZIP2, ZRT1, ZUO1 |

| | |
|---|---|
| **glu4** | AAD10, AAD4, ABD1, ABP1, ACS1, ADE1, ADE57, ADF1, ADH4, ADH6, ADY3, AGP3, AHC2, AIF1, AIM2, AIM6, ALD4, ALG7, ALR2, AME1, AMF1, ANK1, APA1, APE3, APM3, AQY1, AQY3, ARC40, ARN1, ARN2, ARO4, ATF1, ATG12, ATP15, ATS1, AVT2, AYT1, BAS1, BAT2, BDH1, BDH2, BGL2, BIO2, BIT2, BOL1, BOL3, BRR6, BSC5, BSD2, BUD14, CAB4, CAN1, CBP2, CCR4, CDC15, CDC19, CDC24, CDC39, CDC50, CHA1, CHK1, CIN8, CLN3, CNE1, COQ21, COS1, COS10, COS12, COS4, COS5, COS6, COS7, COS8, COX14, CSE1, CSM1, CSS1, CSS3, CTP1, CTR9, CWC22, CYC3, CYS3, DAD3, DAK2, DAL1, DAL2, DAL3, DAL4, DAL5, DAL7, DAL81, DAN1, DAN4, DCG1, DCP1, DDI2, DEP1, DFP1, DFP2, DFP4, DIA1, DLD3, DOC1, DPB3, DRS2, DSE4, DSF1, DUG2, DUT1, ECM1, ECM34, EFB1, EFG1, EFM1, EFM2, ELP6, EMA35, EMC1, EMC4, EMP47, ENB1, ENP1, ERG13, ERO1, ERP1, ERP2, ERR2, ERR3, ERT1, ERV15, ERV29, ERV46, ESL2, FDH1, FET4, FEX1, FEX2, FIG2, FIT2, FIT3, FLC2, FLO1, FLO10, FLO11, FLO5, FLO9, FMP32, FRE2, FRE3, FRE4, FRE5, FRE7, FRT2, FUN12, FUN14, FUN19, FUN26, FUN30, FYV5, FZF1, GAB1, GCV3, GDH1, GDH3, GEM1, GEX1, GEX2, GIP4, GIT1, GLC8, GPB1, GPB2, GPX2, GRE2, GTR1, GTT1, GTT2, GUD1, GUS1, HAB1, HAP2, HFM1, HIR3, HIS7, HMLALPHA2, HMRA1, HMRA2, HMS2, HOM6, HPA2, HPA3, HPC2, HRA1, HSM3, HSP32, HSU1, HXK2, HXT13, HXT15, HXT16, HXT17, HXT9, HYR1, ICR1, IMA1, IMA2, IMA3, IMA4, IMD2, IML1, INA22, IPA1, IRC24, IRC7, ISW1, JLP1, KAP114, KAR4, KAR9, KIN3, KIN82, KRR1, KTD1, LDS1, LRE1, LRG1, LSR1, LTE1, LYS1, MAK16, MAL12, MAL13, MAL31, MAL32, MAL33, MAN2, MATALPHA1, MCH2, MCM22, MCX1, MDL2, MDM10, MET5, MET8, MGA2, MGM101, MGR1, MHT1, MIC10, MIC12, MLC2, MLP1, MMP1, MND2, MNT2, MPH3, MRC1, MRPL27, MRPL37, MRPS5, MRS1, MRS6, MSC1, MSH3, MST28, MTC4, MTO1, MTW1, MUP3, MYO4, NBP1, NDD1, NDI1, NFT1, NGR1, NIP1, NMD5, NOP8, NPR2, NRE1, NTG1, NUD1, NUP60, OAF1, OCA4, OM14, OPT2, PAF1, PAU1, PAU10, PAU12, PAU13, PAU14, PAU15, PAU18, PAU19, PAU20, PAU21, PAU24, PAU4, PAU6, PAU7, PAU8, PAU9, PBI1, PBN1, PBP2, PCA1, PCC1, PCK1, PCS60, PDB1, PDE1, PDE2, PDP3, PDR18, PEX11, PEX22, PEX34, PGA3, PGU1, PHO11, PHO12, PHO84, PHO89, PHR1, PIP2, PMT2, PMT4, POF1, POP4, POP5, PPS1, PRD1, PRE10, PRE5, PRM9, PRP45, PRT1, PSF3, PSK1, PTA1, PUL3, PUL4, PWR1, PXP3, PXR1, PYC2, PZF1, QCR2, RAD17, RAI1, RBG1, RCF3, RDR1, RDT1, REI1, RFA1, RGD1, RGD2, RIB4, RIB5, RIF1, RMD6, RME3, RMR1, RNH70, ROT2, RPC82, RPO26, RPS12, RPS4A, RRP40, RRT2, RSC9, RTF1, RTG2, RTT102, SAF1, SAM3, SAM4, SAW1, SBP1, SCP1, SCW4, SDH8, SDS24, SEC11, SEC15, SEN34, SEO1, SGM1, SHG1, SHM1, SIR1, SIT1, SKG1, SKI3, SLH1, SLM6, SLX1, SNC1, SNF5, SNO2, SNO4, SNR18, SNZ2, SOR1, SPB1, SPC72, SPO23, SPO7, SPS22, SPT20, SRB6, SRB8, SSA1, SSH1, SUL1, SUP56, SUT532, SWC3, SWC5, SWD1, SWH1, SWP82, SYN8, TAD1, TAE1, TAF1, TAN1, TDA8, TDP1, TFC3, TGA1, TGL3, THI11, THI13, THI2, TIM8, TPD3, TRN1, TRS20, TRX3, TSC10, TTI2, TUB3, TUP1, TYC1, UBP11, UBX7, UIP3, UPA1, UPA2, VAC17, VBA2, VBA3, VBA5, VEL1, VHC1, VMR1, VPS8, VTH1, VTH2, VVS1, XPT1, YAT1, YBP1, YCT1, YOR1, YPS6, YPT10, YRF1-2, YRF1-3, YRF1-4, YRF1-5, YRF1-6, YRF1-7, YRF1-8, YVH1, ZIP2, ZNF1, ZPS1, ZRT1, ZUO1 |
| **glu5** | AAD10, ACS1, ADE1, ADF1, ADH4, AGP3, AIF1, AIM2, AMF1, APA1, AQY3, ARN1, ARN2, ATF1, BDH1, BDH2, BGL2, BIO2, BOL1, BOL3, BSC5, BUD14, CBP2, CDC15, CDC24, CDC39, CDC50, CHA1, CLN3, CNE1, COS1, COS10, COS4, COS5, COS6, COS8, COS9, CSS3, DAK2, DAL1, DAL2, DAL3, DAL4, DAL5, DAL7, DAL81, DAN4, DCG1, DDI2, DFP1, DFP2, DFP4, DLD3, DSE4, ECM1, ECM34, EFB1, EFM1, EMA35, EMC4, ERP1, ERP2, ERR2, ERR3, ERV29, ERV46, FEX1, FEX2, FIT2, FIT3, FLC2, FLO10, FLO5, FRE2, FRE3, FRE5, FUN14, FYV5, FZF1, GCV3, GDH3, GEM1, GEX1, GEX2, GIT1, GPB2, GTT1, HFM1, HMLALPHA2, HMRA1, HMRA2, HSP32, HXK2, HXT17, HYR1, IMA1, IMA3, IMD2, INA22, IRC24, IRC7, JEN1, KAR4, KIN3, KIN82, KRR1, KTD1, LRE1, LYS1, MAL12, MAL13, MAL31, MAL32, MATALPHA1, MCH2, MDM10, MGA2, MIC10, MND2, MNT2, MPH3, MRC1, MRS1, MSH3, MST28, MUP3, NFT1, NRE1, NUP60, OAF1, OCA4, PAU10, PAU12, PAU13, PAU14, PAU15, PAU19, PAU21, PAU24, PAU4, PAU6, PAU7, PAU8, PBN1, PDR18, PEX22, PEX34, PGU1, PHR1, POF1, PRD1, PRM9, PTA1, PUL3, PUL4, PXP3, RDR1, RDT1, RFA1, RMD6, RME3, RMR1, RTG2, SEC11, SEN34, SEO1, SIR1, SNO2, SNO4, SNR18, SNZ2, SOR1, SPB1, SPC72, SPO7, SPS22, SRY1, SSA1, SUP56, SUT532, SWD1, SWH1, TAN1, TFC3, TGA1, THI11, TRN1, UIP3, UPA1, URA1, VAC17, VBA3, VBA5, VEL1, VMR1, VPS8, VTH1, VTH2, YAT1, YPS6, YRF1-2, YRF1-3, YRF1-4, YRF1-6, YRF1-7, YRF1-8, YVH1, ZIP2, ZNF1, ZRT1, ZUO1 |

| | |
|---|---|
| glu6 | AAD10, AAD4, ABP1, ACS1, ACT1, ADE1, ADF1, ADH4, ADH6, AGP3, AIF1, AIM2, ALD4, ALR2, AMF1, APA1, AQY1, AQY3, ARN1, ARN2, ATF1, ATS1, AVT2, AYT1, BAS1, BAT2, BDH1, BDH2, BGL2, BIO2, BNA6, BOL1, BOL3, BSC5, BUD14, CBP2, CCR4, CDC15, CDC19, CDC24, CDC39, CDC50, CHA1, CLN3, CNE1, CNN1, COS1, COS10, COS12, COS4, COS5, COS6, COS7, COS8, COS9, COX14, CSM1, CSS1, CSS3, CYC3, CYS3, DAK2, DAL1, DAL2, DAL3, DAL4, DAL5, DAL7, DAL81, DAN1, DAN4, DCG1, DDI2, DEP1, DFP1, DFP2, DFP4, DIA1, DLD3, DRS2, DSF1, DSN1, DUG1, ECM1, ECM34, EFB1, EFM1, EGH1, EMA35, EMC1, EMP47, ENB1, ERG13, ERJ5, ERO1, ERP1, ERP2, ERR2, ERR3, ERV29, ERV46, FDC1, FDH1, FET4, FET5, FEX1, FEX2, FIG2, FIT1, FIT2, FIT3, FLC2, FLO1, FLO10, FLO11, FLO5, FLO9, FMP32, FRE2, FRE3, FRE5, FRT2, FUN12, FUN14, FUN19, FUN26, FUN30, FYV5, FZF1, GAT4, GCV3, GDH1, GDH3, GEM1, GEX1, GEX2, GIP4, GIT1, GLK1, GPB1, GPB2, GTR1, GTT1, GTT2, HAP5, HFM1, HIR3, HMLALPHA2, HMRA1, HMRA2, HMS2, HOM6, HPA2, HPA3, HRA1, HSP32, HSU1, HXK1, HXK2, HXT11, HXT13, HXT15, HXT16, HXT17, HXT9, HYR1, ICR1, IMA1, IMA2, IMA3, IMA4, IMA5, IMD2, IML1, INA22, IPA1, IRC24, IRC4, IRC6, IRC7, JEN1, JLP1, KAR4, KEG1, KGD4, KIN3, KIN82, KRR1, KTD1, LAM5, LDS1, LRE1, LTE1, LYS1, MAK16, MAL12, MAL13, MAL31, MAL32, MAL33, MAN2, MATALPHA1, MCH2, MCM22, MDM10, MET28, MET5, MGA2, MGM101, MGR1, MHT1, MIC10, MMP1, MND2, MNS1, MNT2, MPH3, MRC1, MRS1, MRS6, MRX20, MSC1, MSH3, MSL1, MST28, MTW1, MUP3, MYO4, NDD1, NDI1, NFT1, NGL3, NMD5, NRE1, NTG1, NUD1, NUP60, OAF1, OCA4, OPT2, OTU1, PAD1, PAU1, PAU10, PAU11, PAU12, PAU13, PAU14, PAU15, PAU18, PAU19, PAU20, PAU21, PAU24, PAU4, PAU6, PAU7, PAU8, PAU9, PBN1, PCK1, PDE2, PDI1, PDR18, PEX22, PEX34, PGA3, PGU1, PHO11, PHO12, PHO84, PHO89, PHR1, PIP2, PMT2, PMT4, POF1, POP5, PRD1, PRE10, PRE4, PRE5, PRI1, PRM9, PRP45, PRT1, PSK1, PTA1, PUG1, PWR1, PXP3, PXR1, QCR2, RAD17, RBG1, RDR1, RDT1, RET2, RFA1, RGD2, RMD6, RMD8, RME3, RMR1, RPN12, RPR2, RPS12, RPS4A, RSC9, RTG2, RUF21, SAM3, SAM4, SAW1, SCP1, SCW4, SEC11, SEC53, SEN34, SEO1, SGM1, SIR1, SIT1, SKG1, SNC1, SNO2, SNO4, SNR18, SNX3, SNZ2, SOR1, SPB1, SPC72, SPO7, SPS22, SQT1, SRY1, SSA1, STL1, STS1, SUP56, SUT532, SWC3, SWD1, SWH1, SWP82, SYN8, TDA8, TFC3, TGA1, THI11, THI13, TIM8, TPD3, TRN1, TTI2, TUB2, TUB3, TYC1, UBP11, UIP3, UPA1, URA1, VAC17, VBA3, VBA5, VEL1, VLD1, VPS8, VTH1, VTH2, VTS1, XPT1, YAP5, YAT1, YOR1, YPS6, YPT1, YRF1-2, YRF1-3, YRF1-4, YRF1-5, YRF1-6, YRF1-7, YRF1-8, YVH1, ZIP2, ZNF1, ZPS1, ZRT1, ZUO1 |
| gal1 | AAC3, ACH1, ADH5, AGP2, AIM3, AKL1, ALG1, ALG14, ALK2, AMN1, APD1, APL3, APN2, ARA1, ARL1, ATG14, ATG42, ATP3, BAP2, BMT2, CBP6, CCZ1, CDC28, CDS1, CHS2, CHS3, CKS1, CMD1, CNM1, CNS1, COQ1, COR1, CPP1, CSG2, CSH1, CST26, CYC8, DSF2, DTR1, ECM13, ECM15, ECM2, ECM31, ECM33, ECM8, EDE1, EDS1, EHT1, ERD2, ETR1, EXO5, EXO84, FAT1, FES1, FIG1, FLR1, FMP23, FMT1, FUI1, FUR4, FUS3, FZO1, GAL1, GAL10, GAL7, GIP1, GPI18, GRS1, GRX7, HAP3, HEK2, HHF1, HHT1, HIR1, HMT1, HSL7, HSP26, HTA2, HTB2, ICS2, IFA38, IML3, IPP1, IRA1, IST2, KAP104, LAA2, LDB7, LSM2, MAK5, MBA1, MCM2, MEC1, MEO1, MIN6, MIS1, MMS4, MNC1, MNN2, MOH1, MRPL16, MRPL36, MRPS9, MRX18, MUD1, MUM2, NCL1, NHP6B, NPL4, NRG2, NTH2, OLA1, OPY1, ORC2, PBY1, PDR3, PDX3, PEP1, PET9, PEX32, PFF1, PHO3, PHO5, PIM1, PIN4, POA1, POL12, POL30, POP7, POP8, PRE7, PRP6, PSY4, PTC4, QDR3, RCR1, RDH54, REB1, REG2, RER2, RFC5, RFS1, RFT1, RIB1, RIB7, RKM3, RPB5, RPG1, RPL19A, RPL19B, RPL4A, RPS11B, RPS6B, RRN10, RRN6, RRT1, RTC2, RXT2, SAS3, SCO1, SCO2, SCT1, SEC17, SEC18, SEC66, SHE1, SHE3, SIF2, SLA1, SLI15, SLM4, SMP1, SMY2, SND3, SNR161, SPP381, SPT7, SSE2, STU1, SUP45, SUS1, SWD3, TAT1, TBRT, TBS1, TCM62, TEC1, TFC1, TIM12, TIP1, TLC1, TPS1, TRM7, TSC3, TYR1, UBC4, UBP14, UBS1, UGA2, UMP1, URA7, UTP20, VID24, VMA2, VPS15, YMC2, YPC1, YPK3, YRO2, YSA1, YSW1, YSY6, ZTA1 |

| | |
|---|---|
| **gal2** | AAC3, ACH1, ADH5, AGP2, AIM3, AKL1, ALG1, ALG14, ALK2, AMN1, APD1, APL3, APN2, ARA1, ARL1, ATG14, ATG42, ATP3, BAP2, BMT2, CBP6, CCZ1, CDC28, CDS1, CHS2, CHS3, CKS1, CMC2, CMD1, CNM1, CNS1, COQ1, COR1, CPP1, CSG2, CSH1, CST26, CYC8, DSF2, ECM13, ECM15, ECM2, ECM33, ECM8, EDE1, EDS1, ERD2, ETR1, EXO5, EXO84, FAT1, FES1, FIG1, FLR1, FMP23, FMT1, FUI1, FUR4, FUS3, GAL1, GAL10, GAL7, GIP1, GPI18, GRS1, GRX7, HAP3, HEK2, HHF1, HHT1, HIR1, HMT1, HSL7, HSP26, HTA2, HTB2, IAI11, ICS2, IFA38, IML3, IPP1, IRA1, IST2, KAP104, KIP1, LAA2, LDB7, LSM2, MAK5, MCM2, MEC1, MEO1, MIN6, MIS1, MMS4, MNC1, MNN2, MOH1, MRPL16, MRPL36, MRPS9, MRX18, MUD1, MUM2, NCL1, NHP6B, NPL4, NRG2, NTH2, OLA1, OPY1, ORC2, PBY1, PDR3, PDX3, PEP1, PET9, PEX32, PFF1, PHO3, PHO5, PIM1, PIN4, POA1, POL12, POL30, POP7, POP8, PRE7, PRP6, PRS4, PRX1, PSY4, PTC3, PTC4, PTH2, QDR3, RCR1, RDH54, REB1, REG2, RER2, RFC5, RFS1, RFT1, RIB1, RIB7, RKM3, RPB5, RPG1, RPL19A, RPL19B, RPL4A, RPS11B, RRN10, RRN6, RRT1, RTC2, RXT2, SAS3, SCO1, SCO2, SCT1, SEC17, SEC18, SEC66, SEF1, SHE1, SHE3, SHP1, SIF2, SKT5, SLA1, SLI15, SLM4, SND3, SNR161, SPP381, SPT7, SSE2, STU1, SUP45, SUS1, TAT1, TBRT, TBS1, TCM62, TEC1, TFC1, TIM12, TIP1, TLC1, TOD6, TPS1, TRM7, TSC3, TYR1, UBC4, UBP13, UBP14, UBS1, UGA2, URA7, UTP20, VID24, VMA2, VPS15, YEL1, YMC2, YPK3, YRO2, YSA1, YSW1, YSY6, ZTA1 |
| **gal3** | AAC3, ACH1, ADH5, AGP2, AIM3, AKL1, ALG1, ALG14, ALK2, AMN1, APD1, APL3, APN2, ARA1, ARL1, ATG14, ATG42, ATP3, BAP2, BMT2, CBP6, CCZ1, CDC28, CDS1, CHS2, CHS3, CKS1, CMD1, CNM1, CNS1, COQ1, COR1, CPP1, CSG2, CSH1, CST26, CYC8, DSF2, DTR1, ECM13, ECM15, ECM2, ECM31, ECM33, ECM8, EDE1, EDS1, EHT1, ERD2, ETR1, EXO5, EXO84, FAT1, FES1, FIG1, FLR1, FMP23, FMT1, FUI1, FUR4, FUS3, FZO1, GAL1, GAL10, GAL7, GIP1, GPI18, GRS1, GRX7, HAP3, HEK2, HHF1, HHT1, HIR1, HMT1, HSL7, HSP26, HTA2, HTB2, ICS2, IFA38, IML3, IPP1, IRA1, IST2, KAP104, LAA2, LDB7, LSM2, MAK5, MBA1, MCM2, MEC1, MEO1, MIN6, MIS1, MMS4, MNC1, MNN2, MOH1, MRPL16, MRPL36, MRPS9, MRX18, MUD1, MUM2, NCL1, NHP6B, NPL4, NRG2, NTH2, OLA1, OPY1, ORC2, PBY1, PDR3, PDX3, PEP1, PET9, PEX32, PFF1, PHO3, PHO5, PIM1, PIN4, POA1, POL12, POL30, POP7, POP8, PRE7, PRP6, PSY4, PTC4, QDR3, RCR1, RDH54, REB1, REG2, RER2, RFC5, RFS1, RFT1, RIB1, RIB7, RKM3, RPB5, RPG1, RPL19A, RPL19B, RPL4A, RPS11B, RPS6B, RRN10, RRN6, RRT1, RTC2, RXT2, SAS3, SCO1, SCO2, SCT1, SEC17, SEC18, SEC66, SHE1, SHE3, SIF2, SLA1, SLI15, SLM4, SMP1, SMY2, SND3, SNR161, SPP381, SPT7, SSE2, STU1, SUP45, SUS1, SWD3, TAT1, TBRT, TBS1, TCM62, TEC1, TFC1, TIM12, TIP1, TLC1, TPS1, TRM7, TSC3, TYR1, UBC4, UBP14, UBS1, UGA2, UMP1, URA7, UTP20, VID24, VMA2, VPS15, YMC2, YPC1, YPK3, YRO2, YSA1, YSW1, YSY6, ZTA1 |
| **gal4** | AAC3, ACH1, ADH5, AGP2, AIM3, AKL1, ALG1, ALG14, ALK2, AMN1, APD1, APL3, APN2, ARA1, ARL1, ATG14, ATG42, ATP3, BAP2, BMT2, CBP6, CCZ1, CDC28, CDS1, CHS2, CHS3, CKS1, CMD1, CNM1, CNS1, COQ1, COR1, CPP1, CSG2, CSH1, CST26, CYC8, DSF2, DTR1, ECM13, ECM15, ECM2, ECM31, ECM33, ECM8, EDE1, EDS1, EHT1, ERD2, ETR1, EXO5, EXO84, FAT1, FES1, FIG1, FLR1, FMP23, FMT1, FUI1, FUR4, FUS3, FZO1, GAL1, GAL10, GAL7, GDT1, GIP1, GPI18, GRS1, GRX7, HAP3, HEK2, HHF1, HHT1, HIR1, HMT1, HSL7, HSP26, HTA2, HTB2, ICS2, IFA38, IML3, IPP1, IRA1, IST2, KAP104, LAA2, LDB7, LSM2, MAK5, MBA1, MCM2, MEC1, MEO1, MIN6, MIS1, MMS4, MNC1, MNN2, MOH1, MRPL16, MRPL36, MRPS9, MRX18, MUD1, MUM2, NCL1, NHP6B, NPL4, NRG2, NTC20, NTH2, OLA1, OPY1, ORC2, PBY1, PCH2, PDR3, PDX3, PEP1, PET9, PEX32, PFF1, PHO3, PHO5, PIM1, PIN4, POA1, POL12, POL30, POP7, POP8, PRE7, PRP6, PSY4, PTC4, QDR3, RCR1, RDH54, REB1, REG2, RER2, RFC5, RFS1, RFT1, RIB1, RIB7, RIM2, RKM3, RPB5, RPG1, RPL19A, RPL19B, RPL21A, RPL4A, RPS11B, RPS6B, RPS9B, RRN10, RRN6, RRT1, RTC2, RXT2, SAS3, SCO1, SCO2, SCT1, SEC17, SEC18, SEC66, SHE1, SHE3, SIF2, SLA1, SLI15, SLM4, SMP1, SMY2, SND3, SNR161, SPP381, SPT7, SSE2, STU1, SUP45, SUS1, SWD3, TAT1, TBRT, TBS1, TCM62, TEC1, TFC1, TIM12, TIP1, TLC1, TPS1, TRM7, TSC3, TYR1, UBC4, UBP14, UBS1, UGA2, UMP1, URA7, UTP20, VID24, VMA2, VPS15, YMC2, YPC1, YPK3, YRO2, YSA1, YSW1, YSY6, ZTA1 |

| | |
|---|---|
| gal5 | AAC3, ACH1, ADH5, AGP2, AIM3, AKL1, ALG1, ALG14, ALK2, AMN1, APD1, APL3, APN2, ARA1, ARL1, ATG14, ATG42, ATP3, BAP2, BMT2, CBP6, CCZ1, CDC28, CDS1, CHS2, CHS3, CKS1, CMD1, CNM1, CNS1, COQ1, COR1, CPP1, CSG2, CSH1, CST26, CYC8, DSF2, DTR1, ECM13, ECM15, ECM2, ECM31, ECM33, ECM8, EDE1, EDS1, EHT1, ERD2, ETR1, EXO5, EXO84, FAT1, FES1, FIG1, FLR1, FMP23, FMT1, FUI1, FUR4, FUS3, FZO1, GAL1, GAL10, GAL7, GDT1, GIP1, GPI18, GRS1, GRX7, HAP3, HEK2, HHF1, HHT1, HIR1, HMT1, HSL7, HSP26, HTA2, HTB2, ICS2, IFA38, IML3, IPP1, IRA1, IST2, KAP104, LAA2, LDB7, LSM2, MAK5, MBA1, MCM2, MEC1, MEO1, MIN6, MIS1, MMS4, MNC1, MNN2, MOH1, MRPL16, MRPL36, MRPS9, MRX18, MUD1, MUM2, NCL1, NHP6B, NPL4, NRG2, NTC20, NTH2, OLA1, OPY1, ORC2, PBY1, PCH2, PDR3, PDX3, PEP1, PET9, PEX32, PFF1, PHO3, PHO5, PIM1, PIN4, POA1, POL12, POL30, POP7, POP8, PRE7, PRP6, PSY4, PTC4, QDR3, RCR1, RDH54, REB1, REG2, RER2, RFC5, RFS1, RFT1, RIB1, RIB7, RIM2, RKM3, RPB5, RPG1, RPL19A, RPL19B, RPL21A, RPL4A, RPS11B, RPS6B, RPS9B, RRN10, RRN6, RRT1, RTC2, RXT2, SAS3, SCO1, SCO2, SCT1, SEC17, SEC18, SEC66, SHE1, SHE3, SIF2, SLA1, SLI15, SLM4, SMP1, SMY2, SND3, SNR161, SPP381, SPT7, SSE2, STU1, SUP45, SUS1, SWD3, TAT1, TBRT, TBS1, TCM62, TEC1, TFC1, TIM12, TIP1, TLC1, TPS1, TRM7, TSC3, TYR1, UBC4, UBP14, UBS1, UGA2, UMP1, URA7, UTP20, VID24, VMA2, VPS15, YMC2, YPC1, YPK3, YRO2, YSA1, YSW1, YSY6, ZTA1 |
| gal6 | AAC3, ACH1, ADH5, AGP2, AIM3, AKL1, ALG1, ALG14, ALK2, AMN1, APD1, APL3, APN2, ARA1, ARL1, ATG14, ATG42, ATP3, BAP2, BMT2, CBP6, CCZ1, CDC28, CDS1, CHS2, CHS3, CKS1, CMD1, CNM1, CNS1, COQ1, COR1, CPP1, CSG2, CSH1, CST26, CYC8, DSF2, DTR1, ECM13, ECM15, ECM2, ECM31, ECM33, ECM8, EDE1, EDS1, EHT1, ERD2, ETR1, EXO5, EXO84, FAT1, FES1, FIG1, FLR1, FMP23, FMT1, FUI1, FUR4, FUS3, FZO1, GAL1, GAL10, GAL7, GDT1, GIP1, GPI18, GRS1, GRX7, HAP3, HEK2, HHF1, HHT1, HIR1, HMT1, HSL7, HSP26, HTA2, HTB2, ICS2, IFA38, IML3, IPP1, IRA1, IST2, KAP104, LAA2, LDB7, LSM2, MAK5, MBA1, MCM2, MEC1, MEO1, MIN6, MIS1, MMS4, MNC1, MNN2, MOH1, MRPL16, MRPL36, MRPS9, MRX18, MUD1, MUM2, NCL1, NHP6B, NPL4, NRG2, NTC20, NTH2, OLA1, OPY1, ORC2, PBY1, PCH2, PDR3, PDX3, PEP1, PET9, PEX32, PFF1, PHO3, PHO5, PIM1, PIN4, POA1, POL12, POL30, POP7, POP8, PRE7, PRP6, PSY4, PTC4, QDR3, RCR1, RDH54, REB1, REG2, RER2, RFC5, RFS1, RFT1, RIB1, RIB7, RIM2, RKM3, RPB5, RPG1, RPL19A, RPL19B, RPL21A, RPL4A, RPS11B, RPS6B, RPS9B, RRN10, RRN6, RRT1, RTC2, RXT2, SAS3, SCO1, SCO2, SCT1, SEC17, SEC18, SEC66, SHE1, SHE3, SIF2, SLA1, SLI15, SLM4, SMP1, SMY2, SND3, SNR161, SPP381, SPT7, SSE2, STU1, SUP45, SUS1, SWD3, TAT1, TBRT, TBS1, TCM62, TEC1, TEF2, TFC1, TIM12, TIP1, TLC1, TPS1, TRM7, TSC3, TYR1, UBC4, UBP14, UBS1, UGA2, UMP1, URA7, UTP20, VID24, VMA2, VPS15, YMC2, YPC1, YPK3, YRO2, YSA1, YSW1, YSY6, ZTA1 |

**600 generations**

| Replicate | Gene list |
|---|---|
| glu1 | ACS1, ADF1, AIM2, BDH1, BDH2, CDC50, CHA1, CNE1, COS6, ECM1, FLC2, FLO5, FYV5, GDH3, GEX1, GPB2, HMLALPHA2, HMRA1, HMRA2, IMD2, KAR4, KRR1, MATALPHA1, MIC10, MRC1, OAF1, OCA4, PAU10, PAU24, PAU4, PAU6, PAU8, PEX22, PEX34, PRD1, RDT1, RMD6, SEO1, VAC17, VBA3, YRF1-6 |

| | |
|---|---|
| **glu2** | AAD10, AAD15, AAD4, AAR2, ACS1, ADE1, ADF1, ADH4, ADH7, ADY3, AGP3, AIF1, AIM2, ALG3, ALG7, APA1, APE3, APL3, APM3, AQY3, ARN1, ARN2, ARO4, AST1, ATG8, ATP1, AVT2, AVT5, BAS1, BAT2, BDH1, BDH2, BGL2, BIO2, BIT2, BNA4, BOI1, BRN1, BRR6, BSC5, BSD2, BUD14, CAB4, CAN1, CBP2, CDC15, CDC27, CDC39, CDC50, CHA1, CHK1, CMC2, CNE1, COR1, COS1, COS10, COS4, COS5, COS6, COS7, COS8, COS9, CSS3, CTP1, CWC22, DAK2, DAL2, DAL3, DAL4, DAL5, DAL7, DAN1, DAN4, DCG1, DDI2, DFP1, DFP2, DFP4, DLD3, DPB3, DSE4, DSF1, DUG2, DUT1, ECM1, ECM13, ECM15, ECM21, ECM34, EDE1, EFB1, EFG1, EFM1, EFM2, EMA35, EMC4, ENB1, ENP1, ERD2, ERO1, ERP1, ERP2, ERR2, ERR3, ERV29, ESL2, FDC1, FEX1, FEX2, FIT2, FIT3, FLC2, FLO10, FLO5, FMP27, FRE2, FRE3, FRE4, FRE5, FUI1, FUN14, FYV5, FZF1, GAB1, GDH3, GEM1, GEX1, GEX2, GIT1, GPB2, GPX2, GTT1, HAB1, HFM1, HIR3, HIS7, HMLALPHA2, HMRA1, HMRA2, HMS2, HOM6, HPA3, HSM3, HSP32, HTA2, HTB2, HXK2, HXT13, HXT15, HXT17, HYR1, IAI11, ILS1, IMA1, IMA3, IMD2, IML1, IPA1, IRC24, IRC4, IRC7, ISW1, KAR4, KIN3, KIN82, KIP1, KRR1, KTD1, KTI11, LRE1, LRG1, LYS1, MAL12, MAL13, MAL31, MAL32, MAL33, MAP2, MATALPHA1, MCH2, MCM22, MDM10, MET5, MGA2, MGM101, MIC10, MIC12, MIN6, MIX23, MLP1, MNS1, MNT2, MNT4, MOH1, MPH3, MRC1, MRP21, MRPL16, MRPL27, MRPL37, MRPS5, MRX3, MSH3, MST28, MTC4, MUP3, NBP1, NFT1, NMD5, NPR2, NRE1, NTH2, NUP170, NUP60, OAF1, OCA4, PAD1, PAF1, PAU10, PAU11, PAU12, PAU13, PAU14, PAU15, PAU18, PAU19, PAU20, PAU21, PAU24, PAU3, PAU4, PAU6, PAU7, PAU8, PAU9, PBN1, PCA1, PCC1, PCK1, PDE1, PDP3, PDR18, PDR3, PET112, PEX22, PEX34, PGU1, PHO89, PHR1, PIN4, PKC1, PMT4, POF1, POL12, POP4, PPS1, PRD1, PRE7, PRM9, PRS4, PRX1, PSY4, PTC3, PTH2, PUL3, PUL4, PXP3, PXR1, RAI1, RCF3, RDS1, RDT1, REI1, RFA1, RGD1, RIB5, RIF1, RIF2, RMD6, RME3, RMR1, RNH70, ROX3, RPL23A, RPL32, RPS4A, RPS8A, RRT1, RRT2, RTG2, RTG3, RTT102, SAF1, SAS3, SCS22, SCW4, SDH8, SEA4, SEC17, SEF1, SEN34, SEO1, SFT2, SGM1, SHG1, SHM1, SHP1, SIR1, SIT1, SKG1, SKT5, SLH1, SLM6, SNF5, SNO2, SNO4, SNR18, SNR56, SNZ2, SOR1, SPB1, SPO23, SPO7, SPS22, SRB6, SRO77, SRY1, SSA1, SSA3, SSH1, SST2, STR2, SUL1, SUP56, SUT532, SWD1, TAE1, TAF1, TAN1, TEL1, TFC3, TGA1, THI11, THI13, TIM8, TOD6, TRN1, TRS20, TSC10, TTI2, TYC1, UBP11, UBP13, UBX7, UIP3, UPA1, URA7, UTP20, VAC17, VBA2, VBA3, VBA5, VEL1, VMR1, VPS8, VTH1, VTH2, XPT1, YEL1, YOR1, YPS6, YPT10, YRF1-2, YRF1-6, ZIP2, ZNF1, ZRT1, ZUO1 |
| **glu3** | AAC3, ABD1, ACS1, ADE1, ADH5, AGP2, AIM2, AIM4, ALG7, AME1, AMN1, APD1, APE3, APM3, ARA1, ARC40, ARL1, ARO4, ATG12, ATG42, BDH1, BDH2, BEM1, BIT2, BMT2, BSD2, BUD14, CCZ1, CDC15, CDC28, CHK1, CKS1, CNE1, CNS1, COQ21, COS111, CSH1, CSS1, CTP1, DAD3, DER1, DFP1, DFP2, DPB3, DTR1, DUG2, DUR12, DUT1, ECM1, ECM31, EFB1, EFM2, EHT1, ENP1, ERP1, ERP2, ERT1, ERV15, EXO5, EXO84, FES1, FLC2, FLO1, FLO5, FTH1, FUN14, FZO1, GDH3, GDT1, GEM1, GPB2, GPX2, HAB1, HIS7, HPC2, HSL7, HSM3, HXT9, ICS2, IFA38, IMA2, IMA4, IRA1, IST2, ISW1, KIN3, KTD1, KTR3, KTR4, LDH1, LSR1, MAK5, MAL33, MBA1, MCM7, MCX1, MDM10, MEC1, MED8, MET8, MIC12, MIN7, MIS1, MMS4, MRPL27, MRPL37, MRPS5, MRPS9, MSI1, MST28, MTC4, NGR1, NHP6B, NPL4, NTC20, NUP60, OAF1, OM14, PAF1, PAU15, PAU18, PAU20, PAU7, PAU8, PAU9, PBP2, PBY1, PCA1, PCH2, PCS60, PDB1, PEX22, PEX32, PGI1, PHO3, PHO5, PHO89, POL30, POP4, POP7, PPS1, PRM9, PYC2, RCF3, REI1, RFA1, RFC5, RGD1, RIB5, RIB7, RIF1, RIM2, ROT2, RPB5, RPL19A, RPL21A, RPS6B, RPS9B, RRT2, RTC2, RXT2, SAF1, SDH8, SDS24, SEC66, SEN34, SEO1, SHG1, SHM1, SIF2, SLI15, SLM6, SLX1, SMP1, SMY2, SNF5, SNR18, SPO23, SPO7, SPP381, SRB6, SSA1, SSE2, SSH1, SUL1, SUP45, SUP56, SWC5, SWD1, SWD3, SWH1, TAE1, TAF5, TBS1, TDP1, TEC1, TFC3, TGA1, THI2, TIM12, TRN1, TRS20, TSC10, TYC1, TYR1, UBS1, UBX7, UIP3, UMP1, VBA2, VHC1, VID24, VPS15, VPS8, VTH1, VVS1, YAT1, YBP1, YMC2, YPC1, YPT10, YSW1, YSY6 |

| | |
|---|---|
| **glu4** | AAC3, ABD1, ACS1, ADE1, ADF1, ADH5, AGP2, AIM2, AIM3, ALG1, ALG7, AME1, AMN1, APD1, APE3, APM3, ARA1, ARC40, ARL1, ARO4, ATG12, ATG14, ATG42, BDH1, BDH2, BEM1, BGL2, BIO2, BIT2, BMT2, BSD2, BUD14, CBP6, CCZ1, CDC15, CDC28, CDC50, CHA1, CHK1, CKS1, CMD1, CNE1, CNS1, COQ21, COS111, COS6, CSH1, CTP1, CYC8, CYS3, DAD3, DEP1, DER1, DFP1, DFP2, DPB3, DUG2, DUR12, DUT1, ECM1, EFB1, EFM2, EMA35, ENP1, ERP1, ERP2, ERT1, ERV15, ERV29, EXO5, EXO84, FES1, FLC2, FLO5, FTH1, FUN14, FYV5, GDH3, GEM1, GEX1, GIT1, GPB2, GPX2, GRS1, GTT1, HAB1, HIS7, HMLALPHA2, HMRA1, HMRA2, HPC2, HSL7, HSM3, HYR1, ICS2, IFA38, IMA1, IMD2, IML3, IRA1, IST2, ISW1, KAR4, KIN3, KRR1, KTD1, KTR3, KTR4, LDH1, LSR1, LYS2, MAK5, MAL12, MAL13, MAL31, MAL32, MAL33, MATALPHA1, MCH2, MCM7, MCX1, MDM10, MEC1, MEO1, MET8, MIC10, MIC12, MIN7, MMS4, MRC1, MRPL27, MRPL36, MRPL37, MRPS5, MRPS9, MST28, MTC4, MUD1, NGR1, NHP6B, NPL4, NTG1, NUP60, OAF1, OCA4, OM14, OPY1, PAF1, PAU10, PAU12, PAU19, PAU24, PAU4, PAU6, PAU7, PAU8, PAU9, PBP2, PBY1, PCA1, PCS60, PDB1, PEX22, PEX32, PEX34, PHO3, PHO5, PHO89, POL30, POP4, POP7, PPS1, PRD1, PRM9, PSK1, PTC4, PYC2, RAD16, RCF3, RDT1, REI1, RFA1, RFC5, RGD1, RIB5, RIB7, RIF1, RMD6, ROT2, RPB5, RRT2, RTC2, RXT2, SAF1, SDH8, SDS24, SEC66, SEN34, SEO1, SHE3, SHG1, SHM1, SIF2, SLI15, SLM6, SLX1, SND3, SNF5, SNR18, SPB1, SPO23, SPO7, SPP381, SRB6, SSA1, SSE2, SSH1, SUL1, SUP45, SUP56, SUS1, SWC3, SWC5, SWD1, SWH1, SYN8, TAE1, TBS1, TDP1, TEF2, TFC1, TFC3, TGA1, THI2, TIM12, TKL2, TPD3, TPS1, TRN1, TRS20, TSC10, TYC1, TYR1, UBS1, UBX7, UIP3, UPA1, VAC17, VBA2, VBA3, VHC1, VID24, VMA2, VPS15, VPS8, VTH2, VVS1, YAT1, YBP1, YMC2, YPS6, YPT10, YRF1-4, YRF1-6, YSA1, YSW1, YSY6, ZUO1 |
| **glu5** | AAD10, AAD4, ACS1, ADE1, ADF1, ADH4, AGP3, AIF1, AIM2, ALD4, ALR2, AMF1, APA1, AQY3, ATF1, AVT2, BAS1, BAT2, BDH1, BDH2, BGL2, BIO2, BRR6, BSC5, BUD14, CAN1, CDC15, CDC39, CDC50, CHA1, CNE1, COS1, COS10, COS4, COS5, COS6, COS7, COX14, CSS3, DAK2, DAL5, DAN1, DAN4, DDI2, DFP1, DFP2, DLD3, DSE4, DSF1, ECM1, EFB1, EMA35, EMC4, EMP47, ENB1, ERO1, ERP1, ERP2, ERR2, ERR3, ERV29, ESL2, FDC1, FEX1, FIT2, FIT3, FLC2, FLO10, FLO5, FRE2, FRE3, FRE4, FRE5, FUN14, FYV5, FZF1, GDH1, GDH3, GEM1, GEX1, GEX2, GIT1, GPB1, GPB2, HFM1, HIR3, HMLALPHA2, HMRA1, HMRA2, HMS2, HOM6, HPA3, HSP32, HXK2, HXT11, HXT13, HXT15, HXT17, HXT8, IMA1, IMA2, IMA5, IMD2, IML1, IPA1, IRC4, IRC7, KAR4, KIN3, KIN82, KRR1, KTD1, LRE1, LRG1, MAL12, MAL13, MAL31, MAL32, MATALPHA1, MCH2, MCM22, MDM10, MET5, MGM101, MIC10, MIX23, MNS1, MNT2, MPH3, MRC1, MRS6, MSH3, MST28, NDD1, NFT1, NMD5, NPR2, NUD1, NUP60, OAF1, OCA4, PAD1, PAU10, PAU11, PAU12, PAU14, PAU15, PAU19, PAU21, PAU24, PAU4, PAU6, PAU7, PAU8, PBN1, PCC1, PCK1, PDE1, PDE2, PDR18, PEX22, PEX34, PGU1, PHR1, PIP2, PKC1, PMT4, PRD1, PRE10, PRM9, PRT1, PUL3, PUL4, RAD17, RAI1, RDR1, RDT1, RFA1, RMD6, RME3, RMR1, RPS12, RPS4A, RTG2, RTG3, SCP1, SEA4, SEN34, SEO1, SFT2, SGM1, SIR1, SIT1, SKG1, SNO2, SNO4, SNR18, SNZ2, SOR1, SPB1, SPO7, SRO77, SSA1, SUP56, SUT532, SWD1, SWH1, SWP82, TAN1, TFC3, TGA1, THI11, THI13, TIM8, TRN1, TTI2, UBP11, UIP3, UPA1, VAC17, VBA3, VBA5, VEL1, VPS8, VTH1, VTH2, XPT1, YAT1, YPS6, YRF1-2, YRF1-4, YRF1-6, YRF1-8, ZIP2, ZNF1, ZRT1, ZUO1 |
| **glu6** | RMD6, SEO1 |
| **gal1** | AAC3, ACH1, ADH5, AGP2, AIM3, AKL1, ALG1, ALG14, ALK2, AMN1, APD1, APL3, APN2, ARA1, ARL1, ATG14, ATG42, ATP3, BAP2, BMT2, CBP6, CCZ1, CDC28, CDS1, CHS2, CHS3, CKS1, CMD1, CNM1, CNS1, COQ1, COR1, CPP1, CSG2, CSH1, CST26, CYC8, DSF2, DTR1, ECM13, ECM15, ECM2, ECM31, ECM33, ECM8, EDE1, EDS1, EHT1, ERD2, ETR1, EXO5, EXO84, FAT1, FES1, FIG1, FLR1, FMP23, FMT1, FUI1, FUR4, FUS3, FZO1, GAL1, GAL10, GAL7, GDT1, GIP1, GPI18, GRS1, GRX7, HAP3, HEK2, HHF1, HHT1, HIR1, HMT1, HSL7, HSP26, HTA2, HTB2, ICS2, IFA38, IML3, IPP1, IRA1, IST2, KAP104, LAA2, LDB7, LSM2, MAK5, MBA1, MCM2, MEC1, MEO1, MIN6, MIS1, MMS4, MNC1, MNN2, MOH1, MRPL16, MRPL36, MRPS9, MRX18, MUD1, MUM2, NCL1, NHP6B, NPL4, NRG2, NTC20, NTH2, OLA1, OPY1, ORC2, PBY1, PCH2, PDR3, PDX3, PEP1, PET9, PEX32, PFF1, PHO3, PHO5, PIM1, PIN4, POA1, POL12, POL30, POP7, POP8, PRE7, PRP6, PSY4, PTC4, QDR3, RCR1, RDH54, REB1, REG2, RER2, RFC5, RFS1, RFT1, RIB1, RIB7, RKM3, RPB5, RPG1, RPL19A, RPL19B, RPL21A, RPL4A, RPS11B, RPS6B, RPS9B, RRN10, RRN6, RRT1, RTC2, RXT2, SCO1, SCO2, SCT1, SEC17, SEC18, SEC66, SHE1, SHE3, SIF2, SLA1, SLI15, SLM4, SMP1, SMY2, SND3, SNR161, SPP381, SPT7, SSE2, STU1, SUP45, SUS1, SWD3, TAT1, TBRT, TBS1, TCM62, TEC1, TFC1, TIM12, TIP1, TLC1, TPS1, TRM7, TSC3, TYR1, UBC4, UBP14, UBS1, UGA2, UMP1, URA7, UTP20, VID24, VMA2, VPS15, YMC2, YPC1, YPK3, YRO2, YSA1, YSW1, YSY6, ZTA1 |

| | |
|---|---|
| **gal2** | AAC3, ACH1, ADH5, AGP2, AIM3, AKL1, ALG1, ALG14, ALK2, AMN1, APD1, APL3, APN2, ARA1, ARL1, ATG14, ATG42, ATP3, BAP2, BMT2, CBP6, CCZ1, CDC28, CDS1, CHS2, CHS3, CKS1, CMD1, CNM1, CNS1, COQ1, COR1, CPP1, CSG2, CSH1, CST26, CYC8, DSF2, DTR1, ECM13, ECM15, ECM2, ECM31, ECM33, ECM8, EDE1, EDS1, EHT1, ERD2, ETR1, EXO5, EXO84, FAT1, FES1, FIG1, FLR1, FMP23, FMT1, FUI1, FUR4, FUS3, FZO1, GAL1, GAL10, GAL7, GDT1, GIP1, GPI18, GRS1, GRX7, HAP3, HEK2, HHF1, HHT1, HIR1, HMT1, HSL7, HSP26, HTA2, HTB2, ICS2, IFA38, IML3, IPP1, IRA1, IST2, KAP104, LAA2, LDB7, LSM2, MAK5, MBA1, MCM2, MEC1, MEO1, MIN6, MIS1, MMS4, MNC1, MNN2, MOH1, MRPL16, MRPL36, MRPS9, MRX18, MUD1, MUM2, NCL1, NHP6B, NPL4, NRG2, NTC20, NTH2, OLA1, OPY1, ORC2, PBY1, PCH2, PDR3, PDX3, PEP1, PET9, PEX32, PFF1, PHO3, PHO5, PIM1, PIN4, POA1, POL12, POL30, POP7, POP8, PRE7, PRP6, PSY4, PTC4, QDR3, RCR1, RDH54, REB1, REG2, RER2, RFC5, RFS1, RFT1, RIB1, RIB7, RIM2, RKM3, RPB5, RPG1, RPL19A, RPL19B, RPL21A, RPL4A, RPS11B, RPS6B, RPS9B, RRN10, RRN6, RRT1, RTC2, RXT2, SAS3, SCO1, SCO2, SCT1, SEC17, SEC18, SEC66, SHE1, SHE3, SIF2, SLA1, SLI15, SLM4, SMP1, SMY2, SND3, SNR161, SPP381, SPT7, SSE2, STU1, SUP45, SUS1, SWD3, TAT1, TBRT, TBS1, TCM62, TEC1, TFC1, TIM12, TIP1, TLC1, TPS1, TRM7, TSC3, TYR1, UBC4, UBP14, UBS1, UGA2, UMP1, URA7, UTP20, VID24, VMA2, VPS15, YMC2, YPC1, YPK3, YRO2, YSA1, YSW1, YSY6, ZTA1 |
| **gal3** | AAC3, ACH1, ADH5, AGP2, AIM3, AIM4, AKL1, ALG1, ALG14, ALK2, AME1, AMN1, APD1, APL3, APN2, ARA1, ARL1, ATG14, ATG42, ATP3, BAP2, BEM1, BMT2, CBP6, CCZ1, CDC28, CDS1, CHS2, CHS3, CKS1, CMD1, CNM1, CNS1, COQ1, COR1, COS111, CPP1, CSG2, CSH1, CST26, CYC8, DER1, DSF2, DTR1, DUR12, ECM13, ECM15, ECM2, ECM31, ECM33, ECM8, EDE1, EDS1, EHT1, ERD2, ERV15, ETR1, EXO5, EXO84, FAT1, FES1, FIG1, FLR1, FMP23, FMT1, FTH1, FUI1, FUR4, FUS3, FZO1, GAL1, GAL10, GAL7, GDT1, GIP1, GPI18, GRS1, GRX7, HAP3, HEK2, HHF1, HHT1, HIR1, HMT1, HSL7, HSP26, HTA2, HTB2, ICS2, IFA38, IML3, IPP1, IRA1, IST2, KAP104, KTR3, KTR4, LAA2, LDB7, LDH1, LSM2, MAK5, MBA1, MCM2, MCM7, MEC1, MED8, MEO1, MIN6, MIN7, MIS1, MMS4, MNC1, MNN2, MOH1, MRPL16, MRPL36, MRPS9, MRX18, MSI1, MUD1, MUM2, NCL1, NHP6B, NPL4, NRG2, NTC20, NTH2, OLA1, OPY1, ORC2, PBY1, PCH2, PDR3, PDX3, PEP1, PET9, PEX32, PFF1, PGI1, PHO3, PHO5, PIM1, PIN4, POA1, POL12, POL30, POP7, POP8, PRE7, PRP6, PSY4, PTC4, QDR3, RCR1, RDH54, REB1, REG2, RER2, RFC5, RFS1, RFT1, RIB1, RIB7, RIM2, RKM3, RPB5, RPG1, RPL19A, RPL19B, RPL21A, RPL4A, RPS11B, RPS6B, RPS9B, RRN10, RRN6, RRT1, RTC2, RXT2, SAS3, SCO1, SCO2, SCT1, SEC17, SEC18, SEC66, SHE1, SHE3, SIF2, SLA1, SLI15, SLM4, SMP1, SMY2, SND3, SNR161, SPP381, SPT7, SSE2, STU1, SUP45, SUS1, SWD3, TAF5, TAT1, TBRT, TBS1, TCM62, TEC1, TFC1, TIM12, TIP1, TLC1, TPS1, TRM7, TSC3, TYR1, UBC4, UBP14, UBS1, UGA2, UMP1, URA7, UTP20, VID24, VMA2, VPS15, YMC2, YPC1, YPK3, YRO2, YSA1, YSW1, YSY6, ZTA1 |
| **gal4** | AAC3, ACH1, ADH5, AGP2, AIM3, AKL1, ALG1, ALG14, ALK2, AMN1, APD1, APL3, APN2, ARA1, ARL1, ATG14, ATG42, ATP3, BAP2, BMT2, CBP6, CCZ1, CDC28, CDS1, CHS2, CHS3, CKS1, CMD1, CNM1, CNS1, COQ1, COR1, CPP1, CSG2, CSH1, CST26, CYC8, DSF2, DTR1, ECM13, ECM15, ECM2, ECM31, ECM33, ECM8, EDE1, EDS1, EHT1, ERD2, ETR1, EXO5, EXO84, FAT1, FES1, FIG1, FLR1, FMP23, FMT1, FUI1, FUR4, FUS3, FZO1, GAL1, GAL10, GAL7, GDT1, GIP1, GPI18, GRS1, GRX7, HAP3, HEK2, HHF1, HHT1, HIR1, HMT1, HSL7, HSP26, HTA2, HTB2, ICS2, IFA38, IML3, IPP1, IRA1, IST2, KAP104, LAA2, LDB7, LSM2, MAK5, MBA1, MCM2, MEC1, MEO1, MIN6, MIS1, MMS4, MNC1, MNN2, MOH1, MRPL16, MRPL36, MRPS9, MRX18, MUD1, MUM2, NCL1, NHP6B, NPL4, NRG2, NTC20, NTH2, OLA1, OPY1, ORC2, PBY1, PCH2, PDR3, PDX3, PEP1, PET9, PEX32, PFF1, PHO3, PHO5, PIM1, PIN4, POA1, POL12, POL30, POP7, POP8, PRE7, PRP6, PSY4, PTC4, QDR3, RCR1, RDH54, REB1, REG2, RER2, RFC5, RFS1, RFT1, RIB1, RIB7, RKM3, RPB5, RPG1, RPL19A, RPL19B, RPL4A, RPS11B, RPS6B, RRN10, RRN6, RRT1, RTC2, RXT2, SAS3, SCO1, SCO2, SCT1, SEC17, SEC18, SEC66, SHE1, SHE3, SIF2, SLA1, SLI15, SLM4, SMP1, SMY2, SND3, SNR161, SPP381, SPT7, SSE2, STU1, SUP45, SUS1, SWD3, TAT1, TBRT, TBS1, TCM62, TEC1, TFC1, TIM12, TIP1, TLC1, TPS1, TRM7, TSC3, TYR1, UBC4, UBP14, UBS1, UGA2, UMP1, URA7, UTP20, VID24, VMA2, VPS15, YMC2, YPC1, YPK3, YRO2, YSA1, YSW1, YSY6, ZTA1 |

| | |
|---|---|
| **gal5** | AAC3, ACH1, ADH5, AGP2, AIM3, AKL1, ALG1, ALG14, ALK2, AMN1, APD1, APL3, APN2, ARA1, ARL1, ATG14, ATG42, ATP3, BAP2, BMT2, CBP6, CCZ1, CDC28, CDS1, CHS2, CHS3, CKS1, CMD1, CNM1, CNS1, COQ1, COR1, CPP1, CSG2, CSH1, CST26, CYC8, DSF2, DTR1, ECM13, ECM15, ECM2, ECM31, ECM33, ECM8, EDE1, EDS1, EHT1, ERD2, ETR1, EXO5, EXO84, FAT1, FES1, FIG1, FLR1, FMP23, FMT1, FUI1, FUR4, FUS3, FZO1, GAL1, GAL10, GAL7, GIP1, GPI18, GRS1, GRX7, HAP3, HEK2, HHF1, HHT1, HIR1, HMT1, HSL7, HSP26, HTA2, HTB2, ICS2, IFA38, IML3, IPP1, IRA1, IST2, KAP104, LAA2, LDB7, LSM2, MAK5, MBA1, MCM2, MEC1, MEO1, MIN6, MIS1, MMS4, MNC1, MNN2, MOH1, MRPL16, MRPL36, MRPS9, MRX18, MUD1, MUM2, NCL1, NHP6B, NPL4, NRG2, NTH2, OLA1, OPY1, ORC2, PBY1, PDR3, PDX3, PEP1, PET9, PEX32, PFF1, PHO3, PHO5, PIM1, PIN4, POA1, POL12, POL30, POP7, POP8, PRE7, PRP6, PSY4, PTC4, QDR3, RCR1, RDH54, REB1, REG2, RER2, RFC5, RFS1, RFT1, RIB1, RIB7, RKM3, RPB5, RPG1, RPL19A, RPL19B, RPL4A, RPS11B, RPS6B, RRN10, RRN6, RRT1, RTC2, RXT2, SAS3, SCO1, SCO2, SCT1, SEC17, SEC18, SEC66, SHE1, SHE3, SIF2, SLA1, SLI15, SLM4, SMP1, SMY2, SND3, SNR161, SPP381, SPT7, SSE2, STU1, SUP45, SUS1, SWD3, TAT1, TBRT, TBS1, TCM62, TEC1, TFC1, TIM12, TIP1, TLC1, TPS1, TRM7, TSC3, TYR1, UBC4, UBP14, UBS1, UGA2, UMP1, URA7, UTP20, VID24, VMA2, VPS15, YMC2, YPC1, YPK3, YRO2, YSA1, YSW1, YSY6, ZTA1 |
| **gal6** | AAC3, ACH1, ADH5, AGP2, AIM3, AKL1, ALG1, ALG14, ALK2, AMN1, APD1, APL3, APN2, ARA1, ARL1, AST1, ATG14, ATG42, ATP3, BAP2, BMT2, CBP6, CCZ1, CDC28, CDS1, CHS2, CHS3, CKS1, CMC2, CMD1, CNM1, CNS1, COQ1, COR1, CPP1, CSG2, CSH1, CST26, CYC8, DSF2, DTR1, ECM13, ECM15, ECM2, ECM31, ECM33, ECM8, EDE1, EDS1, EHT1, ERD2, ETR1, EXO5, EXO84, FAT1, FES1, FIG1, FLR1, FMP23, FMT1, FUI1, FUR4, FUS3, FZO1, GAL1, GAL10, GAL7, GIP1, GPI18, GRS1, GRX7, HAP3, HEK2, HHF1, HHT1, HIR1, HMT1, HSL7, HSP26, HTA2, HTB2, IAI11, ICS2, IFA38, IML3, IPP1, IRA1, IST2, KAP104, KIP1, KTI11, LAA2, LDB7, LSM2, MAK5, MBA1, MCM2, MEC1, MEO1, MIN6, MIS1, MMS4, MNC1, MNN2, MOH1, MRPL16, MRPL36, MRPS9, MRX18, MUD1, MUM2, NCL1, NHP6B, NPL4, NRG2, NTH2, OLA1, OPY1, ORC2, PBY1, PDR3, PDX3, PEP1, PET9, PEX32, PFF1, PHO3, PHO5, PIM1, PIN4, POA1, POL12, POL30, POP7, POP8, PRE7, PRP6, PRS4, PRX1, PSY4, PTC3, PTC4, PTH2, QDR3, RCR1, RDH54, REB1, REG2, RER2, RFC5, RFS1, RFT1, RIB1, RIB7, RKM3, RPB5, RPG1, RPL19A, RPL19B, RPL4A, RPS11B, RPS6B, RPS8A, RRN10, RRN6, RRT1, RTC2, RXT2, SAS3, SCO1, SCO2, SCT1, SEC17, SEC18, SEC66, SEF1, SHE1, SHE3, SHP1, SIF2, SKT5, SLA1, SLI15, SLM4, SMP1, SMY2, SND3, SNR161, SNR56, SPP381, SPT7, SSE2, STU1, SUP45, SUS1, SWD3, TAT1, TBRT, TBS1, TCM62, TEC1, TFC1, TIM12, TIP1, TLC1, TOD6, TPS1, TRM7, TSC3, TYR1, UBC4, UBP13, UBP14, UBS1, UGA2, UMP1, URA7, UTP20, VID24, VMA2, VPS15, YEL1, YMC2, YPC1, YPK3, YRO2, YSA1, YSW1, YSY6, ZTA1 |

**900 generations**

| Replicate | Gene list |
|---|---|
| **glu1** | AAC3, AAR2, ACH1, ADH5, AGP2, AKL1, ALG14, ALK2, AMN1, APD1, APL3, APN2, ARA1, ARL1, AST1, ATG14, ATG42, ATG8, ATP3, BAP2, BMT2, CBP6, CCZ1, CDC28, CHS3, CKS1, CMC2, CNM1, CNS1, COQ1, COR1, CPP1, CSH1, CST26, DSF2, ECM13, ECM15, ECM2, ECM33, ECM8, EDE1, ERD2, ETR1, EXO5, EXO84, FAT1, FES1, FIG1, FLR1, FMP23, FMT1, FUI1, FUR4, FUS3, GAL1, GAL10, GAL7, GIP1, GPI18, GRS1, GRX7, HAP3, HEK2, HHF1, HHT1, HIR1, HSL7, HSP26, HTA2, HTB2, IAI11, ICS2, IFA38, ILS1, IPP1, IRA1, IST2, KAP104, KIP1, KTI11, LAA2, LDB7, LSM2, MAK5, MCM2, MEC1, MEO1, MIN6, MIS1, MIX23, MMS4, MNC1, MNN2, MOH1, MRPL16, MRPL36, MRPS9, MRX18, MUD1, MUM2, NCL1, NHP6B, NPL4, NRG2, NTH2, OLA1, OPY1, ORC2, PBY1, PDR3, PEP1, PET9, PEX32, PFF1, PHO3, PHO5, PIM1, PIN4, PKC1, POA1, POL12, POL30, POP7, POP8, PRE7, PRP6, PRS4, PRX1, PSY4, PTC3, PTC4, PTH2, QDR3, RCR1, RDH54, REB1, REG2, RER2, RFC5, RFS1, RFT1, RIB1, RIB7, RPB5, RPG1, RPL19A, RPL19B, RPL23A, RPS11B, RPS8A, RRN10, RRN6, RRT1, RTC2, RXT2, SAS3, SCO2, SCT1, SEA4, SEC17, SEC18, SEC66, SEF1, SEO1, SHE1, SHE3, SHP1, SKT5, SLA1, SLI15, SLM4, SNR56, SPP381, SPT7, SRO77, SSA3, SSE2, STU1, SUP45, TAT1, TBRT, TBS1, TCM62, TEC1, TFC1, TIM12, TIP1, TOD6, TPS1, TRM7, TSC3, TYR1, UBC4, UBP13, UBP14, UBS1, UGA2, URA7, UTP20, VMA2, VPS15, YEL1, YPK3, YRO2, YSW1, YSY6, ZTA1 |

| | |
|---|---|
| **glu2** | AAC3, AAR2, ABD1, ACH1, ACS1, ADE1, ADH5, AIM2, AKL1, ALG14, ALG3, ALG7, ALK2, APD1, APE3, APL3, APM3, APN2, ARA1, ARC40, ARO4, AST1, ATG12, ATG42, ATG8, ATP1, AVT5, BAP2, BDH1, BDH2, BIT2, BMT2, BNA4, BOI1, BOL1, BOL3, BRN1, BSD2, BUD14, CDC15, CDC19, CDC24, CDC27, CDS1, CHK1, CHS3, CLN3, CMC2, CNE1, CNM1, COQ1, COQ21, COR1, CPP1, CTP1, CYC3, DAD3, DFP2, DPB3, DSF2, DUG2, DUT1, ECM1, ECM13, ECM15, ECM2, ECM21, ECM33, ECM8, EDE1, EDS1, EFB1, EFM2, ENP1, ERD2, ERP1, ERP2, ERT1, ERV46, ETR1, FLC2, FLO5, FLR1, FMT1, FUI1, FUN14, FUR4, FUS3, GAL1, GAL10, GAL7, GCV3, GDH3, GEM1, GPB2, GPI18, GPX2, GRX7, HAB1, HAP3, HEK2, HHF1, HHT1, HIR1, HIS7, HPC2, HSM3, HSP26, HTA2, HTB2, IAI11, ILS1, IMD2, IPP1, IRA1, IST2, ISW1, KAP104, KIN3, KIP1, KTI11, LAA2, LDB7, LSM2, LSR1, MAK5, MAL33, MAP2, MCM2, MCX1, MIC12, MIN6, MIS1, MIX23, MNN2, MOH1, MRP21, MRPL16, MRPL27, MRPL37, MRPS5, MRPS9, MRX3, MST28, MTC4, NCL1, NHP6B, NRG2, NTH2, NUP170, NUP60, OAF1, OLA1, OM14, ORC2, PAF1, PAU8, PAU9, PBP2, PCA1, PCS60, PDB1, PDR3, PEP1, PET112, PET9, PEX22, PFF1, PHO3, PHO5, PHO89, PIM1, PIN4, PKC1, POA1, POL12, POL30, POP4, POP8, PPS1, PRE7, PRM9, PRS4, PRX1, PSY4, PTA1, PTC3, PTH2, PYC2, RBG1, RCF3, RCR1, RDH54, REI1, RER2, RFA1, RFC5, RFT1, RGD1, RIB1, RIB5, RIF1, RKM3, ROT2, ROX3, RPG1, RPL19A, RPL19B, RPL23A, RPL32, RPL4A, RPS8A, RRN10, RRN6, RRT1, RRT2, RTC2, RTG3, SAF1, SAS3, SCO2, SCS22, SCT1, SDH8, SDS24, SEA4, SEC17, SEC18, SEF1, SEN34, SEO1, SFT2, SHE1, SHG1, SHM1, SHP1, SKT5, SLA1, SLM4, SLM6, SLX1, SNF5, SNR18, SNR56, SPC72, SPO23, SPT7, SRB6, SRO77, SSA1, SSA3, SSH1, STU1, SUL1, SUP45, SWC5, SWD1, SWH1, TAE1, TAT1, TBRT, TBS1, TDP1, TEC1, TEL1, TFC3, TGA1, THI2, TIM12, TIP1, TOD6, TRM7, TRN1, TRS20, TSC10, TSC3, TYC1, UBC4, UBP13, UBP14, UBX7, UGA2, URA7, UTP20, VBA2, VHC1, VPS8, VVS1, YAT1, YBP1, YEL1, YPK3, YPT10, YRF1-6, YSW1 |
| **glu3** | AAC3, ADH5, AGP2, AMN1, APD1, APE3, APM3, ARA1, ARL1, ATG42, BMT2, BSD2, CCZ1, CDC28, CHK1, CKS1, CNS1, CSH1, CTP1, DPB3, DTR1, DUG2, DUT1, ECM31, EFM2, EHT1, EXO5, EXO84, FES1, FZO1, GDT1, HAB1, HSL7, HSM3, ICS2, IFA38, IRA1, IST2, MAK5, MAL33, MBA1, MEC1, MIS1, MMS4, MRPL27, MRPS5, MRPS9, MTC4, NHP6B, NPL4, NTC20, OPY1, PAF1, PBY1, PCA1, PCH2, PEX32, PHO3, PHO5, PHO89, POL30, POP4, POP7, PPS1, RCF3, RFC5, RIB5, RIB7, RIF1, RPB5, RPL19A, RPS6B, RTC2, RXT2, SAF1, SEC66, SEO1, SHE3, SHG1, SLI15, SMP1, SMY2, SNF5, SPO23, SPP381, SRB6, SSE2, SSH1, SUL1, SUP45, SWD3, TBS1, TEC1, TIM12, TRS20, TYC1, TYR1, UBC4, UBS1, UBX7, UMP1, VBA2, VPS15, YPC1, YSW1, YSY6 |
| **glu4** | ADE1, ADF1, ADH5, ALG7, AMN1, APD1, APE3, APM3, ARA1, ARL1, ARO4, ATG42, BMT2, BSD2, BUD14, CDC15, CDC28, CDC39, CDC50, CHA1, CHK1, CNS1, CSH1, CTP1, DFP1, DFP2, DPB3, DUG2, DUT1, EFM2, EMA35, ENP1, ERP1, EXO5, EXO84, FES1, FYV5, GEX1, GIT1, GPX2, HAB1, HIS7, HMLALPHA2, HMRA1, HMRA2, HSM3, ICS2, IFA38, IRA1, ISW1, KAR4, KIN3, KRR1, KTD1, MAK5, MAL33, MATALPHA1, MEC1, MIC10, MMS4, MRC1, MRPL27, MRPS5, MRPS9, MSH3, MST28, MTC4, NUP60, OCA4, PAF1, PAU7, PAU9, PBY1, PCA1, PEX32, PEX34, PHO89, POP4, POP7, PPS1, PRD1, PRM9, RCF3, RDT1, RFA1, RIB5, RIB7, RIF1, RPB5, RRT2, RTC2, RXT2, SAF1, SEN34, SEO1, SHG1, SLI15, SNF5, SPO23, SPP381, SRB6, SSE2, SSH1, SUL1, SUP45, SUP56, SWD1, TBS1, TFC3, TGA1, TRS20, TYC1, TYR1, UBS1, UBX7, UIP3, VAC17, VBA2, VBA3, VPS15, VPS8, YSW1, YSY6 |
| **glu5** | ADF1, CDC50, CHA1, EMA35, FYV5, GEX1, GIT1, HMLALPHA2, HMRA1, HMRA2, KAR4, KRR1, MATALPHA1, MIC10, MRC1, OCA4, PEX34, PRD1, RDT1, SEO1, VAC17, VBA3 |

| | |
|---|---|
| **glu6** | AAD10, AAD4, ACS1, ADE1, ADF1, ADH4, AGE1, AGP3, AIF1, AIM2, ALR2, ANK1, APA1, APA2, API2, AQY3, ARN1, ARN2, ATP15, AVT2, BAS1, BDH1, BDH2, BGL2, BIO2, BRR6, BSC5, BUD14, CAB1, CAB4, CAN1, CBP2, CDC15, CDC39, CDC50, CHA1, CIN8, CNE1, COS1, COS10, COS4, COS5, COS6, COS7, COS8, COS9, COX14, CSS3, CWC22, DAK2, DAL2, DAL3, DAL4, DAL5, DAL7, DAN1, DAN4, DCG1, DDI2, DFP1, DFP2, DFP4, DLD3, DOC1, DSE4, DSF1, ECM1, ECM34, EFG1, EFM1, EMA35, ERO1, ERP1, ERR2, ERR3, ERV29, ESL2, FDC1, FDH1, FEX1, FEX2, FIT1, FIT2, FIT3, FLC2, FLO10, FLO5, FPR2, FRE2, FRE3, FRE4, FRE5, FYV5, FZF1, GDH3, GEM1, GEX1, GEX2, GIT1, GPB2, GTT1, GUS1, HFM1, HIR3, HLR1, HMLALPHA2, HMRA1, HMRA2, HOM6, HPA3, HSP31, HSP32, HXK2, HXT13, HXT15, HXT17, HYR1, IMA1, IMD2, IML1, IPA1, IRC24, IRC4, IRC7, JEN1, KAP114, KAR4, KAR9, KIN3, KRE28, KRR1, KTD1, LRE1, LRG1, LYS1, MAL12, MAL13, MAL31, MAL32, MATALPHA1, MCH2, MCM22, MDL2, MET5, MGA2, MIC10, MIX23, MLP1, MNS1, MNT2, MNT4, MPH3, MRC1, MSH3, MST28, MUP3, NFT1, NMD5, NPR2, NRE1, NUP60, OAF1, OCA4, PAD1, PAU10, PAU11, PAU12, PAU13, PAU14, PAU15, PAU19, PAU21, PAU24, PAU4, PAU6, PAU7, PAU8, PBI1, PBN1, PCC1, PCK1, PDE1, PDR18, PEX22, PEX34, PGU1, PHR1, PKC1, PLC1, PMT4, POF1, PRD1, PRM9, PUL3, PUL4, PXP3, PXR1, QCR7, RAI1, RBA50, RDT1, RFA1, RMD6, RME3, RMR1, RNH70, RTF1, RTG2, RTG3, RTT102, SAM3, SAM4, SCW4, SEA4, SEN34, SEO1, SGM1, SIR1, SIT1, SKG1, SLH1, SNA2, SNO2, SNO4, SNR84, SNZ2, SOR1, SPB1, SPS1, SPS2, SPS22, SRO77, SRY1, STL1, SUP56, SUT532, SWD1, SWP82, TAD1, TAF1, TFC3, TGA1, THI11, THI13, TIM8, TTI2, UBP11, UIP3, UPA1, URA1, URC2, VAC17, VBA3, VBA5, VEL1, VMR1, VTH1, VTH2, XPT1, YOR1, YPS6, ZIP2, ZNF1, ZRT1, ZUO1 |
| **gal1** | AAC3, ACH1, ADH5, AGP2, AIM3, AIM4, AKL1, ALG1, ALG14, ALK2, AMN1, APD1, APL3, APN2, ARA1, ARL1, ATG14, ATG42, ATP3, BAP2, BEM1, BMT2, CBP6, CCZ1, CDC28, CDS1, CHS2, CHS3, CKS1, CMD1, CNM1, CNS1, COQ1, COR1, COS111, CPP1, CSG2, CSH1, CST26, CYC8, DER1, DSF2, DTR1, ECM13, ECM15, ECM2, ECM31, ECM33, ECM8, EDE1, EDS1, EHT1, ERD2, ETR1, EXO5, EXO84, FAT1, FES1, FIG1, FLR1, FMP23, FMT1, FTH1, FUI1, FUR4, FUS3, FZO1, GAL1, GAL10, GAL7, GDT1, GIP1, GPI18, GRS1, GRX7, HAP3, HEK2, HHF1, HHT1, HIR1, HMT1, HSL7, HSP26, HTA2, HTB2, ICS2, IFA38, IML3, IPP1, IRA1, IST2, KAP104, KTR3, KTR4, LAA2, LDB7, LDH1, LSM2, MAK5, MBA1, MCM2, MCM7, MEC1, MED8, MEO1, MIN6, MIN7, MIS1, MMS4, MNC1, MNN2, MOH1, MRPL16, MRPL36, MRPS9, MRX18, MSI1, MUD1, MUM2, NCL1, NHP6B, NPL4, NRG2, NTC20, NTH2, OLA1, OPY1, ORC2, PBY1, PCH2, PDR3, PDX3, PEP1, PET9, PEX32, PFF1, PGI1, PHO3, PHO5, PIM1, PIN4, POA1, POL12, POL30, POP7, POP8, PRE7, PRP6, PSY4, PTC4, QDR3, RCR1, RDH54, REB1, REG2, RER2, RFC5, RFS1, RFT1, RIB1, RIB7, RIM2, RKM3, RPB5, RPG1, RPL19A, RPL19B, RPL21A, RPL4A, RPS11B, RPS6B, RPS9B, RRN10, RRN6, RRT1, RTC2, RXT2, SAS3, SCO1, SCO2, SCT1, SEC17, SEC18, SEC66, SHE1, SHE3, SIF2, SLA1, SLI15, SLM4, SMP1, SMY2, SND3, SNR161, SPP381, SPT7, SSE2, STU1, SUP45, SUS1, SWD3, TAF5, TAT1, TBRT, TBS1, TCM62, TEC1, TFC1, TIM12, TIP1, TLC1, TPS1, TRM7, TSC3, TYR1, UBC4, UBP14, UBS1, UGA2, UMP1, URA7, UTP20, VID24, VMA2, VPS15, YMC2, YPC1, YPK3, YRO2, YSA1, YSW1, YSY6, ZTA1 |
| **gal2** | AAC3, ACH1, ADH5, AGP2, AIM3, AKL1, ALG1, ALG14, ALK2, AMN1, APD1, APL3, APN2, ARA1, ARL1, ATG14, ATG42, ATP3, BAP2, BMT2, CBP6, CCZ1, CDC28, CDS1, CHS2, CHS3, CKS1, CMD1, CNM1, CNS1, COQ1, COR1, CPP1, CSG2, CSH1, CST26, DSF2, DTR1, ECM13, ECM15, ECM2, ECM31, ECM33, ECM8, EDE1, EDS1, EHT1, ERD2, ETR1, EXO5, EXO84, FAT1, FES1, FIG1, FLR1, FMP23, FMT1, FUI1, FUR4, FUS3, FZO1, GAL1, GAL10, GAL7, GIP1, GPI18, GRS1, GRX7, HAP3, HEK2, HHF1, HHT1, HIR1, HMT1, HSL7, HSP26, HTA2, HTB2, ICS2, IFA38, IML3, IPP1, IRA1, IST2, KAP104, LAA2, LDB7, LSM2, MAK5, MBA1, MCM2, MEC1, MEO1, MIN6, MIS1, MMS4, MNC1, MNN2, MOH1, MRPL16, MRPL36, MRPS9, MRX18, MUD1, MUM2, NCL1, NHP6B, NPL4, NRG2, NTH2, OLA1, OPY1, ORC2, PBY1, PDR3, PDX3, PEP1, PET9, PEX32, PFF1, PHO3, PHO5, PIM1, PIN4, POA1, POL12, POL30, POP7, POP8, PRE7, PRP6, PSY4, PTC3, PTC4, QDR3, RCR1, RDH54, REB1, REG2, RER2, RFC5, RFS1, RFT1, RIB1, RIB7, RKM3, RPB5, RPG1, RPL19A, RPL19B, RPL4A, RPS11B, RPS6B, RRN10, RRN6, RRT1, RTC2, RXT2, SAS3, SCO1, SCO2, SCT1, SEC17, SEC18, SEC66, SHE1, SHE3, SIF2, SLA1, SLI15, SLM4, SMP1, SMY2, SND3, SNR161, SPP381, SPT7, SSE2, STU1, SUP45, SUS1, SWD3, TAT1, TBRT, TBS1, TCM62, TEC1, TFC1, TIM12, TIP1, TLC1, TOD6, TPS1, TRM7, TSC3, TYR1, UBC4, UBP14, UBS1, UGA2, UMP1, URA7, UTP20, VID24, VMA2, VPS15, YMC2, YPC1, YPK3, YRO2, YSA1, YSW1, YSY6, ZTA1 |

| | |
|---|---|
| **gal3** | AAC3, ACH1, ADH5, AGP2, AIM3, AIM4, AKL1, ALG1, ALG14, ALK2, AMN1, APD1, APL3, APN2, ARA1, ARL1, ATG14, ATG42, ATP3, BAP2, BEM1, BMT2, CBP6, CCZ1, CDC28, CDS1, CHS2, CHS3, CKS1, CMD1, CNM1, CNS1, COQ1, COR1, COS111, CPP1, CSG2, CSH1, CST26, DER1, DSF2, DTR1, ECM13, ECM15, ECM2, ECM31, ECM33, ECM8, EDE1, EDS1, EHT1, ERD2, ETR1, EXO5, EXO84, FAT1, FES1, FIG1, FLR1, FMP23, FMT1, FTH1, FUI1, FUR4, FUS3, FZO1, GAL1, GAL10, GAL7, GDT1, GIP1, GPI18, GRS1, GRX7, HAP3, HEK2, HHF1, HHT1, HIR1, HMT1, HSL7, HSP26, HTA2, HTB2, ICS2, IFA38, IML3, IPP1, IRA1, IST2, KAP104, KTR3, KTR4, LAA2, LDB7, LDH1, LSM2, MAK5, MBA1, MCM2, MCM7, MEC1, MED8, MEO1, MIN6, MIN7, MIS1, MMS4, MNC1, MNN2, MOH1, MRPL16, MRPL36, MRPS9, MRX18, MSI1, MUD1, MUM2, NCL1, NHP6B, NPL4, NRG2, NTC20, NTH2, OLA1, OPY1, ORC2, PBY1, PCH2, PDR3, PDX3, PEP1, PET9, PEX32, PFF1, PGI1, PHO3, PHO5, PIM1, PIN4, POA1, POL12, POL30, POP7, POP8, PRE7, PRP6, PSY4, PTC4, QDR3, RCR1, RDH54, REB1, REG2, RER2, RFC5, RFS1, RFT1, RIB1, RIB7, RIM2, RKM3, RPB5, RPG1, RPL19A, RPL19B, RPL21A, RPL4A, RPS11B, RPS6B, RPS9B, RRN10, RRN6, RRT1, RTC2, RXT2, SAS3, SCO1, SCO2, SCT1, SEC17, SEC18, SEC66, SHE1, SHE3, SIF2, SLA1, SLI15, SLM4, SMP1, SMY2, SND3, SNR161, SPP381, SPT7, SSE2, STU1, SUP45, SUS1, SWD3, TAF5, TAT1, TBRT, TBS1, TCM62, TEC1, TFC1, TIM12, TIP1, TLC1, TPS1, TRM7, TSC3, TYR1, UBC4, UBP14, UBS1, UGA2, UMP1, URA7, UTP20, VID24, VMA2, VPS15, YMC2, YPC1, YPK3, YRO2, YSA1, YSW1, YSY6, ZTA1 |
| **gal4** | AAC3, AAR2, ACH1, ADH5, AGP2, AIM3, AKL1, ALG1, ALG14, ALK2, AMN1, APD1, APL3, APN2, ARA1, ARL1, AST1, ATG14, ATG42, ATG8, ATP3, BAP2, BMT2, CBP6, CCZ1, CDC28, CDS1, CHS2, CHS3, CKS1, CMC2, CMD1, CNM1, CNS1, COQ1, COR1, CPP1, CSG2, CSH1, CST26, DSF2, DTR1, ECM13, ECM15, ECM2, ECM31, ECM33, ECM8, EDE1, EDS1, EHT1, ERD2, ETR1, EXO5, EXO84, FAT1, FES1, FIG1, FLR1, FMP23, FMT1, FUI1, FUR4, FUS3, FZO1, GAL1, GAL10, GAL7, GIP1, GPI18, GRS1, GRX7, HAP3, HEK2, HHF1, HHT1, HIR1, HMT1, HSL7, HSP26, HTA2, HTB2, IAI11, ICS2, IFA38, ILS1, IML3, IPP1, IRA1, IST2, KAP104, KIP1, KTI11, LAA2, LDB7, LSM2, MAK5, MBA1, MCM2, MEC1, MEO1, MIN6, MIS1, MMS4, MNC1, MNN2, MOH1, MRPL16, MRPL36, MRPS9, MRX18, MUD1, MUM2, NCL1, NHP6B, NPL4, NRG2, NTH2, OLA1, OPY1, ORC2, PBY1, PDR3, PDX3, PEP1, PET9, PEX32, PFF1, PHO3, PHO5, PIM1, PIN4, POA1, POL12, POL30, POP7, POP8, PRE7, PRP6, PRS4, PRX1, PSY4, PTC3, PTC4, PTH2, QDR3, RCR1, RDH54, REB1, REG2, RER2, RFC5, RFS1, RFT1, RIB1, RIB7, RKM3, RPB5, RPG1, RPL19A, RPL19B, RPL4A, RPS11B, RPS6B, RPS8A, RRN10, RRN6, RRT1, RTC2, RXT2, SAS3, SCO1, SCO2, SCT1, SEC17, SEC18, SEC66, SEF1, SHE1, SHE3, SHP1, SIF2, SKT5, SLA1, SLI15, SLM4, SMP1, SMY2, SND3, SNR161, SNR56, SPP381, SPT7, SSA3, SSE2, STU1, SUP45, SUS1, SWD3, TAT1, TBRT, TBS1, TCM62, TEC1, TFC1, TIM12, TIP1, TLC1, TOD6, TPS1, TRM7, TSC3, TYR1, UBC4, UBP13, UBP14, UBS1, UGA2, UMP1, URA7, UTP20, VID24, VMA2, VPS15, YEL1, YMC2, YPC1, YPK3, YRO2, YSA1, YSW1, YSY6, ZTA1 |
| **gal5** | AAC3, ACH1, ADH5, AGP2, AIM3, AIM4, AKL1, ALG1, ALG14, ALK2, AMN1, APD1, APL3, APN2, ARA1, ARL1, ATG14, ATG42, ATP3, BAP2, BEM1, BMT2, CBP6, CCZ1, CDC28, CDS1, CHS2, CHS3, CKS1, CMD1, CNM1, CNS1, COQ1, COR1, COS111, CPP1, CSG2, CSH1, CST26, CYC8, DER1, DSF2, DTR1, ECM13, ECM15, ECM2, ECM31, ECM33, ECM8, EDE1, EDS1, EHT1, ERD2, ETR1, EXO5, EXO84, FAT1, FES1, FIG1, FLR1, FMP23, FMT1, FTH1, FUI1, FUR4, FUS3, FZO1, GAL1, GAL10, GAL7, GDT1, GIP1, GPI18, GRS1, GRX7, HAP3, HEK2, HHF1, HHT1, HIR1, HMT1, HSL7, HSP26, HTA2, HTB2, ICS2, IFA38, IML3, IPP1, IRA1, IST2, KAP104, KTR3, KTR4, LAA2, LDB7, LDH1, LSM2, MAK5, MBA1, MCM2, MCM7, MEC1, MED8, MEO1, MIN6, MIN7, MIS1, MMS4, MNC1, MNN2, MOH1, MRPL16, MRPL36, MRPS9, MRX18, MSI1, MUD1, MUM2, NCL1, NHP6B, NPL4, NRG2, NTC20, NTH2, OLA1, OPY1, ORC2, PBY1, PCH2, PDR3, PDX3, PEP1, PET9, PEX32, PFF1, PGI1, PHO3, PHO5, PIM1, PIN4, POA1, POL12, POL30, POP7, POP8, PRE7, PRP6, PSY4, PTC3, PTC4, QDR3, RCR1, RDH54, REB1, REG2, RER2, RFC5, RFS1, RFT1, RIB1, RIB7, RIM2, RKM3, RPB5, RPG1, RPL19A, RPL19B, RPL21A, RPL4A, RPS11B, RPS6B, RPS9B, RRN10, RRN6, RRT1, RTC2, RXT2, SAS3, SCO1, SCO2, SCT1, SEC17, SEC18, SEC66, SHE1, SHE3, SIF2, SLA1, SLI15, SLM4, SMP1, SMY2, SND3, SNR161, SPP381, SPT7, SSE2, STU1, SUP45, SUS1, SWD3, TAF5, TAT1, TBRT, TBS1, TCM62, TEC1, TFC1, TIM12, TIP1, TLC1, TOD6, TPS1, TRM7, TSC3, TYR1, UBC4, UBP14, UBS1, UGA2, UMP1, URA7, UTP20, VID24, VMA2, VPS15, YMC2, YPC1, YPK3, YRO2, YSA1, YSW1, YSY6, ZTA1 |

| | |
|---|---|
| gal6 | AAC3, ACH1, ADH5, AGP2, AIM3, AIM4, AKL1, ALG1, ALG14, ALK2, AMN1, APD1, APL3, APN2, ARA1, ARL1, ATG14, ATG42, ATP3, BAP2, BEM1, BMT2, CBP6, CCZ1, CDC28, CDS1, CHS2, CHS3, CKS1, CMD1, CNM1, CNS1, COQ1, COR1, COS111, CPP1, CSG2, CSH1, CST26, CYC8, DER1, DSF2, DTR1, ECM13, ECM15, ECM2, ECM31, ECM33, ECM8, EDE1, EDS1, EHT1, ERD2, ETR1, EXO5, EXO84, FAT1, FES1, FIG1, FLR1, FMP23, FMT1, FTH1, FUI1, FUR4, FUS3, FZO1, GAL1, GAL10, GAL7, GDT1, GIP1, GPI18, GRS1, GRX7, HAP3, HEK2, HHF1, HHT1, HIR1, HMT1, HSL7, HSP26, HTA2, HTB2, ICS2, IFA38, IML3, IPP1, IRA1, IST2, KAP104, KTR3, KTR4, LAA2, LDB7, LDH1, LSM2, MAK5, MBA1, MCM2, MCM7, MEC1, MED8, MEO1, MIN6, MIN7, MIS1, MMS4, MNC1, MNN2, MOH1, MRPL16, MRPL36, MRPS9, MRX18, MSI1, MUD1, MUM2, NCL1, NHP6B, NPL4, NRG2, NTC20, NTH2, OLA1, OPY1, ORC2, PBY1, PCH2, PDR3, PDX3, PEP1, PET9, PEX32, PFF1, PGI1, PHO3, PHO5, PIM1, PIN4, POA1, POL12, POL30, POP7, POP8, PRE7, PRP6, PSY4, PTC3, PTC4, QDR3, RCR1, RDH54, REB1, REG2, RER2, RFC5, RFS1, RFT1, RIB1, RIB7, RIM2, RKM3, RPB5, RPG1, RPL19A, RPL19B, RPL21A, RPL4A, RPS11B, RPS6B, RPS9B, RRN10, RRN6, RRT1, RTC2, RXT2, SAS3, SCO1, SCO2, SCT1, SEC17, SEC18, SEC66, SHE1, SHE3, SIF2, SLA1, SLI15, SLM4, SMP1, SMY2, SND3, SNR161, SPP381, SPT7, SSE2, STU1, SUP45, SUS1, SWD3, TAF5, TAT1, TBRT, TBS1, TCM62, TEC1, TFC1, TIM12, TIP1, TLC1, TOD6, TPS1, TRM7, TSC3, TYR1, UBC4, UBP14, UBS1, UGA2, UMP1, URA7, UTP20, VID24, VMA2, VPS15, YMC2, YPC1, YPK3, YRO2, YSA1, YSW1, YSY6, ZTA1 |

**1200 generations**

| Replicate | Gene list |
|---|---|
| glu1 | AAC3, AAR2, ACH1, ADH5, AKL1, ALG3, ALK2, AMN1, APD1, APE3, APL3, APM3, APN2, ARA1, ARL1, ARO4, AST1, ATG42, ATG8, ATP1, ATP3, AVT5, BMT2, BNA4, BOI1, BRN1, BSD2, CDC27, CDC28, CHK1, CHS3, CMC2, CNM1, CNS1, COQ1, COR1, CPP1, CSH1, CST26, CTP1, DPB3, DSF2, DUG2, DUT1, ECM13, ECM15, ECM2, ECM21, ECM33, ECM8, EDE1, EFM2, ERD2, ETR1, EXO5, FAT1, FIG1, FLR1, FMP23, FMT1, FUI1, FUR4, FUS3, GAL1, GAL10, GAL7, GIP1, GPI18, GRX7, HAB1, HAP3, HEK2, HHF1, HHT1, HIR1, HSM3, HTA2, HTB2, IAI11, ICS2, IFA38, ILS1, IMD2, IPP1, IRA1, IST2, KAP104, KIP1, KTI11, LAA2, LDB7, LSM2, MAK5, MAL33, MAP2, MCM2, MIN6, MIS1, MIX23, MNC1, MNN2, MOH1, MRP21, MRPL16, MRPL27, MRPS5, MRPS9, MRX18, MRX3, MTC4, MUM2, NCL1, NPL4, NRG2, NTH2, NUP170, OLA1, ORC2, PAF1, PCA1, PDR3, PEP1, PET112, PET9, PEX32, PFF1, PHO89, PIM1, PIN4, PKC1, POA1, POL12, POL30, POP4, POP7, POP8, PPS1, PRE7, PRP6, PRS4, PRX1, PSY4, PTC3, PTH2, QDR3, RCF3, RCR1, REB1, REG2, RER2, RFC5, RFS1, RFT1, RGD1, RIB1, RIB5, RIB7, RIF1, ROX3, RPB5, RPG1, RPL19A, RPL19B, RPL23A, RPL32, RPS11B, RPS8A, RRN10, RRN6, RRT1, RTC2, RTG3, SAF1, SAS3, SCO2, SCS22, SCT1, SEA4, SEC17, SEC18, SEC66, SEF1, SEO1, SFT2, SHE1, SHG1, SHP1, SKT5, SLA1, SLI15, SLM4, SNF5, SNR56, SPO23, SPP381, SPT7, SRB6, SRO77, SSA3, SSE2, SSH1, STU1, SUL1, SUP45, TBS1, TCM62, TEC1, TEL1, TOD6, TRM7, TRS20, TSC3, TYC1, TYR1, UBC4, UBP13, UBP14, UBS1, UBX7, UGA2, URA7, UTP20, VBA2, YEL1, YPK3, YRO2, YSW1, YSY6, ZTA1 |
| glu2 | AAD15, AAR2, ACH1, ACS1, ADE1, ADF1, ADH5, ADH7, AIM2, ALG3, ALK2, APE3, APL3, APM3, APN2, ARA1, ARO4, AST1, ATG42, ATG8, ATP1, ATP3, AVT5, BDH1, BDH2, BMT2, BNA4, BOI1, BRN1, BSD2, BUD14, CDC15, CDC27, CDC50, CHA1, CHK1, CMC2, CNE1, COR1, CST26, CTP1, DFP1, DFP2, DPB3, DUG2, DUT1, ECM1, ECM13, ECM21, EDE1, EFM2, EMA35, ENP1, ERD2, ERP1, FAT1, FIG1, FLC2, FLO5, FMT1, FUI1, FUS3, FYV5, GDH3, GEM1, GEX1, GIP1, GIT1, GPB2, GPX2, HAB1, HAP3, HEK2, HIR1, HIS7, HMLALPHA2, HMRA1, HMRA2, HSM3, IAI11, ILS1, IMD2, IRA1, ISW1, KAR4, KIN3, KIP1, KRR1, KTD1, KTI11, LAA2, LDB7, LSM2, MAK5, MAL33, MAP2, MATALPHA1, MCM2, MEC1, MIC10, MIN6, MIX23, MOH1, MRC1, MRP21, MRPL16, MRPL27, MRPS5, MRPS9, MRX3, MST28, MTC4, NCL1, NUP170, NUP60, OAF1, OCA4, PAF1, PAU3, PAU7, PAU8, PAU9, PCA1, PDR3, PEP1, PET112, PET9, PEX22, PEX34, PHO89, PIM1, PIN4, PKC1, POL12, POP4, POP8, PPS1, PRD1, PRE7, PRM9, PRS4, PRX1, PSY4, PTC3, PTH2, QDR3, RCF3, RDS1, RDT1, RFA1, RFT1, RIB1, RIB5, RIF1, RMD6, ROX3, RPL19B, RPL23A, RPL32, RPS8A, RRN10, RRN6, RRT1, RRT2, RTC2, RTG3, SAF1, SAS3, SCS22, SCT1, SEA4, SEC17, SEF1, SEN34, SEO1, SFT2, SHE1, SHG1, SHP1, SKT5, SLA1, SNF5, SNR56, SPO23, SRB6, SRO77, SSA3, SSH1, STU1, SUL1, SUP45, SUP56, SWD1, SWH1, TBS1, TCM62, TEL1, TFC3, TGA1, TOD6, TRS20, TYC1, UBP13, UBX7, UIP3, URA7, UTP20, VAC17, VBA2, VBA3, VPS8, YAT1, YEL1, YRF1-6, YSW1 |

| | |
|---|---|
| **glu3** | ABD1, ACS1, ADE1, ADF1, ADH5, AGP2, AIM2, ALG7, AME1, AMN1, APD1, APE3, APM3, ARA1, ARC40, ARL1, ARO4, ATG12, ATG14, ATG42, BDH1, BDH2, BIT2, BMT2, BSD2, BUD14, CBP6, CCZ1, CDC15, CDC28, CDC39, CDC50, CHA1, CHK1, CKS1, CNE1, CNS1, COQ21, CSH1, CSS1, CTP1, DAD3, DFP1, DPB3, DUG2, DUR12, DUT1, ECM1, EFM2, EMA35, ENP1, ERP1, ERT1, ERV15, EXO5, EXO84, FES1, FLC2, FLO1, FLO5, FRE2, FTH1, FYV5, GDH3, GEM1, GEX1, GIT1, GPB2, GPX2, GRS1, HAB1, HIS7, HMLALPHA2, HMRA1, HMRA2, HPC2, HSL7, HSM3, HXT9, ICS2, IFA38, IMA4, IMD2, IRA1, ISW1, KAR4, KIN3, KRR1, KTR3, LDH1, LSR1, MAK5, MAL33, MATALPHA1, MCH2, MCX1, MEC1, MEO1, MET8, MIC10, MIC12, MMS4, MRC1, MRPL27, MRPL36, MRPL37, MRPS5, MRPS9, MSH3, MTC4, MUD1, NGR1, NPL4, NUP60, OAF1, OCA4, OM14, OPY1, PAF1, PAU10, PAU15, PAU18, PAU20, PAU7, PAU8, PAU9, PBP2, PBY1, PCA1, PCS60, PDB1, PEX22, PEX32, PEX34, PHO89, POP4, POP7, PPS1, PRD1, PTC4, PYC2, RCF3, RDT1, REI1, RFA1, RGD1, RIB5, RIB7, RIF1, ROT2, RPB5, RRT2, RTC2, RXT2, SAF1, SDH8, SDS24, SEC66, SEN34, SEO1, SHE3, SHG1, SHM1, SLI15, SLM6, SLX1, SNF5, SPO23, SPP381, SRB6, SSE2, SSH1, SUL1, SUP45, SWC5, SWD1, TAE1, TBS1, TDP1, TFC1, TFC3, TGA1, THI2, TPS1, TRS20, TSC10, TYC1, TYR1, UBS1, UBX7, VAC17, VBA2, VBA3, VHC1, VMA2, VPS15, VPS8, VTH1, VVS1, YBP1, YPT10, YSW1, YSY6 |
| **glu4** | AAC3, AAR2, ACH1, ADH5, AIM3, AKL1, ALG1, ALG14, ALG3, ALK2, APL3, APN2, ARA1, AST1, ATG42, ATG8, ATP1, ATP3, AVT5, BAP2, BMT2, BNA4, BOI1, BRN1, CBP6, CDC27, CHS3, CMC2, CMD1, CNM1, COQ1, COR1, CPP1, CST26, CYC8, DSF2, ECM13, ECM15, ECM2, ECM21, ECM33, ECM8, EDE1, ERD2, ETR1, EXO84, FAT1, FES1, FIG1, FLR1, FMT1, FUI1, FUR4, FUS3, GAL1, GAL10, GAL7, GIP1, GPI18, GRS1, GRX7, HAP3, HEK2, HHF1, HHT1, HIR1, HSP26, HTA2, HTB2, IAI11, ILS1, IML3, IPP1, IRA1, IST2, KAP104, KIP1, KTI11, LAA2, LDB7, LSM2, MAK5, MAP2, MCM2, MEC1, MIN6, MIS1, MIX23, MMS4, MNC1, MNN2, MOH1, MRP21, MRPL16, MRPL36, MRPS9, MRX3, MUD1, MUM2, NCL1, NHP6B, NRG2, NTH2, NUP170, OLA1, ORC2, PBY1, PDR3, PEP1, PET112, PET9, PFF1, PHO3, PHO5, PIM1, PIN4, PKC1, POA1, POL12, POL30, POP8, PRE7, PRS4, PRX1, PSY4, PTC3, PTC4, PTH2, QDR3, RCR1, RDH54, RER2, RFC5, RFT1, RIB1, ROX3, RPG1, RPL19A, RPL19B, RPL23A, RPL32, RPS8A, RRN10, RRN6, RRT1, RTC2, RTG3, RXT2, SAS3, SCO2, SCS22, SCT1, SEA4, SEC17, SEC18, SEF1, SFT2, SHE1, SHP1, SIF2, SKT5, SLA1, SLM4, SND3, SNR56, SPT7, SRO77, SSA3, STU1, SUP45, SUS1, TAT1, TBRT, TBS1, TCM62, TEC1, TEF2, TEL1, TFC1, TIM12, TIP1, TOD6, TRM7, TSC3, UBC4, UBP13, UBP14, UGA2, URA7, UTP20, VID24, VPS15, YEL1, YMC2, YSA1, YSW1 |
| **glu5** | AAR2, ACH1, ADF1, ADH5, AKL1, ALG3, ALG7, ALK2, AMN1, APD1, APE3, APL3, APM3, APN2, ARA1, ARL1, ARO4, AST1, ATG42, ATG8, ATP1, ATP3, AVT5, BMT2, BNA4, BOI1, BRN1, BSD2, CDC27, CDC28, CDC50, CDS1, CHA1, CHK1, CHS3, CKS1, CMC2, CNM1, CNS1, COQ1, COR1, CPP1, CSH1, CST26, CTP1, DPB3, DSF2, DUG2, DUT1, ECM13, ECM15, ECM2, ECM21, EDE1, EFM2, ENP1, ERD2, ETR1, EXO5, FAT1, FIG1, FLO5, FLR1, FMT1, FUI1, FUR4, FUS3, FYV5, GAL1, GAL10, GAL7, GEX1, GIP1, GPI18, GPX2, GRX7, HAB1, HAP3, HEK2, HHF1, HHT1, HIR1, HIS7, HMLALPHA2, HMRA1, HMRA2, HSL7, HSM3, HTA2, HTB2, IAI11, ICS2, IFA38, ILS1, IMD2, IPP1, IRA1, ISW1, KAP104, KAR4, KIP1, KRR1, KTI11, LAA2, LDB7, LSM2, MAK5, MAL33, MAP2, MATALPHA1, MCM2, MEC1, MIC10, MIN6, MIX23, MNC1, MNN2, MOH1, MRC1, MRP21, MRPL16, MRPL27, MRPS5, MRPS9, MRX18, MRX3, MTC4, MUM2, NCL1, NTH2, NUP170, OCA4, OLA1, ORC2, PAF1, PAU12, PAU8, PAU9, PCA1, PDR3, PEP1, PET112, PET9, PEX32, PEX34, PHO89, PIM1, PIN4, PKC1, POA1, POL12, POP4, POP7, POP8, PPS1, PRD1, PRE7, PRP6, PRS4, PRX1, PSY4, PTC3, PTH2, QDR3, RCF3, RCR1, RDT1, RER2, RFT1, RGD1, RIB1, RIB5, RIB7, RIF1, RMD6, ROX3, RPB5, RPL19B, RPL23A, RPL32, RPS8A, RRN10, RRN6, RRT1, RRT2, RTC2, RTG3, SAF1, SAS3, SCO2, SCS22, SCT1, SEA4, SEC17, SEF1, SEO1, SFT2, SHE1, SHG1, SHP1, SKT5, SLA1, SLI15, SNF5, SNR56, SPO23, SPP381, SRB6, SRO77, SSA3, SSH1, STU1, SUL1, SUP45, TBS1, TCM62, TEL1, TOD6, TRM7, TRS20, TSC3, TYC1, TYR1, UBP13, UBP14, UBS1, UBX7, UGA2, URA7, UTP20, VAC17, VBA2, VBA3, YEL1, YPK3, YSW1, YSY6 |

| | |
|---|---|
| **glu6** | AAD10, ABP1, ACS1, ADE1, ADF1, ADH4, AGP3, AHC2, AIF1, AIM2, AMF1, APA1, AQY3, ARN1, ARN2, ATF1, ATS1, AYT1, BAS1, BAT2, BDH1, BDH2, BGL2, BIO2, BOL1, BOL3, BSC5, BUD14, CAB4, CCR4, CDC15, CDC19, CDC24, CDC39, CDC50, CHA1, CLN3, CNE1, COS1, COS10, COS12, COS4, COS5, COS6, COS7, COS8, COX14, CSM1, CWC22, CYC3, CYS3, DAK2, DAL1, DAL2, DAL3, DAL4, DAL5, DAL7, DAL81, DAN1, DAN4, DCG1, DDI2, DEP1, DFP1, DFP2, DFP4, DLD3, DRS2, DSE4, ECM1, ECM34, EFB1, EFG1, EMA35, EMC1, ERG13, ERO1, ERP1, ERP2, ERR2, ERR3, ERV29, ERV46, FDC1, FDH1, FEX1, FEX2, FIG2, FIT1, FIT2, FIT3, FLC2, FLO10, FLO11, FLO5, FRE2, FRE3, FRE4, FRE5, FRT2, FUN12, FUN14, FUN19, FUN26, FUN30, FYV5, FZF1, GCV3, GDH1, GDH3, GEM1, GEX1, GEX2, GIP4, GIT1, GPB2, GTR1, GTT1, GTT2, HFM1, HIR3, HMLALPHA2, HMRA1, HMRA2, HMS2, HOM6, HRA1, HSP32, HSU1, HXK2, HXT16, HXT17, HYR1, ICR1, IMA1, IMA3, IMD2, IML1, INA22, IPA1, IRC24, IRC4, IRC7, JLP1, KAR4, KIN3, KIN82, KRR1, KTD1, LDS1, LRE1, LTE1, LYS1, MAK16, MAL12, MAL13, MAL31, MAL32, MAN2, MATALPHA1, MCH2, MCM22, MDM10, MET5, MGA2, MGM101, MGR1, MHT1, MIC10, MMP1, MND2, MNS1, MNT2, MPH3, MRC1, MRS1, MSC1, MSH3, MST28, MTW1, MYO4, NDI1, NFT1, NMD5, NRE1, NTG1, NUP60, OAF1, OCA4, PAD1, PAU1, PAU10, PAU11, PAU12, PAU13, PAU14, PAU15, PAU19, PAU21, PAU24, PAU4, PAU6, PAU7, PAU8, PBN1, PCK1, PDR18, PEX22, PEX34, PGA3, PGU1, PHO11, PHO12, PHO84, PHR1, PMT2, PMT4, POF1, POP5, PRD1, PRM9, PRP45, PSK1, PTA1, PUL3, PUL4, PWR1, PXP3, PXR1, RBG1, RDR1, RDT1, RFA1, RMD6, RME3, RMR1, RNH70, RPS4A, RSC9, RTG2, RTT102, SAM3, SAW1, SCW4, SEC11, SEN34, SEO1, SGM1, SIR1, SKG1, SNC1, SNO2, SNO4, SNR18, SNZ2, SOR1, SPB1, SPC72, SPO7, SPS22, SRB8, SSA1, STL1, SUP56, SUT532, SWC3, SWD1, SWH1, SYN8, TAF1, TFC3, TGA1, THI11, TIM8, TPD3, TRN1, TRX3, TTI2, TUB3, TUP1, UBP11, UIP3, UPA1, VAC17, VBA3, VBA5, VEL1, VPS8, VTH2, XPT1, YAT1, YCT1, YOR1, YPS6, YRF1-2, YRF1-3, YRF1-4, YRF1-5, YRF1-6, YRF1-7, YRF1-8, YVH1, ZIP2, ZNF1, ZRT1, ZUO1 |
| **gal1** | AAC3, ACH1, ADH5, AGP2, AIM3, AIM4, AKL1, ALG1, ALG14, ALK2, AMN1, APD1, APL3, APN2, ARA1, ARL1, ATG14, ATG42, ATP3, BAP2, BEM1, BMT2, CBP6, CCZ1, CDC28, CDS1, CHS2, CHS3, CKS1, CMD1, CNM1, CNS1, COQ1, COR1, COS111, CPP1, CSG2, CSH1, CST26, CYC8, DER1, DSF2, DTR1, ECM13, ECM15, ECM2, ECM31, ECM33, ECM8, EDE1, EDS1, EHT1, ERD2, ETR1, EXO5, EXO84, FAT1, FES1, FIG1, FLR1, FMP23, FMT1, FTH1, FUI1, FUR4, FUS3, FZO1, GAL1, GAL10, GAL7, GDT1, GIP1, GPI18, GRS1, GRX7, HAP3, HEK2, HHF1, HHT1, HIR1, HMT1, HSL7, HSP26, HTA2, HTB2, ICS2, IFA38, IML3, IPP1, IRA1, IST2, KAP104, KTR3, KTR4, LAA2, LDB7, LDH1, LSM2, MAK5, MBA1, MCM2, MCM7, MEC1, MED8, MEO1, MIN6, MIN7, MIS1, MMS4, MNC1, MNN2, MOH1, MRPL16, MRPL36, MRPS9, MRX18, MSI1, MUD1, MUM2, NCL1, NHP6B, NPL4, NRG2, NTC20, NTH2, OLA1, OPY1, ORC2, PBY1, PCH2, PDR3, PDX3, PEP1, PET9, PEX32, PFF1, PGI1, PHO3, PHO5, PIM1, PIN4, POA1, POL12, POL30, POP7, POP8, PRE7, PRP6, PSY4, PTC4, QDR3, RCR1, RDH54, REB1, REG2, RER2, RFC5, RFS1, RFT1, RIB1, RIB7, RIM2, RKM3, RPB5, RPG1, RPL19A, RPL19B, RPL21A, RPL4A, RPS11B, RPS6B, RPS9B, RRN10, RRN6, RRT1, RTC2, RXT2, SAS3, SCO1, SCO2, SCT1, SEC17, SEC18, SEC66, SHE1, SHE3, SIF2, SLA1, SLI15, SLM4, SMP1, SMY2, SND3, SNR161, SPP381, SPT7, SSE2, STU1, SUP45, SUS1, SWD3, TAF5, TAT1, TBRT, TBS1, TCM62, TEC1, TEF2, TFC1, TIM12, TIP1, TKL2, TLC1, TPS1, TRM7, TSC3, TYR1, UBC4, UBP14, UBS1, UGA2, UMP1, URA7, UTP20, VID24, VMA2, VPS15, YMC2, YPC1, YPK3, YRO2, YSA1, YSW1, YSY6, ZTA1 |
| **gal2** | AAC3, ACH1, ADH5, AGP2, AIM3, AKL1, ALG1, ALG14, ALK2, AMN1, APD1, APL3, APN2, ARA1, ARL1, ATG14, ATG42, ATP3, BAP2, BMT2, CBP6, CCZ1, CDC28, CDS1, CHS2, CHS3, CKS1, CMD1, CNM1, CNS1, COQ1, COR1, CPP1, CSG2, CSH1, CST26, CYC8, DSF2, DTR1, ECM13, ECM15, ECM2, ECM31, ECM33, ECM8, EDE1, EDS1, EHT1, ERD2, ETR1, EXO5, EXO84, FAT1, FES1, FIG1, FLR1, FMP23, FMT1, FUI1, FUR4, FUS3, FZO1, GAL1, GAL10, GAL7, GDT1, GIP1, GPI18, GRS1, GRX7, HAP3, HEK2, HHF1, HHT1, HIR1, HMT1, HSL7, HSP26, HTA2, HTB2, ICS2, IFA38, IML3, IPP1, IRA1, IST2, KAP104, LAA2, LDB7, LSM2, MAK5, MBA1, MCM2, MEC1, MEO1, MIN6, MIS1, MMS4, MNC1, MNN2, MOH1, MRPL16, MRPL36, MRPS9, MRX18, MUD1, MUM2, NCL1, NHP6B, NPL4, NRG2, NTC20, NTH2, OLA1, OPY1, ORC2, PBY1, PCH2, PDR3, PDX3, PEP1, PET9, PEX32, PFF1, PHO3, PHO5, PIM1, PIN4, POA1, POL12, POL30, POP7, POP8, PRE7, PRP6, PSY4, PTC3, PTC4, PTH2, QDR3, RCR1, RDH54, REB1, REG2, RER2, RFC5, RFS1, RFT1, RIB1, RIB7, RKM3, RPB5, RPG1, RPL19A, RPL19B, RPL21A, RPL4A, RPS11B, RPS6B, RPS9B, RRN10, RRN6, RRT1, RTC2, RXT2, SAS3, SCO1, SCO2, SCT1, SEC17, SEC18, SEC66, SHE1, SHE3, SHP1, SIF2, SLA1, SLI15, SLM4, SMP1, SMY2, SND3, SNR161, SPP381, SPT7, SSE2, STU1, SUP45, SUS1, SWD3, TAT1, TBRT, TBS1, TCM62, TEC1, TEF2, TFC1, TIM12, TIP1, TKL2, TLC1, TOD6, TPS1, TRM7, TSC3, TYR1, UBC4, UBP14, UBS1, UGA2, UMP1, URA7, UTP20, VID24, VMA2, VPS15, YMC2, YPC1, YPK3, YRO2, YSA1, YSW1, YSY6, ZTA1 |

| | |
|---|---|
| **gal3** | AAC3, ACH1, ADH5, AGP2, AIM3, AIM4, AKL1, ALG1, ALG14, ALK2, AMN1, APD1, APL3, APN2, ARA1, ARL1, ATG14, ATG42, ATP3, BAP2, BEM1, BMT2, CBP6, CCZ1, CDC28, CDS1, CHS2, CHS3, CKS1, CMD1, CNM1, CNS1, COQ1, COR1, COS111, CPP1, CSG2, CSH1, CST26, CYC8, DER1, DSF2, DTR1, ECM13, ECM15, ECM2, ECM31, ECM33, ECM8, EDE1, EDS1, EHT1, ERD2, ETR1, EXO5, EXO84, FAT1, FES1, FIG1, FLR1, FMP23, FMT1, FTH1, FUI1, FUR4, FUS3, FZO1, GAL1, GAL10, GAL7, GDT1, GIP1, GPI18, GRS1, GRX7, HAP3, HEK2, HHF1, HHT1, HIR1, HMT1, HSL7, HSP26, HTA2, HTB2, ICS2, IFA38, IML3, IPP1, IRA1, IST2, KAP104, KTR3, KTR4, LAA2, LDB7, LDH1, LSM2, MAK5, MBA1, MCM2, MCM7, MEC1, MED8, MEO1, MIN6, MIN7, MIS1, MMS4, MNC1, MNN2, MOH1, MRPL16, MRPL36, MRPS9, MRX18, MSI1, MUD1, MUM2, NCL1, NHP6B, NPL4, NRG2, NTC20, NTH2, OLA1, OPY1, ORC2, PBY1, PCH2, PDR3, PDX3, PEP1, PET9, PEX32, PFF1, PGI1, PHO3, PHO5, PIM1, PIN4, POA1, POL12, POL30, POP7, POP8, PRE7, PRP6, PSY4, PTC4, QDR3, RCR1, RDH54, REB1, REG2, RER2, RFC5, RFS1, RFT1, RIB1, RIB7, RIM2, RKM3, RPB5, RPG1, RPL19A, RPL19B, RPL21A, RPL4A, RPS11B, RPS6B, RPS9B, RRN10, RRN6, RRT1, RTC2, RXT2, SCO1, SCO2, SCT1, SEC17, SEC18, SEC66, SHE1, SHE3, SIF2, SLA1, SLI15, SLM4, SMP1, SMY2, SND3, SNR161, SPP381, SPT7, SSE2, STU1, SUP45, SUS1, SWD3, TAF5, TAT1, TBRT, TBS1, TCM62, TEC1, TEF2, TFC1, TIM12, TIP1, TKL2, TLC1, TPS1, TRM7, TSC3, TYR1, UBC4, UBP14, UBS1, UGA2, UMP1, URA7, UTP20, VID24, VMA2, VPS15, YMC2, YPC1, YPK3, YRO2, YSA1, YSW1, YSY6, ZTA1 |
| **gal4** | AAC3, ACH1, ADH5, AGP2, AIM3, AIM4, AKL1, ALG1, ALG14, ALK2, AMN1, APD1, APL3, APN2, ARA1, ARL1, ATG14, ATG42, ATP3, BAP2, BEM1, BMT2, CBP6, CCZ1, CDC28, CDS1, CHS2, CHS3, CKS1, CMD1, CNM1, CNS1, COQ1, COR1, COS111, CPP1, CSG2, CSH1, CST26, CYC8, DER1, DSF2, DTR1, ECM13, ECM15, ECM2, ECM31, ECM33, ECM8, EDE1, EDS1, EHT1, ERD2, ETR1, EXO5, EXO84, FAT1, FES1, FIG1, FLR1, FMP23, FMT1, FTH1, FUI1, FUR4, FUS3, FZO1, GAL1, GAL10, GAL7, GDT1, GIP1, GPI18, GRS1, GRX7, HAP3, HEK2, HHF1, HHT1, HIR1, HMT1, HSL7, HSP26, HTA2, HTB2, IAI11, ICS2, IFA38, IML3, IPP1, IRA1, IST2, KAP104, KTR3, KTR4, LAA2, LDB7, LDH1, LSM2, MAK5, MBA1, MCM2, MCM7, MEC1, MED8, MEO1, MIN6, MIN7, MIS1, MMS4, MNC1, MNN2, MOH1, MRPL16, MRPL36, MRPS9, MRX18, MSI1, MUD1, MUM2, NCL1, NHP6B, NPL4, NRG2, NTC20, NTH2, OLA1, OPY1, ORC2, PBY1, PCH2, PDR3, PDX3, PEP1, PET9, PEX32, PFF1, PGI1, PHO3, PHO5, PIM1, PIN4, POA1, POL12, POL30, POP7, POP8, PRE7, PRP6, PSY4, PTC3, PTC4, PTH2, QDR3, RCR1, RDH54, REB1, REG2, RER2, RFC5, RFS1, RFT1, RIB1, RIB7, RIM2, RKM3, RPB5, RPG1, RPL19A, RPL19B, RPL21A, RPL4A, RPS11B, RPS6B, RPS9B, RRN10, RRN6, RRT1, RTC2, RXT2, SAS3, SCO1, SCO2, SCT1, SEC17, SEC18, SEC66, SHE1, SHE3, SHP1, SIF2, SLA1, SLI15, SLM4, SMP1, SMY2, SND3, SNR161, SPP381, SPT7, SSE2, STU1, SUP45, SUS1, SWD3, TAF5, TAT1, TBRT, TBS1, TCM62, TEC1, TEF2, TFC1, TIM12, TIP1, TKL2, TLC1, TOD6, TPS1, TRM7, TSC3, TYR1, UBC4, UBP14, UBS1, UGA2, UMP1, URA7, UTP20, VID24, VMA2, VPS15, YMC2, YPC1, YPK3, YRO2, YSA1, YSW1, YSY6, ZTA1 |
| **gal5** | AAC3, ACH1, ADH5, AGP2, AIM3, AKL1, ALG1, ALG14, ALK2, AMN1, APD1, APL3, APN2, ARA1, ARL1, ATG14, ATG42, ATP3, BAP2, BMT2, CBP6, CCZ1, CDC28, CDS1, CHS2, CHS3, CKS1, CMD1, CNM1, CNS1, COQ1, COR1, CPP1, CSG2, CSH1, CST26, CYC8, DSF2, DTR1, ECM13, ECM15, ECM2, ECM31, ECM33, ECM8, EDE1, EDS1, EHT1, ERD2, ETR1, EXO5, EXO84, FAT1, FES1, FIG1, FLR1, FMP23, FMT1, FUI1, FUR4, FUS3, FZO1, GAL1, GAL10, GAL7, GDT1, GIP1, GPI18, GRS1, GRX7, HAP3, HEK2, HHF1, HHT1, HIR1, HMT1, HSL7, HSP26, HTA2, HTB2, ICS2, IFA38, IML3, IPP1, IRA1, IST2, KAP104, LAA2, LDB7, LSM2, MAK5, MBA1, MCM2, MEC1, MEO1, MIN6, MIS1, MMS4, MNC1, MNN2, MOH1, MRPL16, MRPL36, MRPS9, MRX18, MUD1, MUM2, NCL1, NHP6B, NPL4, NRG2, NTC20, NTH2, OLA1, OPY1, ORC2, PBY1, PCH2, PDR3, PDX3, PEP1, PET9, PEX32, PFF1, PHO3, PHO5, PIM1, PIN4, POA1, POL12, POL30, POP7, POP8, PRE7, PRP6, PSY4, PTC4, QDR3, RCR1, RDH54, REB1, REG2, RER2, RFC5, RFS1, RFT1, RIB1, RIB7, RIM2, RKM3, RPB5, RPG1, RPL19A, RPL19B, RPL21A, RPL4A, RPS11B, RPS6B, RPS9B, RRN10, RRN6, RRT1, RTC2, RXT2, SAS3, SCO1, SCO2, SCT1, SEC17, SEC18, SEC66, SHE1, SHE3, SIF2, SLA1, SLI15, SLM4, SMP1, SMY2, SND3, SNR161, SPP381, SPT7, SSE2, STU1, SUP45, SUS1, SWD3, TAT1, TBRT, TBS1, TCM62, TEC1, TEF2, TFC1, TIM12, TIP1, TKL2, TLC1, TPS1, TRM7, TSC3, TYR1, UBC4, UBP14, UBS1, UGA2, UMP1, URA7, UTP20, VID24, VMA2, VPS15, YMC2, YPC1, YPK3, YRO2, YSA1, YSW1, YSY6, ZTA1 |

| | |
|---|---|
| **gal6** | AAC3, ACH1, ADH5, AGP2, AIM3, AIM4, AKL1, ALG1, ALG14, ALK2, AMN1, APD1, APL3, APN2, ARA1, ARL1, ATG14, ATG42, ATP3, BAP2, BEM1, BMT2, CBP6, CCZ1, CDC28, CDS1, CHS2, CHS3, CKS1, CMD1, CNM1, CNS1, COQ1, COR1, COS111, CPP1, CSG2, CSH1, CST26, CYC8, DER1, DSF2, DTR1, ECM13, ECM15, ECM2, ECM31, ECM33, ECM8, EDE1, EDS1, EHT1, ERD2, ETR1, EXO5, EXO84, FAT1, FES1, FIG1, FLR1, FMP23, FMT1, FTH1, FUI1, FUR4, FUS3, FZO1, GAL1, GAL10, GAL7, GDT1, GIP1, GPI18, GRS1, GRX7, HAP3, HEK2, HHF1, HHT1, HIR1, HMT1, HSL7, HSP26, HTA2, HTB2, ICS2, IFA38, IML3, IPP1, IRA1, IST2, KAP104, KTR3, KTR4, LAA2, LDB7, LDH1, LSM2, MAK5, MBA1, MCM2, MCM7, MEC1, MED8, MEO1, MIN6, MIN7, MIS1, MMS4, MNC1, MNN2, MOH1, MRPL16, MRPL36, MRPS9, MRX18, MSI1, MUD1, MUM2, NCL1, NHP6B, NPL4, NRG2, NTC20, NTH2, OLA1, OPY1, ORC2, PBY1, PCH2, PDR3, PDX3, PEP1, PET9, PEX32, PFF1, PGI1, PHO3, PHO5, PIM1, PIN4, POA1, POL12, POL30, POP7, POP8, PRE7, PRP6, PSY4, PTC4, QDR3, RCR1, RDH54, REB1, REG2, RER2, RFC5, RFS1, RFT1, RIB1, RIB7, RIM2, RKM3, RPB5, RPG1, RPL19A, RPL19B, RPL21A, RPL4A, RPS11B, RPS6B, RPS9B, RRN10, RRN6, RRT1, RTC2, RXT2, SAS3, SCO1, SCO2, SCT1, SEC17, SEC18, SEC66, SHE1, SHE3, SIF2, SLA1, SLI15, SLM4, SMP1, SMY2, SND3, SNR161, SPP381, SPT7, SSE2, STU1, SUP45, SUS1, SWD3, TAF5, TAT1, TBRT, TBS1, TCM62, TEC1, TFC1, TIM12, TIP1, TLC1, TPS1, TRM7, TSC3, TYR1, UBC4, UBP14, UBS1, UGA2, UMP1, URA7, UTP20, VID24, VMA2, VPS15, YMC2, YPC1, YPK3, YRO2, YSA1, YSW1, YSY6, ZTA1 |

**Table S3:** List of genes that have undergone duplication at each time point in each of the glucose and galactose replicate.

**300 generations**

| Replicate | Gene list |
|---|---|
| **glu1** | ACS2, APS1, ARP7, ASA1, ASR1, ATG26, ATG33, ATG38, BRR1, BUD8, CCC1, CCT6, CDC123, CIS1, CLB4, COA4, COQ9, CPR6, CRR1, DIB1, DIC1, DPH5, EMG1, ENT2, ERV2, FCY1, FKS1, FRE1, GAS2, GLN1, GRS2, HCR1, HMX1, HOS1, HRD3, IDP2, ILV5, IRC16, ISA2, KAP95, LTP1, LUG1, MAS1, MDL1, MDM36, MED1, MMR1, MRL1, MSC3, MSS51, NCW2, NHP6A, NIT3, NMT1, NOT5, NVJ2, OPY2, ORM2, PBA1, PEX13, PNP1, PUS5, PWP1, QRI5, RDS3, RFX1, RNH203, ROX1, RPL26A, RPL37A, RPP0, RPS31, RSA3, RSC2, RVS167, SAM1, SEC13, SEC8, SHH4, SKG3, SMK1, SND1, SNR41, SNR51, SNR70, SPE3, SPO24, SPO77, SRP54, SUA7, SUP16, SWI6, TAG1, TAL1, TEF1, TFB4, TFS1, TKL1, TOS4, TRR4, TUB4, UBA3, UPS1, UPS2, UPS3, UTP13, VMA13, VTA1, YKE2, YMC1 |
| **glu2** | - |
| **glu3** | ACL4, ADA2, ADE8, ADK1, ADR1, AFR1, AHA1, AIM7, AKR1, ALT2, AMD2, APC4, APS1, APT2, ARG82, ARH1, ARO1, ARO10, ARO80, ARP10, ARP7, ARX1, ASA1, ASP1, ATC1, ATG26, ATG38, ATP17, ATP22, ATP5, BCP1, BCS1, BFR2, BMH2, BNA7, BTT1, CAB5, CAD1, CBS2, CCC1, CCC2, CDC1, CDC123, CDC37, CDC40, CFT1, CHL4, CIA1, CLB4, CMI8, CNL1, COA4, COI1, COQ4, COQ9, COX20, COX26, CPP2, CPR1, CPR5, CPR6, CRF1, CRR1, CSN9, CTA1, CTH1, CTS2, CWC15, CWC21, CYM1, DAD4, DFM1, DIB1, DIG2, DIN7, DIT1, DIT2, DNF2, DOA4, DON1, DOP1, DOS2, DOT1, DPB4, DPH5, DPL1, DPP1, DXO1, DYN2, EAF1, EBS1, ECM11, ECM18, EKI1, EMG1, EMT1, ENT2, ENT5, ERD1, ERV2, ESC2, ESF1, EXG2, FCF1, FIN1, FMN1, FMP16, FOB1, FRE1, FRQ1, GCD6, GCN2, GGA1, GIC2, GIR2, GIS1, GLN1, GLO2, GPI11, GPI17, GPI19, GPI8, GRS2, GRX3, GTB1, GUK1, HCR1, HDA2, HEH2, HEL2, HEM1, HIM1, HKR1, HMO1, HMX1, HNT2, HOM2, HPR1, HPT1, HRD3, HRQ1, HSP42, HSP78, HST4, HTA1, HTB1, HXT3, HXT7, ICS3, IDP2, ILT1, INM2, INO2, IPK1, IPT1, IRC16, IRC3, IVY1, IZH1, JIP4, KEI1, KGD2, KIN1, KRE2, LCB2, LRS4, LSM6, LYS4, MAE1, MAK21, MCM21, MDL1, MDM36, MET32, MFA1, MFB1, MGP12, MHR1, MKC7, MMR1, MNN10, MOR1, MRL1, MRP1, MRP20, MRPL1, MRPL28, MRPL35, MRPL7, MRPS28, MRX10, MRX14, MRX16, MRX8, MSC2, MSC3, MSH6, MSN5, MSS116, MSS4, MSS51, MSW1, MTC5, MTH1, MTQ2, MZM1, NBP2, NCB2, NCW2, NGG1, NHA1, NHX1, NKP1, NMT1, NPL3, NSE3, NUM1, NUP42, NVJ3, OCA6, OMS1, OPY2, PAA1, PAC11, PAL1, PAM1, PBA1, PCF11, PDC2, PDR15, PDS1, PEP7, PET100, PEX10, PEX13, PEX29, PEX3, PEX5, PEX7, PFA5, PFU1, PGD1, PHM6, PHO8, PHO92, PIB1, PIB2, PKH1, PKH3, PLP1, PMP3, PMT7, PNP1, PPH3, PPM1, PPN1, PPZ2, PRO1, PRP28, PRP3, PRP42, PRY1, PRY3, PWP1, QRI5, RAD30, RAD34, RAD55, RAD9, RAV2, REF2, RFX1, RGA2, RGP1, RIB3, RKM2, RKM4, RKM5, RLI1, RMD5, RMT2, RNH202, RPA14, RPB7, RPL12B, RPL27B, RPL37A, RPN9, RPP2B, RPS13, RPS17B, RPS18A, RPS31, RPT3, RQC1, RRG1, RRN5, RRP1, RRP17, RRP45, RRP8, RSC3, RSM24, RTN1, RTR2, RTT103, RUB1, SAC3, SAC6, SAC7, SAM1, SAN1, SAS4, SBE2, SCC2, SDC1, SDH4, SDH6, SEC1, SEC13, SEC26, SEC7, SED1, SEM1, SHE9, SHU2, SIP1, SIR4, SIZ1, SKG3, SKP1, SLD5, SLS1, SLU7, SNF1, SNF11, SNM1, SNR13, SNU56, SNX41, SPC110, SPC19, SPO24, SPO71, SPP41, SPR28, SPT3, SRB7, SRP101, SRP54, SSD1, SSF2, SSN2, SSS1, SSY1, STB3, STE14, STE5, STN1, STP1, SUA7, SUF3, SUM1, SUP2, SUP35, SUR2, SVF1, SWA2, SWF1, SWI5, SWI6, SWM1, SWR1, SXM1, SYF1, TAF10, TAF12, TAG1, TCP1, TEF1, TFA1, TFB1, TFB3, TFB5, TFC6, TFS1, THI74, TIF35, TIM11, TIP41, TKL1, TLD1, TLG1, TMA64, TMN2, TMS1, TOM1, TOS4, TPS2, TRM1, TRM82, TRP4, TRR1, TRS120, TRS23, TRS31, TRS85, TSA2, TUB4, TVP15, TVP23, UBA2, UBC1, UBC13, UBC5, UBX5, UGO1, UME6, UPC2, UPS1, UPS2, URH1, UTP4, UTP5, UTP6, VBA4, VHS1, VMA13, VPS41, VPS52, VPS60, VPS64, VPS72, VPS74, VTA1, VTC5, WIP1, XRS2, YAP6, YCF1, YCG1, YFT2, YHP1, YKE2, YPQ2, YPR1, YPS7, YRA1, YSP2, ZIP1 |

| | |
|---|---|
| **glu4** | ACL4, ADA2, ADE8, ADK1, ADR1, AFR1, AHA1, AIM7, AKR1, ALT2, AMD2, APC4, APT2, ARG82, ARH1, ARO1, ARO10, ARO80, ARP10, ARX1, ASP1, ATC1, ATO3, ATP17, ATP22, ATP5, BCP1, BCS1, BFR2, BMH2, BNA7, BTT1, CAB5, CAD1, CBS2, CCC2, CCT6, CDC1, CDC34, CDC37, CDC40, CFT1, CHL4, CIA1, CIN10, CMI8, CNL1, COI1, COQ4, COX20, COX26, CPP2, CPR1, CPR5, CRF1, CSN9, CTA1, CTH1, CTS2, CWC15, CWC21, CYM1, DAD4, DFM1, DIG2, DIN7, DIT1, DIT2, DNF2, DOA4, DON1, DOP1, DOS2, DOT1, DPB4, DPL1, DPP1, DXO1, DYN2, EAF1, EBS1, ECM11, ECM18, EFT1, EKI1, EMC10, EMT1, ENT5, ERD1, ESC2, ESF1, EXG2, FCF1, FIN1, FMN1, FMP16, FOB1, FRQ1, GCD6, GCN2, GGA1, GIC2, GIR2, GIS1, GLO2, GPI11, GPI17, GPI19, GPI8, GRX3, GTB1, GUK1, HDA2, HEH2, HEL2, HEM1, HIM1, HKR1, HMO1, HNT2, HOM2, HPR1, HPT1, HRQ1, HSP42, HSP78, HST4, HTA1, HTB1, HXT3, HXT7, ILT1, INM2, INO2, IPK1, IPT1, IRC3, IVY1, JIP4, KEI1, KGD2, KIN1, KRE2, LCB2, LRS4, LSM6, LYS4, MAK21, MCM21, MET32, MFA1, MFB1, MGP12, MHR1, MKC7, MNN10, MOR1, MRP1, MRP20, MRPL1, MRPL28, MRPL35, MRPL7, MRPS28, MRX10, MRX14, MRX16, MRX8, MSC2, MSH6, MSN5, MSS116, MSS4, MSW1, MTC5, MTH1, MTQ2, MUS81, NBP2, NCB2, NGG1, NHX1, NKP1, NPL3, NSE3, NUM1, NUP42, NVJ3, OCA6, OMS1, PAA1, PAC11, PAL1, PAM1, PCF11, PDC2, PDR15, PDS1, PEP7, PET100, PEX10, PEX29, PEX3, PEX5, PEX7, PFA5, PFU1, PHM6, PHO8, PHO92, PIB1, PKH3, PLP1, PMP3, PMT7, PPH3, PPM1, PPN1, PPZ2, PRO1, PRP28, PRP3, PRP42, PST1, RAD30, RAD34, RAD55, RAD9, RAV2, REF2, RGA2, RGP1, RIB3, RKM2, RKM4, RLI1, RMD5, RMT2, RNH202, RPA14, RPB7, RPL12B, RPL27B, RPN9, RPP2B, RPS13, RPS17B, RPS18A, RPT3, RQC1, RRG1, RRP1, RRP17, RRP45, RRP8, RSC3, RSM24, RTN1, RTR2, RTT103, RUB1, RVB1, RVS167, SAC3, SAC6, SAC7, SAN1, SAS4, SBE2, SCC2, SDC1, SDH4, SDH6, SEC1, SEC26, SEC7, SED1, SEM1, SHE9, SHU2, SIP1, SIR4, SIZ1, SKP1, SLD5, SLU7, SLY1, SND1, SNF1, SNF11, SNM1, SNR13, SNU56, SNX41, SPC110, SPC19, SPO71, SPP41, SPR28, SPT3, SRB7, SRP101, SSD1, SSF2, SSN2, SSS1, SSY1, STB3, STE14, STE5, STN1, STP1, SUF3, SUM1, SUP2, SUP35, SUR2, SVF1, SWA2, SWF1, SWI5, SWM1, SWR1, SXM1, SYF1, TAF10, TAF12, TCP1, TFB1, TFB3, TFB5, TFC6, TGL2, THI74, TIF35, TIM11, TLD1, TLG1, TMA64, TMN2, TMS1, TOM1, TPS2, TRM1, TRM82, TRP4, TRR1, TRS120, TRS23, TRS31, TRS85, TSA2, TVP15, TVP23, UBA2, UBC1, UBC13, UBC5, UBX5, UGO1, UME6, UPC2, UPS3, URH1, UTP4, UTP5, UTP6, VBA4, VHS1, VPS41, VPS52, VPS60, VPS64, VPS72, VPS74, VTC5, WIP1, XRS2, YAP6, YCF1, YCG1, YFT2, YHP1, YOS9, YPQ2, YPR1, YPS7, YRA1, YSP2, ZIP1 |
| **glu5** | ACS2, APS1, ATG38, CCC1, CDC123, CLB4, COA4, COQ9, CPR6, CRR1, DPH5, ENT2, ERV2, FRE1, GLN1, HCR1, HMX1, HRD3, IDP2, IRC16, MAS1, MSC3, MSS51, NCW2, NMT1, PBA1, PEX13, PNP1, PUS5, PWP1, QRI5, RFX1, RNH203, RPS31, RSA3, SEC13, SHH4, SPO24, TAG1, TFS1, TUB4, UPS1, UPS2, UTP13, VMA13, YKE2 |
| **glu6** | ACS2, AIM7, APS1, ATG26, ATG38, CCC1, CDC123, CDC34, CLB4, COA4, COQ9, CPR6, CRR1, DOA4, DOS2, DPH5, EMC10, EMG1, ENT2, FMP16, FRE1, HCR1, HMX1, HRD3, IDP2, IFH1, IPT1, LCB2, MAK21, MAS1, MDL1, MMR1, MSC3, MSS51, NCW2, NMT1, OCA6, PAA1, PBA1, PDC2, PET100, PEX13, PNP1, PPH3, PST1, PUS5, PWP1, QRI5, RAD55, RFX1, RNH203, RPL37A, RPS13, RPS31, RRG1, RSA3, RTR2, SAM1, SEC13, SED1, SHH4, SHU2, SKG3, SNF11, STN1, SWI6, TAG1, TFB5, TFS1, TGL2, TOS4, TPS2, TUB4, UBC5, UCC1, UPS1, UPS2, UTP13, VPS41, VTA1, YKE2, YOS9 |
| **gal1** | AAD14, ACM1, ACO2, ACP1, ACT1, ADD66, ADE57, ADH2, ADH6, ADK2, ADY3, AFG1, AGA1, AGE1, AIM17, AIM18, AIM46, AIM6, ALD4, ALR1, ALR2, AMF1, ANK1, AOS1, APA2, API2, APL6, APM1, APM2, AQY1, AQY2, ARC35, ARG8, ARG81, ARN1, ASG7, ATF1, ATG10, ATG13, ATG18, ATG22, ATG27, ATG32, ATG36, ATG41, ATG7, ATP11, ATP12, ATP15, ATR1, AVT2, AXL2, BAS1, BAT1, BBP1, BCK2, BEM2, BET2, BIK1, BIO3, BIO4, BIO5, BMH1, BNA6, BRE4, BRE5, BRR2, BRR6, BSC5, BSC6, BSP1, BUD32, BUL2, BUR6, BXI1, CAB1, CAB4, CAC2, CAF40, CAN1, CAR2, CBP1, CBP2, CBT1, CCA1, CCT2, CDC14, CDC26, CDC33, CDC6, CHD1, CIN8, CIS3, CLA4, CLN2, CNA1, CNN1, COF1, COG3, COQ2, COQ5, COQ6, CPS1, CRG1, CSA1, CSE1, CSM2, CSS1, CSS2, CTF8, CTK3, CTR2, CTR9, CUB1, CUE4, CUS2, CWC21, CWC22, DAD2, DAL82, DAT1, DBP6, DBP8, DCP1, DIA1, DIF1, DIG2, DIM1, DIP5, DMC1, DNF1, DOA1, DOC1, DPB2, DPH2, DPM1, DSE4, DUG1, DUR3, EAP1, ECM25, ECM29, ECM30, ECM32, ECM7, ECO1, EFG1, EFM1, EGD2, EGO4, EGT2, ELO1, ELP6, EMC1, EMC3, EMC4, EMI1, EMI2, EMP47, EMW1, ENO1, ENO2, ENT4, ERG20, ERG9, ERJ5, ESF2, ESL2, ETP1, EUG1, FAB1, FAU1, FDC1, FDH1, FET4, FET5, FIG4, FIT1, FKH1, FKS3, FLO1, FLO10, FLO9, FLX1, FMO1, FMP10, FMP27, FMP32, FMP33, FOL2, FPK1, FPR2, FPR4, FPS1, FRD1, FRE4, FRE6, FRE7, FRM2, FUM1, FUS1, GAB1, GAL4, GAS1, GAS4, GCG1, GCN5, GDA1, GDB1, GDH1, GFD2, GID7, GIM5, GIN4, GLC8, GLK1, GLY1, GMC1, GMC2, GND1, GND2, GNP1, GON7, GOS1, GPB1, GPI16, GRC3, GRE2, GRH1, GRX1, GRX2, GRX4, GTA1, GTR1, GUD1, GUS1, GUT1, GYP5, GYP7, HAL5, HAP2, HAT2, HBN1, HBS1, HDA3, HFI1, HFM1, HHY1, HIS2, HIS4, HLR1, HMG2, HO, HOL1, HPA2, HPA3, HRP1, HRT1, HSP150, HSP31, HUA1, HUT1, HXK1, HXT11, HXT14, |

| | |
|---|---|
| | HXT16, HXT8, HXT9, IES6, IGD1, IKI1, IMA2, IMA4, IMA5, IMD3, IRC5, IRC6, IRC7, ISC10, ITR1, IZH1, JEN1, JIP4, JIP5, JJJ2, KAP114, KAR9, KEG1, KEL3, KGD4, KOG1, KRE1, KRE2, KRE28, KRE9, KRI1, LAA1, LAG1, LAM5, LCD1, LDB18, LEM3, LEU3, LNP1, LOS1, LPP1, LRG1, LSB3, LSB5, LSM3, LYS9, MAK10, MBB1, MCH4, MCM10, MCO14, MDH2, MDJ2, MDL2, MDM1, MDM31, MED7, MES1, MET10, MET16, MFG1, MGR1, MIA40, MID1, MIN10, MLC2, MLP1, MLP2, MMS1, MNL1, MNN11, MNN4, MNN5, MNT4, MOB2, MON2, MRP2, MRP4, MRPL20, MRPL4, MRPS12, MRPS18, MRX15, MRX20, MRX6, MSA2, MSB3, MSO1, MST1, MTC3, MTD1, MTG2, MTM1, MTO1, MUP3, MVD1, MXR2, MYG1, MZM1, NAB6, NBL1, NBP1, NCE101, NDD1, NDI1, NGK1, NGL3, NIP1, NMD3, NOG2, NOP19, NOP8, NPR2, NPR3, NUC1, NUD1, NUP133, NUP188, NUT2, NVJ1, OCA5, OM45, OMA1, OPI1, OPT1, OPT2, OST4, OST5, OSW7, OTU1, OTU2, OXA1, OXP1, OYE2, PAB1, PAC11, PAD1, PAN6, PAU2, PAU20, PBI1, PCC1, PCK1, PCL1, PCM1, PDA1, PDE1, PDI1, PDP3, PES4, PET122, PET494, PEX1, PEX11, PEX2, PEX29, PEX6, PFA3, PFD1, PFK27, PFS1, PFS2, PGA3, PHA2, PHO11, PHO13, PHO4, PHO8, PHO84, PHO90, PIR5, PKH1, PLC1, PLM2, PML39, POF1, POL5, POP2, PPM2, PPX1, PRB1, PRE4, PRE5, PRE8, PRP16, PRP19, PRP21, PRP4, PRS3, PSE1, PSF3, PSP1, PTH1, PTK1, PTP1, PTR2, PTR3, PUF6, PUG1, PUL3, PUL4, PUP2, PUS4, PXL1, PXR1, PZF1, QCR2, QCR6, QCR7, QCR8, RAD10, RAD2, RAD24, RAD3, RAD4, RAI1, RBA50, RBD2, RBH1, RCY1, RDR1, RET2, REV7, RFA2, RFA3, RFC3, RGD2, RGD3, RHO1, RIB3, RIB4, RIF2, RIM101, RIM21, RIM4, RIX1, RMD8, RME3, RML2, RMR1, RNH70, RNP1, RNQ1, ROF1, ROG3, RPC82, RPD3, RPF2, RPH1, RPL12A, RPL16A, RPL17B, RPL18A, RPL18B, RPL22B, RPL25, RPL29, RPL2A, RPL36B, RPL37B, RPL39, RPL40A, RPL40B, RPL6B, RPL8A, RPL8B, RPN10, RPN12, RPO26, RPO41, RPS14B, RPS19A, RPS19B, RPS1A, RPS20, RPS22A, RPS4B, RRI2, RRP40, RRP7, RRT5, RSC8, RSM19, RSM28, RTC1, RTF1, RTT102, RUF21, RUF22, RUF23, SAC1, SAM2, SAM3, SAM4, SAP155, SAP4, SAY1, SBP1, SCH9, SCW10, SCW4, SDD1, SDH2, SDH7, SDS22, SEC15, SEC20, SEC21, SEC23, SEC39, SEC53, SEC65, SET2, SET5, SHE10, SHU1, SIR1, SIR3, SIT1, SKG1, SKI3, SKN7, SKP2, SLD5, SLF1, SLH1, SLN1, SLO1, SMC2, SMF1, SMN1, SMT3, SMX3, SNA2, SNF1, SNF6, SNM1, SNR191, SNR40, SNR49, SNR53, SNR64, SNR67, SNR68, SNR80, SNR84, SNR85, SOM1, SOP4, SPC97, SPG3, SPO11, SPS1, SPS2, SPT2, SPT20, SRL3, SRO9, SRP40, SRP68, SRY1, SSB1, SSL2, SSP1, SST2, STB1, STB5, STE20, STE50, STE6, STL1, SUF6, SUP6, SUT390, SVP26, SWE1, SWI3, SWP82, TAD1, TAF1, TAF13, TAF8, TAN1, TCA17, TDA8, TGL3, TGL4, THI21, THI5, TMA108, TMT1, TNA1, TOG1, TOH1, TOR2, TOS6, TPK1, TPM2, TRF5, TRM11, TRM112, TRM13, TRP3, TRT2, TRZ1, TSL1, TSR2, TUB2, TUB3, TVP38, UBA1, UBI4, UBP11, UBP12, UBP15, ULI1, UPA2, URA1, URA5, URC2, UTP14, UTP9, VAN1, VHS2, VID30, VIK1, VMA6, VMA8, VMR1, VNX1, VPS13, VPS3, VPS4, VPS501, VPS52, VPS60, VPS68, VPS72, VPS9, WSC4, YAH1, YAP3, YBT1, YGK3, YKT6, YLF2, YPD1, YPT1, YPT11, YRA2, YTA7, ZDS2, ZIM17, ZIP2, ZOD1, ZPS1, ZRG17 |
| **gal2** | AAD14, AAD4, ACM1, ACO2, ACP1, ACS1, ACT1, ADD66, ADE57, ADH6, ADK2, ADY3, AFG1, AGA1, AGE1, AIM17, AIM18, AIM2, AIM46, AIM6, ALD4, ALR1, AMF1, ANK1, AOS1, APA2, API2, APL6, APM1, APM2, AQY1, ARC35, ARG8, ATF1, ATG13, ATG22, ATG32, ATG36, ATG41, ATG7, ATP11, ATP15, ATS1, AXL2, BAT1, BBP1, BCK2, BDH1, BDH2, BEM2, BET2, BGL2, BIO3, BIO4, BIO5, BMH1, BNA6, BOL1, BOL3, BRE4, BRE5, BRR2, BSC6, BSP1, BUD32, BUR6, BXI1, CAB1, CAB4, CAC2, CAF40, CAR2, CBP1, CBT1, CCA1, CCR4, CCT2, CDC14, CDC19, CDC24, CDC26, CDC33, CDC6, CHD1, CLA4, CLN2, CLN3, CNB1, CNE1, CNN1, COG3, COQ2, COQ6, CRG1, CSA1, CSE1, CSS1, CTF8, CTR2, CTR9, CUB1, CUE4, CUS2, CWC21, CWC22, CYC3, DAL82, DBP6, DBP8, DCP1, DIA1, DIF1, DIG2, DIM1, DIP5, DMC1, DNF1, DOA1, DOC1, DPB2, DPH2, DPM1, DUG1, DUR3, EAP1, ECM1, ECM25, ECM29, ECM32, ECM7, ECO1, EFG1, EGD2, EGO4, EGT2, ELO1, EMC3, EMI1, EMI2, EMW1, ENO1, ENO2, ERG9, ERJ5, ERV29, ERV46, ESF2, EUG1, FAU1, FDH1, FET5, FIG4, FIT1, FLC2, FLO1, FLO9, FLX1, FMO1, FMP10, FMP27, FMP32, FMP45, FOL2, FPK1, FPR2, FPR4, FRD1, FRE7, FUM1, FUN12, FUN19, FUN26, GAB1, GAL4, GAS4, GCG1, GCS1, GCV3, GDB1, GDH1, GDH3, GEM1, GFD2, GID7, GIN4, GLK1, GLY1, GMC1, GMC2, GND1, GND2, GNP1, GON7, GOS1, GPB2, GPI16, GRE2, GRH1, GRX1, GRX2, GRX4, GTA1, GUD1, GUS1, GUT1, GYP5, GYP7, HAP2, HAT2, HBT1, HDA3, HFI1, HHY1, HIS2, HLR1, HMG2, HO, HOL1, HPA2, HRP1, HRT1, HSP31, HUA1, HUT1, HXK1, HXT11, HXT14, HXT8, HXT9, IES6, IKI1, IMA2, IMA5, IRC5, IRC6, ISC10, ITR1, IZH1, JEN1, JIP5, KAP114, KAR9, KEG1, KEL3, KGD4, KOG1, KRE1, KRE2, KRE28, KRI1, LAA1, LAM5, LCD1, LEM3, LEU3, LNP1, LOS1, LPP1, LRG1, LSB3, LSB5, LSM3, LSM5, LYS9, MAK10, MBB1, MCM10, MCO14, MDH2, MDJ2, MDL2, MDM31, MED7, MES1, MET10, MET16, MFG1, MGR1, MIA40, MID1, MIN10, MLC2, MLP2, MMS1, MNL1, MNN4, MNN5, MNT4, MON2, MRP2, MRPL20, MRPL4, MRPS12, MRPS18, MRX15, MRX20, MRX6, MSB3, MSO1, MST1, MTG2, MTM1, MTO1, MTW1, MVD1, MXR2, MYG1, MZM1, NBL1, NBP1, NCE101, NGK1, NMD3, NOG2, NOP8, NPR3, NUC1, NUD1, NUT2, NVJ1, OAF1, OCA5, OM45, OMA1, OPI1, OPT1, |

| | |
|---|---|
| | OPT2, OST4, OSW7, OTU1, OXA1, OXP1, OYE2, PAB1, PAC11, PAD1, PAN6, PAU2, PAU20, PBI1, PCL1, PCM1, PDA1, PDI1, PDP3, PEA2, PET122, PET494, PEX1, PEX11, PEX2, PEX22, PEX29, PEX6, PFA3, PFK27, PFS1, PFS2, PHA2, PHO11, PHO12, PHO13, PHO4, PHO8, PHO90, PKH1, PLC1, PLM2, POL5, POP2, POP5, PPM2, PPX1, PRB1, PRE4, PRP16, PRP21, PRP4, PRP45, PSF3, PSP1, PTA1, PTH1, PTK1, PTP1, PTR2, PTR3, PUF6, PUG1, PUS4, PXL1, PXR1, PZF1, QCR2, QCR6, QCR7, RAD2, RAD24, RAD3, RAD4, RBA50, RBD2, RBG1, RCY1, RDR1, RET2, REV7, RFA2, RFC3, RGD3, RHO1, RIB3, RIB4, RIF2, RIM101, RIM21, RIM4, RIX1, RMD8, RML2, RNH70, ROF1, RPC82, RPD3, RPH1, RPL12A, RPL18B, RPL25, RPL29, RPL2A, RPL36B, RPL37B, RPL39, RPL40A, RPL6B, RPL8A, RPN10, RPN12, RPO26, RPS14B, RPS19B, RPS1A, RPS22A, RPS4B, RRP40, RRT5, RSC8, RSM19, RSM28, RTC1, RTF1, RTT102, RUF23, SAC1, SAM2, SAM3, SAM4, SAP155, SAY1, SBP1, SCC4, SCH9, SCW4, SDD1, SDH7, SDS22, SEC20, SEC21, SEC23, SEC39, SEC53, SET5, SHS1, SIR3, SKI3, SKN7, SKP2, SLD5, SLF1, SLH1, SLN1, SLO1, SMC2, SMF1, SMN1, SMT3, SMX3, SNA2, SNF1, SNF6, SNM1, SNR191, SNR40, SNR49, SNR53, SNR64, SNR67, SNR68, SNR80, SNR84, SOM1, SOP4, SPC72, SPC97, SPG3, SPI1, SPO11, SPS1, SPS2, SPT15, SPT2, SPT20, SRL3, SRO9, SRP40, SSB1, SSL2, SSP1, SST2, STB1, STB5, STE6, STL1, SUF6, SUP6, SUT390, SVP26, SWE1, TAD1, TAF1, TCA17, TDA8, TGL4, THI21, THI5, TMA108, TMT1, TNA1, TOG1, TOR2, TOS6, TPM2, TRF5, TRM11, TRM112, TRM13, TRP3, TRT2, TSL1, TVP38, UBA1, UBP12, UBP3, ULI1, UPA1, URA1, URC2, UTP9, VHS2, VIK1, VMA6, VMA8, VNX1, VPS3, VPS4, VPS52, VPS60, VPS68, VPS72, WHI4, WSC4, YAH1, YAT1, YGK3, YKT6, YOR1, YPD1, YPT1, YPT11, YRA2, YRF1-7, YTA7, ZIM17, ZPS1, ZRG17 |
| **gal3** | AAD14, AAD4, ACM1, ACO2, ACP1, ACS1, ACT1, ADD66, ADE57, ADH6, ADK2, ADY3, AFG1, AGA1, AGE1, AIF1, AIM17, AIM18, AIM2, AIM46, AIM6, ALD4, ALR1, ALR2, AMF1, ANK1, AOS1, APA2, API2, APL6, APM1, APM2, AQY1, AQY2, ARC35, ARG8, ARG81, ARN1, ATF1, ATG10, ATG13, ATG22, ATG27, ATG32, ATG36, ATG41, ATG7, ATP11, ATP12, ATP15, ATR1, AVT2, AXL2, BAS1, BAT1, BAT2, BBP1, BCK2, BEM2, BET2, BGL2, BIO2, BIO3, BIO4, BIO5, BMH1, BNA6, BOL1, BOL2, BOL3, BRE4, BRE5, BRR2, BRR6, BSC5, BSC6, BSP1, BUD32, BUL2, BUR6, BXI1, CAB1, CAB4, CAC2, CAF40, CAN1, CAR2, CBP1, CBP2, CBT1, CCA1, CCT2, CDC14, CDC19, CDC23, CDC24, CDC26, CDC33, CDC6, CHD1, CIN8, CLA4, CLN2, CLN3, CNN1, COF1, COG1, COG3, COQ2, COQ5, COQ6, COS10, COS9, CPD1, CPS1, CRG1, CSA1, CSE1, CSS1, CSS3, CTF8, CTK3, CTR2, CTR9, CUB1, CUE4, CUS2, CWC21, CWC22, CYC3, DAD2, DAK2, DAL5, DAL82, DAN1, DAN4, DAT1, DBP6, DBP8, DCP1, DFP4, DIA1, DIF1, DIG2, DIM1, DIP5, DMC1, DNF1, DOA1, DOC1, DPB2, DPH2, DPM1, DSE4, DUG1, DUR3, EAP1, ECM25, ECM29, ECM32, ECM34, ECM7, ECO1, EDC1, EFG1, EFM1, EGD2, EGO4, EGT2, ELO1, ELP6, EMC1, EMC3, EMC4, EMI1, EMI2, EMP47, EMW1, ENB1, ENO1, ENO2, ENT4, ERG13, ERG9, ERJ5, ERR2, ERR3, ERV29, ERV46, ESF2, ESL2, ETP1, EUG1, FAU1, FDC1, FDH1, FET4, FET5, FEX1, FEX2, FIG4, FIT1, FIT2, FIT3, FKS3, FLC2, FLO1, FLO10, FLO9, FLX1, FMO1, FMP10, FMP27, FMP32, FMP45, FOL2, FPK1, FPR2, FPR4, FPS1, FRD1, FRE3, FRE4, FRE5, FRE6, FRE7, FUM1, FUN12, FUN19, FZF1, GAB1, GAL4, GAS1, GAS4, GCG1, GCN5, GCS1, GCV3, GDA1, GDB1, GDH1, GEM1, GFD2, GID7, GIN4, GIP4, GLC8, GLK1, GLY1, GMC1, GMC2, GND1, GND2, GNP1, GON7, GOS1, GPB1, GPH1, GPI16, GRE2, GRH1, GRX1, GRX2, GRX4, GTA1, GTR1, GUD1, GUS1, GUT1, GYP5, GYP7, HAP2, HAT2, HBS1, HBT1, HDA3, HFI1, HFM1, HHY1, HLR1, HMG2, HMS2, HO, HOL1, HPA2, HPA3, HRP1, HRT1, HSP31, HSP32, HUA1, HUT1, HXK1, HXK2, HXT11, HXT13, HXT14, HXT15, HXT16, HXT17, HXT8, HXT9, IES6, IKI1, IMA1, IMA2, IMA4, IMA5, IPA1, IRC5, IRC6, IRC7, ISC10, ITR1, IZH1, JEN1, JIP5, KAP114, KAR9, KEG1, KEL3, KGD4, KIP3, KOG1, KRE1, KRE2, KRE28, KRE9, KRI1, LAA1, LAM5, LCD1, LDB18, LEM3, LEU3, LNP1, LOS1, LPP1, LRG1, LSB5, LSM3, LYS9, MAK10, MAL13, MAN2, MBB1, MCH4, MCO14, MDH2, MDJ2, MDL2, MDM1, MDM31, MDM34, MED7, MES1, MET10, MET16, MFG1, MGA1, MGM101, MGR1, MIA40, MID1, MIG2, MIN10, MLC2, MLP1, MMS1, MNL1, MNN11, MNN4, MNN5, MNT4, MON2, MRP2, MRPL20, MRPL4, MRPS12, MRPS18, MRS6, MRX15, MRX20, MRX6, MSA2, MSB3, MSO1, MST1, MTC3, MTD1, MTG2, MTM1, MTO1, MTW1, MUP3, MVD1, MXR2, MYG1, MZM1, NAB6, NBL1, NBP1, NCE101, NCS6, NDD1, NDI1, NGK1, NGL3, NIF3, NIP1, NMD3, NOG2, NOP19, NOP8, NPR2, NPR3, NUC1, NUD1, NUP133, NUP188, NUT2, NVJ1, OAF1, OCA5, OM45, OMA1, OPI1, OPT1, OPT2, ORC4, OST4, OST5, OSW7, OTU1, OTU2, OXA1, OXP1, OYE2, PAB1, PAC11, PAD1, PAN6, PAU15, PAU18, PAU19, PAU2, PAU20, PAU21, PBI1, PCC1, PCK1, PCL1, PCM1, PDA1, PDE1, PDI1, PDP3, PDR18, PET122, PET494, PEX1, PEX11, PEX2, PEX29, PEX6, PFA3, PFD1, PFK27, PFS1, PFS2, PGA3, PGU1, PHA2, PHO11, PHO12, PHO13, PHO4, PHO8, PHO84, PHO90, PHR1, PKH1, PLC1, PLM2, PML39, PMT4, POF1, POL5, POP2, POP5, PPM2, PPX1, PRB1, PRE4, PRE5, PRP16, PRP19, PRP21, PRP4, PRP45, PRS3, PSE1, PSF3, PSP1, PTA1, PTH1, PTK1, PTP1, PTR2, PTR3, PUF6, PUG1, PUL3, PUL4, PUP2, PUS4, PXL1, PXP3, PXR1, PZF1, QCR2, QCR6, QCR7, RAD2, RAD24, RAD3, RAD4, RAI1, RBA50, RBD2, RBG1, RBH1, RCY1, RDR1, RET2, REV7, RFA2, RFA3, RFC3, RGD2, RGD3, RHO1, RIB3, RIB4, RIE1, RIF2, RIM101, |

| | |
|---|---|
| | RIM21, RIM4, RIX1, RMD8, RME3, RML2, RMR1, RNH70, RNP1, ROF1, RPC82, RPD3, RPF2, RPH1, RPL12A, RPL17B, RPL18A, RPL18B, RPL25, RPL29, RPL2A, RPL36B, RPL37B, RPL39, RPL40A, RPL40B, RPL6B, RPL8A, RPL8B, RPN10, RPN12, RPO26, RPS12, RPS14B, RPS19A, RPS19B, RPS1A, RPS20, RPS22A, RPS4A, RPS4B, RRP40, RRT5, RSC8, RSM19, RSM28, RTC1, RTF1, RTG2, RTT102, RUF21, RUF23, SAC1, SAM2, SAM3, SAM4, SAP155, SAP4, SAY1, SBP1, SCH9, SCW4, SDD1, SDH2, SDH7, SDS22, SDT1, SEC15, SEC20, SEC21, SEC23, SEC39, SEC53, SEC65, SET5, SGV1, SHE10, SHS1, SHU1, SIR1, SIR3, SIT1, SKG1, SKI3, SKI8, SKN7, SKP2, SLD5, SLF1, SLH1, SLN1, SLO1, SMC2, SMF1, SMN1, SMT3, SMX3, SNA2, SNC1, SNF1, SNF6, SNM1, SNO4, SNR191, SNR40, SNR49, SNR53, SNR64, SNR67, SNR68, SNR80, SNR84, SNR85, SOL4, SOM1, SOP4, SPC72, SPC97, SPG3, SPO11, SPS1, SPS2, SPS22, SPT2, SPT20, SRL3, SRO9, SRP40, SRY1, SSB1, SSL2, SSP1, SST2, STB1, STB5, STE20, STE6, STL1, SUF6, SUP6, SUT390, SVP26, SWE1, SWI3, SWP82, TAD1, TAF1, TAF13, TAF8, TAN1, TCA17, TDA8, TGL3, TGL4, THI13, THI21, THI5, THP2, TIF3, TMA108, TMT1, TNA1, TOG1, TOH1, TOR2, TOS6, TPM2, TRF5, TRM11, TRM112, TRM13, TRP3, TRT2, TRZ1, TSL1, TUB2, TUB3, TVP38, UBA1, UBI4, UBP11, UBP12, UPA1, UPA2, URA1, URA5, URC2, UTP9, UTR2, VAM7, VAN1, VHS2, VID30, VIK1, VMA6, VMA8, VMR1, VNX1, VPS13, VPS3, VPS4, VPS501, VPS52, VPS60, VPS68, VPS72, VRG4, VTH1, WHI4, WSC4, YAH1, YAP3, YBT1, YEF1, YGK3, YKT6, YLF2, YOR1, YPD1, YPT1, YPT11, YPT32, YRA2, YTA7, ZDS2, ZIM17, ZIP2, ZNF1, ZPS1, ZRG17, ZRT1, ZUO1 |
| **gal4** | AAD14, ABZ1, ACM1, ACO2, ACP1, ACT1, ADD66, ADE57, ADH6, ADK2, AFG1, AGA1, AGE1, AGP1, AGX1, AIM17, AIM18, AIM46, AIM6, ALD4, ALG12, ALR1, AMF1, ANK1, AOS1, APA2, API2, APL6, APM1, APM2, AQY1, AQY2, ARC35, ARG8, ARG81, ARN1, ARN2, ATF1, ATG10, ATG13, ATG18, ATG22, ATG32, ATG36, ATG41, ATG7, ATP11, ATP15, ATR1, AXL2, BAS1, BAT1, BBP1, BCK2, BEM2, BET2, BIK1, BIO3, BIO4, BIO5, BMH1, BNA6, BOL1, BOL2, BOL3, BRE4, BRE5, BRR2, BRR6, BSC6, BSP1, BUD17, BUD32, BUL2, BUR6, BXI1, CAB1, CAB4, CAC2, CAF40, CAR2, CBP1, CBP2, CBT1, CCA1, CCT2, CDC14, CDC19, CDC23, CDC24, CDC26, CDC33, CDC6, CHD1, CLA4, CLN2, CLN3, CNB1, CNN1, COF1, COG3, COQ2, COQ5, COQ6, COS7, COS8, COX14, CPR8, CRG1, CSA1, CSE1, CSM2, CSS1, CSS2, CTF8, CTK3, CTR2, CTR9, CUB1, CUE4, CUS2, CWC21, CWC22, CYC3, CYC7, DAL82, DAT1, DBP6, DBP8, DCP1, DFP4, DIA1, DIF1, DIG2, DIM1, DIP5, DMC1, DNF1, DOA1, DOC1, DPB2, DPH2, DPM1, DUG1, DUR3, EAP1, ECM25, ECM29, ECM32, ECM34, ECM7, ECO1, EFG1, EFM1, EGD2, EGO4, EGT2, ELO1, ELP6, EMC3, EMI1, EMI2, EMW1, ENO1, ENO2, ENT4, ERG13, ERG9, ERJ5, ERO1, ERV46, ESF2, ESL1, ESL2, ETP1, EUG1, FAB1, FAU1, FDH1, FET4, FET5, FIG4, FIT1, FKH1, FLO1, FLO10, FLO9, FLX1, FMO1, FMP10, FMP27, FMP45, FOL2, FPK1, FPR2, FPR4, FPS1, FRD1, FRE6, FRE7, FRM2, FUM1, FUN12, FUN19, FUS1, GAB1, GAL4, GAS4, GCG1, GCN5, GCS1, GCV3, GDA1, GDB1, GDH1, GFD2, GID7, GIN4, GLC8, GLK1, GLY1, GMC1, GMC2, GND1, GND2, GNP1, GON7, GOS1, GPB1, GPH1, GPI16, GRE2, GRH1, GRX1, GRX2, GRX4, GTA1, GTR1, GUD1, GUS1, GUT1, GYP5, GYP7, HAC1, HAP2, HAT2, HBN1, HBT1, HDA3, HFI1, HHY1, HIS2, HIS4, HLR1, HMG2, HO, HOL1, HPA2, HRP1, HRT1, HSP31, HUA1, HUB1, HUT1, HXK1, HXT11, HXT14, HXT16, HXT8, HXT9, IES6, IGD1, IKI1, IMA4, IMA5, IRC5, IRC6, IRC7, ISC10, ITR1, IZH1, JEN1, JIP5, KAP114, KAR9, KEG1, KEL3, KGD4, KIP3, KOG1, KRE1, KRE2, KRE28, KRI1, LAA1, LAM5, LCD1, LDB18, LEM3, LEU3, LNP1, LOS1, LPP1, LSB3, LSB5, LSM3, LYS9, MAK10, MBB1, MCM10, MCO14, MDH2, MDJ2, MDL2, MDM1, MDM31, MDM34, MED7, MES1, MET10, MET16, MFG1, MGR1, MIA40, MID1, MIG2, MIL1, MIN10, MLC2, MLP1, MLP2, MMS1, MNL1, MNN11, MNN4, MNN5, MOB2, MON2, MRP2, MRPL20, MRPL4, MRPS12, MRPS18, MRS6, MRX15, MRX20, MRX6, MSB3, MSC1, MSO1, MST1, MTG2, MTM1, MTO1, MTW1, MUP3, MVD1, MXR2, MYG1, MZM1, NAB6, NBL1, NBP1, NCE101, NCS6, NDD1, NDI1, NGK1, NGL3, NIF3, NMD3, NOG2, NOP19, NOP8, NPR3, NUC1, NUD1, NUP188, NUT2, NVJ1, OCA5, OM45, OMA1, OPI1, OPT1, OPT2, ORC4, OST4, OSW7, OTU1, OTU2, OXA1, OXP1, OYE2, PAB1, PAC11, PAN6, PAU13, PAU18, PAU2, PAU20, PBI1, PCC1, PCK1, PCL1, PCM1, PDA1, PDE1, PDI1, PDP3, PES4, PET122, PET494, PEX1, PEX11, PEX2, PEX29, PEX6, PFA3, PFK27, PFS1, PFS2, PGA3, PHA2, PHO11, PHO12, PHO13, PHO4, PHO8, PHO84, PHO90, PKH1, PLC1, PLM2, PML39, POL5, POP2, POP5, PPG1, PPM2, PPX1, PRB1, PRE4, PRE5, PRP16, PRP19, PRP21, PRP4, PRS3, PSF3, PSP1, PTA1, PTH1, PTK1, PTP1, PTR2, PTR3, PUF6, PUG1, PUP2, PUS4, PXL1, PXP3, PXR1, PZF1, QCR2, QCR6, QCR7, RAD17, RAD2, RAD23, RAD24, RAD3, RAD4, RAI1, RBA50, RBD2, RBG1, RCY1, RDR1, RET2, REV7, RFA2, RFC3, RGD3, RHO1, RIB3, RIB4, RIF2, RIM101, RIM15, RIM21, RIM4, RIX1, RMD8, RML2, RNH70, RNP1, RNQ1, ROF1, ROG3, RPC82, RPD3, RPH1, RPL12A, RPL16A, RPL18B, RPL22B, RPL25, RPL29, RPL2A, RPL36B, RPL37B, RPL39, RPL40A, RPL40B, RPL6B, RPL8A, RPL8B, RPN10, RPN12, RPO26, RPO41, RPS12, RPS14B, RPS19A, RPS19B, RPS1A, RPS20, RPS22A, RPS4B, RRP40, RRP7, RRT5, RSC8, RSC9, RSM19, RSM28, RTC1, RTF1, RTT102, RUF21, RUF22, RUF23, SAC1, SAM2, SAM3, SAM4, SAP155, SAY1, SBP1, SCH9, SCW4, SDD1, SDH2, SDH7, SDS22, SEC12, SEC20, SEC21, SEC23, SEC39, SEC65, SET5, |

| | |
|---|---|
| | SGV1, SHS1, SHU1, SIR1, SIR3, SKG1, SKI3, SKI8, SKN7, SKP2, SLD5, SLF1, SLH1, SLN1, SLO1, SMC2, SMF1, SMN1, SMT3, SMX3, SNA2, SNF1, SNF6, SNM1, SNR191, SNR40, SNR49, SNR53, SNR64, SNR67, SNR68, SNR80, SNR84, SNR85, SOL1, SOM1, SOP4, SPC97, SPG3, SPO11, SPS1, SPS2, SPT2, SPT20, SRL3, SRO9, SRP40, SRY1, SSB1, SSK2, SSL2, SSP1, SST2, STB1, STB5, STE20, STE50, STE6, SUF6, SUP6, SUT390, SVP26, SWE1, TAD1, TAF1, TAF13, TAF8, TCA17, TDA8, TGL3, TGL4, THI21, THI5, THP2, TIF3, TMA108, TMT1, TNA1, TOG1, TOR2, TOS6, TPM2, TRF5, TRM11, TRM112, TRM13, TRP3, TRT2, TSL1, TUB2, TUB3, TVP38, UBA1, UBI4, UBP11, UBP12, ULI1, UPA2, URA1, URA5, URC2, UTP9, UTR2, UTR4, VAM7, VAN1, VHS2, VIK1, VMA6, VMA8, VMR1, VNX1, VPR1, VPS13, VPS3, VPS4, VPS52, VPS60, VPS68, VPS72, WHI4, WSC4, YAH1, YAP3, YBT1, YEF1, YGK3, YKT6, YLF2, YPD1, YPT1, YPT11, YPT32, YRA2, YRF1-2, YRF1-5, YRF1-6, YRF1-7, YTA7, ZDS2, ZIM17, ZNG1, ZPS1, ZRG17 |
| **gal5** | AAD10, AAD14, AAD4, ABP1, ABZ1, ACM1, ACO2, ACP1, ACS1, ACT1, ADD66, ADE57, ADH2, ADH6, ADK2, ADY3, AFG1, AGA1, AGE1, AGP1, AGP3, AGX1, AIM17, AIM18, AIM2, AIM46, AIM6, ALD4, ALR1, ALR2, AMF1, ANK1, ANP1, AOS1, APA2, API2, APL6, APM1, APM2, AQY1, AQY2, AQY3, ARC35, ARG8, ARG81, ARN1, ATF1, ATG10, ATG13, ATG22, ATG27, ATG32, ATG36, ATG41, ATG7, ATP11, ATP12, ATP15, ATR1, ATS1, AVT2, AXL2, BAS1, BAT1, BAT2, BBP1, BCK2, BDH1, BDH2, BEM2, BET2, BGL2, BIK1, BIO2, BIO3, BIO4, BIO5, BMH1, BNA6, BOL1, BOL2, BOL3, BRE4, BRE5, BRR2, BRR6, BSC5, BSC6, BSP1, BUD32, BUL2, BUR6, BXI1, CAB1, CAB4, CAC2, CAF40, CAN1, CAR2, CBP1, CBP2, CBT1, CCA1, CCR4, CCT2, CDC13, CDC14, CDC19, CDC24, CDC26, CDC33, CDC6, CHD1, CIN2, CIN8, CLA4, CLN2, CLN3, CNB1, CNE1, CNN1, COF1, COG1, COG3, COQ2, COQ5, COQ6, COS4, CPS1, CRG1, CSA1, CSE1, CSM1, CSM2, CSS1, CTF8, CTK3, CTR2, CTR9, CUB1, CUE4, CUS2, CWC22, CYC3, CYC7, DAK2, DAL5, DAL82, DAN1, DAN4, DAT1, DBP6, DBP8, DCP1, DDI2, DFP4, DIA1, DIF1, DIM1, DIP5, DMC1, DNF1, DOA1, DOC1, DPB2, DPH2, DPM1, DRS2, DSE4, DUG1, DUR3, EAP1, ECM1, ECM25, ECM29, ECM32, ECM34, ECM7, ECO1, EDC1, EFG1, EFM1, EGD2, EGO4, EGT2, ELO1, ELP6, EMC3, EMC4, EMI1, EMI2, EMP47, EMW1, ENO1, ENO2, ENT4, ENV7, ERG9, ERJ5, ERR2, ERV29, ERV46, ESF2, ESL2, ETP1, EUG1, FAS2, FAU1, FDH1, FET4, FET5, FEX1, FEX2, FIG4, FIT1, FIT2, FIT3, FKS3, FLC2, FLO1, FLO10, FLO9, FLX1, FMO1, FMP10, FMP27, FMP32, FMP45, FOL2, FPK1, FPR2, FPR4, FPS1, FRD1, FRE3, FRE4, FRE5, FRE6, FRE7, FRM2, FRT2, FUM1, FUN12, FUN19, FUN26, FUS1, FZF1, GAB1, GAL4, GAS1, GAS4, GCG1, GCN5, GCS1, GCV3, GDA1, GDB1, GDH1, GEM1, GFD2, GID7, GIN4, GIP4, GLC8, GLK1, GLY1, GMC1, GMC2, GND1, GND2, GNP1, GON7, GOS1, GPB1, GPB2, GPH1, GPI16, GRE2, GRH1, GRX1, GRX2, GRX4, GTA1, GTR1, GUD1, GUS1, GUT1, GYP5, GYP7, HAC1, HAP2, HAT2, HBN1, HBT1, HDA3, HFI1, HFM1, HHY1, HIS4, HLR1, HMG2, HMS2, HO, HOL1, HPA2, HPA3, HRA1, HRP1, HRT1, HSP31, HSP32, HSP82, HUA1, HUB1, HUT1, HXK1, HXK2, HXT11, HXT13, HXT14, HXT16, HXT17, HXT8, HXT9, HYP2, IES6, IKI1, IMA1, IMA4, IMA5, IPA1, IQG1, IRC5, IRC6, ISC10, ITR1, IZH1, JEN1, JIP5, KAP114, KAR9, KEG1, KEL3, KGD4, KIP3, KOG1, KRE1, KRE28, KRE9, KRI1, LAA1, LAM5, LCD1, LDB18, LEM3, LEU3, LNP1, LOS1, LPP1, LRG1, LSB5, LSM3, LSM5, LTE1, LYS9, MAK10, MAK16, MAL13, MAN2, MBB1, MCH4, MCM10, MCO14, MDH2, MDJ2, MDL2, MDM1, MDM31, MDM34, MED7, MES1, MET10, MET16, MFG1, MGA1, MGM101, MGR1, MIA40, MID1, MIG2, MIL1, MIN10, MLC2, MLP1, MLP2, MMS1, MNL1, MNN11, MNN4, MNN5, MNT4, MOB2, MON2, MRP2, MRPL20, MRPL4, MRPS12, MRPS18, MRX15, MRX20, MRX6, MSB3, MSO1, MST1, MTC3, MTC7, MTG2, MTM1, MTO1, MTW1, MUP3, MVD1, MXR2, MYG1, MYO4, MZM1, NAB6, NBL1, NBP1, NCE101, NCS6, NDD1, NDI1, NGK1, NGL3, NIF3, NIP1, NMD3, NOG2, NOP19, NOP8, NPR2, NPR3, NSL1, NUC1, NUD1, NUP188, NUT2, NVJ1, OAF1, OCA5, OM45, OMA1, OPI1, OPT1, OPT2, ORC4, OST4, OST5, OSW7, OTU1, OTU2, OXA1, OXP1, OYE2, PAB1, PAC11, PAN6, PAU2, PAU20, PAU21, PBI1, PCC1, PCK1, PCL1, PCM1, PDA1, PDE1, PDI1, PDP3, PDR18, PEA2, PET122, PET494, PEX1, PEX11, PEX2, PEX22, PEX6, PFA3, PFD1, PFK27, PFS1, PFS2, PGA3, PGU1, PHA2, PHO11, PHO13, PHO4, PHO84, PHO90, PHR1, PKH1, PLC1, PLM2, PML39, PMT2, PMT4, POL5, POP2, POP5, PPG1, PPM2, PPX1, PRB1, PRE4, PRE5, PRP16, PRP19, PRP21, PRP4, PRP45, PRS3, PSE1, PSF3, PSP1, PTA1, PTH1, PTK1, PTP1, PTR2, PTR3, PUF6, PUG1, PUL3, PUL4, PUP2, PUS4, PXL1, PXP3, PXR1, PZF1, QCR2, QCR6, QCR7, RAD2, RAD23, RAD24, RAD3, RAD4, RAI1, RBA50, RBD2, RBG1, RBH1, RCY1, RDR1, RET2, REV7, RFA2, RFA3, RFC3, RGD2, RGD3, RHO1, RIB3, RIB4, RIE1, RIF2, RIM101, RIM15, RIM21, RIM4, RIX1, RMD8, RME3, RML2, RMR1, RNH70, RNP1, RNQ1, ROF1, RPC82, RPD3, RPH1, RPL12A, RPL16A, RPL17B, RPL18A, RPL18B, RPL22B, RPL25, RPL29, RPL2A, RPL36B, RPL37B, RPL39, RPL40A, RPL40B, RPL6B, RPL8A, RPL8B, RPN10, RPN12, RPO26, RPO41, RPS14B, RPS19A, RPS19B, RPS1A, RPS20, RPS22A, RPS4A, RPS4B, RRP40, RRP7, RRT5, RSC8, RSM19, RSM28, RTC1, RTF1, RTG2, RTT102, RUF21, RUF23, RVB2, SAC1, SAM2, SAM3, SAM4, SAP155, SAP4, SAW1, SAY1, SBP1, SCC4, SCH9, SCW10, SCW4, SDD1, SDH2, SDH7, SDS22, SDT1, SEC15, SEC20, SEC21, SEC23, SEC39, SEC53, SEC65, SET5, SGV1, |

| | |
|---|---|
| | SHE10, SHS1, SHU1, SIP2, SIR1, SIR3, SIT1, SKG1, SKI3, SKI8, SKN7, SKP2, SLD5, SLF1, SLH1, SLN1, SLO1, SMC2, SMF1, SMN1, SMT3, SMX3, SNA2, SNC1, SNF6, SNO2, SNR191, SNR40, SNR49, SNR53, SNR64, SNR67, SNR68, SNR80, SNR84, SNR85, SNZ2, SOL1, SOL4, SOM1, SOP4, SPC72, SPC97, SPG3, SPI1, SPO11, SPS1, SPS2, SPT15, SPT2, SPT20, SRL3, SRO9, SRP40, SRP68, SRY1, SSB1, SSL2, SSO1, SSP1, SST2, STB1, STB5, STE20, STE50, STE6, STL1, SUF6, SUI3, SUP6, SUT390, SVP26, SWE1, SWI3, SWP82, TAD1, TAF1, TAF13, TAF8, TAN1, TCA17, TDA8, TGL3, TGL4, THI11, THI13, THI21, THI5, TIF3, TMA108, TMT1, TNA1, TOG1, TOH1, TOR2, TOS6, TPM2, TRF5, TRM11, TRM112, TRM13, TRP3, TRT2, TSL1, TUB2, TUB3, TVP38, UBA1, UBI4, UBP11, UBP12, UBP15, UBP3, UPA1, UPA2, URA1, URA5, URC2, UTP9, UTR2, UTR4, UTR5, VAM7, VAN1, VHS2, VID30, VIK1, VMA11, VMA6, VMA8, VMR1, VNX1, VPS13, VPS3, VPS4, VPS68, VRG4, WHI4, WSC4, YAH1, YAP3, YAR1, YBT1, YEF1, YGK3, YKT6, YLF2, YOR1, YPD1, YPT1, YPT11, YPT32, YRA2, YRF1-5, YRF1-7, YTA7, ZDS2, ZIM17, ZIP2, ZNF1, ZPS1, ZRG17, ZRT1, ZUO1 |
| **gal6** | AAD10, AAD14, AAD4, ABP1, ABZ1, ACM1, ACO2, ACP1, ACS1, ACT1, ADD66, ADE4, ADE57, ADH2, ADH4, ADH6, ADK2, ADY3, AFG1, AGA1, AGE1, AGP3, AIF1, AIM17, AIM18, AIM2, AIM29, AIM33, AIM46, AIM6, ALD4, ALO1, ALR1, ALR2, AMF1, ANK1, AOS1, APA1, APA2, API2, APL6, APM1, APM2, AQY1, AQY2, AQY3, ARC35, ARG8, ARG81, ARN1, ARN2, ATF1, ATG10, ATG13, ATG22, ATG32, ATG36, ATG41, ATG7, ATM1, ATP11, ATP15, ATP18, ATR1, AVT2, AXL2, AYT1, BAS1, BAT1, BAT2, BBP1, BCK2, BDH1, BDH2, BEM2, BET2, BGL2, BIO2, BIO3, BIO4, BIO5, BMH1, BNA6, BOL1, BOL2, BOL3, BRE4, BRE5, BRR2, BRR6, BSC5, BSC6, BSP1, BUD32, BUL2, BUR6, BXI1, CAB1, CAB4, CAC2, CAF40, CAN1, CAR2, CBP1, CBP2, CBT1, CCA1, CCT2, CDC14, CDC19, CDC24, CDC26, CDC33, CDC6, CHD1, CIN8, CLA4, CLN2, CLN3, CNE1, CNN1, COF1, COG1, COG3, COQ2, COQ5, COQ6, COS1, COS10, COS12, COS4, COS5, COS7, COS8, COX14, CPD1, CRG1, CSA1, CSE1, CSS1, CSS3, CTF8, CTK3, CTR2, CTR9, CUB1, CUE4, CUS2, CWC21, CWC22, CYC3, DAD2, DAK2, DAL1, DAL2, DAL4, DAL5, DAL7, DAL81, DAL82, DAN1, DAN4, DAT1, DBP6, DBP8, DCG1, DCP1, DDI2, DFP4, DIA1, DIF1, DIG2, DIM1, DIP5, DMC1, DNF1, DOA1, DOC1, DPB2, DPH2, DPM1, DRS2, DSE4, DSF1, DSN1, DUG1, DUR3, DYN3, EAP1, ECM1, ECM25, ECM29, ECM32, ECM34, ECM4, ECM7, ECO1, EDC1, EFG1, EFM1, EGD2, EGH1, EGO4, EGT2, ELO1, ELP6, EMC1, EMC3, EMC4, EMI1, EMI2, EMP47, EMW1, ENB1, ENO1, ENO2, ENT4, ERG13, ERG9, ERJ5, ERO1, ERR2, ERR3, ERV29, ERV46, ESF2, ESL2, ETP1, EUG1, FAU1, FDC1, FDH1, FET4, FET5, FEX1, FEX2, FIG2, FIG4, FIT1, FIT2, FIT3, FKS3, FLC2, FLO1, FLO10, FLO11, FLO9, FLX1, FMO1, FMP10, FMP27, FMP32, FMP45, FOL2, FPK1, FPR2, FPR4, FPS1, FRA1, FRD1, FRE3, FRE4, FRE5, FRE6, FRE7, FRT2, FUM1, FUN12, FUN19, FZF1, GAB1, GAL4, GAS1, GAS4, GAT4, GCG1, GCN5, GCS1, GCV3, GDA1, GDB1, GDH1, GEM1, GEX2, GFD2, GID7, GIM5, GIN4, GIP4, GLC8, GLK1, GLY1, GMC1, GMC2, GND1, GND2, GNP1, GON7, GOS1, GPB1, GPB2, GPH1, GPI13, GPI16, GRC3, GRE2, GRH1, GRX1, GRX2, GRX4, GSR1, GTA1, GTR1, GTT2, GUD1, GUS1, GUT1, GYP5, GYP7, HAP2, HAT2, HBS1, HBT1, HDA3, HFI1, HFM1, HHY1, HIR3, HIS2, HLR1, HMG2, HMS2, HO, HOL1, HPA2, HPA3, HRA1, HRP1, HRT1, HSP31, HSP32, HSU1, HUA1, HUB1, HUT1, HXK1, HXK2, HXT11, HXT13, HXT14, HXT15, HXT16, HXT17, HXT8, HXT9, ICR1, IES6, IKI1, IMA1, IMA2, IMA4, IMA5, INA22, IPA1, IRC19, IRC4, IRC5, IRC6, IRC7, ISA1, ISC10, ITR1, IZH1, JEN1, JIP5, JLP1, KAP114, KAR9, KEG1, KEL3, KGD4, KIN82, KIP3, KOG1, KRE1, KRE2, KRE28, KRI1, LAA1, LAG1, LAM5, LCB1, LCD1, LDB18, LEM3, LEU3, LIP1, LNP1, LOS1, LPP1, LRE1, LRG1, LSB3, LSB5, LSM3, LYS9, MAK10, MAK16, MAL13, MAN2, MBB1, MCH4, MCO14, MDH2, MDJ2, MDL2, MDM1, MDM31, MDM34, MED7, MES1, MET10, MET16, MET28, MFG1, MGA1, MGM101, MGR1, MHT1, MIA40, MID1, MIG2, MIN10, MLC2, MLP1, MMP1, MMS1, MND2, MNL1, MNN4, MNN5, MNT2, MNT4, MOB2, MON2, MPH3, MRP2, MRP4, MRPL20, MRPL4, MRPS12, MRPS18, MRS1, MRS6, MRX15, MRX20, MRX6, MSA2, MSB3, MSC1, MSH3, MSL1, MSO1, MST1, MTC3, MTD1, MTG2, MTM1, MTO1, MTW1, MUP3, MVD1, MXR2, MYG1, MYO4, MZM1, NAB6, NBL1, NBP1, NCE101, NCS6, NDD1, NDI1, NFT1, NGK1, NGL3, NIF3, NIP1, NMD3, NOG2, NOP19, NOP8, NPR2, NPR3, NUC1, NUD1, NUP133, NUP188, NUT2, NVJ1, OAF1, OCA5, OM45, OMA1, OPI1, OPT1, OPT2, ORC4, OST4, OST5, OSW7, OTU1, OTU2, OXA1, OXP1, OYE2, PAB1, PAC11, PAD1, PAN6, PAU1, PAU11, PAU13, PAU14, PAU15, PAU18, PAU19, PAU2, PAU20, PAU21, PBI1, PCC1, PCK1, PCL1, PCM1, PDA1, PDE1, PDI1, PDP3, PDR18, PET122, PET494, PEX1, PEX11, PEX2, PEX22, PEX29, PEX6, PFA3, PFK27, PFS1, PFS2, PGA3, PGU1, PHA2, PHO11, PHO12, PHO13, PHO4, PHO8, PHO84, PHO90, PHR1, PIP2, PKH1, PLC1, PLM2, PML39, PMT4, POF1, POL5, POP2, POP5, PPG1, PPM2, PPX1, PRB1, PRC1, PRE10, PRE4, PRE5, PRE8, PRI1, PRP16, PRP19, PRP21, PRP4, PRP45, PRS3, PSE1, PSF3, PSP1, PTA1, PTH1, PTK1, PTP1, PTR2, PTR3, PUF6, PUG1, PUL3, PUL4, PUP2, PUS4, PWR1, PXL1, PXP3, PXR1, PZF1, QCR2, QCR6, QCR7, RAD10, RAD17, RAD2, RAD24, RAD3, RAD4, RAI1, RBA50, RBD2, RBG1, RCY1, RDR1, RET2, REV7, RFA2, RFC3, RGD2, RGD3, RHO1, RIB3, RIB4, RIE1, RIF2, |

| | |
|---|---|
| | RIM101, RIM21, RIM4, RIX1, RIX7, RMD8, RME3, RML2, RMR1, RNH70, RNP1, ROF1, RPC82, RPD3, RPF2, RPH1, RPL12A, RPL18A, RPL18B, RPL25, RPL29, RPL2A, RPL36B, RPL37B, RPL39, RPL40A, RPL40B, RPL6B, RPL8A, RPL8B, RPN10, RPN12, RPO26, RPO41, RPR2, RPS12, RPS14B, RPS19A, RPS19B, RPS1A, RPS20, RPS22A, RPS4A, RPS4B, RRP40, RRT5, RRT7, RSC8, RSC9, RSM19, RSM28, RTC1, RTF1, RTG2, RTT102, RUF21, RUF23, SAC1, SAM2, SAM3, SAM4, SAP155, SAP4, SAW1, SAY1, SBP1, SCH9, SCP1, SCW10, SCW4, SDD1, SDH2, SDH7, SDS22, SDT1, SEC11, SEC15, SEC20, SEC21, SEC23, SEC39, SEC53, SEC65, SET5, SGV1, SHE10, SHS1, SHU1, SIR1, SIR3, SIT1, SKG1, SKI3, SKI8, SKN7, SKP2, SLD5, SLF1, SLH1, SLN1, SLO1, SMC2, SMF1, SMN1, SMT3, SMX3, SNA2, SNC1, SNF1, SNF6, SNM1, SNO2, SNO4, SNR191, SNR40, SNR49, SNR53, SNR64, SNR67, SNR68, SNR80, SNR84, SNR85, SNZ2, SOL1, SOL4, SOM1, SOP4, SOR1, SPC72, SPC97, SPG3, SPO11, SPS1, SPS2, SPS22, SPT2, SPT20, SQT1, SRL3, SRO9, SRP40, SSB1, SSL2, SSP1, SST2, STB1, STB5, STE20, STE6, STL1, STS1, SUF6, SUP6, SUT390, SUT532, SVP26, SWE1, SWP82, TAD1, TAF1, TAF13, TAF8, TAN1, TCA17, TDA8, TGL3, TGL4, THI11, THI13, THI21, THI5, THP2, TIF3, TMA108, TMT1, TNA1, TOG1, TOR2, TOS6, TPM2, TPO1, TRF5, TRM11, TRM112, TRM13, TRP3, TRT2, TRZ1, TSL1, TUB1, TUB2, TUB3, TVP38, UBA1, UBI4, UBP11, UBP12, UBP15, UFO1, ULI1, UPA1, UPA2, URA1, URA5, URC2, UTP14, UTP9, UTR2, VAM7, VAN1, VBA5, VEL1, VHS2, VID30, VIK1, VLD1, VMA6, VMA8, VMR1, VNX1, VPS13, VPS3, VPS4, VPS501, VPS52, VPS60, VPS68, VPS72, VPS9, VRG4, VTH1, WHI4, WSC4, YAH1, YAP3, YAP5, YBT1, YCT1, YEF1, YGK3, YKT6, YLF2, YME2, YOR1, YPD1, YPT1, YPT11, YPT32, YRA2, YRF1-2, YRF1-3, YRF1-5, YRF1-7, YRF1-8, YTA7, YVH1, ZDS2, ZIM17, ZIP2, ZNF1, ZOD1, ZPS1, ZRG17, ZRT1, ZUO1 |

**600 generations**

| Replicate | Gene list |
|---|---|
| **glu1** | APS1, ATG38, CCC1, CDC123, CLB4, COA4, COQ9, CPR6, CRR1, DPH5, ENT2, FRE1, HCR1, HMX1, HRD3, IDP2, MSC3, MSS51, NCW2, NMT1, PBA1, PEX13, PNP1, PWP1, QRI5, RFX1, RPS31, RSA3, SEC13, TAG1, TUB4, UPS1, UPS2, YKE2 |
| **glu2** | AAC1, ACE2, ACF2, ACS2, ADY4, ALK1, APC2, APJ1, APS1, ARL3, ARP7, ARP8, ASA1, ATG11, ATG21, ATG26, ATG33, ATG38, BAG7, BNA5, BRR1, BUB2, BUD8, BUR2, CAM1, CAR1, CCC1, CCT6, CDC123, CDC42, CIN10, CIS1, CKB1, CKI1, CLB4, COA4, COQ9, CPR6, CQD1, CRR1, CSR1, DCN1, DIB1, DIC1, DIG1, DIP2, DPC25, DPH5, DPH6, ECM22, EEB1, EFT1, EFT2, EGD1, ELC1, ELP3, ELP4, EMG1, ENT2, ERC1, ERI1, ERV2, ESS1, EST1, FBP1, FCY1, FKH2, FKS1, FMP30, FRE1, GAS2, GDE1, GET1, GID11, GLN1, GLR1, GRS2, GYP6, HCR1, HMX1, HOS1, HRD3, HTS1, ICS3, IDH2, IDP2, IFH1, ILV3, ILV5, IMP4, INA17, IRC16, ISA2, ISM1, JAC1, KAP95, KTR6, LAT1, LCL1, LEE1, LGE1, LPX1, LSM8, LTP1, LUG1, MAE1, MAK3, MAS1, MCM16, MDL1, MDM36, MED1, MET31, MGR2, MHP1, MKS1, MLF3, MMR1, MNN9, MRL1, MRX11, MSC3, MSD1, MSF1, MSK1, MSS51, MSY1, MUS81, NCW2, NHA1, NHP6A, NIS1, NIT3, NMT1, NOG1, NOP4, NOT5, OAZ1, OPY2, ORM2, OST3, PBA1, PDC5, PEP3, PEX13, PGD1, PIB2, PNG1, PNP1, PRY1, PRY3, PSY3, PUF2, PUS5, PUT1, PWP1, QRI5, REC107, RFX1, RKM5, RLM1, RMP1, RNH201, RNH203, ROX1, RPL16B, RPL26A, RPL37A, RPL43A, RPL9B, RPP0, RPS31, RPS6A, RRN5, RSA3, RSC2, RUP1, RVS167, SAM1, SEC13, SEC16, SEC61, SEC62, SEC8, SFL1, SGF11, SHH4, SIA1, SKG3, SLS1, SLX4, SMD3, SMK1, SND1, SPE3, SPE4, SPO24, SPO77, SRP54, SSE1, SSN3, SSU1, STP3, STT3, SUA7, SUN4, SUP16, SWI6, SYH1, TAG1, TAH18, TAL1, TEF1, TES1, TFA1, TFB4, TFS1, THP3, TIF5, TIP41, TIS11, TKL1, TMA22, TOM7, TOP3, TOS4, TPM1, TRR4, TUB4, UBA3, UCC1, UPS1, UPS2, UPS3, USB1, UTP13, VMA13, VPS16, VRP1, VTA1, WHI5, YDC1, YKE2, YMC1, YPS3, ZRT2 |

| | |
|---|---|
| **glu3** | ACL4, ADA2, ADE8, ADK1, ADR1, AFR1, AGE1, AHA1, AIM7, AKR1, ALT2, AMD2, APA2, APC4, API2, APT2, ARG82, ARH1, ARO1, ARO10, ARO80, ARP10, ARX1, ASP1, ATO3, ATP17, ATP22, ATP5, BCP1, BCS1, BFR2, BMH2, BNA7, BTT1, CAD1, CCC2, CDC34, CDC37, CDC40, CFT1, CHL4, CIA1, CIN10, CMI8, CNL1, COI1, COQ4, COX20, COX26, CPP2, CPR1, CPR5, CRF1, CSN9, CTA1, CTH1, CTS2, CWC15, CWC21, CYM1, DAD4, DBF4, DET1, DFM1, DIG2, DIN7, DIT1, DIT2, DNF2, DOA4, DON1, DOP1, DOS2, DOT1, DPB4, DPL1, DPP1, DXO1, DYN2, EAF1, EBS1, ECM11, ECM18, EFT1, EKI1, EMC10, EMI1, EMI2, EMT1, ENT5, ERD1, ESC2, ESF1, EUG1, EXG2, FCF1, FIN1, FMN1, FMP16, FOB1, FPR2, FRQ1, GCD6, GCN2, GGA1, GIC2, GIN4, GIR2, GIS1, GLO2, GMC1, GNP1, GPI11, GPI17, GPI19, GPI8, GRH1, GRX2, GRX3, GTB1, GUK1, HDA2, HEH2, HEL2, HEM1, HIM1, HKR1, HLR1, HMO1, HNT2, HOM2, HPR1, HPT1, HRQ1, HSP42, HSP78, HTA1, HTB1, HXT3, HXT7, ILT1, INM2, INO2, IPK1, IPT1, IRC3, ITR1, IVY1, IZH1, JIP4, KEI1, KGD2, KIN1, KRE2, LCB2, LCD1, LPP1, LRS4, LSM6, LYS4, MAK21, MCM21, MET32, MFA1, MFB1, MGP12, MHR1, MKC7, MNN10, MOR1, MRP1, MRP20, MRPL1, MRPL28, MRPL35, MRPL7, MRPS28, MRX10, MRX14, MRX16, MRX8, MSC2, MSH6, MSN5, MSS4, MSW1, MTC5, MTH1, MTQ2, MUS81, MZM1, NBP2, NCB2, NGG1, NHX1, NKP1, NPL3, NSE3, NUM1, NVJ3, OCA6, OMS1, PAA1, PAC11, PAL1, PAM1, PCF11, PDC2, PDR15, PDS1, PEP7, PET100, PEX10, PEX29, PEX3, PEX5, PEX7, PFA5, PFU1, PHM6, PHO8, PHO92, PIB1, PKH1, PKH3, PLM2, PMP3, PMT7, PPH3, PPM1, PPN1, PPZ2, PRO1, PRP28, PRP3, PRP42, PSP1, PST1, PUF6, QCR7, RAD30, RAD34, RAD55, RAD9, RAV2, RBA50, RGA2, RGP1, RIB3, RKM4, RLI1, RMD5, RMT2, RNH202, RPA14, RPB7, RPL12B, RPL27B, RPL37B, RPN9, RPP2B, RPS13, RPS17B, RPS18A, RPT3, RQC1, RRG1, RRP1, RRP17, RRP45, RRP8, RSC3, RSM24, RSM28, RTN1, RTR2, RTT103, RUB1, RVS167, SAC3, SAC6, SAC7, SAM2, SAN1, SBE2, SCC2, SDC1, SDH4, SDH6, SDH7, SEC1, SEC20, SEC26, SEC7, SED1, SEM1, SHE9, SHU2, SIP1, SIR4, SIZ1, SKP1, SLD5, SLF1, SLU7, SMT3, SNA2, SNF1, SNF11, SNM1, SNR13, SNR84, SNU56, SNX41, SPC110, SPC19, SPG3, SPO71, SPP41, SPR28, SPS1, SPS2, SPT3, SRB7, SRP101, SSD1, SSF2, SSN2, SSS1, SSY1, STB3, STE14, STE5, STN1, STP1, SUF3, SUM1, SUP2, SUP35, SUR2, SVF1, SWA2, SWF1, SWI5, SWM1, SWR1, SXM1, SYF1, TAF10, TAF12, TCP1, TFB1, TFB3, TFB5, TFC6, TGL2, THI74, TIF35, TIM11, TLD1, TLG1, TMA64, TMN2, TMS1, TOM1, TPI1, TPS2, TRM1, TRM82, TRP4, TRR1, TRS120, TRS23, TRS31, TRS85, TSA2, TVP15, TVP23, UBA2, UBC1, UBC13, UBC5, UBX5, UGO1, UME6, UPC2, URC2, URH1, UTP4, UTP5, UTP6, VBA4, VHS1, VPS3, VPS41, VPS52, VPS60, VPS64, VPS72, VPS74, VTC5, WIP1, XRS2, YAP6, YCF1, YCG1, YFT2, YHP1, YOS9, YPQ2, YPR1, YPS7, YRA1, YSP2, ZIP1 |
| **glu4** | ACL4, ADA2, ADE8, ADK1, ADR1, AFR1, AHA1, AIM7, AKR1, ALT2, AMD2, APC4, APS1, APT2, ARG82, ARH1, ARO1, ARO10, ARO80, ARP10, ARX1, ASP1, ATC1, ATO3, ATP17, ATP22, ATP5, BCP1, BCS1, BFR2, BMH2, BNA7, BTT1, CAB5, CAD1, CBS2, CCC2, CCT6, CDC1, CDC37, CDC40, CFT1, CHL4, CIA1, CIN10, CMI8, CNL1, COI1, COQ4, COX20, COX26, CPP2, CPR1, CPR5, CRF1, CSN9, CTA1, CTH1, CTS2, CWC15, CWC21, CYM1, DAD4, DFM1, DIG2, DIN7, DIT1, DIT2, DNF2, DOA4, DON1, DOP1, DOS2, DOT1, DPB4, DPH5, DPL1, DPP1, DXO1, DYN2, EAF1, EBS1, ECM11, ECM18, EFT1, EKI1, EMC10, EMT1, ENT5, ERD1, ESC2, ESF1, EXG2, FCF1, FIN1, FMN1, FMP16, FOB1, FRQ1, GCD6, GCN2, GGA1, GIC2, GIR2, GIS1, GLO2, GPI11, GPI17, GPI19, GPI8, GRX3, GTB1, GUK1, HDA2, HEH2, HEL2, HEM1, HIM1, HKR1, HMO1, HNT2, HOM2, HPR1, HPT1, HRQ1, HSP42, HSP78, HST4, HTA1, HTB1, HXT3, HXT7, IDP2, ILT1, INM2, INO2, IPK1, IPT1, IRC3, IVY1, IZH1, JIP4, KEI1, KGD2, KIN1, KRE2, LCB2, LRS4, LSM6, LYS4, MAK21, MAS1, MCM21, MET32, MFA1, MFB1, MGP12, MHR1, MKC7, MNN10, MOR1, MRP1, MRP20, MRPL1, MRPL28, MRPL35, MRPL7, MRPS28, MRX10, MRX14, MRX16, MRX8, MSC2, MSH6, MSN5, MSS116, MSS4, MSW1, MTC5, MTH1, MTQ2, MUS81, MZM1, NBP2, NCB2, NGG1, NHX1, NKP1, NPL3, NSE3, NUM1, NUP42, NVJ3, OCA6, OMS1, PAA1, PAC11, PAL1, PAM1, PCF11, PDC2, PDR15, PDS1, PEP7, PET100, PEX10, PEX29, PEX3, PEX5, PEX7, PFA5, PFU1, PHM6, PHO8, PHO92, PIB1, PKH1, PKH3, PLP1, PMP3, PMT7, PPH3, PPM1, PPN1, PPZ2, PRO1, PRP28, PRP3, PRP42, PUS5, RAD30, RAD34, RAD55, RAD9, RAV2, REF2, RFX1, RGA2, RGP1, RIB3, RKM2, RKM4, RLI1, RMD5, RMT2, RNH202, RPA14, RPB7, RPL12B, RPL27B, RPN9, RPP2B, RPS13, RPS17B, RPS18A, RPS31, RPT3, RQC1, RRG1, RRP1, RRP17, RRP45, RRP8, RSC3, RSM24, RTN1, RTR2, RTT103, RUB1, RVB1, RVS167, SAC3, SAC6, SAC7, SAN1, SAS4, SBE2, SCC2, SDC1, SDH4, SDH6, SEC1, SEC26, SEC7, SED1, SEM1, SHE9, SHH4, SHU2, SIP1, SIR4, SIZ1, SKP1, SLD5, SLU7, SLY1, SND1, SNF1, SNF11, SNM1, SNR13, SNU56, SNX41, SPC110, SPC19, SPO71, SPP41, SPR28, SPT3, SRB7, SRP101, SSD1, SSF2, SSN2, SSS1, SSY1, STB3, STE14, STE5, STN1, STP1, SUF3, SUM1, SUP2, SUP35, SUR2, SVF1, SWA2, SWF1, SWI5, SWM1, SWR1, SXM1, SYF1, TAF10, TAF12, TAG1, TCP1, TFB1, TFB3, TFB5, TFC6, TFS1, TGL2, THI74, TIF35, TIM11, TLD1, TLG1, TMA64, TMN2, TMS1, TOM1, TPS2, TRM1, TRM82, TRP4, TRR1, TRS120, TRS23, TRS31, TRS85, TSA2, TVP15, TVP23, UBA2, UBC1, UBC13, UBC5, UBX5, |

| | |
|---|---|
| | UGO1, UME6, UPC2, UPS2, UPS3, URH1, UTP4, UTP5, UTP6, VBA4, VHS1, VPS41, VPS52, VPS60, VPS64, VPS72, VPS74, VTC5, WIP1, XRS2, YAP6, YCF1, YCG1, YFT2, YHP1, YOS9, YPQ2, YPR1, YPS7, YRA1, YSP2, ZIP1 |
| **glu5** | ACE2, ACF2, ACS2, AIM20, AIM7, APC2, APS1, ARL3, ARO10, ARP7, ATG26, ATG38, ATG44, ATO3, BAP3, CAM1, CCC1, CCT6, CDC123, CDC34, CIN10, CKI1, CLB4, COA1, COA4, COI1, COQ9, CPR6, CRR1, DBF4, DCN1, DET1, DIG1, DIP2, DOA4, DOS2, DPH5, DPH6, EFT1, ELC1, EMC10, EMG1, ENT2, ERV2, FMP16, FRE1, GID11, GLN1, GUT2, HCR1, HEM12, HEM13, HMX1, HRD3, HTS1, IDP2, IMP21, IPT1, IRC16, KTR6, LCB2, LCL1, LEE1, LGE1, LSM6, MAK21, MAS1, MDL1, MMR1, MNN9, MSC3, MSS51, MUS81, NCW2, NHA1, NKP1, NMT1, OAZ1, OCA6, PAA1, PBA1, PDC5, PEP3, PEX13, PNP1, PPH3, PST1, PUS5, PUT1, PWP1, QRI5, RAD55, RFX1, RGA2, RKM5, RMP1, RNH203, RPC11, RPL37A, RPP2B, RPS13, RPS31, RRG1, RRN5, RTR2, RVS167, SAM1, SDH6, SEC13, SED1, SGF11, SHH4, SKG3, SLS1, SLX4, SMD3, SND1, SNF11, SPE4, SPO24, SRO7, SWI6, TAG1, TFS1, TGL2, TIP41, TIS11, TOS4, TPI1, TPS2, TUB4, UBC5, UBP7, UPS1, UPS2, UPS3, USB1, VMA13, VMS1, VPS16, VTA1, YKE2, YOS9, YPS3, YRA1, ZRT2 |
| **glu6** | ACS2, APS1, CAR1, CAR2, COQ9, DPH5, EMG1, ENO2, ENT2, ERV2, FMP46, GLN1, HMX1, HRD3, ICS3, IDP2, IMP21, IRC16, LPX1, LSM3, MAS1, MRPL4, MSS51, NAP1, NCW2, NMT1, PBA1, PLN1, PRY1, PRY3, PUS5, PWP1, QRI5, RFX1, RNH203, RPL37A, RPP0, RPS31, SAM1, SCS3, SHB17, SHH4, SKG3, SPO24, SWI6, TAG1, TFS1, TOS4, UIP5, UPS1, UPS2, UTH1, VMA13, VTA1, WHI5, YKE2 |
| **gal1** | AAD10, AAD14, AAD4, ABP1, ABZ1, ABZ2, ACM1, ACO2, ACP1, ACT1, ADD66, ADE4, ADE57, ADH2, ADH4, ADH6, ADK2, ADY3, AFG1, AGA1, AGE1, AGP1, AGP3, AIF1, AIM17, AIM18, AIM21, AIM33, AIM46, AIM6, ALD4, ALO1, ALR1, ALR2, AMF1, ANK1, ANP1, AOS1, APA1, APA2, API2, APL6, APM1, APM2, AQY1, AQY2, AQY3, ARC35, ARG8, ARG81, ARN1, ARN2, ASG7, ASH1, ASN1, ATF1, ATG10, ATG13, ATG18, ATG22, ATG27, ATG32, ATG36, ATG41, ATG7, ATM1, ATP11, ATP12, ATP15, ATP18, ATR1, AVT2, AXL2, AYT1, BAS1, BAT1, BAT2, BBP1, BCK2, BEM2, BET2, BGL2, BIK1, BIO2, BIO3, BIO4, BIO5, BMH1, BNA6, BOL1, BOL2, BOL3, BRE4, BRE5, BRR2, BRR6, BSC5, BSC6, BSP1, BUD32, BUL2, BUR6, BXI1, CAB1, CAB4, CAC2, CAF120, CAF40, CAN1, CAR2, CBP1, CBP2, CBT1, CCA1, CCT2, CDC13, CDC14, CDC19, CDC23, CDC24, CDC26, CDC33, CDC6, CHD1, CIN2, CIN8, CIS3, CLA4, CLN2, CLN3, CNB1, CNN1, COF1, COG1, COG3, COQ2, COQ5, COQ6, COS1, COS10, COS12, COS4, COS5, COS7, COS8, COX14, CPD1, CPS1, CRG1, CSA1, CSE1, CSM1, CSM2, CSS1, CSS2, CSS3, CTF8, CTK3, CTR2, CTR9, CUB1, CUE4, CUR1, CUS2, CWC21, CWC22, CYC3, CYC7, DAD2, DAK2, DAL1, DAL2, DAL3, DAL4, DAL5, DAL7, DAL81, DAL82, DAN1, DAN4, DAT1, DBP6, DBP8, DCG1, DCP1, DDI1, DDI2, DFP4, DIA1, DIA4, DIF1, DIG2, DIM1, DIP5, DJP1, DLD3, DMC1, DNA2, DNF1, DOA1, DOC1, DPB2, DPH2, DPM1, DSE4, DSF1, DSN1, DSS1, DTD1, DUG1, DUR3, DUS1, DYN3, EAP1, ECM25, ECM29, ECM32, ECM34, ECM7, ECO1, EDC1, EFG1, EFM1, EGD2, EGH1, EGO4, EGT2, ELO1, ELP6, EMC1, EMC3, EMC4, EMI1, EMI2, EMP47, EMW1, ENB1, ENO1, ENO2, ENT4, ENV7, ERG11, ERG13, ERG20, ERG24, ERG9, ERJ5, ERO1, ERR2, ERR3, ERV29, ERV46, ESF2, ESL2, ETP1, EUG1, FAB1, FAS2, FAT3, FAU1, FDC1, FDH1, FET4, FET5, FEX1, FEX2, FIG2, FIG4, FIT1, FIT2, FIT3, FKS3, FLO1, FLO10, FLO11, FLO9, FLX1, FMO1, FMP10, FMP27, FMP32, FMP33, FMP45, FOL2, FPK1, FPR2, FPR4, FPS1, FRA1, FRD1, FRE3, FRE4, FRE5, FRE6, FRE7, FRM2, FTR1, FUM1, FUN12, FUN19, FUS1, FZF1, GAB1, GAL4, GAS1, GAS4, GAT4, GCG1, GCN5, GCS1, GCV3, GDA1, GDB1, GDH1, GDH2, GEX2, GFD2, GID7, GIM5, GIN4, GLC8, GLK1, GLY1, GMC1, GMC2, GND1, GND2, GNP1, GON7, GOS1, GOT1, GPA1, GPB1, GPH1, GPI13, GPI16, GRC3, GRE2, GRH1, GRX1, GRX2, GRX4, GSR1, GTA1, GTR1, GTT1, GTT2, GUD1, GUS1, GUT1, GYP5, GYP7, HAL5, HAP2, HAS1, HAT2, HBN1, HBS1, HBT1, HCH1, HDA3, HER2, HFI1, HFM1, HHY1, HIR3, HIS2, HIS4, HLR1, HMG2, HMS2, HO, HOL1, HOM6, HPA2, HPA3, HRP1, HRT1, HSE1, HSH155, HSP150, HSP31, HSP32, HSP82, HSU1, HUA1, HUB1, HUT1, HXK1, HXK2, HXT11, HXT13, HXT14, HXT15, HXT16, HXT17, HXT8, HXT9, HYM1, HYP2, HYR1, ICR1, IES6, IGD1, IKI1, IMA1, IMA2, IMA3, IMA4, IMA5, IML1, INA22, IPA1, IQG1, IRC19, IRC24, IRC4, IRC5, IRC6, IRC7, ISA1, ISC10, IST3, ITR1, IZH1, JEN1, JIP4, JIP5, JJJ2, JLP1, JNM1, KAP114, KAR9, KCC4, KEG1, KEL3, KGD4, KIN82, KIP3, KOG1, KRE1, KRE2, KRE28, KRE6, KRE9, KRI1, LAA1, LAG1, LAM5, LCB1, LCD1, LDB18, LEM3, LEU2, LEU3, LEU5, LIP1, LNP1, LOS1, LOT5, LPP1, LRE1, LRG1, LSB3, LSB5, LSM3, LSM5, LYS1, LYS9, MAK10, MAL12, MAL13, MAN2, 46166, MBB1, MCH4, MCM10, MCO14, MDH2, MDJ2, MDL2, MDM1, MDM31, MDM34, MDY2, MED7, MES1, MET10, MET16, MET28, MET5, MFG1, MGA1, MGA2, MGM101, MGR1, MHT1, MIA40, MID1, MIG2, MIN10, MLC2, MLP1, MLP2, MMP1, MMS1, MND2, MNL1, MNN11, MNN4, MNN5, MNT2, MNT4, MOB2, MON2, MPC2, MPH1, MPH3, MRP2, MRP4, MRPL10, MRPL20, MRPL33, MRPL4, MRPS12, MRPS18, MRS1, MRS6, MRX15, MRX20, MRX6, MSA2, MSB3, MSB4, MSC1, MSH3, MSL1, MSN1, MSO1, MST1, MTC3, MTC7, MTD1, MTG2, MTM1, MTO1, MTR2, MTW1, MUP3, MVD1, MXR2, MYG1, |

| | |
|---|---|
| | MZM1, NAB6, NBL1, NBP1, NCA2, NCE101, NCE102, NCS6, NDD1, NDI1, NEM1, NFS1, NFT1, NGK1, NGL3, NIF3, NIP1, NMD3, NOG2, NOP19, NOP6, NOP8, NPR2, NPR3, NRE1, NSL1, NUC1, NUD1, NUP133, NUP188, NUT2, NVJ1, OCA5, OM45, OMA1, OPI1, OPT1, OPT2, ORC4, OSH7, OST4, OST5, OSW7, OTU1, OTU2, OXA1, OXP1, OYE2, PAB1, PAC11, PAD1, PAN1, PAN6, PAP2, PAU1, PAU11, PAU12, PAU13, PAU14, PAU15, PAU18, PAU19, PAU2, PAU20, PAU21, PAU4, PBI1, PBN1, PCC1, PCK1, PCL1, PCM1, PDA1, PDE1, PDI1, PDP3, PDR18, PEA2, PES4, PET122, PET494, PEX1, PEX11, PEX2, PEX29, PEX6, PFA3, PFD1, PFK27, PFS1, PFS2, PGA3, PGU1, PHA2, PHO11, PHO12, PHO13, PHO4, PHO8, PHO84, PHO90, PHR1, PIN3, PIP2, PIR5, PKH1, PKH3, PLC1, PLM2, PML39, PMT4, POF1, POL5, POP2, POP3, PPG1, PPM2, PPX1, PRB1, PRC1, PRE10, PRE4, PRE5, PRE8, PRI1, PRM1, PRP16, PRP19, PRP21, PRP3, PRP4, PRP8, PRR2, PRS1, PRS3, PSE1, PSF3, PSP1, PTA1, PTH1, PTH4, PTK1, PTP1, PTR2, PTR3, PUF6, PUG1, PUL3, PUL4, PUP2, PUS4, PWR1, PXA2, PXL1, PXP3, PXR1, PZF1, QCR10, QCR2, QCR6, QCR7, QCR8, RAD10, RAD17, RAD2, RAD23, RAD24, RAD3, RAD4, RAI1, RBA50, RBD2, RBG1, RBH1, RCY1, RDR1, RET2, REV7, RFA2, RFA3, RFC3, RGD2, RGD3, RHO1, RIB3, RIB4, RIE1, RIF2, RIM101, RIM21, RIM4, RIX1, RIX7, RMD6, RMD8, RME3, RML2, RMR1, RMT2, RNH70, RNP1, RNQ1, ROF1, ROG3, RPC82, RPD3, RPF2, RPH1, RPL12A, RPL14B, RPL16A, RPL17B, RPL18A, RPL18B, RPL25, RPL27A, RPL27B, RPL29, RPL2A, RPL36B, RPL37B, RPL39, RPL40A, RPL40B, RPL6B, RPL8A, RPL8B, RPN10, RPN12, RPO26, RPO41, RPR2, RPS12, RPS14B, RPS19A, RPS19B, RPS1A, RPS20, RPS22A, RPS4A, RPS4B, RRI1, RRI2, RRP40, RRP7, RRT5, RRT7, RSC8, RSC9, RSM19, RSM28, RTC1, RTF1, RTG2, RTT102, RUF21, RUF22, RUF23, RVB2, SAC1, SAM2, SAM3, SAM4, SAP155, SAP4, SAY1, SBP1, SCC4, SCH9, SCP1, SCW10, SCW4, SDC1, SDD1, SDH2, SDH7, SDS22, SDT1, SEC11, SEC15, SEC20, SEC21, SEC23, SEC39, SEC53, SEC65, SET2, SET5, SGV1, SHE10, SHR3, SHR5, SHS1, SHU1, SIP2, SIR1, SIR3, SIT1, SKG1, SKI3, SKI8, SKM1, SKN7, SKP2, SLD5, SLF1, SLH1, SLN1, SLO1, SMC2, SMF1, SMN1, SMT3, SMX3, SNA2, SNF1, SNF6, SNM1, SNO2, SNO4, SNR13, SNR191, SNR40, SNR49, SNR53, SNR64, SNR67, SNR68, SNR80, SNR84, SNR85, SNZ2, SOD2, SOL1, SOL3, SOL4, SOM1, SOP4, SOR1, SPC97, SPE1, SPG3, SPI1, SPO11, SPS1, SPS2, SPS22, SPT15, SPT16, SPT2, SPT20, SQT1, SRL3, SRO9, SRP40, SRP68, SRY1, SSB1, SSL2, SSO1, SSP1, SST2, STB1, STB5, STE20, STE50, STE6, STL1, STP2, STS1, SUE1, SUF6, SUI3, SUP53, SUP6, SUT390, SUT532, SVP26, SWE1, SWI3, SWP82, TAD1, TAF1, TAF13, TAF8, TAN1, TCA17, TCD1, TDA1, TDA3, TDA6, TDA8, TDA9, TGL3, TGL4, THI11, THI13, THI21, THI5, THP2, TIF3, TIM10, TIM22, TIM8, TLG1, TMA108, TMT1, TNA1, TOG1, TOH1, TOR2, TOS6, TPK1, TPM2, TPO1, TPO3, TRF5, TRM11, TRM112, TRM13, TRP3, TRS31, TRT2, TRZ1, TSL1, TTI2, TUB1, TUB2, TUB3, TVP38, UBA1, UBI4, UBP11, UBP12, UBP15, UBP3, UBP5, UFO1, UGO1, ULI1, UPA1, UPA2, URA1, URA5, URC2, URN1, UTP14, UTP9, UTR2, UTR4, UTR5, VAM7, VAN1, VBA5, VEL1, VHS2, VID30, VIK1, VLD1, VMA11, VMA6, VMA8, VMR1, VNX1, VPS13, VPS29, VPS3, VPS4, VPS501, VPS52, VPS60, VPS68, VPS72, VPS9, VRG4, VTH1, VTH2, WHI4, WSC2, WSC4, YAH1, YAP1801, YAP3, YAP5, YAR1, YBT1, YCT1, YEF1, YGK3, YKT6, YLF2, YME2, YOR1, YPD1, YPT1, YPT11, YPT32, YRA2, YRF1-2, YRF1-3, YRF1-5, YRF1-7, YRF1-8, YTA7, YVH1, ZDS2, ZIM17, ZIP2, ZNF1, ZOD1, ZPS1, ZRG17, ZRT1, ZUO1 |
| **gal2** | AAD10, AAD14, AAD4, ABP1, ABZ1, ACM1, ACO2, ACP1, ACS1, ACT1, ADD66, ADE4, ADE57, ADH2, ADH4, ADH6, ADK2, ADY3, AFG1, AGA1, AGE1, AGP1, AGP3, AGX1, AHC2, AIF1, AIM17, AIM18, AIM2, AIM21, AIM33, AIM46, AIM6, AIP5, ALD4, ALG12, ALO1, ALR1, ALR2, AMF1, ANK1, ANP1, AOS1, APA1, APA2, API2, APL6, APM1, APM2, AQY1, AQY2, AQY3, ARC35, ARE2, ARG8, ARG81, ARN1, ARN2, ASG7, ASH1, ASN1, ATF1, ATG10, ATG13, ATG18, ATG22, ATG27, ATG32, ATG36, ATG41, ATG7, ATM1, ATP11, ATP12, ATP15, ATP18, ATP23, ATR1, AVT2, AXL2, AYT1, BAS1, BAT1, BAT2, BBP1, BCK2, BDH1, BDH2, BEM2, BET2, BGL2, BIK1, BIO2, BIO3, BIO4, BIO5, BMH1, BNA6, BOL1, BOL2, BOL3, BRE4, BRE5, BRF1, BRR2, BRR6, BSC5, BSC6, BSP1, BUD17, BUD32, BUL2, BUR6, BXI1, CAB1, CAB4, CAC2, CAF120, CAF40, CAN1, CAR2, CBP1, CBP2, CBT1, CCA1, CCT2, CDC14, CDC19, CDC23, CDC24, CDC26, CDC33, CDC6, CHD1, CIN2, CIN8, CIS3, CLA4, CLN2, CLN3, CNB1, CNE1, CNN1, COF1, COG1, COG3, COQ2, COQ5, COQ6, COS1, COS10, COS12, COS4, COS5, COS7, COS8, COX14, CPD1, CPR4, CPR8, CPS1, CRG1, CSA1, CSE1, CSM1, CSM2, CSS1, CSS2, CSS3, CTF8, CTK3, CTR2, CTR9, CUB1, CUE4, CUR1, CUS2, CWC21, CWC22, CYC3, CYC7, DAK2, DAL1, DAL2, DAL3, DAL4, DAL5, DAL7, DAL81, DAL82, DAN1, DAN4, DAT1, DBP6, DBP8, DCG1, DCP1, DDI2, DFP4, DIA1, DIF1, DIG2, DIM1, DIP5, DJP1, DMC1, DNA2, DNF1, DOA1, DOC1, DPB2, DPH2, DPM1, DSE4, DSF1, DSN1, DUG1, DUR3, DYN3, EAP1, ECM1, ECM25, ECM29, ECM32, ECM34, ECM7, ECO1, EDC1, EFG1, EFM1, EGD2, EGH1, EGO2, EGO4, EGT2, ELO1, ELP6, EMC1, EMC3, EMC4, EMI1, EMI2, EMP47, EMW1, ENB1, ENO1, ENO2, ENT4, ENV7, ERG13, ERG20, ERG24, ERG9, ERJ5, ERO1, ERR2, ERR3, ERS1, ERV29, ERV46, ESF2, ESL1, ESL2, ETP1, EUG1, FAB1, FAS2, FAT3, FAU1, FDC1, FDH1, FET4, FET5, FEX1, FEX2, FIG2, FIG4, FIT1, FIT2, FIT3, FKH1, FKS3, FLC2, FLO1, FLO10, FLO11, FLO9, FLX1, |

| | |
|---|---|
| | FMO1, FMP10, FMP27, FMP32, FMP33, FMP45, FOL2, FPK1, FPR2, FPR4, FPS1, FRA1, FRD1, FRE3, FRE4, FRE5, FRE6, FRE7, FRM2, FTR1, FUB1, FUM1, FUN12, FUN19, FUS1, FZF1, GAB1, GAL4, GAS1, GAS4, GAT4, GCG1, GCN5, GCS1, GCV3, GDA1, GDB1, GDH1, GEM1, GEX2, GFD2, GID7, GIM5, GIN4, GLC8, GLK1, GLY1, GMC1, GMC2, GND1, GND2, GNP1, GON7, GOS1, GOT1, GPB1, GPB2, GPH1, GPI13, GPI16, GRC3, GRE2, GRH1, GRX1, GRX2, GRX4, GSR1, GTA1, GTR1, GTT1, GTT2, GUD1, GUS1, GUT1, GYP5, GYP7, HAC1, HAL5, HAP2, HAT2, HBN1, HBT1, HCH1, HDA3, HER2, HFI1, HFM1, HHY1, HIS2, HIS4, HLR1, HMG2, HMS2, HO, HOL1, HPA2, HPA3, HRP1, HRT1, HSP150, HSP31, HSP32, HSP82, HSU1, HUA1, HUB1, HUT1, HXK1, HXK2, HXT11, HXT13, HXT14, HXT15, HXT16, HXT17, HXT8, HXT9, HYM1, HYP2, HYR1, ICR1, IES6, IGD1, IKI1, IMA1, IMA2, IMA3, IMA4, IMA5, IMG2, INA22, IQG1, IRC19, IRC24, IRC4, IRC5, IRC6, IRC7, ISA1, ISC10, IST3, ITR1, IZH1, JEN1, JIP4, JIP5, JJJ2, JLP1, JNM1, KAP114, KAR9, KEG1, KEL3, KGD4, KIP3, KOG1, KRE1, KRE2, KRE28, KRE6, KRE9, KRI1, LAA1, LAG1, LAM5, LCB1, LCD1, LDB18, LEM3, LEU3, LIP1, LNP1, LOS1, LOT5, LPP1, LRE1, LRG1, LSB3, LSB5, LSC2, LSM3, LSM5, LYS1, LYS9, MAK10, MAL12, MAL13, MAN2, 46166, MBB1, MCM10, MCO14, MDH2, MDJ2, MDL2, MDM1, MDM31, MDM34, MED7, MES1, MET10, MET16, MET28, MFG1, MGA1, MGA2, MGM101, MGR1, MHT1, MIA40, MID1, MIG2, MIL1, MIN10, MLC2, MLP1, MLP2, MMP1, MMS1, MND2, MNL1, MNN11, MNN4, MNN5, MNT2, MNT4, MOB2, MON2, MPC2, MPC3, MPH3, MPP6, MRP2, MRP4, MRPL10, MRPL20, MRPL4, MRPL50, MRPS12, MRPS18, MRS1, MRS6, MRX15, MRX20, MRX6, MSB3, MSC1, MSL1, MSO1, MST1, MTC3, MTC7, MTG2, MTM1, MTO1, MTR2, MTW1, MUP3, MVD1, MXR2, MYG1, MZM1, NAB6, NBL1, NBP1, NCA2, NCE101, NCE102, NCS6, NDD1, NDI1, NFT1, NGK1, NGL3, NIF3, NIP1, NMD3, NOG2, NOP19, NOP8, NPR2, NPR3, NRE1, NSL1, NUC1, NUD1, NUP188, NUT2, NVJ1, OAF1, OCA5, OM45, OMA1, OPI1, OPT1, OPT2, ORC4, OST4, OST5, OSW7, OTU1, OTU2, OXA1, OXP1, OYE2, PAB1, PAC11, PAD1, PAN1, PAN6, PAT1, PAU1, PAU11, PAU12, PAU13, PAU14, PAU15, PAU18, PAU19, PAU2, PAU20, PAU21, PAU4, PBI1, PCC1, PCK1, PCL1, PCM1, PDA1, PDE1, PDI1, PDP3, PDR18, PEA2, PES4, PET122, PET494, PEX1, PEX11, PEX18, PEX2, PEX21, PEX22, PEX29, PEX6, PFA3, PFD1, PFK1, PFK27, PFS1, PFS2, PGA3, PGU1, PHA2, PHO11, PHO12, PHO13, PHO4, PHO8, PHO84, PHO90, PHR1, PIN3, PIR5, PKH1, PLC1, PLM2, PML39, PMT4, POF1, POL5, POP2, POP3, POP5, PPG1, PPM2, PPX1, PRB1, PRC1, PRE4, PRE5, PRE8, PRI1, PRM1, PRP16, PRP19, PRP21, PRP4, PRP45, PRP8, PRS3, PSE1, PSF3, PSP1, PTA1, PTC6, PTH1, PTK1, PTP1, PTR2, PTR3, PUF6, PUG1, PUL3, PUL4, PUP2, PUS4, PWR1, PXA2, PXL1, PXP3, PXR1, PZF1, QCR2, QCR6, QCR7, QCR8, RAD10, RAD2, RAD23, RAD24, RAD3, RAD4, RAI1, RBA50, RBD2, RBG1, RBH1, RCF2, RCY1, RDR1, RET2, REV7, RFA2, RFA3, RFC3, RGD2, RGD3, RHO1, RIB3, RIB4, RIE1, RIF2, RIM101, RIM15, RIM21, RIM4, RIX1, RIX7, RMD8, RME3, RML2, RMR1, RNH70, RNP1, RNQ1, ROF1, ROG3, RPC82, RPD3, RPH1, RPL12A, RPL16A, RPL17B, RPL18B, RPL22B, RPL25, RPL29, RPL2A, RPL36B, RPL37B, RPL39, RPL40A, RPL40B, RPL6B, RPL8A, RPL8B, RPN10, RPN12, RPO26, RPO41, RPR2, RPS14B, RPS19A, RPS19B, RPS1A, RPS20, RPS22A, RPS4A, RPS4B, RRP40, RRP7, RRT5, RRT7, RSA4, RSC8, RSC9, RSM19, RSM28, RTC1, RTF1, RTG2, RTT102, RUF21, RUF22, RUF23, RVB2, SAC1, SAM2, SAM3, SAM4, SAP155, SAP4, SAY1, SBP1, SCC4, SCH9, SCW10, SCW4, SDA1, SDD1, SDH2, SDH7, SDS22, SDT1, SEC11, SEC12, SEC15, SEC20, SEC21, SEC23, SEC39, SEC53, SEC65, SET2, SET5, SGV1, SHE10, SHS1, SHU1, SIR1, SIR3, SIT1, SKG1, SKI3, SKI8, SKN7, SKP2, SLD5, SLF1, SLH1, SLN1, SLO1, SMC2, SMF1, SMN1, SMT3, SMX3, SNA2, SNF1, SNF12, SNF6, SNM1, SNO2, SNO4, SNR191, SNR40, SNR45, SNR49, SNR53, SNR64, SNR67, SNR68, SNR80, SNR84, SNR85, SNZ2, SOL1, SOL2, SOL3, SOL4, SOM1, SOP4, SOR1, SPC72, SPC97, SPE1, SPG3, SPI1, SPO11, SPS1, SPS2, SPS22, SPT15, SPT2, SPT20, SQT1, SRB8, SRL3, SRO9, SRP40, SRP68, SSB1, SSK2, SSK22, SSL2, SSO1, SSP1, SST2, STB1, STB5, STE20, STE50, STE6, STL1, STS1, SUE1, SUF6, SUI3, SUP6, SUT390, SUT532, SVP26, SWE1, SWI3, SWP82, TAD1, TAF1, TAF13, TAF8, TAN1, TCA17, TDA1, TDA11, TDA6, TDA8, TGL3, TGL4, THI11, THI13, THI21, THI5, THP2, TIF3, TIM23, TMA108, TMT1, TNA1, TOG1, TOH1, TOR2, TOS6, TPK1, TPM2, TPO1, TPO3, TRF5, TRM11, TRM112, TRM13, TRP3, TRT2, TRX3, TSL1, TUB1, TUB2, TUB3, TUP1, TVP38, UBA1, UBI4, UBP11, UBP12, UBP15, UBP3, UFO1, ULI1, UPA1, UPA2, URA1, URA5, URC2, URN1, USV1, UTP14, UTP9, UTR2, UTR4, UTR5, VAM7, VAN1, VBA5, VEL1, VHS2, VID30, VIK1, VLD1, VMA11, VMA6, VMA8, VMR1, VNX1, VPR1, VPS13, VPS3, VPS4, VPS52, VPS60, VPS68, VPS72, VPS9, VRG4, VTH1, VTH2, WHI4, WSC2, WSC4, YAH1, YAP1801, YAP1802, YAP3, YAP5, YAR1, YBT1, YCT1, YEF1, YGK3, YKT6, YLF2, YME2, YOR1, YPD1, YPT1, YPT11, YPT32, YRA2, YRF1-2, YRF1-3, YRF1-5, YRF1-7, YRF1-8, YTA7, YVH1, ZDS2, ZIM17, ZIP2, ZNF1, ZNG1, ZOD1, ZPS1, ZRG17, ZRT1, ZUO1 |

| | |
|---|---|
| **gal3** | AAD10, AAD14, AAD4, ABP1, ABZ1, ACM1, ACO2, ACP1, ACS1, ACT1, ADD66, ADE4, ADE57, ADH2, ADH4, ADH6, ADK2, ADY3, AFG1, AGA1, AGE1, AGP1, AGP3, AGX1, AIF1, AIM17, AIM18, AIM2, AIM21, AIM33, AIM46, AIM6, ALD4, ALG12, ALO1, ALR1, ALR2, AMF1, ANK1, ANP1, AOS1, APA1, APA2, API2, APL6, APM1, APM2, AQY1, AQY2, AQY3, ARC35, ARG8, ARG81, ARN1, ARN2, ASG7, ATF1, ATG10, ATG13, ATG18, ATG22, ATG27, ATG32, ATG36, ATG41, ATG7, ATM1, ATP11, ATP12, ATP15, ATP18, ATR1, AVT2, AXL2, AYT1, BAS1, BAT1, BAT2, BBP1, BCK2, BDH1, BDH2, BEM2, BET2, BGL2, BIK1, BIO2, BIO3, BIO4, BIO5, BMH1, BNA6, BOL1, BOL2, BOL3, BRE4, BRE5, BRR2, BRR6, BSC5, BSC6, BSP1, BUD17, BUD32, BUL2, BUR6, BXI1, CAB1, CAB4, CAC2, CAF40, CAN1, CAR2, CBP1, CBP2, CBT1, CCA1, CCT2, CDC13, CDC14, CDC19, CDC23, CDC24, CDC26, CDC33, CDC39, CDC50, CDC6, CHD1, CIN2, CIN8, CLA4, CLN2, CLN3, CNB1, CNE1, CNN1, COF1, COG1, COG3, COQ2, COQ5, COQ6, COS1, COS10, COS12, COS4, COS5, COS6, COS7, COS8, COS9, COX14, CPD1, CPR8, CPS1, CRG1, CSA1, CSE1, CSM1, CSS1, CSS2, CSS3, CTF8, CTK3, CTR2, CTR9, CUB1, CUE4, CUS2, CWC21, CWC22, CYC3, CYC7, DAD2, DAK2, DAL1, DAL2, DAL3, DAL4, DAL5, DAL7, DAL81, DAL82, DAN1, DAN4, DAT1, DBP6, DBP8, DCG1, DCP1, DDI2, DFP4, DIA1, DIF1, DIG2, DIM1, DIP5, DJP1, DLD3, DMC1, DNF1, DOA1, DOC1, DPB2, DPH2, DPM1, DSE4, DSF1, DSN1, DTD1, DUG1, DUR3, DYN3, EAP1, ECM1, ECM25, ECM29, ECM32, ECM34, ECM4, ECM7, ECO1, EDC1, EFG1, EFM1, EGD2, EGH1, EGO4, EGT2, ELO1, ELP6, EMA35, EMC1, EMC3, EMC4, EMI1, EMI2, EMP47, EMW1, ENB1, ENO1, ENO2, ENT4, ENV7, ERG13, ERG20, ERG9, ERJ5, ERO1, ERR2, ERR3, ERV29, ERV46, ESF2, ESL2, ETP1, EUG1, FAB1, FAS2, FAU1, FDC1, FDH1, FET4, FET5, FEX1, FEX2, FIG2, FIG4, FIT1, FIT2, FIT3, FKS3, FLC2, FLO1, FLO10, FLO11, FLO9, FLX1, FMO1, FMP10, FMP27, FMP32, FMP45, FOL2, FPK1, FPR2, FPR4, FPS1, FRA1, FRD1, FRE2, FRE3, FRE4, FRE5, FRE6, FRE7, FRM2, FUM1, FUN12, FUN19, FUS1, FZF1, GAB1, GAL4, GAS1, GAS4, GAT4, GCG1, GCN5, GCS1, GCV3, GDA1, GDB1, GDH1, GDH2, GDH3, GEM1, GEX2, GFD2, GID7, GIM5, GIN4, GIP4, GIT1, GLC8, GLK1, GLY1, GMC1, GMC2, GND1, GND2, GNP1, GON7, GOS1, GOT1, GPB1, GPB2, GPH1, GPI13, GPI16, GRC3, GRE2, GRH1, GRX1, GRX2, GRX4, GSR1, GTA1, GTR1, GTT1, GTT2, GUD1, GUS1, GUT1, GYP5, GYP7, HAC1, HAL5, HAP2, HAT2, HBN1, HBS1, HBT1, HDA3, HER2, HFI1, HFM1, HHY1, HIR3, HIS2, HIS4, HLR1, HMG2, HMRA1, HMRA2, HMS2, HO, HOL1, HOM6, HPA2, HPA3, HRP1, HRT1, HSE1, HSP31, HSP32, HSP82, HSU1, HUA1, HUB1, HUT1, HXK1, HXK2, HXT11, HXT13, HXT14, HXT15, HXT16, HXT17, HXT8, HXT9, HYP2, HYR1, ICR1, IES6, IGD1, IKI1, IMA1, IMA2, IMA3, IMA4, IMA5, IML1, INA22, INO4, IPA1, IQG1, IRC19, IRC24, IRC4, IRC5, IRC6, IRC7, ISA1, ISC10, IST3, ITR1, IZH1, JEN1, JIP4, JIP5, JLP1, JNM1, KAP114, KAR9, KEG1, KEL3, KGD4, KIN82, KIP3, KOG1, KRE1, KRE2, KRE28, KRE9, KRI1, LAA1, LAG1, LAM5, LCB1, LCD1, LDB18, LEM3, LEU3, LIP1, LNP1, LOS1, LPP1, LRE1, LRG1, LSB3, LSB5, LSM3, LSM5, LYS1, LYS9, MAK10, MAL12, MAL13, MAL31, MAL32, MAN2, MBB1, MCH2, MCH4, MCO14, MDH2, MDJ2, MDL2, MDM1, MDM31, MDM34, MDY2, MED7, MES1, MET10, MET16, MET28, MFG1, MGA1, MGA2, MGM101, MGR1, MHT1, MIA40, MID1, MIG2, MIL1, MIN10, MLC2, MLP1, MMP1, MMS1, MND2, MNL1, MNN11, MNN4, MNN5, MNT2, MNT4, MOB2, MON2, MPH1, MPH3, MRP2, MRP4, MRPL20, MRPL4, MRPS12, MRPS18, MRS1, MRS6, MRX15, MRX20, MRX6, MSA2, MSB3, MSB4, MSC1, MSH3, MSL1, MSN1, MSO1, MST1, MTC3, MTC7, MTD1, MTG2, MTM1, MTO1, MTW1, MUP3, MVD1, MXR2, MYG1, MYO4, MZM1, NAB6, NBL1, NBP1, NCE101, NCS6, NDD1, NDI1, NFT1, NGK1, NGL3, NIF3, NIP1, NMD3, NOG2, NOP19, NOP8, NPR2, NPR3, NRE1, NSL1, NUC1, NUD1, NUP133, NUP188, NUT2, NVJ1, OAF1, OCA4, OCA5, OM45, OMA1, OPI1, OPT1, OPT2, ORC4, OST4, OST5, OSW7, OTU1, OTU2, OXA1, OXP1, OYE2, PAB1, PAC11, PAD1, PAN1, PAN6, PAP2, PAU1, PAU10, PAU11, PAU12, PAU13, PAU14, PAU15, PAU18, PAU19, PAU2, PAU20, PAU21, PAU24, PAU4, PBI1, PCC1, PCK1, PCL1, PCM1, PDA1, PDE1, PDI1, PDP3, PDR18, PEA2, PES4, PET122, PET494, PEX1, PEX11, PEX2, PEX22, PEX29, PEX6, PFA3, PFD1, PFK27, PFS1, PFS2, PGA3, PGU1, PHA2, PHO11, PHO12, PHO13, PHO4, PHO8, PHO84, PHO90, PHR1, PIP2, PKH1, PLC1, PLM2, PML39, PMT4, POF1, POL5, POP2, POP5, PPG1, PPM2, PPX1, PRB1, PRC1, PRE10, PRE4, PRE5, PRE8, PRI1, PRP16, PRP19, PRP21, PRP4, PRP45, PRR2, PRS3, PSE1, PSF3, PSP1, PTA1, PTH1, PTH4, PTK1, PTP1, PTR2, PTR3, PUF6, PUG1, PUL3, PUL4, PUP2, PUS4, PWR1, PXL1, PXP3, PXR1, PZF1, QCR2, QCR6, QCR7, QCR8, RAD10, RAD17, RAD2, RAD23, RAD24, RAD3, RAD4, RAI1, RBA50, RBD2, RBG1, RBH1, RCY1, RDR1, RET2, REV7, RFA2, RFA3, RFC3, RGD2, RGD3, RHO1, RIB3, RIB4, RIE1, RIF2, RIM101, RIM15, RIM21, RIM4, RIX1, RIX7, RMD6, RMD8, RME3, RML2, RMR1, RNH70, RNP1, RNQ1, ROF1, ROG3, RPC82, RPD3, RPF2, RPH1, RPL12A, RPL17B, RPL18A, RPL18B, RPL22B, RPL25, RPL29, RPL2A, RPL36B, RPL37B, RPL39, RPL40A, RPL40B, RPL6B, RPL8A, RPL8B, RPN10, RPN12, RPO26, RPO41, RPR2, RPS12, RPS14B, RPS19A, RPS19B, RPS1A, RPS20, RPS22A, RPS4A, RPS4B, RRI1, RRI2, RRP40, RRP7, RRT5, RRT7, RSC8, RSC9, RSM19, RSM28, RTC1, RTF1, RTG2, RTT102, RUF21, RUF22, RUF23, RVB2, SAC1, SAM2, SAM3, SAM4, SAP155, SAP4, SAY1, SBP1, SCC4, SCH9, SCP1, SCW10, SCW4, SDD1, SDH2, |

| | |
|---|---|
| | SDH7, SDS22, SDT1, SEC11, SEC12, SEC15, SEC20, SEC21, SEC23, SEC39, SEC53, SEC65, SET2, SET5, SGV1, SHE10, SHR5, SHS1, SHU1, SIP2, SIR1, SIR3, SIT1, SKG1, SKI3, SKI8, SKM1, SKN7, SKP2, SLD5, SLF1, SLH1, SLN1, SLO1, SMC2, SMF1, SMN1, SMT3, SMX3, SNA2, SNC1, SNF1, SNF6, SNM1, SNO2, SNO4, SNR191, SNR40, SNR49, SNR53, SNR64, SNR67, SNR68, SNR80, SNR84, SNR85, SNZ2, SOL1, SOL4, SOM1, SOP4, SOR1, SPC72, SPC97, SPG3, SPI1, SPO11, SPS1, SPS2, SPS22, SPT15, SPT2, SPT20, SQT1, SRL3, SRO9, SRP40, SRP68, SRY1, SSB1, SSK2, SSL2, SSO1, SSP1, SST2, STB1, STB5, STE20, STE50, STE6, STL1, STS1, SUF1, SUF6, SUI3, SUP6, SUT390, SUT532, SVP26, SWE1, SWI3, SWP82, TAD1, TAF1, TAF13, TAF8, TAN1, TCA17, TDA8, TGL3, TGL4, THI11, THI13, THI21, THI5, THP2, TIF3, TIM22, TMA108, TMT1, TNA1, TOG1, TOH1, TOR2, TOS6, TPK1, TPM2, TPO1, TRF5, TRM11, TRM112, TRM13, TRP3, TRT2, TRZ1, TSL1, TUB1, TUB2, TUB3, TVP38, UBA1, UBI4, UBP11, UBP12, UBP15, UBP3, UFO1, ULI1, UPA1, UPA2, URA1, URA5, URC2, UTP14, UTP9, UTR2, UTR4, UTR5, VAM7, VAN1, VBA5, VEL1, VHS2, VID30, VIK1, VLD1, VMA11, VMA6, VMA8, VMR1, VNX1, VPS13, VPS3, VPS4, VPS501, VPS52, VPS60, VPS68, VPS72, VPS9, VRG4, VTH1, VTH2, WHI4, WSC4, YAH1, YAP3, YAP5, YAR1, YBT1, YCT1, YEF1, YGK3, YKT6, YLF2, YME2, YOR1, YPD1, YPT1, YPT11, YPT32, YRA2, YRF1-2, YRF1-3, YRF1-4, YRF1-5, YRF1-7, YRF1-8, YTA7, YVH1, ZDS2, ZEO1, ZIM17, ZIP2, ZNF1, ZNG1, ZOD1, ZPS1, ZRG17, ZRT1, ZUO1 |
| **gal4** | AAD10, AAD14, AAD4, ABP1, ABZ1, ACC1, ACM1, ACO2, ACP1, ACS1, ACT1, ADD66, ADE4, ADE57, ADH2, ADH4, ADH6, ADK2, ADY3, AFG1, AGA1, AGE1, AGP1, AGP3, AGX1, AHC2, AIF1, AIM17, AIM18, AIM2, AIM21, AIM29, AIM33, AIM46, AIM6, AIP5, ALD4, ALG12, ALG5, ALO1, ALR1, ALR2, AMF1, ANK1, ANP1, AOS1, APA2, API2, APL6, APM1, APM2, AQY1, AQY2, AQY3, ARC35, ARE1, ARE2, ARG8, ARG81, ARN1, ARN2, ASG7, ASH1, ASN1, ATF1, ATG10, ATG13, ATG15, ATG18, ATG22, ATG23, ATG27, ATG32, ATG36, ATG41, ATG7, ATM1, ATP11, ATP12, ATP15, ATP18, ATP23, ATR1, ATS1, AVT2, AXL2, AYT1, BAS1, BAT1, BAT2, BBP1, BCK2, BDH1, BDH2, BEM2, BET2, BET3, BGL2, BIK1, BIO2, BIO3, BIO4, BIO5, BMH1, BNA6, BOL1, BOL2, BOL3, BRE4, BRE5, BRF1, BRR2, BRR6, BSC5, BSC6, BSP1, BUD17, BUD23, BUD31, BUD32, BUL2, BUR6, BXI1, CAB1, CAB4, CAC2, CAF120, CAF40, CAN1, CAR2, CBP1, CBP2, CBT1, CCA1, CCR4, CCT2, CDC13, CDC14, CDC19, CDC23, CDC24, CDC26, CDC33, CDC39, CDC50, CDC6, CET1, CHC1, CHD1, CHP1, CIN2, CIN8, CIS3, CLA4, CLN2, CLN3, CMK1, CNA1, CNB1, CNE1, CNN1, COF1, COG1, COG3, COQ2, COQ5, COQ6, COS1, COS10, COS12, COS4, COS5, COS6, COS7, COS8, COS9, COX14, CPD1, CPR4, CPR8, CPS1, CRG1, CRN1, CSA1, CSE1, CSM1, CSM2, CSS1, CSS2, CSS3, CTF8, CTK3, CTR2, CTR86, CTR9, CUB1, CUE4, CUR1, CUS2, CWC2, CWC21, CWC22, CYC3, CYC7, CYS3, DAD2, DAK2, DAL1, DAL2, DAL3, DAL4, DAL5, DAL7, DAL81, DAL82, DAN1, DAN4, DAT1, DBP6, DBP8, DCG1, DCP1, DDI2, DEP1, DFP4, DIA1, DIF1, DIG2, DIM1, DIP5, DJP1, DLD3, DMC1, DNA2, DNF1, DOA1, DOC1, DPB2, DPH2, DPM1, DRE2, DRS2, DSE4, DSF1, DSN1, DTD1, DUG1, DUR3, DYN3, EAP1, ECM1, ECM25, ECM29, ECM30, ECM32, ECM34, ECM4, ECM7, ECO1, EDC1, EFG1, EFM1, EGD2, EGH1, EGO2, EGO4, EGT2, ELO1, ELP6, EMA35, EMC3, EMC4, EMI1, EMI2, EMP47, EMW1, ENB1, ENO1, ENO2, ENT4, ENV7, ERG13, ERG20, ERG24, ERG9, ERJ5, ERO1, ERP2, ERR2, ERR3, ERS1, ERV29, ERV46, ESF2, ESL1, ESL2, ETP1, EUG1, FAB1, FAS2, FAT3, FAU1, FDC1, FDH1, FET4, FET5, FEX1, FEX2, FIG2, FIG4, FIT1, FIT2, FIT3, FKH1, FKS3, FLC2, FLO1, FLO10, FLO11, FLO9, FLX1, FMO1, FMP10, FMP27, FMP32, FMP33, FMP40, FMP45, FOL2, FPK1, FPR2, FPR4, FPS1, FRD1, FRE2, FRE3, FRE4, FRE5, FRE6, FRE7, FRM2, FRT2, FTR1, FUB1, FUM1, FUN12, FUN14, FUN19, FUN26, FUN30, FUS1, FZF1, GAB1, GAL4, GAS1, GAS4, GAT4, GCG1, GCN5, GCS1, GCV3, GDA1, GDB1, GDH1, GDH2, GDH3, GEM1, GEX2, GFD2, GID7, GIM5, GIN4, GIP4, GIT1, GLC8, GLK1, GLY1, GMC1, GMC2, GND1, GND2, GNP1, GON7, GOS1, GPB1, GPB2, GPH1, GPI16, GPT2, GRC3, GRE1, GRE2, GRH1, GRX1, GRX2, GRX4, GSR1, GSY1, GTA1, GTR1, GTT1, GTT2, GUD1, GUS1, GUT1, GYP5, GYP7, HAC1, HAL5, HAP2, HAP5, HAT2, HBN1, HBS1, HBT1, HCH1, HCM1, HDA3, HFI1, HFM1, HHY1, HIR3, HIS2, HIS4, HLR1, HMG2, HMRA1, HMRA2, HMS2, HO, HOL1, HOM6, HPA2, HPA3, HRA1, HRP1, HRT1, HSP150, HSP31, HSP32, HSP82, HSU1, HUA1, HUB1, HUT1, HXK1, HXK2, HXT11, HXT13, HXT14, HXT15, HXT16, HXT17, HXT8, HXT9, HYM1, HYP2, HYR1, ICR1, IES6, IGD1, IKI1, IMA1, IMA2, IMA3, IMA4, IMA5, IMD3, IMG1, IMG2, INA22, IOC3, IPA1, IQG1, IRC24, IRC4, IRC5, IRC6, IRC7, ISC10, IST3, ITR1, IZH1, JEN1, JIP4, JIP5, JJJ2, JLP1, KAP114, KAR9, KEG1, KEL3, KGD4, KIN82, KIP3, KOG1, KRE1, KRE2, KRE28, KRE6, KRE9, KRI1, LAA1, LAG1, LAM5, LCB1, LCD1, LDB18, LDS1, LEM3, LEU3, LIP1, LNP1, LOS1, LOT5, LPP1, LRG1, LSB3, LSB5, LSC2, LSM3, LSM5, LTE1, LYS1, LYS9, MAG2, MAK10, MAK16, MAL12, MAL13, MAL31, MAL32, MAN2, 46166, MBB1, MCH2, MCH4, MCM10, MCO14, MDH2, MDJ2, MDL2, MDM1, MDM10, MDM31, MDM34, MDY2, MED7, MES1, MET1, MET10, MET16, MET28, MFG1, MGA1, MGA2, MGM101, MGR1, MHT1, MIA40, MID1, MIG2, MIL1, MIN10, MLC2, MLP1, MLP2, MMP1, MMS1, MMT2, MND2, MNL1, MNN11, |

| | |
|---|---|
| | MNN4, MNN5, MNT2, MNT4, MOB2, MON2, MPC2, MPC3, MPH3, MPP6, MRP2, MRP4, MRPL10, MRPL20, MRPL4, MRPL50, MRPS12, MRPS18, MRS1, MRS6, MRX15, MRX20, MRX6, MSA2, MSB3, MSB4, MSC1, MSH3, MSL1, MSN1, MSO1, MST1, MTC3, MTC7, MTD1, MTG2, MTM1, MTO1, MTR2, MTW1, MUP3, MVD1, MXR2, MYG1, MYO4, MZM1, NAB6, NBL1, NBP1, NCA2, NCE101, NCE102, NCS6, NDD1, NDI1, NEW1, NFT1, NGK1, NGL3, NIF3, NIP1, NMD3, NOG2, NOP19, NOP6, NOP8, NPR2, NPR3, NRE1, NSL1, NTG1, NUC1, NUD1, NUP133, NUP188, NUT2, NVJ1, OAF1, OCA4, OCA5, OM45, OMA1, OPI1, OPT1, OPT2, ORC4, OST4, OST5, OSW7, OTU1, OTU2, OXA1, OXP1, OYE2, PAB1, PAC11, PAD1, PAN1, PAN6, PAP2, PAT1, PAU1, PAU10, PAU11, PAU12, PAU13, PAU14, PAU15, PAU18, PAU19, PAU2, PAU20, PAU21, PAU24, PAU4, PAU6, PBI1, PCC1, PCK1, PCL1, PCM1, PDA1, PDE1, PDE2, PDI1, PDP3, PDR18, PEA2, PES4, PET122, PET494, PEX1, PEX11, PEX18, PEX2, PEX21, PEX22, PEX29, PEX6, PFA3, PFD1, PFK1, PFK27, PFS1, PFS2, PGA3, PGU1, PHA2, PHO11, PHO12, PHO13, PHO4, PHO8, PHO84, PHO90, PHR1, PIN3, PIP2, PIR5, PKH1, PLC1, PLM2, PML39, PMT2, PMT4, POL5, POP2, POP3, POP5, PPG1, PPM2, PPX1, PRB1, PRC1, PRE10, PRE4, PRE5, PRE8, PRI1, PRM1, PRP16, PRP19, PRP21, PRP4, PRP45, PRP8, PRR2, PRS3, PRT1, PSE1, PSF3, PSK1, PSP1, PTA1, PTC6, PTH1, PTH4, PTK1, PTP1, PTR2, PTR3, PUF6, PUG1, PUL3, PUL4, PUP2, PUS4, PWP2, PWR1, PXA2, PXL1, PXP3, PXR1, PZF1, QCR2, QCR6, QCR7, QCR8, RAD10, RAD17, RAD18, RAD2, RAD23, RAD24, RAD3, RAD4, RAI1, RBA50, RBD2, RBG1, RBH1, RCF2, RCY1, RDR1, RET2, REV7, RFA2, RFA3, RFC3, RGD2, RGD3, RHO1, RIB3, RIB4, RIE1, RIF2, RIM101, RIM15, RIM21, RIM4, RIX1, RMD6, RMD8, RME3, RML2, RMR1, RNH70, RNP1, RNQ1, ROF1, ROG3, RPC82, RPD3, RPF2, RPH1, RPL12A, RPL16A, RPL17B, RPL18A, RPL18B, RPL22B, RPL25, RPL29, RPL2A, RPL36B, RPL37B, RPL39, RPL40A, RPL40B, RPL6B, RPL8A, RPL8B, RPN10, RPN12, RPO26, RPO41, RPR2, RPS12, RPS14B, RPS19A, RPS19B, RPS1A, RPS20, RPS22A, RPS4A, RPS4B, RRI1, RRI2, RRP40, RRP7, RRT12, RRT5, RSA4, RSC6, RSC8, RSC9, RSM19, RSM28, RTC1, RTF1, RTG2, RTT102, RUF21, RUF22, RUF23, RVB2, SAC1, SAM2, SAM3, SAM4, SAP155, SAP4, SAW1, SAY1, SBP1, SCC4, SCH9, SCP1, SCW10, SCW4, SDA1, SDD1, SDH2, SDH7, SDS22, SDT1, SEC11, SEC12, SEC15, SEC20, SEC21, SEC23, SEC39, SEC53, SEC65, SED4, SEN1, SET2, SET5, SGV1, SHE10, SHR3, SHR5, SHS1, SHU1, SIP2, SIR1, SIR3, SIS2, SIT1, SKG1, SKI3, SKI8, SKM1, SKN7, SKP2, SLD5, SLF1, SLH1, SLN1, SLO1, SMC2, SMF1, SMM1, SMN1, SMT3, SMX3, SNA2, SNC1, SNF1, SNF12, SNF6, SNM1, SNO2, SNO4, SNR191, SNR40, SNR49, SNR53, SNR64, SNR67, SNR68, SNR80, SNR84, SNR85, SNX3, SNZ2, SOL1, SOL2, SOL3, SOL4, SOM1, SOP4, SOR1, SPC72, SPC97, SPE1, SPG3, SPI1, SPO11, SPO7, SPS1, SPS2, SPT15, SPT16, SPT2, SPT20, SQT1, SRB8, SRL3, SRO9, SRP40, SRP68, SRY1, SSA1, SSB1, SSK2, SSK22, SSL2, SSO1, SSP1, SST2, STB1, STB5, STE20, STE50, STE6, STL1, STS1, SUE1, SUF6, SUI3, SUP6, SUP61, SUT390, SUT532, SVP26, SWC3, SWE1, SWI3, SWP82, SYN8, TAD1, TAF1, TAF13, TAF8, TAH1, TAN1, TCA17, TDA11, TDA5, TDA6, TDA8, TDA9, TGL3, TGL4, THI11, THI13, THI21, THI5, THP2, THR4, TIF3, TIM22, TIM23, TMA108, TMT1, TNA1, TOG1, TOH1, TOR2, TOS6, TPD3, TPK1, TPM2, TPO3, TRF5, TRM11, TRM112, TRM13, TRN1, TRP3, TRT2, TRX3, TRZ1, TSL1, TSR2, TUB1, TUB2, TUB3, TUP1, TUS1, TVP38, TVS1, UBA1, UBI4, UBP11, UBP12, UBP15, UBP3, UBP5, UFO1, UGA4, ULI1, UPA1, UPA2, URA1, URA5, URC2, URN1, USV1, UTP14, UTP9, UTR2, UTR4, UTR5, VAM7, VAN1, VBA5, VEL1, VHS2, VID30, VIK1, VLD1, VMA11, VMA6, VMA8, VMR1, VNX1, VPS13, VPS3, VPS4, VPS501, VPS52, VPS60, VPS68, VPS72, VPS9, VRG4, VTH1, VTH2, VTS1, WHI4, WSC2, WSC4, YAH1, YAP1801, YAP1802, YAP3, YAP5, YAR1, YBT1, YCT1, YEF1, YGK3, YIH1, YKT6, YLF2, YME2, YOR1, YPD1, YPS6, YPT1, YPT11, YPT32, YRA2, YRF1-2, YRF1-3, YRF1-4, YRF1-5, YRF1-6, YRF1-7, YRF1-8, YTA7, YVH1, ZDS2, ZEO1, ZIM17, ZIP2, ZNF1, ZNG1, ZOD1, ZPS1, ZRG17, ZRT1, ZUO1, UPA2, URA1, URA5, URC2, URN1, USV1, UTP14, UTP9, UTR2, UTR4, UTR5, VAM7, VAN1, VBA5, VEL1, VHS2, VID30, VIK1, VLD1, VMA11, VMA6, VMA8, VMR1, VNX1, VPR1, VPS13, VPS3, VPS4, VPS501, VPS52, VPS60, VPS68, VPS72, VPS8, VPS9, VRG4, VTH1, VTH2, VTS1, WHI4, WSC2, WSC3, WSC4, YAH1, YAP1801, YAP1802, YAP3, YAP5, YAR1, YAT1, YBT1, YCT1, YEF1, YGK3, YIH1, YKT6, YLF2, YME2, YOR1, YPD1, YPS6, YPT1, YPT11, YPT32, YRA2, YRF1-1, YRF1-2, YRF1-3, YRF1-5, YRF1-6, YRF1-7, YRF1-8, YTA7, YVH1, ZDS2, ZEO1, ZIM17, ZIP2, ZNF1, ZNG1, ZOD1, ZPS1, ZRG17, ZRT1, ZUO1 |

| | |
|---|---|
| gal5 | AAD14, AAD4, ACM1, ACO2, ACS1, ACT1, ADD66, ADE57, ADH2, ADH4, ADH6, ADK2, ADY3, AFG1, AGA1, AGE1, AIF1, AIM17, AIM18, AIM2, AIM46, AIM6, ALD4, ALR1, ALR2, AMF1, ANK1, ANP1, AOS1, APA2, API2, APL6, APM1, APM2, AQY1, AQY2, ARC35, ARG8, ARG81, ARN1, ARN2, ATF1, ATG10, ATG13, ATG22, ATG32, ATG36, ATG41, ATG7, ATP11, ATP15, ATR1, AVT2, AXL2, AYT1, BAS1, BAT1, BAT2, BBP1, BCK2, BDH1, BDH2, BEM2, BET2, BGL2, BIK1, BIO2, BIO3, BIO4, BIO5, BMH1, BNA6, BOL1, BOL3, BRE4, BRE5, BRR2, BRR6, BSC5, BSC6, BSP1, BUD32, BUL2, BUR6, BXI1, CAB1, CAB4, CAC2, CAF40, CAN1, CAR2, CBP1, CBP2, CBT1, CCA1, CCT2, CDC14, CDC19, CDC24, CDC26, CDC33, CDC6, CHD1, CIN8, CLA4, CLN2, CLN3, CNE1, CNN1, COF1, COG3, COQ2, COQ5, COQ6, COS1, COS10, COS12, COS5, COS7, COS8, COX14, CRG1, CSA1, CSE1, CSS1, CSS3, CTF8, CTK3, CTR2, CTR9, CUB1, CUE4, CUS2, CWC21, CWC22, CYC3, CYC7, DAL5, DAL82, DAN1, DAN4, DAT1, DBP6, DBP8, DCP1, DFP4, DIA1, DIF1, DIG2, DIM1, DIP5, DLD3, DMC1, DNF1, DOA1, DOC1, DPB2, DPM1, DSE4, DSF1, DUG1, DUR3, EAP1, ECM1, ECM25, ECM29, ECM32, ECM34, ECM7, ECO1, EFG1, EFM1, EGD2, EGO4, EGT2, ELO1, ELP6, EMC1, EMC3, EMC4, EMI1, EMI2, EMP47, EMW1, ENB1, ENO1, ENO2, ENT4, ERG13, ERG9, ERJ5, ERO1, ERR2, ERR3, ERV29, ERV46, ESF2, ESL2, ETP1, EUG1, FAU1, FDC1, FDH1, FET4, FET5, FEX1, FEX2, FIG4, FIT1, FIT2, FIT3, FKS3, FLC2, FLO1, FLO10, FLO9, FLX1, FMO1, FMP10, FMP27, FMP32, FMP45, FOL2, FPK1, FPR2, FPR4, FPS1, FRD1, FRE3, FRE4, FRE5, FRE6, FRE7, FUM1, FUN12, FUN19, FZF1, GAB1, GAL4, GAS1, GAS4, GCG1, GCN5, GCS1, GCV3, GDA1, GDB1, GDH1, GEM1, GEX2, GFD2, GID7, GIM5, GIN4, GIP4, GLC8, GLK1, GLY1, GMC1, GMC2, GND1, GND2, GNP1, GON7, GOS1, GPB1, GPB2, GPH1, GPI16, GRC3, GRE2, GRH1, GRX1, GRX2, GRX4, GTA1, GTR1, GTT2, GUD1, GUS1, GUT1, GYP5, GYP7, HAP2, HAT2, HBT1, HDA3, HFI1, HFM1, HHY1, HIS2, HIS4, HLR1, HMG2, HMS2, HO, HOL1, HPA2, HPA3, HRP1, HRT1, HSP31, HSP32, HSU1, HUA1, HUT1, HXK1, HXK2, HXT11, HXT13, HXT14, HXT15, HXT16, HXT17, HXT8, HXT9, IES6, IKI1, IMA1, IMA2, IMA3, IMA4, IMA5, IRC4, IRC5, IRC6, IRC7, ISC10, ITR1, IZH1, JEN1, JIP5, JLP1, KAP114, KAR9, KEG1, KEL3, KGD4, KOG1, KRE1, KRE2, KRE28, KRI1, LAA1, LAM5, LCD1, LDB18, LEM3, LEU3, LNP1, LOS1, LPP1, LRG1, LSB3, LSB5, LSM3, LYS9, MAK10, MAL12, MAL13, MAN2, MBB1, MCH4, MCM10, MCO14, MDH2, MDJ2, MDL2, MDM1, MDM31, MED7, MES1, MET10, MET16, MFG1, MGM101, MGR1, MHT1, MID1, MIN10, MLC2, MLP1, MLP2, MMP1, MMS1, MNL1, MNN4, MNN5, MNT2, MNT4, MON2, MPH3, MRP2, MRPL20, MRPL4, MRPS12, MRPS18, MRS6, MRX15, MRX20, MRX6, MSB3, MSC1, MSN1, MSO1, MTC3, MTG2, MTM1, MTO1, MTW1, MUP3, MVD1, MXR2, MYG1, MZM1, NAB6, NBL1, NBP1, NCE101, NDD1, NDI1, NFT1, NGK1, NGL3, NIP1, NMD3, NOG2, NOP19, NOP8, NPR2, NPR3, NUC1, NUD1, NUP188, NUT2, NVJ1, OAF1, OCA5, OM45, OMA1, OPI1, OPT1, OPT2, ORC4, OST4, OST5, OSW7, OTU1, OTU2, OXA1, OXP1, OYE2, PAB1, PAC11, PAD1, PAN6, PAU1, PAU11, PAU12, PAU13, PAU14, PAU15, PAU18, PAU19, PAU2, PAU20, PAU21, PBI1, PCC1, PCK1, PCL1, PCM1, PDA1, PDE1, PDI1, PDP3, PDR18, PET122, PET494, PEX11, PEX2, PEX22, PEX29, PEX6, PFA3, PFK27, PFS1, PFS2, PGA3, PGU1, PHA2, PHO11, PHO12, PHO13, PHO4, PHO8, PHO84, PHO90, PHR1, PKH1, PLC1, PLM2, PML39, PMT4, POF1, POL5, POP2, POP5, PPM2, PPX1, PRB1, PRE4, PRE5, PRE8, PRP16, PRP19, PRP21, PRP4, PRP45, PRS3, PSE1, PSF3, PSP1, PTA1, PTH1, PTK1, PTP1, PTR2, PTR3, PUF6, PUG1, PUL3, PUL4, PUP2, PUS4, PXL1, PXP3, PXR1, PZF1, QCR2, QCR6, QCR7, RAD10, RAD2, RAD23, RAD24, RAD3, RAD4, RAI1, RBA50, RBD2, RBG1, RCY1, RDR1, RET2, REV7, RFA2, RFC3, RGD2, RGD3, RHO1, RIB3, RIB4, RIF2, RIM101, RIM21, RIM4, RIX1, RMD6, RMD8, RME3, RML2, RMR1, RNH70, RNP1, ROF1, RPC82, RPD3, RPH1, RPL12A, RPL18A, RPL18B, RPL25, RPL29, RPL2A, RPL36B, RPL37B, RPL39, RPL40A, RPL40B, RPL6B, RPL8A, RPL8B, RPN10, RPN12, RPO26, RPS12, RPS14B, RPS19A, RPS19B, RPS1A, RPS20, RPS22A, RPS4A, RPS4B, RRI2, RRP40, RRP7, RRT5, RSC8, RSC9, RSM19, RSM28, RTC1, RTF1, RTG2, RTT102, RUF21, RUF23, SAC1, SAM2, SAM3, SAM4, SAP155, SAP4, SAY1, SBP1, SCH9, SCW10, SCW4, SDD1, SDH2, SDH7, SEC15, SEC20, SEC21, SEC23, SEC39, SEC53, SEC65, SET5, SGV1, SHE10, SHS1, SHU1, SIR1, SIR3, SIT1, SKG1, SKI3, SKN7, SKP2, SLD5, SLF1, SLH1, SLN1, SLO1, SMC2, SMF1, SMN1, SMT3, SMX3, SNA2, SNC1, SNF6, SNO4, SNR191, SNR40, SNR49, SNR53, SNR64, SNR67, SNR68, SNR80, SNR84, SNR85, SOL1, SOM1, SOP4, SOR1, SPC72, SPC97, SPG3, SPO11, SPS1, SPS2, SPS22, SPT2, SPT20, SRL3, SRO9, SRP40, SRY1, SSB1, SSL2, SSP1, SST2, STB1, STB5, STE20, STE50, STE6, STL1, SUF6, SUP6, SUT390, SUT532, SVP26, SWE1, SWP82, TAD1, TAF1, TAF13, TAF8, TAN1, TCA17, TDA8, TGL3, TGL4, THI13, THI21, THI5, TIF3, TMA108, TMT1, TNA1, TOG1, TOR2, TOS6, TPM2, TRF5, TRM11, TRM112, TRM13, TRP3, TRT2, TSL1, TUB2, TUB3, TVP38, UBA1, UBI4, UBP11, UBP12, UBP15, ULI1, UPA1, UPA2, URA1, URA5, URC2, UTP14, UTP9, UTR2, UTR4, VAN1, VBA5, VEL1, VHS2, VID30, VIK1, VMA6, VMA8, VMR1, VNX1, VPS13, VPS3, VPS4, VPS52, VPS60, VPS68, VPS72, VPS9, VTH1, WHI4, WSC4, YAH1, YAP3, YBT1, YCT1, YEF1, YGK3, YLF2, YOR1, YPD1, YPT1, YPT11, YRA2, YRF1-2, YRF1-3, YRF1-4, YRF1-5, YRF1-6, YRF1-7, YRF1-8, YTA7, ZDS2, ZIM17, ZIP2, ZOD1, ZPS1, ZRG17, ZRT1, ZUO1 |

| | |
|---|---|
| **gal6** | AAD10, AAD14, AAD4, ABP1, ACM1, ACO2, ACS1, ACT1, ADD66, ADE57, ADH4, ADH6, ADK2, ADY3, AFG1, AGA1, AGE1, AGP3, AIF1, AIM17, AIM18, AIM2, AIM46, AIM6, ALD4, ALR1, ALR2, AMF1, ANK1, AOS1, APA2, API2, APL6, APM1, APM2, AQY1, AQY3, ARC35, ARG8, ARN1, ATF1, ATG13, ATG27, ATG32, ATG36, ATG41, ATG7, ATP11, ATP12, ATP15, ATR1, AVT2, AXL2, BAS1, BAT1, BAT2, BBP1, BCK2, BDH1, BDH2, BEM2, BET2, BGL2, BIO2, BIO3, BIO4, BIO5, BMH1, BNA6, BOL1, BOL2, BOL3, BRE4, BRE5, BRR2, BRR6, BSC5, BSC6, BSP1, BUD32, BUL2, BUR6, BXI1, CAB1, CAB4, CAC2, CAF40, CAN1, CAR2, CBP1, CBP2, CBT1, CCA1, CCT2, CDC13, CDC14, CDC19, CDC24, CDC26, CDC33, CDC6, CHD1, CIN2, CIN8, CLA4, CLN2, CLN3, CNE1, CNN1, COG1, COG3, COQ2, COQ5, COQ6, COS1, COS10, COS4, COS7, COX14, CRG1, CSA1, CSE1, CSM1, CSS1, CSS3, CTF8, CTK3, CTR2, CTR9, CUB1, CUE4, CUS2, CWC21, CWC22, CYC3, DAK2, DAL1, DAL2, DAL3, DAL4, DAL5, DAL7, DAL81, DAL82, DAN1, DAN4, DAT1, DBP6, DBP8, DCG1, DCP1, DDI2, DFP4, DIA1, DIF1, DIG2, DIM1, DIP5, DJP1, DMC1, DNF1, DOA1, DOC1, DPB2, DPM1, DSE4, DSN1, DTD1, DUG1, DUR3, EAP1, ECM1, ECM25, ECM29, ECM32, ECM34, ECM7, EDC1, EFG1, EFM1, EGD2, EGH1, EGO4, EGT2, ELO1, ELP6, EMC3, EMC4, EMI1, EMI2, EMP47, EMW1, ENB1, ENO1, ENO2, ENV7, ERG13, ERG9, ERJ5, ERO1, ERR2, ERR3, ERV29, ERV46, ESF2, ESL2, EUG1, FAS2, FAU1, FDC1, FDH1, FET4, FET5, FEX1, FEX2, FIG2, FIG4, FIT1, FIT2, FIT3, FKS3, FLC2, FLO1, FLO10, FLO11, FLO9, FLX1, FMO1, FMP10, FMP27, FMP32, FMP45, FOL2, FPK1, FPR2, FPR4, FRD1, FRE3, FRE4, FRE5, FRE7, FUM1, FUN12, FUN19, FZF1, GAB1, GAL4, GAS1, GAS4, GAT4, GCG1, GCS1, GCV3, GDB1, GDH1, GDH3, GEM1, GEX2, GIN4, GLC8, GLY1, GMC1, GMC2, GND1, GND2, GNP1, GON7, GOS1, GPB1, GPB2, GPI16, GRE2, GRH1, GRX2, GRX4, GTR1, GUD1, GUS1, GUT1, GYP5, GYP7, HAP2, HAT2, HBT1, HDA3, HFI1, HFM1, HHY1, HLR1, HMG2, HMS2, HO, HOL1, HPA2, HPA3, HRP1, HRT1, HSP31, HSP32, HSP82, HUA1, HUT1, HXK1, HXK2, HXT11, HXT13, HXT14, HXT15, HXT16, HXT17, HXT8, HXT9, ICR1, IES6, IKI1, IMA1, IMA2, IMA4, IMA5, INA22, IQG1, IRC24, IRC4, IRC5, IRC6, IRC7, ISC10, IST3, ITR1, IZH1, JEN1, JIP4, JIP5, KAP114, KAR9, KEG1, KEL3, KGD4, KIP3, KOG1, KRE1, KRE2, KRE28, KRE9, KRI1, LAA1, LAM5, LCD1, LEM3, LEU3, LNP1, LOS1, LPP1, LRG1, LSM3, LYS1, LYS9, MAK10, MAN2, MBB1, MCM10, MCO14, MDH2, MDJ2, MDL2, MDM1, MDM31, MDM34, MED7, MES1, MET10, MET16, MET28, MFG1, MGA2, MGM101, MID1, MIG2, MIN10, MLC2, MLP1, MLP2, MMS1, MND2, MNL1, MNN11, MNN4, MNN5, MNT2, MNT4, MON2, MPH3, MRP2, MRPL20, MRPL4, MRPS12, MRPS18, MRS1, MRS6, MRX15, MRX20, MRX6, MSB3, MSC1, MSL1, MSO1, MTC3, MTG2, MTM1, MTO1, MTW1, MUP3, MVD1, MYG1, MZM1, NAB6, NBL1, NBP1, NCE101, NCS6, NDD1, NDI1, NFT1, NGK1, NGL3, NIF3, NIP1, NMD3, NOG2, NOP8, NPR2, NPR3, NRE1, NSL1, NUC1, NUD1, NUP188, NUT2, NVJ1, OAF1, OCA5, OM45, OMA1, OPI1, OPT1, OPT2, OST4, OST5, OSW7, OTU1, OXA1, OXP1, OYE2, PAB1, PAC11, PAD1, PAN1, PAN6, PAU11, PAU14, PAU15, PAU18, PAU19, PAU2, PAU20, PAU21, PBI1, PCC1, PCK1, PCL1, PCM1, PDA1, PDE1, PDP3, PDR18, PET122, PET494, PEX11, PEX2, PEX22, PEX29, PEX6, PFA3, PFD1, PFK27, PFS1, PFS2, PGA3, PGU1, PHA2, PHO11, PHO12, PHO13, PHO4, PHO8, PHO84, PHO90, PHR1, PKH1, PLC1, PLM2, PML39, POL5, POP2, POP5, PPM2, PPX1, PRB1, PRE4, PRE5, PRI1, PRP16, PRP21, PRP4, PRP45, PSE1, PSF3, PSP1, PTA1, PTH1, PTK1, PTP1, PTR2, PTR3, PUF6, PUG1, PUL3, PUL4, PUP2, PUS4, PWR1, PXL1, PXP3, PXR1, PZF1, QCR2, QCR6, QCR7, RAD17, RAD2, RAD24, RAD3, RAD4, RAI1, RBA50, RBD2, RBG1, RBH1, RCY1, RDR1, RET2, REV7, RFA2, RFA3, RFC3, RGD2, RGD3, RHO1, RIB3, RIB4, RIF2, RIM101, RIM21, RIM4, RIX1, RMD8, RME3, RML2, RMR1, RNH70, ROF1, RPC82, RPD3, RPH1, RPL12A, RPL17B, RPL18B, RPL25, RPL29, RPL2A, RPL36B, RPL37B, RPL39, RPL40A, RPL40B, RPL6B, RPL8A, RPN10, RPN12, RPO26, RPR2, RPS12, RPS14B, RPS19B, RPS1A, RPS22A, RPS4A, RPS4B, RRI1, RRP40, RRT5, RSC8, RSC9, RSM19, RSM28, RTC1, RTF1, RTG2, RTT102, RUF23, RVB2, SAC1, SAM2, SAM3, SAM4, SAP155, SAP4, SAY1, SBP1, SCH9, SCW4, SDD1, SDH7, SDT1, SEC11, SEC15, SEC20, SEC21, SEC23, SEC39, SEC53, SEC65, SET5, SHE10, SHS1, SIR1, SIR3, SIT1, SKG1, SKI3, SKI8, SKN7, SKP2, SLD5, SLF1, SLH1, SLN1, SLO1, SMC2, SMF1, SMN1, SMT3, SMX3, SNA2, SNF1, SNF6, SNM1, SNO2, SNO4, SNR191, SNR40, SNR49, SNR53, SNR64, SNR67, SNR68, SNR80, SNR84, SNR85, SNZ2, SOL1, SOM1, SOP4, SOR1, SPC72, SPC97, SPG3, SPO11, SPS1, SPS2, SPT2, SPT20, SQT1, SRL3, SRP40, SRP68, SSB1, SSL2, SSO1, SSP1, SST2, STB1, STB5, STE6, STL1, STS1, SUF6, SUI3, SUP6, SUT390, SUT532, SVP26, SWE1, SWI3, SWP82, TAD1, TAF1, TAF8, TAN1, TCA17, TDA8, TGL3, TGL4, THI11, THI13, THI21, THI5, TIM22, TMA108, TMT1, TNA1, TOG1, TOR2, TOS6, TPM2, TRF5, TRM11, TRM112, TRM13, TRP3, TRT2, TUB3, TVP38, UBA1, UBP11, UBP12, UPA1, UPA2, URA1, URA5, URC2, UTP9, VAM7, VAN1, VBA5, VEL1, VHS2, VID30, VIK1, VLD1, VMA11, VMA6, VMA8, VMR1, VNX1, VPS3, VPS4, VPS52, VPS60, VPS68, VPS72, VRG4, VTH1, WHI4, WSC4, YAH1, YAP5, YAR1, YGK3, YOR1, YPD1, YPT1, YPT11, YPT32, YRA2, YRF1-2, YRF1-5, YRF1-7, YTA7, YVH1, ZDS2, ZIM17, ZIP2, ZNF1, ZPS1, ZRG17, ZRT1, ZUO1 |

**900 generations**

| Replicate | Gene list |
|---|---|
| glu1 | AIM18, AIM20, AIM46, ARP7, ATG23, ATG33, ATG44, ATP11, BAT1, BCD1, BDF1, BER1, BLS1, BNR1, BRL1, BUD8, BXI1, CAF40, CAR2, CNA1, COA1, CRN1, CSR1, CTF3, CTF8, CTR2, CTR3, CUS2, DAL82, DIF1, DUS3, DUS4, ECM30, EGD2, EMW1, ENO2, ERC1, ERG9, ERV2, FBP1, FMO1, FTR1, GAG1, GLN1, GND1, GPI16, GUT2, HPF1, HXT14, IKI1, IKI3, ILV5, IMD3, IMP21, INA1, IRC16, KOG1, KRI1, LNP1, LSM3, LSM5, LUG1, MAG2, MDM31, MID1, MNL1, MRPL4, MRPS18, MSB3, MSC7, NAM2, NBL1, NCP1, NEL1, NIT1, NIT3, NVJ1, ORM2, OXA1, OYE2, PCL1, PEA2, PET122, PFS1, PFS2, PHA2, PIH1, POT1, PPX1, PSY3, PTH1, PUN1, PUS4, PUT2, RFA2, RFC3, RGD3, RIM21, RIX1, ROF1, RPL18B, RPL31B, RPN10, RPS19B, RPS4B, RRD1, RRF1, RSC2, RVS167, SCC4, SCH9, SCS3, SEC21, SEC61, SEI1, SEN1, SET5, SFP1, SKN7, SKP2, SMC6, SMN1, SMU2, SNR40, SOA1, SPI1, SPO24, SPT15, SRB2, SSP1, STB1, STB5, STP3, SUC2, SUF6, SVP26, SWC7, TAL1, TOS6, TRF5, TSR2, UBP3, UBP7, UTP21, UTP9, VIP1, VMA10, VMA13, VNX1, VPR1, VPS36, YPT11, ZIM17 |
| glu2 | APS1, ATG26, ATG38, CCC1, CDC123, CLB4, COA4, COQ9, CPR6, CRR1, DFR1, DPH5, EMG1, ENT2, FRE1, HCR1, HMX1, HRD3, IDP2, IRC13, MAS1, MDL1, MMR1, MSC3, MSS51, NCW2, NMT1, PBA1, PEX13, PNP1, PUS5, PWP1, QRI5, RFX1, RNH203, RPL33B, RPL37A, RPS31, SAM1, SEC13, SHH4, SKG3, SNR17A, SWI6, TAG1, TFS1, TOS4, TUB4, UPS1, UPS2, VTA1, YKE2 |
| glu3 | ACL4, ADA2, ADE8, ADK1, ADR1, AFR1, AGE1, AHA1, AIM20, AIM7, AKR1, ALT2, AMD2, APA2, APC4, API2, APT2, ARG82, ARH1, ARO1, ARO10, ARO80, ARP10, ARX1, ASP1, ATC1, ATG44, ATO3, ATP17, ATP22, ATP5, BCP1, BCS1, BFR2, BMH2, BNA7, BNR1, BTT1, BUD8, CAB1, CAB5, CAD1, CBS2, CCC2, CDC1, CDC37, CDC40, CFT1, CHL4, CIA1, CIN10, CMI8, CNL1, COA1, COI1, COQ4, COX20, COX26, CPP2, CPR1, CPR5, CRF1, CSN9, CSR1, CTA1, CTH1, CTS2, CWC15, CWC21, CYM1, DAD4, DFM1, DIG2, DIN7, DIT1, DIT2, DNF2, DOA4, DON1, DOP1, DOS2, DOT1, DPB4, DPL1, DPP1, DXO1, DYN2, EAF1, EBS1, ECM11, ECM18, EFT1, EKI1, EMI1, EMI2, EMT1, ENT5, ERD1, ERV2, ESC2, ESF1, EUG1, EXG2, FBP1, FCF1, FDC1, FIN1, FIT1, FMN1, FMP16, FMP46, FOB1, FPR2, FRQ1, GCD6, GCN2, GGA1, GIC2, GIN4, GIR2, GIS1, GLN1, GLO2, GMC1, GNP1, GPI11, GPI17, GPI19, GPI8, GRH1, GRX2, GRX3, GTB1, GUK1, HDA2, HEH2, HEL2, HEM1, HIM1, HKR1, HLR1, HMO1, HNT2, HOM2, HPF1, HPR1, HPT1, HRQ1, HSP31, HSP42, HSP78, HST4, HTA1, HTB1, HXT3, HXT7, ILT1, ILV5, INM2, INO2, IPK1, IPT1, IRC16, IRC3, IRC4, ITR1, IVY1, IZH1, JIP4, KEI1, KGD2, KIN1, KRE2, KRE28, LCB2, LCD1, LPP1, LRS4, LSM6, LUG1, LYS4, MAK21, MCM21, MET32, MFA1, MFB1, MGP12, MHR1, MKC7, MNN10, MOR1, MRP1, MRP20, MRPL1, MRPL28, MRPL35, MRPL7, MRPS28, MRX10, MRX14, MRX16, MRX8, MSC2, MSH6, MSN5, MSS116, MSS4, MSW1, MTC5, MTH1, MTQ2, MUS81, MZM1, NAP1, NBP2, NCB2, NGG1, NHX1, NIT1, NIT3, NKP1, NPL3, NSE3, NUM1, NUP42, NVJ3, OCA6, OMS1, ORM2, PAA1, PAC11, PAD1, PAL1, PAM1, PAU11, PCF11, PDC2, PDR15, PDS1, PEP7, PET100, PEX10, PEX29, PEX3, PEX5, PEX7, PFA5, PFU1, PHM6, PHO8, PHO92, PIB1, PKH1, PKH3, PLM2, PLN1, PLP1, PMP3, PMT7, POT1, PPH3, PPM1, PPN1, PPZ2, PRO1, PRP28, PRP3, PRP42, PSP1, PSY3, PUF6, QCR7, RAD30, RAD34, RAD55, RAD9, RAV2, RBA50, REF2, RGA2, RGP1, RIB3, RKM2, RKM4, RLI1, RMD5, RMT2, RNH202, RPA14, RPB7, RPL12B, RPL27B, RPL37B, RPN9, RPP2B, RPS13, RPS17B, RPS18A, RPT3, RQC1, RRG1, RRP1, RRP17, RRP45, RRP8, RSC3, RSM24, RSM28, RTN1, RTR2, RTT103, RUB1, RVB1, SAC3, SAC6, SAC7, SAM2, SAN1, SAS4, SBE2, SCC2, SDC1, SDH4, SDH6, SDH7, SEC1, SEC20, SEC26, SEC61, SEC7, SED1, SEM1, SHB17, SHE9, SHU2, SIP1, SIR4, SIZ1, SKP1, SLD5, SLF1, SLU7, SLY1, SMT3, SMU2, SNA2, SND1, SNF1, SNF11, SNM1, SNR13, SNR84, SNU56, SNX41, SOA1, SPC110, SPC19, SPG3, SPO24, SPO71, SPP41, SPR28, SPS1, SPS2, SPT3, SRB7, SRP101, SSD1, SSF2, SSN2, SSS1, SSY1, STB3, STE14, STE5, STL1, STN1, STP1, STP3, SUC2, SUF3, SUM1, SUP2, SUP35, SUR2, SVF1, SWA2, SWF1, SWI5, SWM1, SWR1, SXM1, SYF1, TAF10, TAF12, TAL1, TCP1, TFB1, TFB3, TFB5, TFC6, THI74, TIF35, TIM11, TLD1, TLG1, TMA64, TMN2, TMS1, TOM1, TPS2, TRM1, TRM82, TRP4, TRR1, TRS120, TRS23, TRS31, TRS85, TSA2, TVP15, TVP23, UBA2, UBC1, UBC13, UBC5, UBP7, UBX5, UGO1, UIP5, UME6, UPC2, UPS3, URC2, URH1, UTH1, UTP4, UTP5, UTP6, VBA4, VHS1, VMA13, VPS3, VPS41, VPS52, VPS60, VPS64, VPS72, VPS74, VTC5, WIP1, XRS2, YAP6, YCF1, YCG1, YFT2, YHP1, YPQ2, YPR1, YPS7, YRA1, YSP2, ZIP1 |

| | |
|---|---|
| **glu4** | ACL4, ACS2, ADA2, ADK1, ADR1, AFR1, AGE1, AHA1, AIM7, AKR1, ALT2, AMD2, APA2, APC4, API2, APS1, APT2, ARE1, ARG82, ARH1, ARO1, ARO10, ARO80, ARP10, ARX1, ASP1, ATC1, ATG26, ATG33, ATG38, ATP17, ATP22, ATP5, BAP3, BCP1, BCS1, BFR2, BMH2, BNA7, BTT1, BUD23, BUD8, CAB1, CAB5, CAD1, CBS2, CCC1, CCC2, CDC1, CDC123, CDC34, CDC37, CDC40, CFT1, CHL4, CIA1, CIS1, CLB4, CMI8, CNL1, COA4, COQ4, COQ9, COX26, CPP2, CPR1, CPR5, CPR6, CRF1, CRR1, CSN9, CSR1, CTA1, CTF3, CTH1, CTS2, CWC15, CWC21, CYM1, DAD4, DBF4, DET1, DIC1, DIG2, DIN7, DNF2, DOA4, DON1, DOP1, DOS2, DOT1, DPB4, DPH5, DPL1, DPP1, DXO1, DYN2, EAF1, EBS1, ECM11, ECM18, EKI1, EMC10, EMG1, EMI1, EMI2, EMT1, ENT2, ENT5, ERD1, ESC2, ESF1, EUG1, EXG2, FBP1, FCF1, FIN1, FIT1, FKH2, FKS1, FMN1, FMP16, FMP46, FOB1, FPR2, FRE1, FRQ1, GAS2, GCD6, GCN2, GGA1, GIC2, GIN4, GIR2, GIS1, GLO2, GMC1, GNP1, GPI11, GPI17, GPI19, GPI8, GRH1, GRX2, GRX3, GTB1, GUK1, GYP6, HCR1, HDA2, HEH2, HEL2, HEM12, HEM13, HIM1, HKR1, HLR1, HMO1, HMX1, HNT2, HOM2, HPF1, HPR1, HRD3, HRQ1, HSP31, HSP42, HSP78, HST4, HTA1, HTB1, HXT3, HXT7, ICS3, IDP2, ILT1, ILV5, IMG1, INM2, INO2, IPK1, IPT1, IRC3, ITR1, IZH1, JIP4, KAP95, KEI1, KGD2, KIN1, KRE2, KRE28, LAT1, LCB2, LCD1, LPP1, LRS4, LSM6, LUG1, MAK21, MAS1, MCM21, MDL1, MET32, MFA1, MFB1, MGP12, MHP1, MHR1, MKC7, MMR1, MNN10, MOR1, MRL1, MRP1, MRPL1, MRPL28, MRPL35, MRPL7, MRPS28, MRX10, MRX14, MRX16, MRX8, MSC2, MSC3, MSH6, MSK1, MSN5, MSS116, MSS4, MSS51, MSW1, MTC5, MTH1, MTQ2, MZM1, NAM2, NAP1, NBP2, NCW2, NGG1, NHA1, NHX1, NIT3, NMT1, NPL3, NSE3, NUM1, NUP42, NVJ3, OCA6, OMS1, OPY2, ORM2, PAA1, PAC11, PAL1, PAM1, PBA1, PCF11, PDC2, PDS1, PEP7, PET100, PEX10, PEX13, PEX29, PEX3, PEX5, PEX7, PFA5, PFU1, PGD1, PHM6, PHO8, PHO92, PIB1, PIB2, PKH1, PKH3, PLM2, PLN1, PLP1, PMP3, PMT7, PNP1, PPH3, PPM1, PPN1, PPZ2, PRO1, PRP28, PRP3, PRP42, PRY1, PRY3, PSP1, PST1, PSY3, PUF6, PUS5, PWP1, QCR7, QRI5, RAD30, RAD34, RAD55, RAD9, RAV2, RBA50, REF2, RFX1, RGA2, RGP1, RIB3, RKM2, RKM4, RKM5, RLI1, RMD5, RMT2, RNH201, RNH202, RNH203, RPA14, RPC11, RPL12B, RPL16B, RPL26A, RPL27B, RPL37A, RPL37B, RPL9B, RPN9, RPP0, RPS13, RPS17B, RPS18A, RPS31, RQC1, RRG1, RRN5, RRP1, RRP17, RRP45, RRP8, RSC3, RSM24, RSM28, RTR2, RTT103, RUB1, RVB1, SAC3, SAC6, SAM1, SAM2, SAN1, SAS4, SBE2, SCC2, SDC1, SDH4, SDH6, SDH7, SEC1, SEC13, SEC20, SEC26, SEC61, SEC7, SEM1, SHB17, SHH4, SHU2, SIP1, SIR4, SKG3, SKP1, SLD5, SLF1, SLS1, SLU7, SLX4, SLY1, SMT3, SNA2, SNF1, SNF11, SNM1, SNR13, SNR84, SNU56, SNX41, SPC110, SPC19, SPG3, SPO71, SPO77, SPP41, SPR28, SPS1, SPS2, SRB7, SRP101, SSD1, SSF2, SSN2, SSS1, SSY1, STB3, STE5, STN1, STP1, STP3, STT3, SUM1, SUN4, SUP2, SUP35, SUR2, SVF1, SWA2, SWF1, SWI5, SWI6, SWM1, SWR1, SYF1, TAF10, TAF12, TAG1, TAL1, TCP1, TEF1, TFB1, TFB3, TFB5, TFC6, TFS1, TGL2, THI74, TIF35, TIM11, TIS11, TKL1, TLD1, TLG1, TMA64, TMN2, TMS1, TOM1, TOM7, TOS4, TPI1, TPS2, TRM1, TRM82, TRP4, TRR1, TRR4, TRS23, TRS31, TRS85, TSA2, TUB4, TVP15, TVP23, UBC1, UBC13, UBC5, UBX5, UGO1, UIP5, UME6, UPC2, UPS1, UPS2, URC2, UTH1, UTP4, UTP6, VBA4, VHS1, VMS1, VPS3, VPS41, VPS52, VPS60, VPS64, VPS72, VPS74, VTA1, VTC5, WIP1, XRS2, YAP6, YAT1, YCF1, YCG1, YFT2, YHP1, YKE2, YOS9, YPQ2, YPR1, YPS7, YSP2, ZIP1 |
| **glu5** | AIM20, ARE1, ATG38, ATG44, AXL2, BNR1, BUD23, BUD8, CAM1, CCC1, CCT2, CDC123, CDC26, CIN10, CIS1, CLB4, COA1, COA3, COA4, CPR6, CRR1, CSR1, DIC1, DIG1, DLS1, EFT1, ENT2, ERG3, ERV2, ESS1, FBP1, FKS1, FLX1, FMP46, FRE1, GAS2, GLN1, GPA2, GYP6, HMX1, HPF1, HRD3, ICS3, ILV3, ILV5, IMG1, IRC16, ISC1, KAP95, LSM8, LUG1, MHP1, MIN10, MNL2, MPM1, MRL1, MRPL8, MSC3, MUS81, NAP1, NCP1, NDC1, NHA1, NIT1, NIT3, OM45, OPY2, ORM2, PGD1, PHO4, PIB2, PLN1, PNP1, POT1, PRY1, PRY3, PSY3, QCR6, QRI5, RAD52, RCF1, REC107, REV7, REX2, RKM5, RPL26A, RPL29, RPL2A, RPP0, RRN5, RRT5, RSC8, RUF23, RVS167, SBH2, SEC13, SEC61, SGF11, SHB17, SHM2, SLS1, SMU2, SNR68, SOA1, SPC25, SPO24, SPO77, SPT8, SRB2, STP3, STT3, SUC2, TAL1, TES1, TFS1, TIP41, TIS11, TKL1, TMA108, TPM2, TRR4, TSA1, TUB4, UBP7, UIP5, USA1, UTH1, VHS2, VMA13, YOX1, ZWF1 |

| | |
|---|---|
| **glu6** | ACE2, ACF2, ACS2, APC2, APJ1, APS1, ARL3, ARP7, ATG26, ATG33, ATG38, BAP3, BNA5, BUD8, CAM1, CCC1, CDC123, CDC34, CDC42, CIN10, CIS1, CKI1, CLB4, COA3, COA4, COQ9, CPR6, CRR1, CSR1, DBF4, DCN1, DET1, DIC1, DIG1, DIP2, DLS1, DPH5, DPH6, DSC3, EFT1, EGD1, ELC1, EMC10, EMG1, ENT2, EOS1, ERC1, ERG3, ERV2, EST1, FAD1, FBP1, FCY1, FKH2, FKS1, FMP46, FRE1, GAS2, GID11, GLN1, GYP6, HCR1, HEM12, HEM13, HER1, HMX1, HOS1, HRD3, HTS1, ICS3, IDP2, ILV5, IMP4, IRC16, ISA2, ISM1, ISU2, KAP95, KIN4, KNH1, KTR6, LAT1, LCL1, LEE1, LGE1, LTP1, LUG1, MAS1, MCP1, MDL1, MED1, MET31, MGE1, MHP1, MKK1, MKS1, MLF3, MMR1, MNL2, MNN9, MPM1, MRL1, MRP10, MRPL8, MRX11, MSC3, MSK1, MSS51, MTF2, MUS81, NAP1, NAT1, NCW2, NHA1, NIS1, NIT3, NMT1, NOP4, NOT5, NPC2, OAZ1, OPY2, ORM2, PBA1, PDC5, PEP3, PET122, PEX13, PGD1, PIB2, PLN1, PNP1, PRM7, PRP11, PRY1, PRY3, PST1, PSY3, PUS5, PUT1, PWP1, QRI5, REX2, RFX1, RKM5, RMP1, RNH201, RNH203, ROX1, RPB8, RPC11, RPL16B, RPL26A, RPL33B, RPL37A, RPL9B, RPP0, RPS31, RRN5, RSC2, RVS167, SAM1, SEC13, SEC61, SGF11, SHB17, SHH4, SHM2, SIR2, SIT4, SKG3, SLS1, SLX4, SMD3, SND1, SNR35, SPE3, SPE4, SPO24, SPO77, SPT8, SRO7, SSN3, STP3, STP4, SUN4, SUP16, SWI6, TAG1, TAL1, TFS1, TGL2, TIP41, TIS11, TKL1, TOM7, TOP3, TOS4, TPI1, TPM1, TRR4, TUB4, UBA3, UBC5, UIP5, UPS1, UPS2, UPS3, USB1, UTH1, VMA13, VMS1, VPS16, VRP1, VTA1, WTM1, WTM2, YKE2, YOS9, YPS1, YPS3, ZRT2, ZWF1 |
| **gal1** | AAD10, AAD14, AAD15, AAD4, ABP1, ACM1, ACO2, ACP1, ACS1, ACT1, ADD66, ADE4, ADE57, ADF1, ADH2, ADH4, ADH6, ADH7, ADK2, ADY3, AFG1, AGA1, AGE1, AGP1, AGP3, AHC2, AIF1, AIM17, AIM2, AIM21, AIM33, AIM6, ALD4, ALO1, ALR1, ALR2, AMF1, ANK1, ANP1, AOS1, APA1, APA2, API2, APL6, APM1, APM2, AQY1, AQY2, AQY3, ARC35, ARG8, ARG81, ARN1, ARN2, ASG7, ASH1, ATF1, ATG10, ATG13, ATG18, ATG22, ATG27, ATG32, ATG36, ATG41, ATG7, ATM1, ATP11, ATP12, ATP15, ATR1, AVT2, AXL2, AYT1, BAS1, BAT2, BBP1, BCK2, BDH1, BDH2, BEM2, BET2, BGL2, BIK1, BIO2, BIO3, BIO4, BIO5, BMH1, BNA6, BOL1, BOL2, BOL3, BRE4, BRE5, BRR2, BRR6, BSC5, BSC6, BSP1, BUD32, BUL2, BUR6, BXI1, CAB1, CAB4, CAC2, CAF120, CAF40, CAN1, CAR2, CBP1, CBP2, CBT1, CCA1, CCT2, CDC13, CDC14, CDC19, CDC23, CDC24, CDC26, CDC33, CDC39, CDC50, CDC6, CHA1, CHD1, CIN2, CIN8, CIS3, CLA4, CLN2, CLN3, CNA1, CNB1, CNE1, CNN1, COF1, COG1, COG3, COQ2, COQ5, COQ6, COS1, COS10, COS12, COS4, COS5, COS6, COS7, COS8, COS9, COX14, CPD1, CPS1, CSA1, CSE1, CSM1, CSS1, CSS2, CSS3, CTK3, CTR2, CTR9, CUB1, CUE4, CUR1, CUS2, CWC21, CWC22, CYC3, CYC7, DAD2, DAK2, DAL1, DAL2, DAL3, DAL4, DAL5, DAL7, DAL81, DAL82, DAN1, DAN4, DAT1, DBP6, DBP8, DCG1, DCP1, DDI2, DFP4, DIA1, DIF1, DIG2, DIM1, DIP5, DJP1, DLD3, DMC1, DNF1, DOA1, DOC1, DPB2, DPH2, DPM1, DSE4, DSF1, DSN1, DTD1, DUG1, DUR3, DYN3, EAP1, ECM1, ECM25, ECM29, ECM30, ECM32, ECM34, ECM7, ECO1, EDC1, EFG1, EFM1, EGH1, EGO2, EGO4, EGT2, ELO1, ELP6, EMA35, EMC1, EMC3, EMC4, EMI1, EMI2, EMP47, EMW1, ENB1, ENO1, ENO2, ENT4, ENV7, ERG13, ERG20, ERG24, ERJ5, ERO1, ERR2, ERR3, ERS1, ERV29, ERV46, ESF2, ESL1, ESL2, ETP1, EUG1, FAB1, FAS2, FAT3, FAU1, FDC1, FDH1, FET4, FET5, FEX1, FEX2, FIG2, FIG4, FIT1, FIT2, FIT3, FKS3, FLC2, FLO1, FLO10, FLO11, FLO5, FLO9, FLX1, FMO1, FMP10, FMP27, FMP32, FMP33, FMP45, FOL2, FPK1, FPR2, FPR4, FPS1, FRA1, FRD1, FRE2, FRE3, FRE4, FRE5, FRE6, FRE7, FRM2, FUB1, FUM1, FUN12, FUN19, FUS1, FYV5, FZF1, GAB1, GAL4, GAS1, GAS4, GAT4, GCG1, GCN5, GCS1, GCV3, GDA1, GDB1, GDH1, GDH3, GEM1, GEX1, GEX2, GFD2, GID7, GIM5, GIN4, GIT1, GLC8, GLK1, GLY1, GMC1, GMC2, GND1, GND2, GNP1, GON7, GOS1, GPB1, GPB2, GPH1, GPI13, GRC3, GRE2, GRH1, GRX1, GRX2, GRX4, GSR1, GTA1, GTR1, GTT1, GTT2, GUD1, GUS1, GUT1, GYP5, GYP7, HAL5, HAP2, HAT2, HBN1, HBS1, HBT1, HCH1, HDA3, HFI1, HFM1, HHY1, HIR3, HIS2, HIS4, HLR1, HMG2, HMLALPHA2, HMRA1, HMRA2, HMS2, HO, HOL1, HPA2, HPA3, HRP1, HRT1, HSP150, HSP31, HSP32, HSP82, HSU1, HUA1, HUT1, HXK1, HXK2, HXT11, HXT13, HXT14, HXT15, HXT16, HXT17, HXT8, HXT9, HYM1, HYP2, HYR1, ICR1, IES6, IGD1, IKI1, IMA1, IMA2, IMA3, IMA4, IMA5, IMD2, IMD3, INA22, IPA1, IQG1, IRC19, IRC24, IRC4, IRC5, IRC6, IRC7, ISA1, ISC10, IST3, ITR1, IZH1, JEN1, JIP4, JIP5, JJJ2, JLP1, KAP114, KAR4, KAR9, KCC4, KEG1, KEL3, KGD4, KIN82, KIP3, KOG1, KRE1, KRE2, KRE28, KRE6, KRE9, KRI1, KRR1, LAA1, LAG1, LAM5, LCB1, LCD1, LDB18, LEM3, LEU2, LEU3, LIP1, LOS1, LOT5, LPP1, LRE1, LRG1, LSB3, LSB5, LSM3, LSM5, LYS1, LYS9, MAK10, MAL12, MAL13, MAL31, MAL32, MAN2, MATALPHA1, 46166, MBB1, MCH2, MCH4, MCM10, MCO14, MDH2, MDJ2, MDL2, MDM1, MDM34, MED7, MES1, MET10, MET16, MET28, MFG1, MGA1, MGA2, MGM101, MGR1, MHT1, MIA40, MIC10, MID1, MIG2, MIN10, MLC2, MLP1, MLP2, MMP1, MMS1, MND2, MNN11, MNN4, MNN5, MNT2, MNT4, MON2, MPH3, MRC1, MRP2, MRP4, MRPL10, MRPL20, MRPL4, MRPS12, MRPS18, MRS1, MRS6, MRX15, MRX20, MRX6, MSB3, MSC1, MSH3, MSL1, MSO1, MST1, MTC3, MTC7, MTD1, MTG2, MTM1, MTO1, MTR2, MTW1, MUP3, MVD1, MXR2, MYG1, MZM1, NAB6, NBP1, NCA2, NCE101, NCE102, NCS6, NDD1, NDI1, NFS1, NFT1, NGK1, NGL3, NIF3, NIP1, NMD3, NOG2, |

| | |
|---|---|
| | NOP19, NOP8, NPR2, NPR3, NRE1, NSL1, NUC1, NUD1, NUP133, NUP188, NUT2, OAF1, OCA4, OCA5, OM45, OMA1, OPI1, OPT1, OPT2, ORC4, OST4, OST5, OSW7, OTU1, OTU2, OXA1, OXP1, OYE2, PAB1, PAC11, PAD1, PAN1, PAN6, PAT1, PAU1, PAU10, PAU11, PAU12, PAU13, PAU14, PAU15, PAU18, PAU19, PAU2, PAU20, PAU21, PAU24, PAU3, PAU4, PAU6, PAU9, PBI1, PBN1, PCC1, PCK1, PCL1, PCM1, PDA1, PDE1, PDI1, PDP3, PDR18, PEA2, PES4, PET122, PET494, PEX1, PEX11, PEX2, PEX22, PEX29, PEX34, PEX6, PFA3, PFD1, PFK27, PFS1, PFS2, PGA3, PGU1, PHA2, PHO11, PHO12, PHO13, PHO4, PHO8, PHO84, PHO90, PHR1, PIN3, PIR5, PKH1, PLC1, PLM2, PML39, PMT4, POF1, POL5, POP2, POP3, POP5, PPM2, PRB1, PRC1, PRD1, PRE4, PRE5, PRE8, PRI1, PRM1, PRP16, PRP19, PRP21, PRP4, PRS3, PSE1, PSF3, PSP1, PTA1, PTC6, PTK1, PTP1, PTR2, PTR3, PUF6, PUG1, PUL3, PUL4, PUP2, PUS4, PWR1, PXA2, PXL1, PXP3, PXR1, PZF1, QCR2, QCR6, QCR7, QCR8, RAD10, RAD17, RAD2, RAD23, RAD24, RAD3, RAD4, RAI1, RBA50, RBD2, RBG1, RBH1, RCY1, RDR1, RDS1, RDT1, RET2, REV7, RFA2, RFA3, RFC3, RGD2, RGD3, RHO1, RIB3, RIB4, RIE1, RIF2, RIM101, RIM21, RIM4, RIX7, RMD6, RMD8, RME3, RML2, RMR1, RNH70, RNP1, RNQ1, ROF1, ROG3, RPC82, RPD3, RPF2, RPH1, RPL12A, RPL16A, RPL17B, RPL18A, RPL18B, RPL25, RPL29, RPL2A, RPL36B, RPL37B, RPL39, RPL40A, RPL40B, RPL6B, RPL8A, RPL8B, RPN12, RPO26, RPR2, RPS12, RPS14B, RPS19A, RPS19B, RPS1A, RPS20, RPS22A, RPS4A, RRP40, RRP7, RRT5, RRT7, RSC8, RSC9, RSM19, RSM28, RTC1, RTF1, RTG2, RTT102, RUF21, RUF22, RUF23, RVB2, SAC1, SAM2, SAM3, SAM4, SAP155, SAP4, SAY1, SBP1, SCC4, SCP1, SCW10, SCW4, SDD1, SDH2, SDH7, SDS22, SDT1, SEC11, SEC15, SEC20, SEC21, SEC23, SEC39, SEC53, SEC65, SET2, SGV1, SHE10, SHS1, SHU1, SIP2, SIR1, SIR3, SIT1, SKG1, SKI3, SKI8, SKP2, SLD5, SLF1, SLH1, SLN1, SLO1, SMC2, SMF1, SMT3, SMX3, SNA2, SNF1, SNF6, SNM1, SNO2, SNO4, SNR191, SNR40, SNR49, SNR53, SNR64, SNR67, SNR68, SNR80, SNR84, SNR85, SNZ2, SOL1, SOL2, SOL4, SOM1, SOP4, SOR1, SPB1, SPC72, SPC97, SPE1, SPG3, SPI1, SPO11, SPS1, SPS2, SPS22, SPT15, SPT2, SPT20, SQT1, SRB8, SRL3, SRO9, SRP40, SRP68, SRY1, SSB1, SSL2, SSO1, SSP1, SST2, STB1, STB5, STE20, STE50, STE6, STL1, STS1, SUE1, SUF6, SUI3, SUP53, SUP6, SUT390, SUT532, SVP26, SWE1, SWI3, SWP82, TAD1, TAF1, TAF13, TAF8, TAN1, TCA17, TDA6, TGL3, TGL4, THI11, THI13, THI21, THI5, THP2, TIF3, TMA108, TMT1, TNA1, TOG1, TOH1, TOR2, TOS6, TPK1, TPM2, TPO1, TPO3, TRF5, TRM11, TRM112, TRM13, TRP3, TRT2, TRX3, TSL1, TSR2, TUB1, TUB2, TUB3, TUP1, TVP38, UBA1, UBI4, UBP11, UBP12, UBP15, UBP3, UFO1, ULI1, UPA1, UPA2, URA1, URA5, URC2, URN1, UTP14, UTR2, UTR4, UTR5, VAC17, VAM7, VAN1, VBA3, VBA5, VEL1, VHS2, VID30, VIK1, VLD1, VMA11, VMA6, VMA8, VMR1, VNX1, VPR1, VPS13, VPS3, VPS4, VPS52, VPS60, VPS68, VPS72, VPS9, VRG4, VTH1, VTH2, WHI4, WSC2, WSC4, YAH1, YAP3, YAP5, YAR1, YBT1, YCT1, YEF1, YGK3, YKT6, YLF2, YME2, YOR1, YPD1, YPS6, YPT1, YPT11, YPT32, YRA2, YRF1-2, YRF1-3, YRF1-4, YRF1-5, YRF1-6, YRF1-7, YRF1-8, YTA7, YVH1, ZDS2, ZIM17, ZIP2, ZNF1, ZOD1, ZPS1, ZRG17, ZRT1, ZUO1 |
| **gal2** | AAD10, AAD14, AAD15, AAD4, ABP1, ACM1, ACO2, ACP1, ACS1, ACT1, ADD66, ADE4, ADE57, ADF1, ADH2, ADH4, ADH6, ADH7, ADK2, ADY3, AFG1, AGA1, AGE1, AGP1, AGP3, AIF1, AIM17, AIM18, AIM2, AIM21, AIM33, AIM46, AIM6, ALD4, ALO1, ALR1, ALR2, AMF1, ANK1, ANP1, AOS1, APA1, APA2, API2, APL6, APM1, APM2, AQY1, AQY2, AQY3, ARC35, ARG8, ARG81, ARN1, ARN2, ASG7, ASH1, ATF1, ATG10, ATG13, ATG18, ATG22, ATG27, ATG32, ATG36, ATG41, ATG7, ATM1, ATP11, ATP12, ATP15, ATR1, AVT2, AXL2, AYT1, BAS1, BAT1, BAT2, BBP1, BCK2, BDH1, BDH2, BEM2, BET2, BGL2, BIK1, BIO2, BIO3, BIO4, BIO5, BMH1, BNA6, BOL1, BOL2, BOL3, BRE4, BRE5, BRR2, BRR6, BSC5, BSC6, BSP1, BUD32, BUL2, BUR6, BXI1, CAB1, CAB4, CAC2, CAF120, CAF40, CAN1, CAR2, CBP1, CBP2, CBT1, CCA1, CCT2, CDC13, CDC14, CDC19, CDC23, CDC24, CDC26, CDC33, CDC39, CDC50, CDC6, CHA1, CHD1, CIN8, CIS3, CLA4, CLN2, CLN3, CNA1, CNB1, CNE1, CNN1, COF1, COG1, COG3, COQ2, COQ5, COQ6, COS1, COS10, COS12, COS4, COS5, COS6, COS7, COS8, COS9, COX14, CPD1, CPS1, CRG1, CSA1, CSE1, CSM1, CSM2, CSS1, CSS2, CSS3, CTF8, CTK3, CTR2, CTR9, CUB1, CUE4, CUR1, CUS2, CWC21, CWC22, CYC3, CYC7, DAK2, DAL1, DAL2, DAL3, DAL4, DAL5, DAL7, DAL81, DAL82, DAN1, DAN4, DAT1, DBP6, DBP8, DCG1, DCP1, DDI2, DFP4, DIA1, DIF1, DIG2, DIM1, DIP5, DJP1, DLD3, DMC1, DNF1, DOA1, DOC1, DPB2, DPH2, DPM1, DSE4, DSF1, DSN1, DTD1, DUG1, DUR3, DYN3, EAP1, ECM1, ECM25, ECM29, ECM30, ECM32, ECM34, ECM7, ECO1, EDC1, EFG1, EFM1, EGD2, EGH1, EGO4, EGT2, ELO1, ELP6, EMA35, EMC1, EMC3, EMC4, EMI1, EMI2, EMP47, EMW1, ENB1, ENO1, ENO2, ENT4, ERG13, ERG20, ERG24, ERG9, ERJ5, ERO1, ERR2, ERR3, ERV29, ERV46, ESF2, ESL1, ESL2, ETP1, EUG1, FAB1, FAT3, FAU1, FDC1, FDH1, FET4, FET5, FEX1, FEX2, FIG2, FIG4, FIT1, FIT2, FIT3, FKH1, FKS3, FLC2, FLO1, FLO10, FLO11, FLO5, FLO9, FLX1, FMO1, FMP10, FMP27, FMP32, FMP33, FMP45, FOL2, FPK1, FPR2, FPR4, FPS1, FRD1, FRE2, FRE3, FRE4, FRE5, FRE6, FRE7, FRM2, FUM1, FUN12, FUN19, FUS1, FYV5, FZF1, GAB1, GAL4, GAS1, GAS4, GAT4, GCG1, GCN5, GCS1, GCV3, GDA1, GDB1, GDH1, GDH3, GEM1, GEX1, GEX2, GFD2, GID7, GIM5, GIN4, GIT1, GLC8, GLK1, |

| | |
|---|---|
| | GLY1, GMC1, GMC2, GND1, GND2, GNP1, GON7, GOS1, GPB1, GPB2, GPH1, GPI16, GRC3, GRE2, GRH1, GRX1, GRX2, GRX4, GSR1, GTA1, GTR1, GTT1, GTT2, GUD1, GUS1, GUT1, GYP5, GYP7, HAL5, HAP2, HAT2, HBN1, HBT1, HCH1, HDA3, HFI1, HFM1, HHY1, HIS2, HIS4, HLR1, HMG2, HMLALPHA2, HMRA1, HMRA2, HMS2, HO, HOL1, HPA2, HPA3, HRP1, HRT1, HSP150, HSP31, HSP32, HSU1, HUA1, HXK1, HXK2, HXT11, HXT13, HXT14, HXT15, HXT16, HXT17, HXT8, HXT9, HYM1, HYP2, HYR1, ICR1, IES6, IGD1, IKI1, IMA1, IMA2, IMA3, IMA4, IMA5, IMD2, IMD3, INA22, IRC24, IRC4, IRC5, IRC6, IRC7, ISC10, IST3, ITR1, IZH1, JEN1, JIP4, JIP5, JJJ2, JLP1, KAP114, KAR4, KAR9, KEG1, KEL3, KGD4, KIN82, KIP3, KOG1, KRE1, KRE2, KRE28, KRE6, KRE9, KRI1, KRR1, LAA1, LAM5, LCB1, LCD1, LDB18, LEM3, LEU3, LIP1, LNP1, LOS1, LOT5, LPP1, LRE1, LRG1, LSB3, LSB5, LSM3, LSM5, LYS1, LYS9, MAK10, MAL12, MAL13, MAL31, MAL32, MAN2, MATALPHA1, 46166, MBB1, MCH2, MCH4, MCM10, MCO14, MDH2, MDJ2, MDL2, MDM1, MDM31, MDM34, MED7, MES1, MET10, MET16, MET28, MFG1, MGA1, MGA2, MGM101, MGR1, MHT1, MIA40, MIC10, MID1, MIG2, MIN10, MLC2, MLP1, MLP2, MMP1, MMS1, MND2, MNL1, MNN11, MNN4, MNN5, MNT2, MNT4, MON2, MPH1, MPH3, MRC1, MRP2, MRP4, MRPL10, MRPL20, MRPL4, MRPS12, MRPS18, MRS1, MRS6, MRX15, MRX20, MRX6, MSB3, MSC1, MSH3, MSL1, MSO1, MST1, MTC3, MTC7, MTG2, MTM1, MTO1, MTR2, MTW1, MUP3, MVD1, MXR2, MYG1, MZM1, NAB6, NBL1, NBP1, NCA2, NCE101, NCS6, NDD1, NDI1, NFT1, NGK1, NGL3, NIF3, NIP1, NMD3, NOG2, NOP19, NOP8, NPR2, NPR3, NRE1, NUC1, NUD1, NUP188, NUT2, NVJ1, OAF1, OCA4, OCA5, OM45, OMA1, OPI1, OPT1, OPT2, ORC4, OST4, OST5, OSW7, OTU1, OTU2, OXA1, OXP1, OYE2, PAB1, PAC11, PAD1, PAN1, PAN6, PAU1, PAU10, PAU11, PAU12, PAU13, PAU14, PAU15, PAU18, PAU19, PAU2, PAU20, PAU21, PAU24, PAU3, PAU4, PAU6, PAU9, PBI1, PBN1, PCC1, PCK1, PCL1, PCM1, PDA1, PDE1, PDI1, PDP3, PDR18, PEA2, PES4, PET122, PET494, PEX1, PEX11, PEX2, PEX22, PEX29, PEX34, PEX6, PFA3, PFD1, PFK27, PFS1, PFS2, PGA3, PGU1, PHA2, PHO11, PHO12, PHO13, PHO4, PHO8, PHO84, PHO90, PHR1, PIN3, PIR5, PKH1, PLC1, PLM2, PML39, POF1, POL5, POP2, POP3, POP5, PPM2, PPX1, PRB1, PRC1, PRD1, PRE4, PRE5, PRE8, PRI1, PRM1, PRP16, PRP19, PRP21, PRP4, PRS3, PSE1, PSF3, PSP1, PTA1, PTH1, PTK1, PTP1, PTR2, PTR3, PUF6, PUG1, PUL3, PUL4, PUP2, PUS4, PWR1, PXA2, PXL1, PXP3, PXR1, PZF1, QCR2, QCR6, QCR7, QCR8, RAD10, RAD17, RAD2, RAD23, RAD24, RAD3, RAD4, RAI1, RBA50, RBD2, RBG1, RBH1, RCY1, RDR1, RDS1, RDT1, RET2, REV7, RFA2, RFA3, RFC3, RGD2, RGD3, RHO1, RIB3, RIB4, RIE1, RIF2, RIM101, RIM21, RIM4, RIX1, RMD6, RMD8, RME3, RML2, RMR1, RNH70, RNP1, RNQ1, ROF1, ROG3, RPC82, RPD3, RPH1, RPL12A, RPL16A, RPL17B, RPL18A, RPL18B, RPL25, RPL29, RPL2A, RPL36B, RPL37B, RPL39, RPL40A, RPL40B, RPL6B, RPL8A, RPL8B, RPN10, RPN12, RPO26, RPR2, RPS12, RPS14B, RPS19A, RPS19B, RPS1A, RPS20, RPS22A, RPS4A, RPS4B, RRP40, RRP7, RRT5, RSC8, RSC9, RSM19, RSM28, RTC1, RTF1, RTG2, RTT102, RUF22, RUF23, SAC1, SAM2, SAM3, SAM4, SAP155, SAP4, SAY1, SBP1, SCC4, SCH9, SCW10, SCW4, SDD1, SDH2, SDH7, SDS22, SDT1, SEC11, SEC15, SEC20, SEC21, SEC23, SEC39, SEC53, SEC65, SET2, SET5, SGV1, SHE10, SHS1, SHU1, SIR1, SIR3, SIT1, SKG1, SKI3, SKI8, SKN7, SKP2, SLD5, SLF1, SLH1, SLN1, SLO1, SMC2, SMF1, SMN1, SMT3, SMX3, SNA2, SNF1, SNF6, SNM1, SNO2, SNO4, SNR191, SNR40, SNR49, SNR53, SNR64, SNR67, SNR68, SNR80, SNR84, SNR85, SNZ2, SOL1, SOL4, SOM1, SOP4, SOR1, SPB1, SPC72, SPC97, SPE1, SPG3, SPI1, SPO11, SPS1, SPS2, SPS22, SPT15, SPT2, SPT20, SQT1, SRL3, SRO9, SRP40, SRY1, SSB1, SSL2, SSP1, SST2, STB1, STB5, STE20, STE50, STE6, STL1, STS1, SUF6, SUP6, SUT390, SUT532, SVP26, SWE1, SWI3, SWP82, TAD1, TAF1, TAF13, TAF8, TAN1, TCA17, TDA6, TDA8, TGL3, TGL4, THI11, THI13, THI21, THI5, THP2, TIF3, TMA108, TMT1, TNA1, TOG1, TOH1, TOR2, TOS6, TPK1, TPM2, TPO3, TRF5, TRM11, TRM112, TRM13, TRP3, TRT2, TSL1, TSR2, TUB1, TUB3, TVP38, UBA1, UBI4, UBP11, UBP12, UBP15, UBP3, UFO1, ULI1, UPA1, UPA2, URA1, URA5, URC2, URN1, UTP14, UTP9, UTR2, UTR4, UTR5, VAC17, VAM7, VAN1, VBA3, VBA5, VEL1, VHS2, VID30, VIK1, VLD1, VMA6, VMA8, VMR1, VNX1, VPR1, VPS13, VPS3, VPS4, VPS52, VPS60, VPS68, VPS72, VPS9, VRG4, VTH1, VTH2, WHI4, WSC2, WSC4, YAH1, YAP3, YAP5, YAT1, YBT1, YCT1, YEF1, YGK3, YKT6, YLF2, YME2, YOR1, YPD1, YPS6, YPT1, YPT11, YPT32, YRA2, YRF1-2, YRF1-3, YRF1-4, YRF1-5, YRF1-6, YRF1-7, YRF1-8, YTA7, YVH1, ZDS2, ZIM17, ZIP2, ZNF1, ZOD1, ZPS1, ZRG17, ZRT1, ZUO1 |

| | |
|---|---|
| gal3 | AAD10, AAD14, AAD15, AAD4, ABP1, ABZ1, ACM1, ACO2, ACP1, ACS1, ACT1, ADD66, ADE4, ADE57, ADF1, ADH2, ADH4, ADH6, ADH7, ADK2, ADY3, AFG1, AGA1, AGE1, AGP1, AGP3, AGX1, AHC2, AIF1, AIM17, AIM2, AIM21, AIM33, AIM6, ALD4, ALG12, ALO1, ALR1, ALR2, AMF1, ANK1, ANP1, APA1, APA2, API2, APL6, APM1, APM2, AQY1, AQY2, AQY3, ARC35, ARE2, ARG8, ARG81, ARN1, ARN2, ASG7, ASH1, ASN1, ATF1, ATG10, ATG15, ATG18, ATG22, ATG23, ATG27, ATG32, ATG36, ATG41, ATG7, ATM1, ATP12, ATP15, ATP18, ATP23, ATR1, AVT2, AXL2, AYT1, BAS1, BAT2, BBP1, BCK2, BDH1, BDH2, BEM2, BGL2, BIK1, BIO2, BIO3, BIO4, BIO5, BMH1, BNA6, BOL1, BOL2, BOL3, BRE4, BRE5, BRF1, BRR2, BRR6, BSC5, BSC6, BSP1, BUD17, BUD32, BUL2, BUR6, CAB1, CAB4, CAC2, CAF120, CAF40, CAN1, CAR2, CBP1, CBP2, CBT1, CCA1, CCT2, CDC13, CDC14, CDC19, CDC23, CDC24, CDC26, CDC33, CDC39, CDC50, CDC6, CHA1, CHD1, CIN2, CIN8, CIS3, CLA4, CLN2, CLN3, CNA1, CNB1, CNE1, CNN1, COF1, COG1, COG3, COQ2, COQ5, COQ6, COS1, COS10, COS12, COS4, COS5, COS6, COS7, COS8, COS9, COX14, CPD1, CPR4, CPR8, CPS1, CSE1, CSM1, CSM2, CSS1, CSS2, CSS3, CTK3, CTR2, CTR9, CUB1, CUE4, CUR1, CUS2, CWC21, CWC22, CYC3, CYC7, DAK2, DAL1, DAL2, DAL3, DAL4, DAL5, DAL7, DAL81, DAN1, DAN4, DAT1, DBP6, DBP8, DCG1, DCP1, DDI2, DFP4, DIA1, DIF1, DIG2, DIM1, DIP5, DJP1, DLD3, DMC1, DNF1, DOA1, DOC1, DPH2, DSE4, DSF1, DSN1, DTD1, DUG1, DUR3, DYN3, EAP1, ECM1, ECM25, ECM29, ECM30, ECM32, ECM34, ECM7, ECO1, EDC1, EFG1, EFM1, EGH1, EGO2, EGO4, EGT2, ELO1, ELP6, EMA35, EMC1, EMC3, EMC4, EMI1, EMI2, EMP47, ENB1, ENO1, ENO2, ENT4, ENV7, ERG13, ERG20, ERG24, ERJ5, ERO1, ERR2, ERR3, ERS1, ERV29, ERV46, ESF2, ESL1, ESL2, ETP1, EUG1, FAB1, FAS2, FAT3, FAU1, FDC1, FDH1, FET4, FET5, FEX1, FEX2, FIG2, FIG4, FIT1, FIT2, FIT3, FKH1, FKS3, FLC2, FLO1, FLO10, FLO11, FLO5, FLO9, FLX1, FMO1, FMP10, FMP27, FMP32, FMP33, FMP45, FOL2, FPK1, FPR2, FPR4, FPS1, FRA1, FRD1, FRE2, FRE3, FRE4, FRE5, FRE6, FRE7, FRM2, FTR1, FUB1, FUM1, FUN12, FUN19, FUS1, FYV5, FZF1, GAB1, GAL4, GAS1, GAS4, GAT4, GCG1, GCN5, GCS1, GCV3, GDA1, GDH1, GDH3, GEM1, GEX1, GEX2, GFD2, GID7, GIM5, GIN4, GIT1, GLC8, GLK1, GLY1, GMC1, GMC2, GND1, GND2, GNP1, GON7, GOS1, GPB1, GPB2, GPH1, GPI13, GRC3, GRE2, GRH1, GRX1, GRX2, GRX4, GSR1, GTA1, GTR1, GTT1, GTT2, GUD1, GUS1, GUT1, GYP5, GYP7, HAC1, HAL5, HAP2, HAP5, HAT2, HBN1, HBT1, HCH1, HFI1, HFM1, HHY1, HIS2, HIS4, HLR1, HMG2, HMLALPHA2, HMRA1, HMRA2, HMS2, HO, HOL1, HPA2, HPA3, HRP1, HRT1, HSP104, HSP150, HSP31, HSP32, HSP82, HSU1, HUA1, HUB1, HUT1, HXK1, HXK2, HXT11, HXT13, HXT15, HXT16, HXT17, HXT8, HXT9, HYM1, HYP2, HYR1, ICR1, IES6, IGD1, IKI1, IMA1, IMA2, IMA3, IMA4, IMA5, IMD2, IMD3, IMG2, INA22, IQG1, IRC19, IRC24, IRC4, IRC5, IRC6, IRC7, ISA1, ISC10, IST3, ITR1, IZH1, JEN1, JIP4, JIP5, JJJ2, JLP1, KAP114, KAR4, KAR9, KCC4, KEG1, KEL3, KGD4, KIN82, KIP3, KOG1, KRE2, KRE28, KRE6, KRE9, KRR1, LAA1, LAG1, LAM5, LCB1, LCD1, LDB18, LEU2, LEU3, LIP1, LOS1, LOT5, LPP1, LRE1, LRG1, LSB3, LSB5, LSC2, LSM3, LSM5, LYS1, LYS9, MAK10, MAL12, MAL13, MAL31, MAL32, MAN2, MATALPHA1, 46166, MBB1, MCH2, MCH4, MCM10, MCO14, MDH2, MDJ2, MDL2, MDM1, MDM34, MED7, MES1, MET10, MET16, MET28, MFG1, MGA1, MGA2, MGM101, MGR1, MHT1, MIA40, MIC10, MID1, MIG2, MIL1, MIN10, MLC2, MLP1, MLP2, MMP1, MMS1, MND2, MNN11, MNN4, MNN5, MNT2, MNT4, MOB2, MON2, MPC3, MPH3, MPP6, MRC1, MRP2, MRP4, MRPL10, MRPL20, MRPL4, MRPL50, MRPS12, MRS1, MRS6, MRX15, MRX20, MRX6, MSB3, MSC1, MSH3, MSL1, MSO1, MST1, MTC3, MTC7, MTG2, MTM1, MTO1, MTR2, MTW1, MUP3, MVD1, MXR2, MYG1, MZM1, NAB6, NBP1, NCA2, NCE101, NCE102, NCS6, NDD1, NDI1, NFS1, NFT1, NGK1, NGL3, NIF3, NIP1, NMD3, NOG2, NOP19, NOP8, NPR2, NPR3, NRE1, NSL1, NUC1, NUD1, NUP188, NUT2, OAF1, OCA4, OCA5, OM45, OMA1, OPI1, OPT1, OPT2, ORC4, OST4, OST5, OSW7, OTU1, OTU2, OXA1, OXP1, OYE2, PAB1, PAC11, PAD1, PAN1, PAN6, PAT1, PAU1, PAU10, PAU11, PAU12, PAU13, PAU14, PAU15, PAU17, PAU18, PAU19, PAU2, PAU20, PAU21, PAU24, PAU3, PAU4, PAU6, PAU9, PBI1, PBN1, PCC1, PCK1, PCL1, PCM1, PDA1, PDE1, PDE2, PDI1, PDP3, PDR18, PEA2, PES4, PET122, PET494, PEX1, PEX11, PEX2, PEX21, PEX22, PEX29, PEX34, PEX6, PFA3, PFD1, PFK1, PFK27, PFS1, PGA3, PGU1, PHO11, PHO12, PHO13, PHO4, PHO8, PHO84, PHO90, PHR1, PIN3, PIP2, PIR5, PKH1, PLC1, PLM2, PML39, POF1, POL5, POM33, POP2, POP3, POP5, PPG1, PPM2, PRB1, PRC1, PRD1, PRE10, PRE4, PRE5, PRE8, PRI1, PRM1, PRP16, PRP19, PRP21, PRP45, PRS3, PRT1, PSE1, PSF3, PSP1, PTA1, PTC6, PTK1, PTP1, PTR2, PTR3, PUF6, PUG1, PUL3, PUL4, PUP2, PUS4, PWR1, PXA2, PXL1, PXP3, PXR1, PZF1, QCR2, QCR6, QCR7, QCR8, RAD10, RAD17, RAD2, RAD23, RAD24, RAD3, RAD4, RAI1, RBA50, RBD2, RBG1, RBH1, RCF2, RCY1, RDR1, RDS1, RDT1, RET2, REV7, RFA3, RFC3, RGD2, RGD3, RHO1, RIB3, RIB4, RIE1, RIF2, RIM101, RIM15, RIM21, RIM4, RIX7, RMD6, RMD8, RME3, RML2, RMR1, RNH70, RNP1, RNQ1, ROF1, ROG3, RPC82, RPD3, RPH1, RPL12A, RPL16A, RPL17B, RPL18A, RPL22B, RPL25, RPL29, RPL2A, RPL36B, RPL37B, RPL39, RPL40A, RPL40B, RPL6B, RPL8A, RPL8B, RPN12, RPO26, RPO41, RPR2, RPS12, RPS14B, RPS19A, RPS1A, RPS20, RPS22A, RPS4A, RRP40, RRP7, RRT5, RRT7, RSA4, RSC8, RSC9, RSM19, RSM28, |

| | |
|---|---|
| | RTC1, RTF1, RTG2, RTT102, RUF21, RUF22, RUF23, RVB2, SAC1, SAM2, SAM3, SAM4, SAP155, SAP4, SAY1, SBP1, SCC4, SCP1, SCW10, SCW4, SDA1, SDD1, SDH2, SDH7, SDS22, SDT1, SEC11, SEC12, SEC15, SEC20, SEC21, SEC39, SEC53, SEC65, SET2, SGV1, SHE10, SHS1, SHU1, SIR1, SIR3, SIT1, SKG1, SKI3, SKI8, SLD5, SLF1, SLH1, SLN1, SLO1, SMC2, SMF1, SMT3, SNA2, SNF1, SNF12, SNF6, SNM1, SNO2, SNO4, SNR191, SNR40, SNR49, SNR53, SNR64, SNR67, SNR68, SNR80, SNR84, SNR85, SNX3, SNZ2, SOL1, SOL2, SOL4, SOM1, SOP4, SOR1, SPB1, SPC72, SPC97, SPE1, SPG3, SPI1, SPO11, SPS1, SPS2, SPS22, SPT15, SPT2, SPT20, SQT1, SRB8, SRL3, SRO9, SRP40, SRP68, SRY1, SSA2, SSB1, SSK2, SSK22, SSL2, SSO1, SSP1, SST2, STB5, STE20, STE50, STE6, STL1, STS1, SUE1, SUF6, SUI3, SUP53, SUP6, SUT390, SUT532, SVP26, SWE1, SWI3, SWP82, TAD1, TAF1, TAF13, TAF8, TAN1, TCA17, TDA6, TGL3, TGL4, THI11, THI13, THI21, THI5, THP2, TIF3, TIM22, TIM23, TMA108, TMT1, TNA1, TOG1, TOH1, TOR2, TOS6, TPK1, TPM2, TPO1, TPO3, TRF5, TRM11, TRM112, TRM13, TRP3, TRT2, TRX3, TSL1, TSR2, TUB1, TUB2, TUB3, TUP1, TVP38, UBA1, UBI4, UBP11, UBP12, UBP15, UBP3, UBP5, UFO1, ULI1, UPA1, UPA2, URA1, URA5, URC2, URN1, UTP14, UTR2, UTR4, UTR5, VAC17, VAM7, VAN1, VBA3, VBA5, VEL1, VHS2, VID30, VIK1, VLD1, VMA11, VMA6, VMA8, VMR1, VPS13, VPS3, VPS52, VPS60, VPS68, VPS72, VPS9, VRG4, VTH1, VTH2, VTS1, WHI4, WSC2, WSC4, YAH1, YAP1802, YAP3, YAP5, YAR1, YBT1, YCT1, YEF1, YGK3, YKT6, YLF2, YME2, YOR1, YPD1, YPS6, YPT1, YPT32, YRA2, YRF1-2, YRF1-3, YRF1-4, YRF1-5, YRF1-6, YRF1-7, YRF1-8, YTA7, YVH1, ZDS2, ZIP2, ZNF1, ZNG1, ZOD1, ZPS1, ZRG17, ZRT1, ZUO1 |
| **gal4** | AAD10, AAD14, AAD4, ACM1, ACO2, ADD66, ADE57, ADH4, ADH6, ADK2, ADY3, AFG1, AGA1, AGE1, AGP3, AIF1, AIM17, AIM18, AIM46, AIM6, ALD4, ALR1, AMF1, ANK1, AOS1, APA2, API2, APL6, APM1, APM2, AQY1, AQY2, AQY3, ARC35, ARG8, ARG81, ARN1, ARN2, ATF1, ATG10, ATG13, ATG32, ATG41, ATP11, ATP15, ATR1, AVT2, AXL2, AYT1, BAS1, BAT1, BAT2, BBP1, BCK2, BEM2, BET2, BGL2, BIO2, BIO3, BIO4, BIO5, BMH1, BNA6, BRE4, BRE5, BRR2, BRR6, BSC5, BSC6, BSP1, BUD32, BUL2, BUR6, BXI1, CAB1, CAB4, CAC2, CAF40, CAN1, CAR2, CBP1, CBP2, CBT1, CCA1, CCT2, CDC26, CDC33, CHD1, CIN8, CLA4, CLN2, CNN1, COF1, COG3, COQ2, COQ5, COQ6, COS1, COS10, COS12, COS4, COS5, COS7, COS8, COX14, CRG1, CSA1, CSE1, CSS1, CSS3, CTF8, CTK3, CTR2, CTR9, CUB1, CUE4, CUS2, CWC22, DAK2, DAL1, DAL2, DAL3, DAL4, DAL5, DAL7, DAL81, DAL82, DAN1, DAN4, DAT1, DBP6, DCG1, DCP1, DDI2, DFP4, DIA1, DIF1, DIM1, DIP5, DLD3, DMC1, DNF1, DOA1, DOC1, DPB2, DPM1, DSE4, DSF1, DUG1, DUR3, ECM25, ECM29, ECM32, ECM34, ECM7, EFG1, EFM1, EGD2, EGT2, EMC3, EMI1, EMI2, EMW1, ENB1, ENO1, ENO2, ERG13, ERG9, ERJ5, ERO1, ERR2, ERR3, ERV29, ESF2, ESL1, EUG1, FAU1, FDC1, FDH1, FET4, FEX1, FEX2, FIG4, FIT1, FIT2, FIT3, FLO1, FLO10, FLO11, FLO9, FLX1, FMO1, FMP10, FMP27, FMP45, FOL2, FPK1, FPR2, FPR4, FPS1, FRD1, FRE3, FRE4, FRE5, FRE6, FRE7, FUM1, FZF1, GAB1, GAL4, GAS4, GAT4, GCG1, GCN5, GCS1, GDB1, GDH1, GEX2, GIN4, GLY1, GMC1, GMC2, GND1, GND2, GNP1, GOS1, GPI16, GRE2, GRH1, GRX2, GRX4, GTR1, GTT1, GTT2, GUD1, GUS1, GUT1, GYP5, GYP7, HAP2, HAT2, HBT1, HDA3, HFI1, HFM1, HHY1, HLR1, HMG2, HMS2, HO, HOL1, HPA2, HPA3, HRT1, HSP31, HSP32, HSU1, HUA1, HUT1, HXK1, HXK2, HXT11, HXT13, HXT14, HXT15, HXT16, HXT17, HXT8, HXT9, HYR1, ICR1, IES6, IKI1, IMA1, IMA2, IMA3, IMA4, IMA5, INA22, IRC24, IRC4, IRC5, IRC6, IRC7, ISC10, ITR1, JEN1, JIP5, JLP1, KAP114, KAR9, KEG1, KEL3, KGD4, KOG1, KRE1, KRE28, KRI1, LAA1, LCD1, LDB18, LEM3, LEU3, LNP1, LPP1, LRG1, LSM3, LYS1, LYS9, MAK10, MAL12, MAL13, MAN2, MBB1, MCM10, MCO14, MDH2, MDJ2, MDL2, MDM1, MDM31, MED7, MES1, MET16, MET28, MFG1, MGA2, MGM101, MHT1, MID1, MIN10, MLC2, MLP2, MMP1, MMS1, MND2, MNL1, MNT2, MNT4, MON2, MPH3, MRP2, MRPL4, MRPS12, MRPS18, MRS1, MRX15, MRX20, MRX6, MSB3, MSC1, MSO1, MTM1, MTO1, MUP3, MVD1, MYG1, NAB6, NBL1, NBP1, NCE101, NDD1, NDI1, NFT1, NGK1, NGL3, NOG2, NOP19, NOP8, NPR2, NPR3, NRE1, NUC1, NUD1, NUP188, NUT2, NVJ1, OCA5, OM45, OPI1, OPT1, OPT2, OST4, OSW7, OXA1, OXP1, OYE2, PAB1, PAD1, PAN6, PAU1, PAU11, PAU12, PAU13, PAU14, PAU15, PAU18, PAU19, PAU2, PAU20, PAU21, PAU4, PAU6, PBI1, PCK1, PCL1, PCM1, PDA1, PDE1, PDP3, PDR18, PET122, PET494, PEX11, PEX2, PEX6, PFA3, PFK27, PFS1, PFS2, PGA3, PGU1, PHA2, PHO11, PHO12, PHO13, PHO4, PHO84, PHO90, PHR1, PLC1, PLM2, PML39, POL5, POP2, PPM2, PPX1, PRB1, PRE4, PRE5, PRP21, PRP4, PSF3, PSP1, PTH1, PTP1, PUG1, PUL3, PUL4, PUP2, PUS4, PWR1, PXP3, PXR1, PZF1, QCR2, QCR6, QCR7, RAD2, RAD24, RAD3, RAD4, RAI1, RBA50, RBD2, RCY1, RDR1, RET2, REV7, RFA2, RFC3, RGD3, RHO1, RIB4, RIF2, RIM101, RIM21, RIM4, RIX1, RMD6, RMD8, RME3, RML2, RMR1, RNH70, RNP1, ROF1, RPC82, RPD3, RPH1, RPL12A, RPL18B, RPL25, RPL29, RPL2A, RPL36B, RPL37B, RPL40A, RPL6B, RPL8A, RPL8B, RPN10, RPN12, RPO26, RPR2, RPS19B, RPS1A, RPS20, RPS4A, RPS4B, RRP40, RRT5, RSC8, RSC9, RSM19, RTC1, RTF1, RTG2, RTT102, RUF23, SAC1, SAM2, SAM3, SAM4, SAP155, SAY1, SBP1, SCH9, SCW4, SDD1, SDH2, SDH7, SEC11, SEC20, SEC21, SEC23, SEC39, SEC65, SET5, SHS1, SIR1, SIR3, SIT1, SKG1, SKI3, |

| | |
|---|---|
| | SKN7, SKP2, SLF1, SLH1, SLN1, SLO1, SMN1, SMT3, SMX3, SNA2, SNF6, SNO2, SNO4, SNR191, SNR40, SNR49, SNR53, SNR64, SNR67, SNR68, SNR80, SNR84, SNR85, SNZ2, SOM1, SOR1, SPG3, SPO11, SPS1, SPS2, SPT2, SPT20, SQT1, SRY1, SSB1, SSL2, SSP1, SST2, STB1, STB5, STE6, STL1, SUF6, SUT390, SUT532, SVP26, TAD1, TAF1, TAF13, TAF8, TCA17, TDA8, THI11, THI13, THI21, THI5, TMA108, TMT1, TNA1, TOG1, TOS6, TPM2, TRF5, TRM112, TRP3, TRT2, TSL1, TUB3, UBA1, UBP11, UPA1, URA1, URA5, URC2, UTP9, VAN1, VBA5, VEL1, VHS2, VIK1, VLD1, VMA6, VMA8, VMR1, VNX1, VPS4, VPS68, VTH1, VTH2, WHI4, WSC4, YAH1, YAP5, YBT1, YCT1, YGK3, YOR1, YPD1, YPS6, YPT11, YRA2, YRF1-2, YRF1-3, YRF1-4, YRF1-5, YRF1-7, YRF1-8, YTA7, YVH1, ZDS2, ZIM17, ZIP2, ZPS1, ZRG17, ZRT1, ZUO1 |
| **gal5** | AAD10, AAD15, AAD4, ABP1, ABZ1, ACM1, ACO2, ACP1, ACS1, ACT1, ADD66, ADE4, ADE57, ADF1, ADH2, ADH4, ADH6, ADH7, ADK2, ADY3, AFG1, AGA1, AGE1, AGP1, AGP3, AHC2, AIF1, AIM17, AIM2, AIM21, AIM33, AIM6, ALD4, ALO1, ALR1, ALR2, AMF1, ANK1, ANP1, APA1, APA2, APE3, API2, APL6, APM1, APM2, APM3, AQY2, AQY3, ARC35, ARG8, ARG81, ARN1, ARN2, ASH1, ATF1, ATG10, ATG15, ATG22, ATG27, ATG32, ATG36, ATG41, ATG7, ATM1, ATP12, ATP15, ATR1, AVT2, AXL2, AYT1, BAS1, BAT2, BBP1, BCK2, BDH1, BDH2, BEM2, BGL2, BIK1, BIO2, BIO3, BIO4, BIO5, BMH1, BNA6, BOL1, BOL2, BOL3, BRE4, BRE5, BRF1, BRR2, BRR6, BSC5, BSC6, BSD2, BSP1, BUD32, BUL2, BUR6, CAB1, CAB4, CAC2, CAF120, CAF40, CAN1, CAR2, CBP1, CBP2, CBT1, CCA1, CCT2, CDC13, CDC14, CDC19, CDC24, CDC26, CDC33, CDC39, CDC50, CDC6, CHA1, CHD1, CIN2, CIN8, CLA4, CLN2, CLN3, CNA1, CNB1, CNE1, CNN1, COF1, COG1, COG3, COQ2, COQ5, COQ6, COS1, COS10, COS12, COS4, COS5, COS6, COS7, COS8, COS9, COX14, CPD1, CPR4, CPS1, CSE1, CSM1, CSM2, CSS3, CTK3, CTP1, CTR2, CTR9, CUB1, CUE4, CUR1, CUS2, CWC2, CWC21, CWC22, CYC3, CYC7, DAK2, DAL1, DAL2, DAL3, DAL4, DAL5, DAL7, DAL81, DAN1, DAN4, DAT1, DBP6, DBP8, DCG1, DCP1, DDI2, DFP4, DIA1, DIF1, DIG2, DIM1, DIP5, DJP1, DLD3, DMC1, DNF1, DOA1, DOC1, DPH2, DSE4, DSF1, DSN1, DTD1, DUG1, DUR3, DYN3, EAP1, ECM1, ECM25, ECM29, ECM30, ECM32, ECM34, ECM7, ECO1, EDC1, EFG1, EFM1, EGH1, EGO2, EGO4, ELO1, ELP6, EMA35, EMC1, EMC3, EMC4, EMI1, EMI2, EMP47, ENB1, ENO1, ENO2, ENT4, ENV7, ERG13, ERG24, ERJ5, ERO1, ERR2, ERR3, ERS1, ERV29, ERV46, ESF2, ESL1, ESL2, ETP1, EUG1, FAS2, FAT3, FAU1, FDC1, FDH1, FET4, FET5, FEX1, FEX2, FIG2, FIT1, FIT2, FIT3, FKS3, FLC2, FLO10, FLO11, FLO5, FLX1, FMO1, FMP10, FMP27, FMP32, FMP45, FOL2, FPK1, FPR2, FPR4, FPS1, FRA1, FRD1, FRE2, FRE3, FRE4, FRE5, FRE6, FRE7, FRM2, FTR1, FUB1, FUM1, FUN12, FUN19, FUS1, FYV5, FZF1, GAB1, GAL4, GAS1, GAS4, GAT4, GCG1, GCN5, GCS1, GCV3, GDA1, GDH1, GDH2, GDH3, GEM1, GEX1, GEX2, GFD2, GID7, GIM5, GIN4, GIT1, GLC8, GLK1, GLY1, GMC1, GMC2, GND1, GND2, GNP1, GON7, GOS1, GPB1, GPB2, GPH1, GPI13, GRC3, GRE2, GRH1, GRX1, GRX2, GRX4, GSR1, GTA1, GTR1, GTT1, GTT2, GUD1, GUS1, GUT1, GYP5, GYP7, HAP2, HAT2, HBN1, HBT1, HCH1, HFI1, HFM1, HHY1, HIR3, HIS2, HIS4, HLR1, HMG2, HMLALPHA2, HMRA1, HMRA2, HMS2, HO, HOL1, HOM6, HPA3, HRP1, HRT1, HSP31, HSP32, HSP82, HSU1, HUA1, HUB1, HUT1, HXK1, HXK2, HXT11, HXT13, HXT15, HXT16, HXT17, HXT8, HYM1, HYP2, HYR1, ICR1, IES6, IKI1, IMA1, IMA2, IMA3, IMA5, IMD2, IMG2, INA22, IPA1, IQG1, IRC19, IRC24, IRC4, IRC5, IRC6, IRC7, ISA1, ISC10, IST3, ITR1, IZH1, JEN1, JIP4, JIP5, JLP1, KAP114, KAR4, KAR9, KEG1, KEL3, KGD4, KIN82, KIP3, KOG1, KRE2, KRE28, KRE6, KRE9, KRR1, LAA1, LAG1, LAM5, LCB1, LCD1, LDB18, LEU3, LIP1, LOS1, LOT5, LPP1, LRE1, LRG1, LSB3, LSB5, LSC2, LSM3, LSM5, LYS1, LYS9, MAK10, MAL12, MAL13, MAL31, MAL32, MAL33, MAN2, MATALPHA1, 46166, MBB1, MCH2, MCH4, MCM10, MCO14, MDH2, MDL2, MDM1, MDM34, MDY2, MED7, MES1, MET10, MET16, MET28, MFG1, MGA1, MGA2, MGM101, MGR1, MHT1, MIA40, MIC10, MID1, MIG2, MIN10, MLP1, MLP2, MMP1, MMS1, MND2, MNN11, MNN4, MNN5, MNT2, MNT4, MON2, MPC3, MPH1, MPH3, MRC1, MRP2, MRP4, MRPL10, MRPL20, MRPL4, MRPS12, MRS1, MRS6, MRX15, MRX20, MRX6, MSB3, MSB4, MSC1, MSH3, MSL1, MSN1, MSO1, MST1, MTC3, MTC7, MTG2, MTM1, MTO1, MTR2, MTW1, MUP3, MVD1, MXR2, MYG1, MZM1, NAB6, NBP1, NCA2, NCE101, NCE102, NCS6, NDD1, NDI1, NFT1, NGK1, NGL3, NIF3, NIP1, NMD3, NOG2, NOP19, NOP6, NOP8, NPR2, NPR3, NRE1, NSL1, NUC1, NUD1, NUP188, NUT2, OAF1, OCA4, OCA5, OM45, OMA1, OPI1, OPT1, ORC4, OST4, OST5, OSW7, OTU1, OTU2, OXA1, OXP1, OYE2, PAB1, PAC11, PAD1, PAN1, PAN6, PAP2, PAT1, PAU1, PAU10, PAU11, PAU12, PAU13, PAU14, PAU15, PAU18, PAU19, PAU2, PAU20, PAU21, PAU24, PAU3, PAU4, PAU6, PAU9, PBI1, PBN1, PCA1, PCC1, PCK1, PCL1, PCM1, PDA1, PDE1, PDI1, PDP3, PDR18, PEA2, PET122, PET494, PEX1, PEX11, PEX2, PEX21, PEX22, PEX29, PEX34, PFD1, PFK1, PFK27, PFS1, PGA3, PGU1, PHO12, PHO13, PHO4, PHO8, PHO84, PHO89, PHO90, PHR1, PIN3, PIP2, PKH1, PLC1, PLM2, PML39, PMT4, POF1, POL5, POP2, POP3, POP5, PPG1, PPM2, PRB1, PRC1, PRD1, PRE4, PRE5, PRE8, PRI1, PRM1, PRP16, PRP19, PRP21, PRP45, PRR2, PRS1, PRS3, PSE1, PSF3, PSP1, PTA1, PTC6, PTH4, PTK1, PTP1, PTR2, PTR3, PUF6, PUG1, PUL3, PUL4, PUP2, PUS4, |

| | |
|---|---|
| | PWR1, PXA2, PXL1, PXP3, PXR1, QCR6, QCR7, RAD10, RAD17, RAD2, RAD23, RAD24, RAD3, RAD4, RAI1, RBA50, RBD2, RBG1, RBH1, RCY1, RDR1, RDS1, RDT1, RET2, REV7, RFA3, RFC3, RGD2, RGD3, RHO1, RIB3, RIB4, RIE1, RIF2, RIM101, RIM21, RIM4, RIX7, RMD6, RMD8, RME3, RML2, RMR1, RNH70, RNP1, RNQ1, ROF1, RPH1, RPL12A, RPL16A, RPL17B, RPL18A, RPL25, RPL29, RPL2A, RPL36B, RPL37B, RPL39, RPL40A, RPL40B, RPL6B, RPL8A, RPL8B, RPN12, RPR2, RPS12, RPS14B, RPS19A, RPS1A, RPS20, RPS22A, RPS4A, RRI1, RRI2, RRP40, RRP7, RRT5, RRT7, RSA4, RSC8, RSC9, RSM19, RSM28, RTC1, RTF1, RTG2, RTT102, RUF21, RUF23, RVB2, SAC1, SAM2, SAM3, SAM4, SAP155, SAP4, SAY1, SBP1, SCC4, SCP1, SCW10, SCW4, SDA1, SDD1, SDH2, SDH7, SDS22, SDT1, SEC11, SEC15, SEC20, SEC21, SEC39, SEC53, SEC65, SGV1, SHE10, SHR3, SHR5, SHS1, SHU1, SIR1, SIR3, SIT1, SKG1, SKI8, SKM1, SLD5, SLF1, SLH1, SLN1, SLO1, SMC2, SMF1, SMT3, SNA2, SNF1, SNF5, SNF6, SNM1, SNO2, SNO4, SNR191, SNR40, SNR49, SNR53, SNR64, SNR67, SNR68, SNR80, SNR84, SNR85, SNZ2, SOL1, SOL2, SOL4, SOM1, SOP4, SOR1, SPB1, SPC72, SPC97, SPE1, SPG3, SPI1, SPO11, SPS1, SPS2, SPS22, SPT15, SPT2, SPT20, SQT1, SRB8, SRL3, SRO9, SRP40, SRP68, SRY1, SSB1, SSK22, SSL2, SSO1, SSP1, SST2, STB5, STE20, STE50, STE6, STL1, STS1, SUE1, SUF6, SUI3, SUL1, SUP6, SUT390, SUT532, SVP26, SWE1, SWI3, SWP82, TAD1, TAF1, TAF13, TAF8, TAN1, TCA17, TDA6, TGL3, TGL4, THI11, THI13, THI21, TIF3, TIM22, TMA108, TMT1, TNA1, TOG1, TOH1, TOR2, TPM2, TPO1, TPO3, TRF5, TRM11, TRM112, TRM13, TRP3, TRT2, TRX3, TSL1, TSR2, TUB1, TUB2, TUB3, TUP1, TVP38, TYC1, UBA1, UBI4, UBP11, UBP12, UBP15, UBP3, UFO1, UGA4, ULI1, UPA1, UPA2, URA1, URA5, URC2, URN1, UTP14, UTR2, UTR4, UTR5, VAC17, VAM7, VAN1, VBA2, VBA3, VBA5, VEL1, VHS2, VID30, VIK1, VLD1, VMA11, VMA6, VMA8, VMR1, VPS13, VPS3, VPS52, VPS60, VPS68, VPS72, VPS9, VRG4, VTH1, VTH2, WHI4, WSC2, WSC4, YAH1, YAP1802, YAP3, YAP5, YAR1, YBT1, YCT1, YEF1, YGK3, YKT6, YLF2, YME2, YOR1, YPD1, YPS6, YPT1, YPT32, YRA2, YRF1-2, YRF1-4, YRF1-5, YRF1-6, YTA7, YVH1, ZDS2, ZIP2, ZNF1, ZOD1, ZPS1, ZRG17, ZRT1, ZUO1 |
| **gal6** | AAD10, AAD14, AAD4, ABZ1, ACM1, ACO2, ACP1, ACS1, ACT1, ADD66, ADE4, ADE57, ADH2, ADH4, ADH6, ADK2, ADY3, AFG1, AGA1, AGE1, AGP3, AIF1, AIM17, AIM18, AIM2, AIM21, AIM33, AIM46, AIM6, ALD4, ALO1, ALR1, ALR2, AMF1, ANK1, ANP1, AOS1, APA2, API2, APL6, APM1, APM2, AQY1, AQY2, AQY3, ARC35, ARG8, ARG81, ARN1, ARN2, ASG7, ATF1, ATG10, ATG13, ATG22, ATG23, ATG27, ATG32, ATG36, ATG41, ATG7, ATM1, ATP11, ATP12, ATP15, ATP18, ATR1, AVT2, AXL2, AYT1, BAS1, BAT1, BAT2, BBP1, BCK2, BDH1, BDH2, BEM2, BET2, BGL2, BIK1, BIO2, BIO3, BIO4, BIO5, BMH1, BNA6, BOL1, BOL2, BOL3, BRE4, BRE5, BRR2, BRR6, BSC5, BSC6, BSP1, BUD32, BUL2, BUR6, BXI1, CAB1, CAB4, CAC2, CAF40, CAN1, CAR2, CBP1, CBP2, CBT1, CCA1, CCT2, CDC13, CDC14, CDC19, CDC24, CDC26, CDC33, CDC6, CHD1, CIN8, CIS3, CLA4, CLN2, CLN3, CNA1, CNB1, CNE1, CNN1, COF1, COG1, COG3, COQ2, COQ5, COQ6, COS1, COS10, COS12, COS4, COS5, COS6, COS7, COS8, COS9, COX14, CPS1, CRG1, CSA1, CSE1, CSM2, CSS1, CSS3, CTF8, CTK3, CTR2, CTR9, CUB1, CUE4, CUR1, CUS2, CWC21, CWC22, CYC3, CYC7, DAK2, DAL1, DAL2, DAL3, DAL4, DAL5, DAL7, DAL81, DAL82, DAN1, DAN4, DAT1, DBP6, DBP8, DCG1, DCP1, DDI2, DFP4, DIA1, DIF1, DIG2, DIM1, DIP5, DJP1, DLD3, DMC1, DNF1, DOA1, DOC1, DPB2, DPH2, DPM1, DSE4, DSF1, DSN1, DUG1, DUR3, DYN3, EAP1, ECM1, ECM25, ECM29, ECM30, ECM32, ECM34, ECM7, ECO1, EDC1, EFG1, EFM1, EGD2, EGH1, EGO4, EGT2, ELO1, ELP6, EMA35, EMC3, EMC4, EMI1, EMI2, EMP47, EMW1, ENB1, ENO1, ENO2, ENT4, ERG13, ERG20, ERG9, ERJ5, ERO1, ERR2, ERR3, ERV29, ERV46, ESF2, ESL1, ESL2, ETP1, EUG1, FAU1, FDC1, FDH1, FET4, FET5, FEX1, FEX2, FIG4, FIT1, FIT2, FIT3, FKH1, FKS3, FLC2, FLO1, FLO10, FLO11, FLO9, FLX1, FMO1, FMP10, FMP27, FMP32, FMP33, FMP45, FOL2, FPK1, FPR2, FPR4, FPS1, FRD1, FRE2, FRE3, FRE4, FRE5, FRE6, FRE7, FRM2, FUM1, FUN12, FUN19, FUS1, FZF1, GAB1, GAL4, GAS1, GAS4, GAT4, GCG1, GCN5, GCS1, GCV3, GDA1, GDB1, GDH1, GDH3, GEM1, GEX2, GFD2, GID7, GIM5, GIN4, GIT1, GLC8, GLK1, GLY1, GMC1, GMC2, GND1, GND2, GNP1, GON7, GOS1, GPB1, GPB2, GPH1, GPI16, GRC3, GRE2, GRH1, GRX1, GRX2, GRX4, GSR1, GTA1, GTR1, GTT1, GTT2, GUD1, GUS1, GUT1, GYP5, GYP7, HAL5, HAP2, HAT2, HBN1, HBT1, HDA3, HFI1, HFM1, HHY1, HIS4, HLR1, HMG2, HMS2, HO, HOL1, HPA2, HPA3, HRT1, HSP150, HSP31, HSP32, HSU1, HUA1, HUB1, HUT1, HXK1, HXK2, HXT11, HXT13, HXT14, HXT15, HXT16, HXT17, HXT8, HXT9, HYP2, HYR1, ICR1, IES6, IKI1, IMA1, IMA2, IMA3, IMA4, IMA5, IMD3, INA22, IRC24, IRC4, IRC5, IRC6, IRC7, ISC10, IST3, ITR1, IZH1, JEN1, JIP4, JIP5, JJJ2, JLP1, KAP114, KAR9, KEG1, KEL3, KGD4, KIP3, KOG1, KRE1, KRE2, KRE28, KRE6, KRE9, KRI1, LAA1, LAM5, LCB1, LCD1, LDB18, LEM3, LEU3, LIP1, LNP1, LOS1, LPP1, LRG1, LSB5, LSM3, LYS1, LYS9, MAK10, MAL12, MAL13, MAL31, MAL32, MAN2, 46166, MBB1, MCH2, MCM10, MCO14, MDH2, MDJ2, MDL2, MDM1, MDM31, MDM34, MED7, MES1, MET10, MET16, MET28, MFG1, MGA1, MGA2, MGM101, MGR1, MHT1, MIA40, MID1, MIG2, MIN10, MLC2, MLP1, MLP2, MMP1, MMS1, MND2, MNL1, MNN11, MNN4, MNN5, MNT2, MNT4, MON2, MPH3, MRP2, MRPL20, MRPL4, MRPS12, MRPS18, MRS1, MRS6, MRX15, MRX20, MRX6, |

| | |
|---|---|
| | MSB3, MSC1, MSL1, MSO1, MST1, MTC3, MTC7, MTG2, MTM1, MTO1, MTW1, MUP3, MVD1, MXR2, MYG1, MZM1, NAB6, NBL1, NBP1, NCA2, NCE101, NCE102, NCS6, NDD1, NDI1, NFT1, NGK1, NGL3, NIF3, NIP1, NMD3, NOG2, NOP19, NOP8, NPR2, NPR3, NRE1, NUC1, NUD1, NUP188, NUT2, NVJ1, OAF1, OCA5, OM45, OMA1, OPI1, OPT1, OPT2, ORC4, OST4, OST5, OSW7, OTU1, OTU2, OXA1, OXP1, OYE2, PAB1, PAC11, PAD1, PAN1, PAN6, PAU1, PAU10, PAU11, PAU12, PAU13, PAU14, PAU15, PAU18, PAU19, PAU2, PAU20, PAU21, PAU24, PAU4, PAU6, PBI1, PCC1, PCK1, PCL1, PCM1, PDA1, PDE1, PDI1, PDP3, PDR18, PET122, PET494, PEX1, PEX11, PEX2, PEX22, PEX29, PEX6, PFA3, PFD1, PFK27, PFS1, PFS2, PGA3, PGU1, PHA2, PHO11, PHO12, PHO13, PHO4, PHO8, PHO84, PHO90, PHR1, PIN3, PIR5, PKH1, PLC1, PLM2, PML39, PMT4, POL5, POP2, POP5, PPG1, PPM2, PPX1, PRB1, PRC1, PRE4, PRE5, PRE8, PRI1, PRP16, PRP19, PRP21, PRP4, PRP45, PRS3, PSE1, PSF3, PSP1, PTA1, PTH1, PTK1, PTP1, PTR2, PTR3, PUF6, PUG1, PUL3, PUL4, PUP2, PUS4, PWR1, PXL1, PXP3, PXR1, PZF1, QCR2, QCR6, QCR7, QCR8, RAD10, RAD2, RAD23, RAD24, RAD3, RAD4, RAI1, RBA50, RBD2, RBG1, RBH1, RCY1, RDR1, RET2, REV7, RFA2, RFA3, RFC3, RGD2, RGD3, RHO1, RIB3, RIB4, RIE1, RIF2, RIM101, RIM21, RIM4, RIX1, RMD6, RMD8, RME3, RML2, RMR1, RNH70, RNP1, RNQ1, ROF1, RPC82, RPD3, RPH1, RPL12A, RPL16A, RPL17B, RPL18B, RPL25, RPL29, RPL2A, RPL36B, RPL37B, RPL39, RPL40A, RPL40B, RPL6B, RPL8A, RPL8B, RPN10, RPN12, RPO26, RPR2, RPS12, RPS14B, RPS19B, RPS1A, RPS20, RPS22A, RPS4A, RPS4B, RRP40, RRP7, RRT5, RSC8, RSC9, RSM19, RSM28, RTC1, RTF1, RTG2, RTT102, RUF21, RUF23, SAC1, SAM2, SAM3, SAM4, SAP155, SAP4, SAY1, SBP1, SCH9, SCW10, SCW4, SDD1, SDH2, SDH7, SDS22, SDT1, SEC11, SEC15, SEC20, SEC21, SEC23, SEC39, SEC53, SEC65, SET2, SET5, SGV1, SHE10, SHS1, SHU1, SIR1, SIR3, SIT1, SKG1, SKI3, SKI8, SKN7, SKP2, SLD5, SLF1, SLH1, SLN1, SLO1, SMC2, SMN1, SMT3, SMX3, SNA2, SNF1, SNF6, SNM1, SNO2, SNO4, SNR191, SNR40, SNR49, SNR53, SNR64, SNR67, SNR68, SNR80, SNR84, SNR85, SNZ2, SOL1, SOL4, SOM1, SOP4, SOR1, SPC72, SPC97, SPG3, SPO11, SPS1, SPS2, SPT2, SPT20, SQT1, SRL3, SRO9, SRP40, SRY1, SSB1, SSL2, SSP1, SST2, STB1, STB5, STE20, STE50, STE6, STL1, STS1, SUE1, SUF6, SUP6, SUT390, SUT532, SVP26, SWE1, SWI3, SWP82, TAD1, TAF1, TAF13, TAF8, TAN1, TCA17, TDA6, TDA8, TGL3, TGL4, THI11, THI13, THI21, THI5, TIF3, TMA108, TMT1, TNA1, TOG1, TOH1, TOR2, TOS6, TPK1, TPM2, TPO3, TRF5, TRM112, TRP3, TRT2, TSL1, TSR2, TUB1, TUB2, TUB3, TVP38, UBA1, UBI4, UBP11, UBP12, UBP15, UFO1, UPA1, UPA2, URA1, URA5, URC2, URN1, UTP14, UTP9, UTR2, UTR4, UTR5, VAM7, VAN1, VBA5, VEL1, VHS2, VID30, VIK1, VLD1, VMA6, VMA8, VMR1, VNX1, VPS13, VPS3, VPS4, VPS52, VPS60, VPS68, VPS72, VPS9, VRG4, VTH1, VTH2, WHI4, WSC4, YAH1, YAP3, YAP5, YBT1, YCT1, YEF1, YGK3, YKT6, YLF2, YME2, YOR1, YPD1, YPS6, YPT1, YPT11, YPT32, YRA2, YRF1-2, YRF1-3, YRF1-4, YRF1-5, YRF1-7, YRF1-8, YTA7, YVH1, ZDS2, ZIM17, ZIP2, ZNF1, ZOD1, ZPS1, ZRG17, ZRT1, ZUO1 |

**1200 generations**

| Replicate | Gene list |
|---|---|
| **glu1** | AAC1, ADE13, AIM20, APS1, ARP7, ATG33, ATG38, ATG44, ATO3, BDF1, BER1, BLS1, BNR1, BUD8, CAR2, CCC1, CDC123, CIN10, CIS1, CLB4, COA1, COA3, COA4, COQ9, CPR6, CRR1, CSR1, CTR3, DIC1, DIF1, DLS1, DPH5, DUS3, DUS4, EFT1, ENT2, ERV2, FBP1, FKS1, FRE1, GAG1, GAS2, GLN1, HCR1, HMX1, HRD3, ICS3, IDP2, ILV5, INA1, IRC16, KAP95, LPX1, LSM3, LUG1, MAE1, MPM1, MRPL4, MRPL8, MSC3, MSS51, MUS81, NCW2, NIT1, NIT3, NMT1, ORM2, OST3, OXA1, PBA1, PET122, PEX13, PGD1, PIB2, PNP1, POT1, PRY1, PRY3, PSY3, PUN1, PWP1, QRI5, RFX1, RPL26A, RPL31B, RPP0, RPS31, RSA3, RSC2, RVS167, SAM1, SEC13, SEC61, SEI1, SFP1, SMU2, SOA1, SPO24, SPO77, STP3, STT3, SUC2, TAG1, TAL1, TFS1, TIP41, TOP3, TRR4, TUB4, UBP7, UPS1, UPS2, UTP21, VIP1, VMA13, VTA1, WHI5, YKE2, ZWF1 |

| | |
|---|---|
| **glu2** | AAC1, ACS2, APS1, ARL3, ARP7, ATG21, ATG26, ATG33, ATG38, ATO3, BAP3, BUB2, BUD8, CAM1, CAR1, CCC1, CCT6, CDC123, CDC34, CIN10, CIS1, CLB4, COA4, COI1, COQ9, CPR6, CQD1, CRR1, CSR1, CTF3, DBF4, DET1, DIC1, DIG1, DPC25, DPH5, EEB1, EFT1, EGD1, ELC1, ELP4, EMC10, EMG1, ENA1, ENT2, ERI1, ERV2, FBP1, FKS1, FMP30, FRE1, GAS2, GDE1, GLN1, GLR1, HCR1, HEM12, HEM13, HMX1, HRD3, HTS1, IDP2, ILV5, INA17, IOC4, IRC16, ISM1, KAP95, KRS1, KTR6, LCL1, LEE1, LGE1, LPX1, LUG1, MAS1, MCM1, MDL1, MET31, MGR2, MMR1, MNN9, MRX11, MSC3, MSD1, MSS51, MSY1, MUS81, NAM2, NCW2, NIT3, NKP1, NMT1, NOG1, NOP4, NRG1, NUP116, OAZ1, ORM2, OST3, PBA1, PEX13, PGD1, PIB2, PNG1, PNP1, PST1, PSY3, PUS5, PWP1, QRI5, RFX1, RLM1, RNH203, RPC11, RPL26A, RPL37A, RPP0, RPP2B, RPS31, RPS6A, RSA3, RSC2, RSM10, RVS167, SAM1, SEC13, SEC61, SEC62, SGF11, SHH4, SKG3, SND1, SNR47, SPO24, SPO77, SSE1, SSN3, SSU1, STP3, STT3, SUR1, SWI6, SYH1, TAG1, TAL1, TFS1, TIP41, TOS4, TPI1, TRR4, TUB4, UPS1, UPS2, UPS3, UTP13, VMA13, VMS1, VPS16, VTA1, WHI5, YKE2, YRA1 |
| **glu3** | ACL4, ADA2, ADE8, ADK1, ADR1, AFR1, AGE1, AHA1, AIM7, AKR1, ALT2, AMD2, APA2, APC4, API2, APT2, ARG82, ARH1, ARO1, ARO10, ARO80, ARP10, ARX1, ASP1, ATC1, ATO3, ATP17, ATP22, ATP5, BCP1, BCS1, BFR2, BMH2, BNA7, BTT1, CAB5, CAD1, CBS2, CCC2, CCT6, CDC1, CDC37, CDC40, CFT1, CHL4, CIA1, CIN10, CMI8, CNL1, COI1, COQ4, COX20, COX26, CPP2, CPR1, CPR5, CRF1, CSN9, CTA1, CTH1, CTS2, CWC15, CWC21, CYM1, DAD4, DFM1, DIG2, DIN7, DIT1, DIT2, DNF2, DOA4, DON1, DOP1, DOS2, DOT1, DPB4, DPL1, DPP1, DXO1, DYN2, EAF1, EBS1, ECM11, ECM18, EFT1, EKI1, EMI1, EMI2, EMT1, ENT5, ERD1, ESC2, ESF1, EUG1, EXG2, FCF1, FIN1, FMN1, FMP16, FOB1, FPR2, FRQ1, GCD6, GCN2, GGA1, GIC2, GIN4, GIR2, GIS1, GLO2, GMC1, GNP1, GPI11, GPI17, GPI19, GPI8, GRH1, GRX2, GRX3, GTB1, GUK1, HDA2, HEH2, HEL2, HEM1, HIM1, HKR1, HLR1, HMO1, HNT2, HOM2, HPR1, HPT1, HRQ1, HSP42, HSP78, HST4, HTA1, HTB1, HXT3, HXT7, ILT1, INM2, INO2, IPK1, IPT1, IRC3, ITR1, IVY1, IZH1, JIP4, KEI1, KGD2, KIN1, KRE2, LCB2, LCD1, LPP1, LRS4, LSM6, LYS4, MAK21, MCM21, MET32, MFA1, MFB1, MGP12, MHR1, MKC7, MNN10, MOR1, MRP1, MRP20, MRPL1, MRPL28, MRPL35, MRPL7, MRPS28, MRX10, MRX14, MRX16, MRX8, MSC2, MSH6, MSN5, MSS116, MSS4, MSW1, MTC5, MTH1, MTQ2, MUS81, MZM1, NBP2, NCB2, NGG1, NHX1, NKP1, NPL3, NSE3, NUM1, NUP42, NVJ3, OCA6, OMS1, PAA1, PAC11, PAL1, PAM1, PCF11, PDC2, PDR15, PDS1, PEP7, PET100, PEX10, PEX29, PEX3, PEX5, PEX7, PFA5, PFU1, PHM6, PHO8, PHO92, PIB1, PKH1, PKH3, PLM2, PLP1, PMP3, PMT7, PPH3, PPM1, PPN1, PPZ2, PRO1, PRP28, PRP3, PRP42, PSP1, PUF6, QCR7, RAD30, RAD34, RAD55, RAD9, RAV2, RBA50, REF2, RGA2, RGP1, RIB3, RKM2, RKM4, RLI1, RMD5, RMT2, RNH202, RPA14, RPB7, RPL12B, RPL27B, RPL37B, RPN9, RPP2B, RPS13, RPS17B, RPS18A, RPT3, RQC1, RRG1, RRP1, RRP17, RRP45, RRP8, RSC3, RSM24, RSM28, RTN1, RTR2, RTT103, RUB1, RVB1, RVS167, SAC3, SAC6, SAC7, SAM2, SAN1, SAS4, SBE2, SCC2, SDC1, SDH4, SDH6, SDH7, SEC1, SEC20, SEC26, SEC7, SED1, SEM1, SHE9, SHU2, SIP1, SIR4, SIZ1, SKP1, SLD5, SLF1, SLU7, SLY1, SMT3, SNA2, SND1, SNF1, SNF11, SNM1, SNR13, SNR84, SNU56, SNX41, SPC110, SPC19, SPG3, SPO71, SPP41, SPR28, SPS1, SPS2, SPT3, SRB7, SRP101, SSD1, SSF2, SSN2, SSS1, SSY1, STB3, STE14, STE5, STN1, STP1, SUF3, SUM1, SUP2, SUP35, SUR2, SVF1, SWA2, SWF1, SWI5, SWM1, SWR1, SXM1, SYF1, TAF10, TAF12, TCP1, TFB1, TFB3, TFB5, TFC6, THI74, TIF35, TIM11, TLD1, TLG1, TMA64, TMN2, TMS1, TOM1, TPS2, TRM1, TRM82, TRP4, TRR1, TRS120, TRS23, TRS31, TRS85, TSA2, TVP15, TVP23, UBA2, UBC1, UBC13, UBC5, UBX5, UGO1, UME6, UPC2, UPS3, URC2, URH1, UTP4, UTP5, UTP6, VBA4, VHS1, VPS3, VPS41, VPS52, VPS60, VPS64, VPS72, VPS74, VTC5, WIP1, XRS2, YAP6, YCF1, YCG1, YFT2, YHP1, YPQ2, YPR1, YPS7, YRA1, YSP2, ZIP1 |
| **glu4** | APS1, ATG38, BER1, BLS1, CCC1, CDC123, CLB4, COA4, CPR6, CRR1, CTR3, DPH5, DUS4, EMG1, FRE1, GAG1, IDP2, MSC3, PDC2, PET100, RFX1, RPL31B, RPL37A, RPS31, RSA3, SAM1, SED1, SEI1, SFP1, SHU2, SKG3, STN1, SWI6, TAG1, TFB5, TFS1, TOS4, TUB4, UPS2, UTP21, VIP1, VPS41, VTA1 |
| **glu5** | AIM20, APS1, ATG32, ATG33, ATG38, ATG44, AXL2, BNR1, BUD8, CCC1, CCT2, CDC123, CIS3, CLB4, COA1, COA4, COQ9, CPR6, CRR1, DIB1, DPH5, ENT2, ERC1, ERV2, ESL1, EST1, FLX1, FRE1, GLN1, GRS2, GUT2, HMX1, HPF1, HRD3, ICS3, IDP2, ILV5, IMP21, IRC16, LUG1, MCM10, MDM36, MHP1, MLP2, MRL1, MSC3, MSS51, NCW2, NIT1, NIT3, NMT1, OM45, OPY2, ORM2, PAN6, PBA1, PNP1, POT1, PRY1, PRY3, PWP1, QRI5, REV7, RFX1, RPL40A, RPP0, RPS31, RRD1, RSA3, RSC2, SCS3, SEC13, SLN1, SMU2, SNR68, SOA1, SPO24, SPO77, SSL2, SUC2, TAG1, TAL1, TEF1, TFS1, TIP41, TKL1, TMA108, TOP3, TPM2, TUB4, UBP7, UPS1, UPS2, VHS2, VMA13, VPR1, YKE2 |

| | |
|---|---|
| **glu6** | ACS2, AIM7, APS1, ATG38, BAP3, CCC1, CDC123, CDC34, CLB4, COA4, COQ9, CPR6, CRR1, DBF4, DET1, DOA4, DOS2, DPH5, EMC10, ENT2, FMP16, FRE1, HCR1, HEM12, HEM13, HMX1, HRD3, IDP2, IPT1, LCB2, MAK21, MAS1, MSC3, MSS51, NCW2, NMT1, NRG1, OCA6, PAA1, PBA1, PDC2, PET100, PEX13, PNP1, PPH3, PST1, PUS5, PWP1, QRI5, RAD55, RFX1, RNH203, RPC11, RPS13, RPS31, RRG1, RSA3, RSM10, RTR2, SEC13, SED1, SHH4, SHU2, SNF11, SNR47, TAG1, TFB5, TGL2, TPI1, TPS2, TUB4, UBC5, UPS1, UPS2, UTP13, VMS1, VPS41, YKE2, YOS9 |
| **gal1** | AAD10, AAD14, AAD15, AAD4, ABP1, ABZ1, ABZ2, ACM1, ACO2, ACP1, ACS1, ACT1, ADD66, ADE4, ADE57, ADF1, ADH2, ADH4, ADH6, ADH7, ADK2, ADY3, AFG1, AGA1, AGE1, AGP1, AGP3, AHC2, AIF1, AIM17, AIM2, AIM21, AIM33, AIM6, ALD4, ALG12, ALO1, ALR1, ALR2, AMF1, ANK1, ANP1, AOS1, APA1, APA2, APE3, API2, APL6, APM1, APM2, APM3, AQY1, AQY2, AQY3, ARC35, ARG8, ARG81, ARN1, ARN2, ASG7, ASH1, ATF1, ATG10, ATG13, ATG18, ATG22, ATG23, ATG27, ATG32, ATG36, ATG41, ATG7, ATM1, ATP12, ATP15, ATP18, ATR1, AVT2, AXL2, AYT1, BAS1, BAT2, BBP1, BCK2, BDH1, BDH2, BEM2, BET1, BET2, BGL2, BIK1, BIO2, BIO3, BIO4, BIO5, BMH1, BNA6, BOL1, BOL2, BOL3, BRE4, BRE5, BRR2, BRR6, BSC5, BSC6, BSD2, BSP1, BUD17, BUD32, BUL2, BUR6, CAB1, CAB4, CAC2, CAF120, CAF40, CAN1, CAR2, CBP1, CBP2, CBT1, CCA1, CCT2, CDC13, CDC14, CDC19, CDC23, CDC24, CDC26, CDC33, CDC39, CDC50, CDC6, CFD1, CFF1, CHA1, CHD1, CIN2, CIN8, CIS3, CLA4, CLN2, CLN3, CMI7, CNA1, CNB1, CNE1, CNN1, COF1, COG1, COG3, COQ2, COQ5, COQ6, COS1, COS10, COS12, COS4, COS5, COS6, COS7, COS8, COS9, COX14, CPD1, CPR8, CPS1, CRN1, CSA1, CSE1, CSM1, CSM2, CSS1, CSS2, CSS3, CTK3, CTP1, CTR2, CTR9, CUB1, CUE4, CUR1, CUS2, CWC2, CWC21, CWC22, CYC3, CYC7, DAK2, DAL1, DAL2, DAL3, DAL4, DAL5, DAL7, DAL81, DAN1, DAN4, DAT1, DBP6, DBP8, DCG1, DCP1, DDI2, DFP4, DIA1, DIF1, DIG2, DIM1, DIP5, DJP1, DLD3, DMC1, DNA2, DNF1, DOA1, DOC1, DPB2, DPB3, DPH2, DPM1, DSE4, DSF1, DSN1, DSS1, DTD1, DUG1, DUG2, DUR3, DUS1, DYN3, EAP1, ECM1, ECM25, ECM29, ECM30, ECM32, ECM34, ECM7, ECO1, EDC1, EFG1, EFM1, EGH1, EGO4, ELO1, ELP6, EMA35, EMC1, EMC3, EMC4, EMI1, EMI2, EMP47, ENB1, ENO1, ENO2, ENT4, ENV7, EPS1, ERG13, ERG20, ERG24, ERJ5, ERO1, ERR2, ERR3, ERV29, ERV46, ESF2, ESL2, ETP1, EUG1, FAB1, FAR1, FAT3, FAU1, FBP26, FDC1, FDH1, FET4, FET5, FEX1, FEX2, FIG2, FIT1, FIT2, FIT3, FKH1, FKS3, FLC2, FLO1, FLO10, FLO11, FLO5, FLO9, FLX1, FMO1, FMP10, FMP27, FMP32, FMP33, FMP45, FOL2, FPK1, FPR2, FPR4, FPS1, FRA1, FRD1, FRE2, FRE3, FRE4, FRE5, FRE6, FRE7, FRM2, FUM1, FUN12, FUN19, FUS1, FYV5, FZF1, GAB1, GAL4, GAS1, GAS4, GAT4, GCG1, GCN5, GCS1, GCV3, GDA1, GDB1, GDH1, GDH2, GDH3, GEM1, GEX1, GEX2, GFD2, GID7, GIM5, GIN4, GIT1, GLC8, GLK1, GLY1, GMC1, GMC2, GND1, GND2, GNP1, GON7, GOS1, GOT1, GPA1, GPB1, GPB2, GPH1, GPI13, GRC3, GRE2, GRH1, GRX1, GRX2, GRX4, GSR1, GTA1, GTR1, GTT1, GTT2, GUD1, GUS1, GUT1, GYP5, GYP7, HAB1, HAL5, HAP2, HAS1, HAT2, HBN1, HBT1, HCH1, HDA3, HER2, HFI1, HFM1, HHY1, HIR3, HIS2, HIS4, HLR1, HMG2, HMLALPHA2, HMRA1, HMRA2, HMS2, HO, HOL1, HOM6, HPA2, HPA3, HRP1, HRT1, HSE1, HSH155, HSP150, HSP31, HSP32, HSP82, HSU1, HUA1, HUB1, HUT1, HXK1, HXK2, HXT11, HXT13, HXT15, HXT16, HXT17, HXT8, HXT9, HYM1, HYR1, ICR1, IES6, IGD1, IMA1, IMA2, IMA3, IMA4, IMA5, IMD2, IMD3, IML1, INA22, INP51, IPA1, IQG1, IRC19, IRC24, IRC4, IRC5, IRC6, IRC7, ISC10, IST3, ITR1, IZH1, JEN1, JIP4, JIP5, JJJ2, JLP1, JNM1, KAP114, KAR4, KAR9, KEG1, KEL3, KGD4, KIN82, KIP3, KRE2, KRE28, KRE6, KRE9, KRR1, LAA1, LAG1, LAM5, LCB1, LCD1, LDB18, LEU3, LEU5, LIP1, LOS1, LOT5, LPP1, LRE1, LRG1, LSB3, LSB5, LSM3, LYS1, LYS9, MAG2, MAK10, MAL12, MAL13, MAL31, MAL32, MAL33, MAN2, MATALPHA1, 46166, MBB1, MCH2, MCM22, MCO14, MDH2, MDL2, MDM1, MDM34, MED7, MES1, MET10, MET16, MET28, MET5, MFG1, MGA1, MGA2, MGM101, MGR1, MHT1, MIA40, MIC10, MID1, MIG2, MIN10, MLC2, MLP1, MMP1, MMS1, MND2, MNN11, MNN4, MNN5, MNT2, MNT4, MON2, MPC2, MPH1, MPH3, MPP6, MRC1, MRP2, MRP4, MRPL10, MRPL27, MRPL33, MRPL4, MRPS12, MRS1, MRS6, MRX15, MRX20, MRX6, MSB3, MSC1, MSH3, MSL1, MSO1, MST1, MTC3, MTG2, MTM1, MTO1, MTR2, MTW1, MUP3, MVD1, MXR2, MYG1, MZM1, NAB6, NBP1, NCA2, NCE101, NCS6, NDD1, NDI1, NEM1, NFT1, NGK1, NGL2, NGL3, NHP2, NIF3, NIP1, NMD3, NOG2, NOP19, NOP6, NOP8, NPR2, NPR3, NRE1, NUC1, NUD1, NUP188, NUT2, OAF1, OCA4, OCA5, OM45, OPI1, OPT1, OPT2, ORC4, OSH7, OST4, OST5, OSW7, OTU1, OTU2, OXA1, OXP1, OYE2, PAB1, PAC11, PAD1, PAF1, PAN1, PAN6, PAU1, PAU10, PAU11, PAU12, PAU13, PAU14, PAU15, PAU18, PAU19, PAU2, PAU20, PAU21, PAU24, PAU3, PAU4, PAU6, PAU9, PBI1, PBN1, PCA1, PCC1, PCK1, PCL1, PCM1, PDA1, PDE1, PDI1, PDP3, PDR18, PES4, PET122, PET494, PEX1, PEX11, PEX18, PEX2, PEX22, PEX29, PEX34, PFD1, PFK27, PFS1, PGA3, PGU1, PHO11, PHO12, PHO13, PHO4, PHO8, PHO84, PHO89, PHO90, PHR1, PIN3, PIP2, PIR5, PKH1, PLC1, PLM2, PML39, PMT4, POF1, POL5, POP2, POP3, PPG1, PPM2, PPS1, PRB1, PRC1, PRD1, PRE10, PRE4, PRE5, PRE8, PRI1, PRM1, PRP19, PRP21, PRP4, PRP8, PRR2, PRS3, PSE1, PSF3, PSP1, PTA1, PTC6, PTK1, PTP1, PTR2, PTR3, PUF6, PUG1, PUL3, PUL4, PUP2, PUS4, PWR1, |

| | |
|---|---|
| | PXA2, PXP3, PXR1, PZF1, QCR10, QCR2, QCR6, QCR7, QCR8, RAD10, RAD17, RAD2, RAD23, RAD24, RAD3, RAD4, RAI1, RBA50, RBD2, RBG1, RBH1, RCY1, RDR1, RDS1, RDT1, RET2, REV7, RFA3, RFC3, RGD2, RGD3, RHO1, RIB3, RIB4, RIE1, RIF1, RIF2, RIM101, RIM21, RIM4, RIX7, RMD6, RMD8, RME3, RML2, RMR1, RNH70, RNP1, RNQ1, ROF1, ROG3, RPC82, RPH1, RPL12A, RPL14B, RPL16A, RPL17B, RPL25, RPL29, RPL2A, RPL36B, RPL37B, RPL39, RPL40B, RPL6B, RPL8A, RPL8B, RPN12, RPO26, RPR2, RPS12, RPS14B, RPS1A, RPS20, RPS22A, RPS4A, RRI1, RRP40, RRP7, RRT5, RRT7, RSC8, RSC9, RSM19, RSM28, RTC1, RTF1, RTG2, RTT102, RUF21, RUF22, RUF23, SAC1, SAF1, SAM2, SAM3, SAM4, SAP155, SAP4, SAY1, SBP1, SCP1, SCW10, SCW4, SDD1, SDH2, SDH7, SDS22, SDT1, SEC11, SEC12, SEC15, SEC20, SEC21, SEC23, SEC39, SEC53, SEC65, SEN1, SET2, SGN1, SGV1, SHE10, SHR3, SHS1, SHU1, SIP2, SIR1, SIR3, SIT1, SKG1, SKI3, SKI8, SLD5, SLF1, SLH1, SLO1, SMC2, SMF1, SMT3, SMX3, SNA2, SNF1, SNF5, SNF6, SNM1, SNO2, SNO4, SNR191, SNR40, SNR49, SNR53, SNR64, SNR67, SNR68, SNR80, SNR84, SNR85, SNZ2, SOL1, SOL3, SOL4, SOM1, SOP4, SOR1, SPB1, SPC72, SPC97, SPE1, SPG3, SPO11, SPS1, SPS2, SPS22, SPT16, SPT2, SPT20, SQT1, SRB8, SRO9, SRP68, SRY1, SSB1, SSH1, SSK2, SSL2, SSP1, SST2, SSY5, STB5, STE20, STE50, STE6, STL1, STP2, STS1, SUF6, SUI3, SUL1, SUP6, SUT390, SUT532, SVP26, SWE1, SWI3, SWP82, TAD1, TAF1, TAF13, TAF8, TAN1, TCA17, TCD1, TDA1, TDA11, TDA5, TDA6, TDA9, TGL3, THI11, THI13, THI21, THI5, THP2, TIF3, TIM10, TIM22, TIM8, TMA108, TMT1, TNA1, TOG1, TOH1, TOR2, TPK1, TPM2, TPO3, TRF5, TRM11, TRM112, TRM13, TRP3, TRT2, TRX3, TSL1, TSR2, TTI2, TUB1, TUB2, TUB3, TUP1, TYC1, UBA1, UBI4, UBP11, UBP12, UBP15, UFO1, UGA4, ULI1, UPA1, UPA2, URA1, URA5, URC2, URN1, UTP14, UTR2, UTR4, UTR5, VAC17, VAM7, VAN1, VBA2, VBA3, VBA5, VEL1, VHS2, VID30, VIK1, VLD1, VMA6, VMA8, VMR1, VPS13, VPS3, VPS4, VPS52, VPS60, VPS68, VPS72, VPS9, VRG4, VTH1, VTH2, WHI4, WSC2, WSC4, YAH1, YAP1801, YAP3, YAP5, YAR1, YBT1, YCT1, YEF1, YGK3, YKT6, YLF2, YME2, YOR1, YPD1, YPS6, YPT1, YPT32, YRA2, YRF1-2, YRF1-3, YRF1-4, YRF1-5, YRF1-7, YRF1-8, YTA7, YVH1, ZDS2, ZIP2, ZNF1, ZNG1, ZOD1, ZPS1, ZRG17, ZRT1, ZUO1 |
| **gal2** | AAD10, AAD15, AAD4, ABP1, ABZ1, ABZ2, ACM1, ACO2, ACP1, ACS1, ACT1, ADD66, ADE4, ADE57, ADF1, ADH2, ADH4, ADH6, ADH7, ADK2, ADY3, AFG1, AGA1, AGP1, AGP3, AHC2, AIF1, AIM17, AIM2, AIM21, AIM33, AIM6, ALD4, ALG12, ALO1, ALR1, ALR2, AMF1, ANK1, ANP1, APA1, APE3, APL6, APM1, APM2, APM3, AQY2, AQY3, ARC35, ARG8, ARG81, ARN1, ARN2, ASG7, ASH1, ASN1, ATF1, ATG10, ATG18, ATG22, ATG23, ATG27, ATG32, ATG36, ATG41, ATG7, ATM1, ATP12, ATP15, ATP18, ATR1, AVT2, AXL2, AYT1, BAS1, BAT2, BBP1, BCK2, BDH1, BDH2, BET1, BET5, BGL2, BIK1, BIO2, BIO3, BIO4, BIO5, BIT2, BMH1, BNA6, BOL1, BOL2, BOL3, BRE4, BRE5, BRF1, BRR2, BRR6, BSC5, BSC6, BSD2, BUD17, BUD32, BUL2, BUR6, CAB4, CAC2, CAF120, CAF40, CAN1, CAR2, CBP1, CBP2, CBT1, CCA1, CCT2, CDC13, CDC14, CDC19, CDC23, CDC24, CDC26, CDC33, CDC39, CDC50, CDC6, CFD1, CFF1, CHA1, CHD1, CHK1, CIN2, CIN8, CIS3, CLA4, CLN2, CLN3, CMI7, CNA1, CNB1, CNE1, CNN1, COF1, COG1, COG3, COQ2, COQ5, COQ6, COS1, COS10, COS12, COS4, COS5, COS6, COS7, COS8, COS9, COX14, CPD1, CPR3, CPR8, CPS1, CRN1, CSE1, CSM1, CSS1, CSS2, CSS3, CTK3, CTP1, CTR2, CTR9, CUB1, CUE4, CUR1, CUS2, CWC2, CWC21, CWC22, CYC3, CYC7, DAD2, DAK2, DAL1, DAL2, DAL3, DAL4, DAL5, DAL7, DAL81, DAN1, DAN4, DAT1, DBP6, DBP8, DCG1, DCP1, DDI2, DFP4, DIA1, DIF1, DIG2, DIM1, DIP5, DJP1, DLD3, DMC1, DNA2, DNF1, DOA1, DOC1, DPB3, DPH2, DSE4, DSF1, DSN1, DSS1, DTD1, DUG1, DUG2, DUR3, DUS1, DYN3, EAP1, ECM1, ECM25, ECM29, ECM30, ECM32, ECM34, ECM7, ECO1, EDC1, EFG1, EFM1, EFM2, EGH1, EGO4, ELO1, ELP6, EMA35, EMC1, EMC3, EMC4, EMP47, ENB1, ENO1, ENO2, ENT4, ENV7, EPS1, ERG13, ERG20, ERG24, ERJ5, ERO1, ERR2, ERR3, ERV29, ERV46, ESF2, ESL2, ETP1, FAB1, FAR1, FAS2, FAT3, FAU1, FDH1, FET4, FET5, FEX1, FEX2, FIG2, FIT2, FIT3, FKS3, FLC2, FLO1, FLO10, FLO11, FLO5, FLX1, FMO1, FMP10, FMP27, FMP32, FMP33, FMP45, FOL2, FPK1, FPR4, FPS1, FRA1, FRD1, FRE2, FRE3, FRE4, FRE5, FRE6, FRE7, FRM2, FUM1, FUN12, FUN19, FUS1, FYV5, FZF1, GAB1, GAL4, GAS1, GAS4, GAT4, GCG1, GCN5, GCS1, GCV3, GDA1, GDH1, GDH2, GDH3, GEM1, GEX1, GEX2, GFD2, GID7, GIM5, GIN4, GIP4, GIT1, GLC8, GLK1, GLY1, GMC1, GMC2, GND1, GND2, GNP1, GON7, GOS1, GOT1, GPA1, GPB1, GPB2, GPH1, GPI13, GRC3, GRE2, GRX1, GRX4, GSR1, GTA1, GTR1, GTT1, GTT2, GUD1, GUS1, GUT1, GYP5, GYP7, HAB1, HAL5, HAP2, HAP5, HAS1, HAT2, HBN1, HBS1, HBT1, HCH1, HER2, HFI1, HFM1, HHY1, HIR3, HIS2, HIS4, HMG2, HMLALPHA2, HMRA1, HMRA2, HMS2, HO, HOL1, HOM6, HPA3, HRP1, HRT1, HSE1, HSH155, HSM3, HSP150, HSP32, HSP82, HSU1, HUA1, HUB1, HUT1, HXK1, HXK2, HXT11, HXT13, HXT15, HXT16, HXT17, HXT8, HXT9, HYM1, HYP2, HYR1, ICR1, IES6, IGD1, IMA1, IMA2, IMA3, IMA4, IMA5, IMD2, IMD3, IML1, INA22, INP51, IPA1, IQG1, IRC19, IRC24, IRC5, IRC6, IRC7, ISA1, ISC10, IST3, ITR1, IZH1, JEN1, JIP4, JIP5, JJJ2, JLP1, JNM1, KAP114, KAR3, KAR4, KAR9, KCC4, KEG1, KEL3, KGD4, KIN82, KIP3, KRE2, KRE6, KRE9, KRR1, LAA1, LAG1, LAM5, LCB1, LCD1, LDB18, LEU3, LEU5, LIP1, LOA1, LOS1, LPP1, LRE1, |

| | |
|---|---|
| | LRG1, LSB3, LSB5, LSM3, LYS1, LYS9, MAG2, MAK10, MAL12, MAL13, MAL31, MAL32, MAL33, MAN2, MATALPHA1, 46166, MBB1, MCH2, MCH4, MCM22, MCM3, MCO14, MDH2, MDL2, MDM1, MDM34, MED7, MEP3, MES1, MET10, MET16, MET28, MET5, MFG1, MGA1, MGA2, MGM101, MGR1, MHT1, MIA40, MIC10, MID1, MIG2, MIL1, MIN10, MLP1, MMP1, MMS1, MND2, MNN11, MNN4, MNN5, MNT2, MNT4, MOB2, MON2, MPC2, MPH1, MPH3, MPP6, MRC1, MRP2, MRP4, MRPL10, MRPL20, MRPL27, MRPL33, MRPL37, MRPL4, MRPL50, MRPS12, MRS1, MRS6, MRX15, MRX20, MRX6, MSA2, MSB3, MSC1, MSH3, MSL1, MSO1, MST1, MTC3, MTC7, MTD1, MTG2, MTM1, MTO1, MTR2, MTW1, MUP3, MVD1, MXR2, MYG1, MZM1, NAB6, NAS2, NBP1, NCA2, NCE101, NCE102, NCS6, NDD1, NDI1, NEM1, NFT1, NGK1, NGL2, NGL3, NHP2, NIF3, NIP1, NMD3, NOC4, NOG2, NOP19, NOP6, NOP8, NPR2, NPR3, NRE1, NSL1, NUC1, NUD1, NUP133, NUP188, NUT2, OAF1, OCA4, OCA5, OM45, OMA1, OPI1, OPT1, ORC4, OSH7, OST4, OST5, OSW7, OTU1, OTU2, OXP1, OYE2, PAB1, PAC11, PAF1, PAN1, PAN6, PAU1, PAU10, PAU12, PAU13, PAU14, PAU15, PAU18, PAU19, PAU2, PAU20, PAU21, PAU24, PAU3, PAU4, PAU6, PAU9, PBI1, PBN1, PCA1, PCC1, PCK1, PCL1, PCM1, PDA1, PDE1, PDE2, PDI1, PDP3, PDR18, PES4, PET494, PEX1, PEX11, PEX2, PEX22, PEX29, PEX34, PFD1, PFK27, PFS1, PGA3, PGU1, PHO11, PHO12, PHO13, PHO4, PHO8, PHO84, PHO89, PHO90, PHR1, PIN3, PIP2, PIR5, PKH1, PKH3, PLC1, PLM2, PML39, PMT4, POF1, POL5, POP2, POP3, POP5, PPG1, PPM2, PPS1, PRB1, PRC1, PRD1, PRE10, PRE4, PRE5, PRE8, PRI1, PRM1, PRP16, PRP19, PRP21, PRP3, PRP45, PRP8, PRR2, PRS3, PRT1, PSE1, PSF3, PSP1, PTA1, PTK1, PTP1, PTR2, PTR3, PUF6, PUG1, PUL3, PUL4, PUP2, PUS4, PWR1, PXA2, PXL1, PXP3, PXR1, QCR10, QCR6, QCR8, RAD10, RAD17, RAD2, RAD23, RAD24, RAD3, RAD4, RAI1, RBD2, RBG1, RBH1, RCY1, RDR1, RDS1, RDT1, RET2, REV7, RFA3, RFC3, RGD2, RGD3, RHO1, RIB3, RIB4, RIE1, RIF1, RIF2, RIM101, RIM15, RIM21, RIM4, RIX7, RMD6, RMD8, RME3, RML2, RMR1, RMT2, RNH70, RNP1, RNQ1, ROF1, ROG3, RPF2, RPH1, RPL12A, RPL14B, RPL16A, RPL17B, RPL18A, RPL22B, RPL25, RPL27B, RPL29, RPL2A, RPL36B, RPL37B, RPL39, RPL40B, RPL6B, RPL8A, RPL8B, RPN12, RPO41, RPR2, RPS12, RPS14B, RPS19A, RPS1A, RPS20, RPS22A, RPS4A, RRI1, RRP15, RRP40, RRP7, RRP9, RRT5, RRT7, RSC8, RSC9, RSM19, RSM28, RTC1, RTF1, RTG2, RTT102, RUF21, RUF22, RUF23, RVB2, SAC1, SAF1, SAM2, SAM3, SAM4, SAP155, SAP4, SAY1, SBP1, SCP1, SCW10, SCW4, SDC1, SDD1, SDH2, SDH8, SDS22, SDT1, SEC11, SEC12, SEC15, SEC20, SEC21, SEC39, SEC53, SEC65, SEN1, SET2, SGM1, SGN1, SGV1, SHE10, SHR3, SHS1, SHU1, SIP2, SIR1, SIR3, SIT1, SKG1, SKI8, SLD5, SLH1, SLO1, SMC2, SMF1, SNC1, SNF1, SNF12, SNF5, SNF6, SNM1, SNO2, SNO4, SNR13, SNR191, SNR40, SNR45, SNR49, SNR53, SNR64, SNR67, SNR68, SNR80, SNR85, SNZ2, SOL1, SOL3, SOL4, SOM1, SOP4, SOR1, SPB1, SPC72, SPC97, SPE1, SPG3, SPO11, SPS22, SPT16, SPT2, SPT20, SQT1, SRL3, SRO9, SRP40, SRP68, SRY1, SSB1, SSH1, SSK2, SSL2, SSO1, SSP1, SST2, STB5, STE20, STE50, STE6, STP2, STS1, SUE1, SUF6, SUI3, SUL1, SUP6, SUP8, SUT390, SUT532, SVP26, SWE1, SWI3, SWP82, TAD1, TAF1, TAF13, TAF8, TAN1, TAZ1, TCA17, TCD1, TDA1, TDA5, TDA6, TDA8, TDA9, TGL3, TGL4, THI11, THI13, THI21, THP2, TIF3, TIM10, TIM22, TIM8, TLG1, TMA108, TMT1, TNA1, TOG1, TOH1, TOR2, TPK1, TPM2, TPO1, TPO3, TRF5, TRM11, TRM112, TRM13, TRP3, TRS31, TRT2, TRX3, TRZ1, TSL1, TSR2, TTI2, TUB1, TUB2, TUB3, TUP1, TUS1, TVP38, TYC1, UBA1, UBI4, UBP11, UBP12, UBP15, UBX7, UFO1, UGA4, UGO1, ULI1, UPA1, UPA2, URA1, URA5, URM1, URN1, USV1, UTP14, UTR2, UTR4, UTR5, VAC17, VAM7, VAN1, VBA2, VBA3, VBA5, VEL1, VHS2, VID30, VIK1, VLD1, VMA11, VMA6, VMA8, VMR1, VPS13, VPS3, VPS501, VPS52, VPS60, VPS68, VPS72, VPS9, VRG4, VTH1, VTH2, VTS1, WAR1, WHI4, WSC2, WSC4, XPT1, YAH1, YAP3, YAP5, YAR1, YBT1, YCT1, YEF1, YGK3, YIA6, YKT6, YKU70, YLF2, YME2, YOR1, YPD1, YPS6, YPT1, YPT32, YRA2, YRF1-2, YRF1-3, YRF1-4, YRF1-5, YRF1-7, YRF1-8, YTA7, YVH1, ZDS2, ZIP2, ZNF1, ZNG1, ZOD1, ZPS1, ZRG17, ZRT1, ZUO1, YPD1, YPS6, YPT1, YPT32, YRA2, YRF1-1, YRF1-2, YRF1-3, YRF1-5, YRF1-6, YRF1-7, YRF1-8, YTA7, YVH1, ZDS2, ZIP2, ZNF1, ZNG1, ZOD1, ZPS1, ZRG17, ZRT1, ZUO1 |
| **gal3** | AAD10, AAD14, AAD15, AAD4, ABP1, ABZ1, ABZ2, ACM1, ACO2, ACP1, ACS1, ACT1, ADD66, ADE4, ADE57, ADF1, ADH2, ADH4, ADH6, ADH7, ADK2, ADY3, AFG1, AGA1, AGE1, AGP1, AGP3, AHC2, AIF1, AIM17, AIM2, AIM21, AIM33, AIM6, ALD4, ALG12, ALO1, ALR1, ALR2, AMF1, ANK1, ANP1, AOS1, APA1, APA2, APE3, API2, APL6, APM1, APM2, APM3, AQY1, AQY2, AQY3, ARC35, ARG8, ARG81, ARN1, ARN2, ASG1, ASG7, ASH1, ASN1, ATF1, ATG10, ATG13, ATG17, ATG18, ATG22, ATG23, ATG27, ATG32, ATG36, ATG41, ATG7, ATM1, ATP12, ATP15, ATP18, ATR1, AVT2, AXL2, AYT1, BAS1, BAT2, BBP1, BCK2, BDH1, BDH2, BEM2, BET1, BET2, BET5, BGL2, BIK1, BIO2, BIO3, BIO4, BIO5, BMH1, BNA6, BOL1, BOL2, BOL3, BRE4, BRE5, BRF1, BRR2, BRR6, BSC5, BSC6, BSD2, BSP1, BUD17, BUD32, BUL2, BUR6, CAB1, CAB4, CAC2, CAF120, CAF40, CAN1, CAR2, CBP1, CBP2, CBT1, CCA1, CCT2, CDC13, CDC14, CDC19, CDC23, CDC24, CDC26, CDC33, CDC39, CDC50, CDC6, CDC73, CFD1, CFF1, CHA1, CHC1, CHD1, CHK1, CIN2, CIN8, CIS3, CLA4, CLN2, CLN3, CMI7, |

| | |
|---|---|
| | CNA1, CNB1, CNE1, CNN1, COF1, COG1, COG3, COQ2, COQ5, COQ6, COS1, COS10, COS12, COS4, COS5, COS6, COS7, COS8, COS9, COX14, CPD1, CPR3, CPR8, CPS1, CRG1, CRN1, CSA1, CSE1, CSM1, CSM2, CSS1, CSS2, CSS3, CTK3, CTP1, CTR2, CTR9, CUB1, CUE4, CUR1, CUS2, CWC2, CWC21, CWC22, CYC3, CYC7, DAD2, DAK2, DAL1, DAL2, DAL3, DAL4, DAL5, DAL7, DAL81, DAN1, DAN4, DAS1, DAT1, DBP6, DBP8, DCG1, DCK1, DCP1, DDI2, DFP4, DIA1, DIF1, DIG2, DIM1, DIP5, DJP1, DLD3, DMC1, DNA2, DNF1, DOA1, DOC1, DPB2, DPB3, DPH2, DPM1, DSE4, DSF1, DSN1, DSS1, DTD1, DUG1, DUG2, DUR3, DUS1, DYN3, EAP1, ECM1, ECM25, ECM29, ECM30, ECM32, ECM34, ECM7, ECO1, EDC1, EFG1, EFM1, EGH1, EGO4, EGT2, ELO1, ELP6, EMA35, EMC1, EMC3, EMC4, EMI1, EMI2, EMP47, ENB1, ENO1, ENO2, ENT4, ENV7, EPS1, ERG13, ERG20, ERG24, ERJ5, ERO1, ERR2, ERR3, ERV29, ERV46, ESF2, ESL2, ETP1, EUG1, FAB1, FAR1, FAS2, FAT3, FAU1, FBP26, FDC1, FDH1, FET4, FET5, FEX1, FEX2, FIG2, FIT1, FIT2, FIT3, FKH1, FKS3, FLC2, FLO1, FLO10, FLO11, FLO5, FLO9, FLX1, FMO1, FMP10, FMP27, FMP32, FMP33, FMP45, FOL2, FPK1, FPR2, FPR4, FPS1, FRA1, FRD1, FRE2, FRE3, FRE4, FRE5, FRE6, FRE7, FRM2, FUM1, FUN12, FUN19, FUS1, FYV5, FZF1, GAB1, GAL4, GAS1, GAS4, GAT4, GCG1, GCN5, GCS1, GCV3, GDA1, GDB1, GDH1, GDH2, GDH3, GEM1, GEX1, GEX2, GFD2, GID7, GIM5, GIN4, GIT1, GLC8, GLK1, GLY1, GMC1, GMC2, GND1, GND2, GNP1, GON7, GOS1, GOT1, GPA1, GPB1, GPB2, GPH1, GPI13, GPI16, GRC3, GRE2, GRH1, GRX1, GRX2, GRX4, GSR1, GTA1, GTR1, GTT1, GTT2, GUD1, GUS1, GUT1, GYP5, GYP7, HAB1, HAL5, HAP2, HAS1, HAT2, HBN1, HBS1, HBT1, HCH1, HDA3, HER2, HFI1, HFM1, HHY1, HIR3, HIS2, HIS4, HLR1, HMG2, HMLALPHA2, HMRA1, HMRA2, HMS2, HO, HOL1, HOM6, HPA2, HPA3, HRP1, HRT1, HSE1, HSH155, HSP150, HSP31, HSP32, HSP82, HSU1, HUA1, HUB1, HUT1, HXK1, HXK2, HXT11, HXT13, HXT15, HXT16, HXT17, HXT8, HXT9, HYM1, HYP2, HYR1, ICR1, IES6, IGD1, IKI1, IMA1, IMA2, IMA3, IMA4, IMA5, IMD2, IMD3, IML1, INA22, INO1, INP51, IPA1, IQG1, IRC19, IRC24, IRC4, IRC5, IRC6, IRC7, ISA1, ISC10, IST3, ITR1, IZH1, JEN1, JIP4, JIP5, JJJ2, JLP1, JNM1, KAP114, KAR4, KAR9, KEG1, KEL3, KEX1, KGD4, KIN82, KIP3, KOG1, KRE2, KRE28, KRE6, KRE9, KRR1, LAA1, LAG1, LAM5, LCB1, LCD1, LDB18, LEU3, LEU5, LIP1, LOS1, LOT5, LPP1, LRE1, LRG1, LSB3, LSB5, LSC2, LSM3, LYS1, LYS9, MAG2, MAK10, MAL12, MAL13, MAL31, MAL32, MAL33, MAN2, MATALPHA1, 46166, MBB1, MCH2, MCH4, MCM22, MCM3, MCO14, MDH2, MDJ2, MDL2, MDM1, MDM34, MED7, MES1, MET10, MET16, MET28, MET5, MFG1, MGA1, MGA2, MGM101, MGR1, MHT1, MIA40, MIC10, MID1, MIG2, MIL1, MIN10, MLC2, MLP1, MMP1, MMS1, MND2, MNN11, MNN4, MNN5, MNT2, MNT4, MOB2, MON2, MPC2, MPC3, MPH1, MPH3, MRC1, MRP2, MRP4, MRPL10, MRPL20, MRPL27, MRPL33, MRPL4, MRPS12, MRS1, MRS6, MRX15, MRX20, MRX6, MSA2, MSB3, MSC1, MSH3, MSL1, MSO1, MST1, MTC3, MTC7, MTD1, MTG2, MTM1, MTO1, MTR2, MTW1, MUP3, MVD1, MXR2, MYG1, MZM1, NAB6, NAS2, NBP1, NCA2, NCE101, NCE102, NCS6, NDD1, NDI1, NEM1, NFT1, NGK1, NGL2, NGL3, NIF3, NIP1, NMD3, NOC4, NOG2, NOP19, NOP6, NOP8, NPR2, NPR3, NRE1, NSL1, NUC1, NUD1, NUP133, NUP188, NUT2, OAF1, OCA4, OCA5, OM45, OMA1, OPI1, OPT1, OPT2, ORC4, OSH7, OST4, OST5, OSW7, OTU1, OTU2, OXA1, OXP1, OYE2, PAB1, PAC11, PAD1, PAF1, PAN1, PAN6, PAU1, PAU10, PAU11, PAU12, PAU13, PAU14, PAU15, PAU18, PAU19, PAU2, PAU20, PAU21, PAU24, PAU3, PAU4, PAU6, PAU9, PBI1, PBN1, PCA1, PCC1, PCK1, PCL1, PCM1, PDA1, PDE1, PDI1, PDP3, PDR18, PES4, PET122, PET494, PEX1, PEX11, PEX18, PEX2, PEX21, PEX22, PEX29, PEX34, PEX6, PFA3, PFD1, PFK1, PFK27, PFS1, PGA3, PGU1, PHO11, PHO12, PHO13, PHO4, PHO8, PHO84, PHO89, PHO90, PHR1, PIN3, PIP2, PIR5, PKH1, PKH3, PLC1, PLM2, PML39, PMT4, POF1, POL5, POP2, POP3, POX1, PPG1, PPM2, PPS1, PRB1, PRC1, PRD1, PRE10, PRE4, PRE5, PRE8, PRI1, PRM1, PRP16, PRP19, PRP21, PRP3, PRP4, PRP8, PRR2, PRS3, PSE1, PSF3, PSP1, PTA1, PTH1, PTK1, PTP1, PTR2, PTR3, PUF6, PUG1, PUL3, PUL4, PUN1, PUP2, PUS4, PWR1, PXA2, PXL1, PXP3, PXR1, PZF1, QCR10, QCR2, QCR6, QCR7, QCR8, RAD10, RAD17, RAD2, RAD23, RAD24, RAD3, RAD4, RAI1, RBA50, RBD2, RBG1, RBH1, RCY1, RDR1, RDS1, RDT1, RET2, REV7, RFA3, RFC3, RGD2, RGD3, RHO1, RIB3, RIB4, RIE1, RIF1, RIF2, RIM101, RIM21, RIM4, RIX7, RMD6, RMD8, RME3, RML2, RMR1, RMT2, RNH70, RNP1, RNQ1, ROF1, ROG3, RPA34, RPC82, RPD3, RPF2, RPH1, RPL12A, RPL14B, RPL16A, RPL17B, RPL18A, RPL22B, RPL25, RPL27B, RPL29, RPL2A, RPL36B, RPL37B, RPL39, RPL40B, RPL6B, RPL8A, RPL8B, RPN12, RPN13, RPO26, RPO41, RPR2, RPS12, RPS14B, RPS19A, RPS1A, RPS20, RPS22A, RPS4A, RRI1, RRP15, RRP40, RRP7, RRT5, RRT7, RSC8, RSC9, RSM19, RSM28, RTC1, RTF1, RTG2, RTT102, RUF21, RUF22, RUF23, RVB2, SAC1, SAF1, SAM2, SAM3, SAM4, SAP155, SAP4, SAY1, SBP1, SCP1, SCW10, SCW4, SDA1, SDC1, SDD1, SDH2, SDH7, SDS22, SDT1, SEC11, SEC12, SEC15, SEC20, SEC21, SEC23, SEC39, SEC53, SEC65, SEN1, SET2, SGM1, SGN1, SGV1, SHE10, SHR3, SHS1, SHU1, SIP2, SIR1, SIR3, SIT1, SKG1, SKI3, SKI8, SLD5, SLF1, SLH1, SLO1, SMC2, SMF1, SMT1, SMT3, SMX3, SNA2, SNA3, SNF1, SNF5, SNF6, SNM1, SNO2, SNO4, SNR128, SNR13, SNR190, SNR191, SNR40, SNR45, SNR49, SNR53, SNR64, SNR67, SNR68, SNR80, SNR84, SNR85, SNZ2, SOL1, SOL3, SOL4, SOM1, SOP4, SOR1, SPB1, |

| | |
|---|---|
| | SPC72, SPC97, SPE1, SPG3, SPI1, SPO11, SPP382, SPS1, SPS2, SPS22, SPT16, SPT2, SPT20, SQT1, SRB8, SRL3, SRO9, SRP40, SRP68, SRY1, SSB1, SSH1, SSK2, SSL2, SSO1, SSP1, SST2, SSY5, STB5, STE20, STE50, STE6, STL1, STS1, SUE1, SUF6, SUI3, SUL1, SUP6, SUP8, SUT390, SUT532, SVP26, SWE1, SWI3, SWP82, TAD1, TAF1, TAF13, TAF8, TAN1, TAO3, TCA17, TCD1, TDA1, TDA11, TDA5, TDA6, TDA8, TDA9, TGL3, TGL4, THI11, THI13, THI21, THI5, THP2, TIF3, TIM10, TIM22, TIM8, TLG1, TMA108, TMT1, TNA1, TOG1, TOH1, TOR2, TPK1, TPM2, TPO1, TPO3, TRF5, TRM11, TRM112, TRM13, TRP3, TRS31, TRT2, TRX3, TRZ1, TSL1, TSR2, TTI2, TUB1, TUB2, TUB3, TUP1, TUS1, TVP38, TYC1, UBA1, UBI4, UBP11, UBP12, UBP15, UBP3, UFO1, UGA4, UGO1, ULI1, UPA1, UPA2, URA1, URA4, URA5, URC2, URM1, URN1, USV1, UTP14, UTR2, UTR4, UTR5, VAC17, VAM7, VAN1, VBA2, VBA3, VBA5, VEL1, VHS2, VID30, VIK1, VLD1, VMA11, VMA6, VMA8, VMR1, VPS13, VPS3, VPS35, VPS36, VPS4, VPS501, VPS52, VPS60, VPS68, VPS72, VPS9, VRG4, VTH1, VTH2, WAR1, WHI4, WSC2, WSC4, XPT1, YAH1, YAP1801, YAP1802, YAP3, YAP5, YAR1, YBT1, YCT1, YEF1, YGK3, YIA6, YKT6, YKU70, YLF2, YME2, YOR1, YPD1, YPS6, YPT1, YPT32, YRA2, YRF1-2, YRF1-3, YRF1-4, YRF1-5, YRF1-6, YRF1-7, YRF1-8, YTA7, YVH1, ZDS2, ZIP2, ZNF1, ZNG1, ZOD1, ZPS1, ZRG17, ZRT1, ZUO1, WSC4, XPT1, YAH1, YAP1801, YAP1802, YAP3, YAP5, YAR1, YAT1, YBT1, YCT1, YEF1, YGK3, YIA6, YKT6, YKU70, YLF2, YME2, YOR1, YPD1, YPS6, YPT1, YPT32, YRA2, YRF1-1, YRF1-2, YRF1-3, YRF1-5, YRF1-6, YRF1-7, YRF1-8, YTA7, YVH1, ZDS2, ZIP2, ZNF1, ZNG1, ZOD1, ZPS1, ZRG17, ZRT1, ZUO1 |
| **gal4** | AAD10, AAD15, AAD4, ABP1, ACM1, ACO2, ACP1, ACS1, ACT1, ADD66, ADE4, ADE57, ADF1, ADH2, ADH4, ADH6, ADH7, ADK2, ADY3, AFG1, AGA1, AGP3, AIF1, AIM17, AIM2, AIM29, AIM33, AIM6, ALD4, ALR1, ALR2, AMF1, ANK1, AOS1, APA1, APE3, APL6, APM1, APM2, APM3, AQY1, AQY2, AQY3, ARC35, ARG8, ARG81, ARN1, ARN2, ASG7, ATF1, ATG10, ATG13, ATG17, ATG18, ATG22, ATG23, ATG27, ATG36, ATG41, ATG7, ATM1, ATP12, ATP15, ATR1, AVT2, AXL2, AYT1, BAS1, BAT2, BBP1, BCK2, BDH1, BDH2, BEM2, BER1, BET2, BGL2, BIO2, BIO3, BIO4, BIO5, BLS1, BMH1, BNA6, BOL1, BOL2, BOL3, BRE4, BRE5, BRR2, BRR6, BSC5, BSC6, BSD2, BSP1, BUD32, BUL2, BUR6, CAB4, CAC2, CAF120, CAF40, CAN1, CAR2, CBP1, CBP2, CBT1, CCA1, CCT2, CDC13, CDC14, CDC19, CDC23, CDC24, CDC26, CDC33, CDC39, CDC50, CDC6, CDC73, CHA1, CHD1, CIN2, CIN8, CLA4, CLN2, CLN3, CNA1, CNB1, CNE1, CNN1, COF1, COG1, COG3, COQ2, COQ5, COQ6, COS1, COS10, COS12, COS4, COS5, COS6, COS7, COS8, COS9, COX14, CPD1, CPS1, CRN1, CSA1, CSE1, CSM2, CSS2, CSS3, CTK3, CTP1, CTR2, CTR3, CTR9, CUB1, CUE4, CUR1, CUS2, CWC2, CWC21, CWC22, CYC3, CYC7, DAD2, DAK2, DAL1, DAL2, DAL3, DAL4, DAL5, DAL7, DAL81, DAN1, DAN4, DAT1, DBP6, DBP8, DCG1, DCK1, DCP1, DDI2, DFP1, DFP2, DFP4, DIA1, DIF1, DIG2, DIM1, DIP5, DLD3, DMC1, DNA2, DNF1, DOA1, DOC1, DPB2, DPB3, DPH2, DPM1, DSE4, DSF1, DSN1, DTD1, DUG1, DUG2, DUR3, DUS4, EAP1, ECM1, ECM25, ECM29, ECM30, ECM32, ECM34, ECM4, ECM7, ECO1, EDC1, EFG1, EFM1, EGH1, EGO4, ELO1, ELP6, EMA35, EMC1, EMC3, EMC4, EMI1, EMP47, ENB1, ENO1, ENO2, ENT4, ENV7, ERG13, ERG20, ERG24, ERJ5, ERO1, ERR2, ERR3, ERV29, ERV46, ESF2, ESL2, ETP1, FAB1, FAS2, FAU1, FDH1, FET4, FET5, FEX1, FEX2, FIG2, FIT2, FIT3, FKS3, FLC2, FLO10, FLO11, FLO5, FLO9, FLX1, FMO1, FMP10, FMP32, FMP45, FOL2, FPK1, FPR4, FPS1, FRD1, FRE2, FRE3, FRE4, FRE5, FRE6, FRE7, FUM1, FUN12, FUN19, FYV5, FZF1, GAG1, GAL4, GAS1, GAS4, GAT4, GCG1, GCN5, GCS1, GCV3, GDA1, GDB1, GDH1, GDH2, GDH3, GEM1, GEX1, GEX2, GID7, GIM5, GIN4, GIP4, GIT1, GLC8, GLK1, GLY1, GMC1, GMC2, GND1, GND2, GNP1, GON7, GOS1, GPB1, GPB2, GPH1, GPI16, GRC3, GRE2, GRX2, GRX4, GTA1, GTR1, GTT1, GTT2, GUD1, GUS1, GUT1, GYP5, GYP7, HAB1, HAL5, HAP2, HAT2, HBS1, HBT1, HCH1, HDA3, HFI1, HFM1, HHY1, HIR3, HIS2, HMLALPHA2, HMRA1, HMRA2, HMS2, HO, HOL1, HOM6, HPA2, HPA3, HRP1, HRT1, HSE1, HSP32, HSP82, HSU1, HUA1, HUT1, HXK1, HXK2, HXT11, HXT13, HXT15, HXT16, HXT17, HXT8, HYM1, HYR1, ICR1, IES6, IGD1, IKI1, IMA1, IMA2, IMA3, IMA5, IMD2, IMD3, IML1, INA1, INA22, INO4, IPA1, IQG1, IRC19, IRC24, IRC5, IRC6, IRC7, ISC10, ITR1, IZH1, JEN1, JIP4, JIP5, JJJ2, JLP1, KAP114, KAR4, KAR9, KEG1, KEL3, KGD4, KIN82, KIP3, KOG1, KRE2, KRE6, KRE9, KRR1, KTD1, LAA1, LAG1, LAM5, LCD1, LDB18, LOS1, LPP1, LRE1, LRG1, LSB3, LSM3, LYS1, LYS9, MAG2, MAK10, MAL12, MAL13, MAL31, MAL32, MAL33, MAN2, MATALPHA1, 46166, MBB1, MCH2, MCH4, MCM22, MCO14, MDH2, MDL2, MDM1, MDM34, MDY2, MED7, MES1, MET10, MET16, MET28, MET5, MFG1, MGA1, MGA2, MGM101, MGR1, MHT1, MIA40, MIC10, MID1, MIG2, MIN10, MLC2, MLP1, MMP1, MMS1, MND2, MNN11, MNN4, MNN5, MNT2, MNT4, MOB2, MON2, MPC2, MPH3, MRC1, MRP2, MRP4, MRPL10, MRPL20, MRPL27, MRPL4, MRPS12, MRS1, MRS6, MRX15, MRX20, MRX6, MSA2, MSB3, MSB4, MSC1, MSH3, MSL1, MSN1, MSO1, MST1, MST28, MTC3, MTD1, MTG2, MTM1, MTO1, MTW1, MUP3, MVD1, MYG1, MZM1, NAB6, NCA2, NCE101, NCE102, NCS6, NDD1, NDI1, NFT1, NGK1, NGL3, NIF3, NIP1, NMD3, NOG2, NOP19, NOP6, NOP8, NPR2, NPR3, NRE1, NSL1, NUC1, NUD1, NUP133, NUP188, NUT2, OAF1, OCA4, OCA5, OM45, OMA1, OPI1, OPT1, OPT2, ORC4, OSH7, OST4, OST5, OSW7, OTU1, |

| | |
|---|---|
| | OTU2, OXA1, OXP1, OYE2, PAB1, PAC11, PAF1, PAP2, PAU1, PAU10, PAU12, PAU13, PAU14, PAU15, PAU19, PAU2, PAU21, PAU24, PAU3, PAU4, PAU6, PAU7, PAU8, PAU9, PBI1, PBN1, PCA1, PCC1, PCK1, PCL1, PCM1, PDA1, PDE1, PDE2, PDI1, PDR18, PES4, PET122, PET494, PEX1, PEX11, PEX2, PEX22, PEX29, PEX34, PFD1, PFK27, PFS1, PGA3, PGU1, PHO11, PHO12, PHO13, PHO4, PHO8, PHO84, PHO89, PHO90, PHR1, PIN3, PIP2, PKH1, PLC1, PLM2, PML39, PMT4, POF1, POL5, POP2, POP3, POP5, PPM2, PPS1, PRB1, PRD1, PRE10, PRE4, PRE5, PRE8, PRI1, PRM1, PRM9, PRP16, PRP19, PRP21, PRP4, PRP45, PRP8, PRR2, PRS3, PRT1, PSE1, PSF3, PSP1, PTA1, PTH1, PTH4, PTK1, PTP1, PTR2, PTR3, PUF6, PUG1, PUL3, PUL4, PUN1, PUP2, PUS4, PWR1, PXA2, PXL1, PXP3, PXR1, PZF1, QCR2, QCR6, QCR8, RAD10, RAD17, RAD2, RAD24, RAD3, RAD4, RAI1, RBD2, RBG1, RBH1, RCY1, RDR1, RDS1, RDT1, RET2, REV7, RFA3, RFC3, RGD2, RGD3, RHO1, RIB3, RIB4, RIE1, RIM101, RIM21, RIM4, RIX7, RMD6, RMD8, RME3, RML2, RMR1, RNH70, RNP1, ROF1, ROG3, RPC82, RPF2, RPH1, RPL12A, RPL14B, RPL16A, RPL17B, RPL18A, RPL22B, RPL25, RPL29, RPL2A, RPL31B, RPL36B, RPL37B, RPL39, RPL40B, RPL6B, RPL8A, RPL8B, RPN12, RPN13, RPO26, RPO41, RPR2, RPS12, RPS14B, RPS19A, RPS1A, RPS20, RPS22A, RPS4A, RRI1, RRI2, RRP40, RRT5, RSC8, RSC9, RSM19, RSM28, RTC1, RTF1, RTG2, RTT102, RUF21, RUF22, RUF23, RVB2, SAC1, SAF1, SAM2, SAM3, SAM4, SAP155, SAP4, SAY1, SBP1, SCP1, SCW10, SCW4, SDD1, SDH2, SDH7, SDS22, SDT1, SEC11, SEC15, SEC20, SEC21, SEC23, SEC39, SEC53, SEC65, SEI1, SEN1, SEO1, SET2, SGV1, SHE10, SHR3, SHR5, SHS1, SHU1, SIP2, SIR1, SIR3, SIT1, SKG1, SKI3, SKI8, SKM1, SLD5, SLH1, SLO1, SMC2, SMF1, SMT3, SMX3, SNC1, SNF1, SNF5, SNF6, SNM1, SNO2, SNO4, SNR191, SNR40, SNR49, SNR53, SNR64, SNR67, SNR68, SNR80, SNR85, SNZ2, SOL1, SOL3, SOL4, SOM1, SOP4, SOR1, SPB1, SPC72, SPC97, SPG3, SPI1, SPO11, SPP382, SPS22, SPT16, SPT2, SPT20, SQT1, SRL3, SRP40, SRP68, SRY1, SSB1, SSH1, SSL2, SSO1, SSP1, STB5, STE20, STE6, STS1, SUE1, SUF1, SUF6, SUI3, SUL1, SUP56, SUP6, SUT390, SUT532, SVP26, SWE1, SWH1, SWI3, SWP82, TAD1, TAF1, TAF13, TAF8, TAN1, TCA17, TDA5, TDA6, TGL3, TGL4, THI11, THI13, THI21, THP2, TIF3, TIM22, TIM8, TMA108, TMT1, TNA1, TOG1, TOH1, TOR2, TPK1, TPM2, TPO3, TRF5, TRM11, TRM112, TRM13, TRP3, TRT2, TRZ1, TSL1, TSR2, TTI2, TUB2, TUB3, TUS1, TVP38, TYC1, UBA1, UBI4, UBP11, UBP12, UBP15, UBP3, UFO1, UGA4, UIP3, ULI1, UPA1, UPA2, URA1, URA4, URA5, URN1, USV1, UTP14, UTP21, UTR2, VAC17, VAM7, VAN1, VBA2, VBA3, VBA5, VEL1, VHS2, VID30, VIK1, VIP1, VLD1, VMA11, VMA6, VMA8, VMR1, VPS13, VPS3, VPS36, VPS4, VPS501, VPS52, VPS60, VPS68, VPS72, VPS9, VRG4, VTH1, VTH2, VTS1, WHI4, WSC2, WSC4, YAH1, YAP3, YAP5, YAR1, YAT1, YBT1, YCT1, YEF1, YGK3, YKT6, YLF2, YME2, YOR1, YPD1, YPS6, YPT1, YPT32, YRA2, YRF1-2, YRF1-3, YRF1-4, YRF1-5, YRF1-6, YRF1-7, YRF1-8, YTA7, YVH1, ZDS2, ZEO1, ZIP2, ZNF1, ZOD1, ZPS1, ZRG17, ZRT1, ZUO1 |
| **gal5** | AAC1, AAD10, AAD14, AAD15, AAD4, ABF2, ABP1, ABZ1, ABZ2, ACM1, ACO2, ACP1, ACS1, ADD66, ADE4, ADE57, ADF1, ADH2, ADH3, ADH4, ADH6, ADH7, ADK2, ADY3, AEP1, AFG1, AGA1, AGE1, AGP1, AGP3, AHC2, AIF1, AIM17, AIM18, AIM2, AIM21, AIM32, AIM33, AIM46, AIM6, AIP1, ALD4, ALO1, ALR1, ALR2, AMD1, AMF1, ANK1, ANP1, AOS1, APA1, APA2, API2, APL6, APM1, APM2, APT1, AQY1, AQY2, AQY3, ARA2, ARC35, ARG7, ARG8, ARG80, ARN1, ARN2, ASC1, ASG7, ASH1, ATF1, ATG10, ATG13, ATG15, ATG18, ATG22, ATG27, ATG32, ATG36, ATG41, ATG7, ATM1, ATP11, ATP12, ATP15, ATP18, ATP25, AVO2, AVT2, AXL2, AYT1, BAS1, BAT1, BAT2, BBP1, BCK2, BDH1, BDH2, BEM2, BET2, BET5, BGL2, BIK1, BIO2, BIO3, BIO4, BIO5, BMH1, BNA6, BOL1, BOL2, BOL3, BRE4, BRE5, BRF1, BRR2, BRR6, BSC5, BSC6, BSP1, BUB2, BUD32, BUR6, BXI1, CAB1, CAB4, CAF120, CAF40, CAN1, CAR2, CAT2, CBP1, CBP2, CBT1, CCA1, CCT2, CDC13, CDC14, CDC19, CDC24, CDC26, CDC33, CDC39, CDC50, CDC6, CFF1, CGI121, CHA1, CHD1, CIN2, CIN8, CIS3, CLA4, CLN2, CLN3, CMP2, CNB1, CNE1, CNN1, COF1, COG1, COG3, COG8, COQ2, COQ6, COS1, COS10, COS12, COS4, COS5, COS6, COS7, COS8, COS9, CPD1, CPR3, CPR4, CPS1, CRG1, CSA1, CSE1, CSM1, CSM2, CSM3, CSS1, CSS2, CSS3, CTF13, CTF18, CTF8, CTR2, CTR9, CUB1, CUR1, CUS2, CWC21, CWC22, CYB2, CYC3, CYC7, DAK1, DAK2, DAL1, DAL2, DAL3, DAL4, DAL5, DAL7, DAL81, DAL82, DAN1, DAN4, DBP6, DBP8, DCG1, DCP1, DDI2, DFP4, DIA1, DIF1, DIG2, DIM1, DIP5, DJP1, DLD3, DMC1, DNF1, DOA1, DOC1, DPB2, DPH2, DPM1, DSE4, DSF1, DSN1, DTD1, DUG1, DUR3, DUS1, DYN3, EAP1, ECM1, ECM25, ECM29, ECM32, ECM34, ECM7, ECO1, EDC1, EFG1, EFM1, EGD2, EGH1, EGO2, EGO4, EGT2, ELO1, ELP6, EMA35, EMC1, EMC3, EMC4, EMI1, EMI2, EMP47, EMW1, ENB1, ENO1, ENO2, ENT4, ENV7, ERB1, ERG20, ERG24, ERG9, ERJ5, ERR2, ERR3, ERS1, ERV25, ERV29, ERV41, ERV46, ESF2, ESL2, ETP1, EUC1, EUG1, FAB1, FAR3, FAS2, FAT3, FAU1, FDC1, FDH1, FET3, FET4, FET5, FEX1, FEX2, FIG2, FIG4, FIT1, FIT2, FIT3, FKH1, FKS3, FLC2, FLO1, FLO10, FLO11, FLO5, FLO9, FLX1, FMO1, FMP10, FMP27, FMP32, FMP33, FMP45, FOL2, FOL3, FPK1, FPR2, FPR3, FPR4, FPS1, FRD1, FRE2, FRE3, FRE4, FRE5, FRE6, FRE7, FRM2, FUB1, FUM1, FUN12, FUN19, FUS1, FYV5, FZF1, GAB1, GAL4, GAL80, GAS1, GAS4, GAT4, GCG1, GCN5, GCS1, GCV3, GDA1, |

| | |
|---|---|
| | GDB1, GDH1, GDH2, GDH3, GEM1, GEX1, GEX2, GFD2, GID7, GIM5, GIN4, GIT1, GLC8, GLK1, GLY1, GMC1, GMC2, GND1, GND2, GNP1, GON7, GOS1, GOT1, GPB1, GPB2, GPH1, GPI16, GRC3, GRE2, GRH1, GRX1, GRX2, GRX4, GSF2, GSR1, GTA1, GTT1, GTT2, GUD1, GUS1, GUT1, GYP5, GYP7, HAL5, HAP2, HAS1, HAT2, HBN1, HBT1, HCH1, HDA3, HER2, HFD1, HFI1, HFM1, HHY1, HIR3, HIS2, HIS4, HLR1, HMG1, HMG2, HMLALPHA2, HMRA1, HMRA2, HMS2, HO, HOL1, HOM6, HPA2, HPA3, HRP1, HRT1, HSP150, HSP31, HSP32, HSP82, HSU1, HUA1, HUB1, HUG1, HUT1, HXK1, HXK2, HXT11, HXT13, HXT14, HXT15, HXT16, HXT17, HXT8, HXT9, HYM1, HYP2, HYR1, ICR1, IES6, IGD1, IKI1, ILV2, IMA1, IMA2, IMA3, IMA4, IMA5, IMD2, IMD4, IMG2, INA22, IOC4, IPA1, IQG1, IRC21, IRC24, IRC4, IRC5, IRC6, IRC7, ISC10, ISF1, IST3, ITR1, ITT1, IZH1, JEN1, JIP4, JIP5, JJJ2, JLP1, JNM1, KAP114, KAR4, KAR5, KAR9, KCC4, KEG1, KEL3, KGD4, KIN82, KIP3, KOG1, KRE1, KRE2, KRE28, KRE6, KRE9, KRI1, KRR1, LAA1, LAF1, LAG1, LAM5, LCB1, LCD1, LDB18, LEM3, LEU2, LEU3, LFT1, LIP1, LNP1, LOS1, LOT5, LPP1, LRE1, LRG1, LSB3, LSB5, LSC2, LSM3, LSM5, LYS1, LYS9, MAK10, MAL12, MAL13, MAL31, MAL32, MAN2, MATALPHA1, 46166, MBB1, MCH2, MCH4, MCM1, MCO14, MDH2, MDJ2, MDL2, MDM31, MDM34, MED11, MED7, MES1, MET10, MET16, MET28, MFG1, MFT1, MGA1, MGA2, MGM101, MGR1, MGR3, MHT1, MIA40, MIC10, MID1, MIG2, MIN10, MLC2, MLP1, MMP1, MMS1, MND2, MNL1, MNN11, MNN4, MNN5, MNT2, MNT4, MON2, MOT3, MPC3, MPH3, MRC1, MRP2, MRP4, MRPL10, MRPL20, MRPL4, MRPS12, MRPS18, MRS1, MRS6, MRX15, MRX20, MRX6, MSB3, MSH3, MSL1, MSO1, MST1, MTC3, MTC7, MTG1, MTG2, MTM1, MTO1, MTR2, MTW1, MUB1, MUP3, MVD1, MXR2, MYG1, MYO5, MZM1, NAM7, NAT4, NBL1, NBP1, NCA2, NCE101, NCE102, NCS6, NDC1, NDD1, NFS1, NFT1, NGK1, NIF3, NIP1, NMD3, NOG2, NOP19, NOP6, NOP8, NPL6, NPR2, NPR3, NRE1, NSE5, NSL1, NTE1, NUC1, NUD1, NUP116, NUT2, NVJ1, OAF1, OCA4, OCA5, OGG1, OM45, OMA1, OPI1, OPT1, OPT2, ORC1, ORC4, OST4, OST5, OST6, OSW7, OTU1, OTU2, OXA1, OXP1, OYE2, PAB1, PAC11, PAD1, PAN1, PAN6, PAT1, PAU1, PAU10, PAU11, PAU12, PAU13, PAU14, PAU15, PAU18, PAU19, PAU2, PAU20, PAU21, PAU24, PAU3, PAU4, PAU6, PAU9, PBI1, PBN1, PCC1, PCK1, PCL1, PCM1, PDA1, PDE1, PDI1, PDL32, PDP3, PDR18, PDS5, PEA2, PES4, PET122, PET494, PEX1, PEX11, PEX2, PEX21, PEX22, PEX29, PEX34, PEX6, PFA3, PFD1, PFK1, PFK27, PFS1, PFS2, PGM2, PGU1, PHA2, PHO11, PHO12, PHO13, PHO4, PHO8, PHO90, PHR1, PIF1, PIN3, PIR5, PKH1, PLC1, PLM2, PMT4, POB3, POF1, POL5, POP2, POP3, POP5, PPG1, PPM2, PPX1, PPZ1, PRB1, PRC1, PRD1, PRE4, PRE5, PRE8, PRI1, PRM1, PRM6, PRP16, PRP19, PRP21, PRP3, PRP39, PRP4, PRP45, PRR2, PRS1, PRS3, PSE1, PSF3, PSP1, PSP2, PTA1, PTC6, PTH1, PTK1, PTP1, PTR2, PTR3, PUF6, PUG1, PUL3, PUL4, PUP2, PUS4, PWR1, PXA2, PXL1, PXP3, PXR1, PZF1, QCR2, QCR6, QCR7, QCR8, RAD10, RAD17, RAD2, RAD23, RAD24, RAD3, RAD33, RAD4, RAD52, RAI1, RBA50, RBD2, RBG1, RBH1, RCF1, RCY1, RDR1, RDS1, RDT1, RET2, REV7, RFA2, RFA3, RFC3, RGD2, RGD3, RHO1, RIB3, RIB4, RIE1, RIF2, RIM101, RIM21, RIM4, RIM9, RIX1, RMD6, RMD8, RME3, RML2, RMR1, RNH70, RNP1, RNQ1, ROF1, ROG3, RPC82, RPD3, RPH1, RPL12A, RPL16A, RPL17B, RPL18A, RPL18B, RPL25, RPL29, RPL2A, RPL36B, RPL37B, RPL39, RPL40A, RPL40B, RPL6A, RPL6B, RPL8A, RPL8B, RPN10, RPN12, RPO26, RPR2, RPS12, RPS14B, RPS17A, RPS18B, RPS19A, RPS19B, RPS1A, RPS1B, RPS20, RPS22A, RPS4A, RPS4B, RRI1, RRN11, RRP40, RRP7, RRT5, RSA4, RSC8, RSE1, RSM19, RSM28, RTC1, RTF1, RTG2, RTT102, RUF22, RUF23, RVB2, SAC1, SAM2, SAM3, SAM37, SAM4, SAP155, SAP4, SAY1, SBP1, SCC4, SCH9, SCP1, SCW10, SCW4, SDA1, SDD1, SDD2, SDH2, SDH7, SDS22, SDT1, SEC11, SEC14, SEC15, SEC20, SEC21, SEC23, SEC39, SEC53, SEG1, SEN15, SET2, SET5, SGV1, SHE10, SHH3, SHR3, SHS1, SHU1, SIR1, SIR3, SIT1, SKG1, SKI3, SKI8, SKN7, SKP2, SLD5, SLF1, SLH1, SLN1, SLO1, SMA2, SMC2, SMF1, SML1, SMN1, SMT3, SMX3, SNA2, SNF1, SNF6, SNM1, SNO1, SNO2, SNO4, SNR191, SNR24, SNR40, SNR49, SNR53, SNR54, SNR64, SNR67, SNR68, SNR80, SNR84, SNZ1, SNZ2, SOL1, SOL2, SOL4, SOM1, SOP4, SOR1, SOV1, SPB1, SPC2, SPC24, SPC72, SPC97, SPE1, SPG3, SPG4, SPI1, SPO11, SPS1, SPS2, SPS22, SPT15, SPT2, SPT20, SQT1, SRB8, SRC1, SRL3, SRO9, SRP40, SRP68, SRT1, SRY1, SSB1, SSK22, SSL2, SSO1, SSP1, SST2, STB1, STB2, STB5, STE20, STE50, STE6, STL1, STS1, STV1, SUE1, SUF6, SUF7, SUI3, SUP5, SUP53, SUP6, SUR7, SUT390, SUT532, SVP26, SWE1, SWI3, SWP82, TAD1, TAF1, TAF11, TAN1, TCA17, TCB3, TDA1, TDA6, TDA8, TDA9, TEM1, TGL3, TGL4, THI11, THI13, THI21, THI5, TIF3, TIM22, TMA108, TMT1, TNA1, TOG1, TOH1, TOR2, TOS6, TPK1, TPM2, TPO3, TRF5, TRM11, TRM112, TRM13, TRM9, TRP3, TRS31, TRT2, TRX3, TSA1, TUB1, TUP1, TVP18, TVP38, UBA1, UBI4, UBP11, UBP12, UBP15, UBP3, UBX2, UBX4, UFO1, ULI1, UNG1, UPA1, UPA2, URA1, URC2, URN1, USA1, UTP14, UTP15, UTP9, UTR2, UTR4, UTR5, VAC17, VAM7, VBA1, VBA3, VBA5, VEL1, VHS2, VID30, VIK1, VLD1, VMA11, VMA6, VMA8, VMR1, VNX1, VPS13, VPS20, VPS3, VPS4, VPS52, VPS60, VPS68, VPS71, VPS72, VRG4, VTH1, VTH2, WAR1, WHI4, WSC2, WSC4, YAH1, YAP1802, YAP3, YAP5, YAR1, YBT1, YCT1, YEF1, YET2, YGK3, YKT6, YKU80, YLF2, YMD8, YME2, YML6, YOR1, YOX1, YPD1, YPK2, YPS6, YPT11, YPT32, YRA2, YRF1-2, YRF1-3, YRF1- |

| | |
|---|---|
| | 4, YRF1-5, YRF1-6, YRF1-7, YRF1-8, YTA12, YTA7, YVH1, ZIM17, ZIP2, ZNF1, ZOD1, ZPS1, ZRG17, ZRT1, ZUO1, YOX1, YPD1, YPK2, YPS6, YPT1, YPT11, YPT32, YRA2, YRF1-1, YRF1-2, YRF1-3, YRF1-5, YRF1-6, YRF1-7, YRF1-8, YTA12, YTA7, YVH1, ZIM17, ZIP2, ZNF1, ZOD1, ZPS1, ZRG17, ZRT1, ZUO1 |
| **gal6** | AAD10, AAD14, AAD15, AAD4, ABP1, ABZ1, ACM1, ACO2, ACP1, ACS1, ACT1, ADD66, ADE4, ADE57, ADF1, ADH2, ADH4, ADH6, ADH7, ADK2, ADY3, AFG1, AGA1, AGE1, AGP1, AGP3, AGX1, AHC2, AIF1, AIM17, AIM2, AIM21, AIM33, AIM6, ALD4, ALG12, ALO1, ALR1, ALR2, AMF1, ANK1, ANP1, AOS1, APA1, APA2, API2, APL6, APM1, APM2, AQY1, AQY2, AQY3, ARC35, ARE2, ARG8, ARG81, ARN1, ARN2, ASG1, ASG7, ASH1, ATF1, ATG10, ATG13, ATG18, ATG22, ATG23, ATG27, ATG32, ATG36, ATG41, ATG7, ATM1, ATP12, ATP15, ATP18, ATP23, ATR1, ATS1, AVT2, AXL2, AYT1, BAS1, BAT2, BBP1, BCK2, BDH1, BDH2, BEM2, BET1, BET2, BGL2, BIK1, BIO2, BIO3, BIO4, BIO5, BMH1, BNA6, BOL1, BOL2, BOL3, BRE4, BRE5, BRF1, BRR2, BRR6, BSC5, BSC6, BSP1, BUD17, BUD32, BUL2, BUR6, CAB1, CAB4, CAC2, CAF120, CAF40, CAN1, CAR2, CBP1, CBP2, CBT1, CCA1, CCR4, CCT2, CDC13, CDC14, CDC19, CDC24, CDC26, CDC33, CDC39, CDC50, CDC6, CFD1, CHA1, CHD1, CIN2, CIN8, CIS3, CLA4, CLN2, CLN3, CMI7, CNA1, CNB1, CNE1, CNN1, COF1, COG1, COG3, COQ2, COQ5, COQ6, COS1, COS10, COS12, COS4, COS5, COS6, COS7, COS8, COS9, COX14, CPD1, CPR8, CPS1, CSA1, CSE1, CSM1, CSM2, CSS1, CSS2, CSS3, CTK3, CTR2, CTR9, CUB1, CUE4, CUR1, CUS2, CWC2, CWC21, CWC22, CYC3, CYC7, CYS3, DAK2, DAL1, DAL2, DAL3, DAL4, DAL5, DAL7, DAL81, DAN1, DAN4, DAT1, DBP6, DBP8, DCG1, DCP1, DDI2, DEP1, DFP2, DFP4, DIA1, DIF1, DIG2, DIM1, DIP5, DJP1, DLD3, DMC1, DNF1, DOA1, DOC1, DPB2, DPH2, DPM1, DRS2, DSE4, DSF1, DSN1, DTD1, DUG1, DUR3, DYN3, EAP1, ECM1, ECM25, ECM29, ECM30, ECM32, ECM34, ECM7, ECO1, EDC1, EFG1, EFM1, EGH1, EGO2, EGO4, EGT2, ELO1, ELP6, EMA35, EMC1, EMC3, EMC4, EMI1, EMI2, EMP47, ENB1, ENO1, ENO2, ENT4, ENV7, EPS1, ERG13, ERG20, ERG24, ERJ5, ERO1, ERR2, ERR3, ERS1, ERV29, ERV46, ESF2, ESL2, ETP1, EUG1, FAB1, FAS2, FAT3, FAU1, FDC1, FDH1, FET4, FET5, FEX1, FEX2, FIG2, FIG4, FIT1, FIT2, FIT3, FKH1, FKS3, FLC2, FLO1, FLO10, FLO11, FLO5, FLO9, FLX1, FMO1, FMP10, FMP27, FMP32, FMP33, FMP45, FOL2, FPK1, FPR2, FPR4, FPS1, FRD1, FRE2, FRE3, FRE4, FRE5, FRE6, FRE7, FRM2, FRT2, FUB1, FUM1, FUN12, FUN19, FUN26, FUN30, FUS1, FYV5, FZF1, GAB1, GAL4, GAS1, GAS4, GAT4, GCG1, GCN5, GCS1, GCV3, GDA1, GDB1, GDH1, GDH2, GDH3, GEM1, GEX1, GEX2, GFD2, GID7, GIM5, GIN4, GIP4, GIT1, GLC8, GLK1, GLY1, GMC1, GMC2, GND1, GND2, GNP1, GON7, GOS1, GPB1, GPB2, GPH1, GRC3, GRE2, GRH1, GRX1, GRX2, GRX4, GSR1, GTA1, GTR1, GTT1, GTT2, GUD1, GUS1, GUT1, GYP5, GYP7, HAC1, HAL5, HAP2, HAT2, HBN1, HBT1, HCH1, HDA3, HFI1, HFM1, HHY1, HIR3, HIS2, HIS4, HLR1, HMG2, HMLALPHA2, HMRA1, HMRA2, HMS2, HO, HOL1, HPA2, HPA3, HRA1, HRP1, HRT1, HSP150, HSP31, HSP32, HSP82, HSU1, HUA1, HUB1, HUT1, HXK1, HXK2, HXT11, HXT13, HXT15, HXT16, HXT17, HXT8, HXT9, HYM1, HYP2, HYR1, ICR1, IES6, IGD1, IKI1, IMA1, IMA2, IMA3, IMA4, IMA5, IMD2, IMD3, INA22, INP51, IPA1, IQG1, IRC24, IRC4, IRC5, IRC6, IRC7, ISC10, IST3, ITR1, IZH1, JEN1, JIP4, JIP5, JJJ2, JLP1, KAP114, KAR4, KAR9, KEG1, KEL3, KGD4, KIN82, KIP3, KOG1, KRE2, KRE28, KRE6, KRE9, KRR1, LAA1, LAG1, LAM5, LCB1, LCD1, LDB18, LDS1, LEU3, LIP1, LOS1, LOT5, LPP1, LRE1, LRG1, LSB3, LSB5, LSC2, LSM3, LTE1, LYS1, LYS9, MAK10, MAK16, MAL12, MAL13, MAL31, MAL32, MAN2, MATALPHA1, 46166, MBB1, MCH2, MCO14, MDH2, MDJ2, MDL2, MDM1, MDM34, MED7, MES1, MET10, MET16, MET28, MFG1, MGA1, MGA2, MGM101, MGR1, MHT1, MIA40, MIC10, MID1, MIG2, MIL1, MIN10, MLC2, MLP1, MLP2, MMP1, MMS1, MND2, MNN11, MNN4, MNN5, MNT2, MNT4, MOB2, MON2, MPC3, MPH1, MPH3, MPP6, MRC1, MRP2, MRP4, MRPL10, MRPL20, MRPL4, MRPL50, MRPS12, MRS1, MRS6, MRX15, MRX20, MRX6, MSB3, MSC1, MSH3, MSL1, MSO1, MST1, MST28, MTC3, MTC7, MTG2, MTM1, MTO1, MTR2, MTW1, MUP3, MVD1, MXR2, MYG1, MYO4, MZM1, NAB6, NAS2, NBP1, NCA2, NCE101, NCE102, NCS6, NDD1, NDI1, NFT1, NGK1, NGL3, NIF3, NIP1, NMD3, NOG2, NOP19, NOP6, NOP8, NPR2, NPR3, NRE1, NSL1, NTG1, NUC1, NUD1, NUP188, NUT2, OAF1, OCA4, OCA5, OM45, OMA1, OPI1, OPT1, OPT2, ORC4, OST4, OST5, OSW7, OTU1, OTU2, OXA1, OXP1, OYE2, PAB1, PAC11, PAD1, PAN1, PAN6, PAT1, PAU1, PAU10, PAU11, PAU12, PAU13, PAU14, PAU15, PAU18, PAU19, PAU2, PAU20, PAU21, PAU24, PAU3, PAU4, PAU6, PAU9, PBI1, PBN1, PCC1, PCK1, PCL1, PCM1, PDA1, PDE1, PDI1, PDP3, PDR18, PES4, PET122, PET494, PEX1, PEX11, PEX2, PEX21, PEX22, PEX29, PEX34, PEX6, PFA3, PFD1, PFK1, PFK27, PFS1, PGA3, PGU1, PHO11, PHO12, PHO13, PHO4, PHO8, PHO84, PHO90, PHR1, PIN3, PIP2, PIR5, PKH1, PLC1, PLM2, PML39, PMT2, PMT4, POF1, POL5, POP2, POP3, POP5, PPG1, PPM2, PRB1, PRC1, PRD1, PRE10, PRE4, PRE5, PRE8, PRI1, PRM1, PRM9, PRP16, PRP19, PRP21, PRP4, PRP45, PRR2, PRS3, PSE1, PSF3, PSK1, PSP1, PTA1, PTC6, PTK1, PTP1, PTR2, PTR3, PUF6, PUG1, PUL3, PUL4, PUP2, PUS4, PWR1, PXA2, PXL1, PXP3, PXR1, PZF1, QCR2, QCR6, QCR7, QCR8, RAD10, RAD17, RAD2, RAD23, RAD24, RAD3, RAD4, RAI1, RBA50, RBD2, RBG1, RBH1, RCF2, RCY1, RDR1, RDS1, RDT1, RET2, REV7, RFA3, RFC3, RGD2, RGD3, RHO1, RIB3, RIB4, |

| | |
|---|---|
| | RIE1, RIF2, RIM101, RIM15, RIM21, RIM4, RMD6, RMD8, RME3, RML2, RMR1, RNH70, RNP1, RNQ1, ROF1, ROG3, RPC82, RPD3, RPH1, RPL12A, RPL16A, RPL17B, RPL22B, RPL25, RPL29, RPL2A, RPL36B, RPL37B, RPL39, RPL40A, RPL40B, RPL6B, RPL8A, RPL8B, RPN12, RPO26, RPO41, RPR2, RPS12, RPS14B, RPS1A, RPS20, RPS22A, RPS4A, RRI1, RRP40, RRP7, RRT5, RSC8, RSC9, RSM19, RSM28, RTC1, RTF1, RTG2, RTT102, RUF21, RUF22, RUF23, RVB2, SAC1, SAM2, SAM3, SAM4, SAP155, SAP4, SAW1, SAY1, SBP1, SCP1, SCW10, SCW4, SDA1, SDD1, SDH2, SDH7, SDS22, SDT1, SEC11, SEC12, SEC15, SEC20, SEC21, SEC23, SEC39, SEC53, SEC65, SET2, SGN1, SGV1, SHE10, SHR3, SHS1, SHU1, SIP2, SIR1, SIR3, SIT1, SKG1, SKI3, SKI8, SLD5, SLF1, SLH1, SLN1, SLO1, SMC2, SMF1, SMT3, SMX3, SNA2, SNC1, SNF1, SNF12, SNF6, SNM1, SNO2, SNO4, SNR191, SNR40, SNR49, SNR53, SNR64, SNR67, SNR68, SNR80, SNR84, SNR85, SNZ2, SOL1, SOL2, SOL4, SOM1, SOP4, SOR1, SPB1, SPC72, SPC97, SPE1, SPG3, SPO11, SPS1, SPS2, SPS22, SPT2, SPT20, SQT1, SRB8, SRL3, SRO9, SRP40, SRP68, SRY1, SSB1, SSK2, SSL2, SSO1, SSP1, SST2, STB5, STE20, STE50, STE6, STL1, STS1, SUE1, SUF6, SUI3, SUP6, SUT390, SUT532, SVP26, SWE1, SWI3, SWP82, SYN8, TAD1, TAF1, TAF13, TAF8, TAN1, TCA17, TDA6, TGL3, TGL4, THI11, THI13, THI21, THI5, THP2, TIF3, TIM22, TIM23, TMA108, TMT1, TNA1, TOG1, TOH1, TOR2, TOS6, TPD3, TPK1, TPM2, TPO3, TRF5, TRM11, TRM112, TRM13, TRP3, TRT2, TRX3, TSL1, TSR2, TUB1, TUB2, TUB3, TUP1, TVP38, UBA1, UBI4, UBP11, UBP12, UBP15, UFO1, UGA4, ULI1, UPA1, UPA2, URA1, URA5, URC2, URN1, UTP14, UTR2, UTR4, UTR5, VAC17, VAM7, VAN1, VBA3, VBA5, VEL1, VHS2, VID30, VIK1, VLD1, VMA11, VMA6, VMA8, VMR1, VPS13, VPS3, VPS4, VPS52, VPS60, VPS68, VPS72, VPS9, VRG4, VTH1, VTH2, WHI4, WSC2, WSC4, YAH1, YAP1802, YAP3, YAP5, YAR1, YAT1, YBT1, YCT1, YEF1, YGK3, YIA6, YKT6, YLF2, YME2, YOR1, YPD1, YPS6, YPT1, YPT32, YRA2, YRF1-2, YRF1-3, YRF1-4, YRF1-5, YRF1-6, YRF1-7, YRF1-8, YTA7, YVH1, ZDS2, ZIP2, ZNF1, ZNG1, ZOD1, ZPS1, ZRG17, ZRT1, ZUO1 |

**Table S4:** Number of unique genes undergoing deletion or duplication in each replicate line.

**Note:**

Genes undergoing deletion or duplication at each time point were merged and only the number of unique genes were counted - overlapping genes were removed. Numbers for each replicate line is listed separately. Total number of genes in the gff file is 6281. This is used to calculate percentage of genes deleted or duplicated.

| Replicate | Number of genes undergoing deletion | Number of genes undergoing duplication | Percentage of genes undergoing deletion | Percentage of genes undergoing duplication |
|---|---|---|---|---|
| glu1 | 526 | 271 | 8.37 | 4.31 |
| glu2 | 548 | 277 | 8.72 | 4.41 |
| glu3 | 351 | 540 | 5.59 | 8.60 |
| glu4 | 726 | 560 | 11.56 | 8.92 |
| glu5 | 490 | 261 | 7.80 | 4.16 |
| glu6 | 450 | 262 | 7.16 | 4.17 |
| gal1 | 235 | 1189 | 3.74 | 18.93 |
| gal2 | 234 | 1228 | 3.73 | 19.55 |
| gal3 | 238 | 1255 | 3.79 | 19.98 |
| gal4 | 256 | 1260 | 4.08 | 20.06 |
| gal5 | 237 | 1262 | 3.77 | 20.09 |
| gal6 | 251 | 1132 | 4.00 | 18.02 |

**Table S5:** Number of unique genes undergoing deletion or duplication in each replicate line.

**Note:**

Genes undergoing deletion or duplication at each time point were counted in each replicate. Numbers for each replicate line, at each time point is listed separately. Total number of genes in the gff file is 6281. This is used to calculate percentage of genes deleted or duplicated.

**Glucose-evolved:**

| Time | Replicate | Number of genes undergoing deletion | Number of genes undergoing duplication | Percentage of genes undergoing deletion | Percentage of genes undergoing duplication |
|---|---|---|---|---|---|
| 300gen | glu1 | 265 | 110 | 4.22 | 1.75 |
| | glu2 | 175 | 0 | 2.79 | 0.00 |
| | glu3 | 233 | 450 | 3.71 | 7.16 |
| | glu4 | 486 | 390 | 7.74 | 6.21 |
| | glu5 | 211 | 46 | 3.36 | 0.73 |
| | glu6 | 391 | 82 | 6.23 | 1.31 |
| 600gen | glu1 | 41 | 34 | 0.65 | 0.54 |
| | glu2 | 378 | 246 | 6.02 | 3.92 |
| | glu3 | 223 | 415 | 3.55 | 6.61 |
| | glu4 | 268 | 402 | 4.27 | 6.40 |
| | glu5 | 237 | 146 | 3.77 | 2.32 |
| | glu6 | 2 | 56 | 0.03 | 0.89 |
| 900gen | glu1 | 207 | 153 | 3.30 | 2.44 |
| | glu2 | 295 | 52 | 4.70 | 0.83 |
| | glu3 | 106 | 463 | 1.69 | 7.37 |
| | glu4 | 123 | 510 | 1.96 | 8.12 |
| | glu5 | 22 | 131 | 0.35 | 2.09 |
| | glu6 | 265 | 223 | 4.22 | 3.55 |
| 1200gen | glu1 | 229 | 119 | 3.65 | 1.89 |
| | glu2 | 215 | 163 | 3.42 | 2.60 |
| | glu3 | 203 | 423 | 3.23 | 6.73 |
| | glu4 | 196 | 43 | 3.12 | 0.68 |
| | glu5 | 237 | 97 | 3.77 | 1.54 |
| | glu6 | 307 | 79 | 4.89 | 1.26 |

**Galactose-evolved:**

| Time | Replicate | Number of genes undergoing deletion | Number of genes undergoing duplication | Percentage of genes undergoing deletion | Percentage of genes undergoing duplication |
|---|---|---|---|---|---|
| 300gen | gal1 | 213 | 722 | 3.39 | 11.49 |
| | gal2 | 215 | 585 | 3.42 | 9.31 |
| | gal3 | 213 | 779 | 3.39 | 12.40 |
| | gal4 | 219 | 708 | 3.49 | 11.27 |
| | gal5 | 219 | 819 | 3.49 | 13.04 |
| | gal6 | 220 | 901 | 3.50 | 14.34 |
| 600gen | gal1 | 217 | 1066 | 3.45 | 16.97 |
| | gal2 | 219 | 1041 | 3.49 | 16.57 |
| | gal3 | 236 | 1029 | 3.76 | 16.38 |
| | gal4 | 216 | 1169 | 3.44 | 18.61 |
| | gal5 | 213 | 804 | 3.39 | 12.80 |
| | gal6 | 230 | 788 | 3.66 | 12.55 |
| 900gen | gal1 | 233 | 1035 | 3.71 | 16.48 |
| | gal2 | 214 | 1014 | 3.41 | 16.14 |
| | gal3 | 232 | 1049 | 3.69 | 16.70 |
| | gal4 | 233 | 661 | 3.71 | 10.52 |
| | gal5 | 235 | 994 | 3.74 | 15.83 |
| | gal6 | 235 | 939 | 3.74 | 14.95 |
| 1200gen | gal1 | 235 | 1053 | 3.74 | 16.76 |
| | gal2 | 224 | 1072 | 3.57 | 17.07 |
| | gal3 | 234 | 1152 | 3.73 | 18.34 |
| | gal4 | 240 | 1004 | 3.82 | 15.98 |
| | gal5 | 221 | 1168 | 3.52 | 18.60 |
| | gal6 | 233 | 1067 | 3.71 | 16.99 |

1. Variant calling pipeline

```
#!/usr/bin/env python3

import os
import subprocess
import argparse
import shutil

def run_command(command, step_folder):
    """Run a shell command with error handling."""
    print(f"Running: {command}")
    subprocess.run(command, shell=True, check=True, cwd=step_folder)

def cleanup_sample_files(sample, output_base, folder_names, steps_to_clean=range(1,
8)):
    """Delete files for the current sample in folders 1-7."""
    for step in steps_to_clean:
        step_folder = os.path.join(output_base, folder_names[step])
        if not os.path.exists(step_folder):
            continue
        for filename in os.listdir(step_folder):
            if sample in filename:  # only delete files for this sample
                file_path = os.path.join(step_folder, filename)
                try:
                    if os.path.isfile(file_path) or os.path.islink(file_path):
                        os.unlink(file_path)
                    elif os.path.isdir(file_path):
                        shutil.rmtree(file_path)
                except Exception as e:
                    print(f"Could not delete {file_path}. Reason: {e}")

def main():
    parser = argparse.ArgumentParser(description="WGS Pipeline with GVCF Output")
    parser.add_argument("--base_dir", required=True, help="Path to input sequencing
data")
    parser.add_argument("--output_base", required=True, help="Path to output
directory")
    parser.add_argument("--reference_genome", required=True, help="Path to reference
genome")
    args = parser.parse_args()

    base_dir = args.base_dir
    output_base = args.output_base
    reference_genome = args.reference_genome
    picard_cmd = "java -jar /home/prachitha/WGS/picard.jar"
    gatk_cmd = "java -jar
/home/prachitha/WGS/gatk-4.4.0.0/gatk-package-4.4.0.0-local.jar"

    samples = [f"A{i}" for i in range(1, 49)]
```

```
    folder_names = {
        1: "1_Trimming_A",
        2: "2_Quality_Check_A",
        3: "3_Adapter_Removal_A",
        4: "4_Alignment_A",
        5: "5_Filtering_A",
        6: "6_Sorting_A",
        7: "7_Read_Group_And_MarkDuplicates_A",
        8: "8_Sorting_And_Indexing_A",
        9: "9_Variant_Calling_A",
        10: "10_Assembled_Reads_A",
        11: "11_GVCF_Calling_A"
    }

    # Create output directories
    for step in folder_names:
        os.makedirs(os.path.join(output_base, folder_names[step]), exist_ok=True)

    # File to track skipped samples
    skipped_file = os.path.join(output_base, "skipped_samples.txt")
    with open(skipped_file, "w") as sf:
        sf.write("Samples skipped due to missing FASTQ files:\n")

    for sample in samples:
        print(f"\n=== Processing {sample} ===")

        r1_file = os.path.join(base_dir, f"{sample}_WGS_BatchEXT24_R1.fastq.gz")
        r2_file = os.path.join(base_dir, f"{sample}_WGS_BatchEXT24_R2.fastq.gz")

        # Skip if input files missing
        if not os.path.exists(r1_file) or not os.path.exists(r2_file):
            print(f"Skipping {sample} – FASTQ files not found.")
            with open(skipped_file, "a") as sf:
                sf.write(f"{sample}\n")
            continue

        # --------------------
        # Step 1: Trimming
        trimmed_r1 = os.path.join(output_base, folder_names[1],
f"{sample}_R1_trimmed.fastq")
        trimmed_r2 = os.path.join(output_base, folder_names[1],
f"{sample}_R2_trimmed.fastq")
        command = f"fastp -i {r1_file} -I {r2_file} -o {trimmed_r1} -O {trimmed_r2}
--html {sample}_fastp_report.html --json {sample}_fastp_report.json --thread 4"
        run_command(command, os.path.join(output_base, folder_names[1]))

        # Step 2: FastQC
        command = f"fastqc {trimmed_r1} {trimmed_r2}"
        run_command(command, os.path.join(output_base, folder_names[2]))
```

```
        # Step 3: Adapter removal
        nextera_trimmed_r1 = os.path.join(output_base, folder_names[3],
f"{sample}_nextera_trimmed_R1.fastq.gz")
        nextera_trimmed_r2 = os.path.join(output_base, folder_names[3],
f"{sample}_nextera_trimmed_R2.fastq.gz")
        command = f"fastp -i {trimmed_r1} -I {trimmed_r2} -o {nextera_trimmed_r1} -O
{nextera_trimmed_r2} --detect_adapter_for_pe -q 20 -l 36"
        run_command(command, os.path.join(output_base, folder_names[3]))

        # Step 4: FastQC after adapter removal
        command = f"fastqc {nextera_trimmed_r1} {nextera_trimmed_r2}"
        run_command(command, os.path.join(output_base, folder_names[2]))

        # Step 5: Alignment using BWA
        raw_sam = os.path.join(output_base, folder_names[4], f"{sample}_raw.sam")
        command = f"bwa mem -t 4 -k 32 -M {reference_genome} {nextera_trimmed_r1}
{nextera_trimmed_r2} > {raw_sam}"
        run_command(command, os.path.join(output_base, folder_names[4]))

        # Step 6: Convert SAM to BAM
        raw_bam = os.path.join(output_base, folder_names[5], f"{sample}_raw.bam")
        command = f"samtools view -bS {raw_sam} > {raw_bam}"
        run_command(command, os.path.join(output_base, folder_names[5]))

        # Step 7: Sorting BAM file
        sorted_bam = os.path.join(output_base, folder_names[6],
f"{sample}_sorted.bam")
        command = f"samtools sort {raw_bam} -o {sorted_bam}"
        run_command(command, os.path.join(output_base, folder_names[6]))

        # Step 8: Add Read Groups and Mark Duplicates
        rg_bam = os.path.join(output_base, folder_names[7], f"{sample}.rg.bam")
        command = f"{picard_cmd} AddOrReplaceReadGroups -I {sorted_bam} -O {rg_bam}
-ID {sample} -LB {sample} -PL ILLUMINA -SM {sample} -PU {sample}"
        run_command(command, os.path.join(output_base, folder_names[7]))

        marked_dup_bam = os.path.join(output_base, folder_names[7],
f"{sample}.marked_dup.bam")
        metrics_file = os.path.join(output_base, folder_names[7],
f"{sample}_metrics_duplicate.txt")
        command = f"{picard_cmd} MarkDuplicates -I {rg_bam} -O {marked_dup_bam} -M
{metrics_file}"
        run_command(command, os.path.join(output_base, folder_names[7]))

        # Step 9: Sorting and Indexing BAM file
        sorted1_bam = os.path.join(output_base, folder_names[8],
f"{sample}_sorted1.bam")
        command = f"{picard_cmd} SortSam -I {marked_dup_bam} -O {sorted1_bam} -SO
coordinate"
```

```
        run_command(command, os.path.join(output_base, folder_names[8]))
        run_command(f"samtools index {sorted1_bam}", os.path.join(output_base,
folder_names[8]))

        # Step 10: Variant Calling using GATK (VCF)
        vcf_raw = os.path.join(output_base, folder_names[9],
f"{sample}_GATK.raw.vcf")
        assembled_reads_bam = os.path.join(output_base, folder_names[10],
f"{sample}_assembled_reads.bam")
        assembled_regions_txt = os.path.join(output_base, folder_names[10],
f"{sample}_assembled_regions.txt")
        command = f"{gatk_cmd} HaplotypeCaller --output {vcf_raw} --input
{sorted1_bam} --reference {reference_genome} --ploidy 2 --bam-output
{assembled_reads_bam} --assembly-region-out {assembled_regions_txt}"
        run_command(command, os.path.join(output_base, folder_names[9]))

        # Step 11: GVCF Generation
        gvcf_file = os.path.join(output_base, folder_names[11], f"{sample}.g.vcf")
        command = f"{gatk_cmd} HaplotypeCaller -R {reference_genome} -I
{sorted1_bam} -O {gvcf_file} -ERC GVCF"
        run_command(command, os.path.join(output_base, folder_names[11]))

        # --------------------
        # CLEANUP: Delete intermediate files for this sample in folders 1-7
        cleanup_sample_files(sample, output_base, folder_names)

if __name__ == "__main__":
    main()
```

2. Extracting AD, DP, AF from vcf files

```
#!/usr/bin/env python3

import os
import pysam
import csv

# ==========================
# USER-DEFINED PATHS
# ==========================
JOINT_VCF = "/NGF_SK1ref.vcf"
OUTPUT_DIR = "/14_AD_AF_extracted"

os.makedirs(OUTPUT_DIR, exist_ok=True)

# ==========================
# SAMPLE GROUP DEFINITIONS
# ==========================
ancestor = "Ancestor2a"
```

```
groups = []
for i in range(1, 13):
    group = [f"A{i}", f"A{i+12}", f"A{i+24}", f"A{i+36}"]
    groups.append(group)

# ==========================
# OPEN VCF
# ==========================
vcf = pysam.VariantFile(JOINT_VCF)

# Sanity check
vcf_samples = list(vcf.header.samples)
if ancestor not in vcf_samples:
    raise ValueError("Ancestor2a not found in VCF header")

# ==========================
# HELPER FUNCTION
# ==========================
def extract_sample_metrics(record, sample):
    """
    Returns:
    DP, REF_AD, ALT1_AD, ALT2_AD, REF_AF, ALT1_AF, ALT2_AF
    """
    data = record.samples[sample]

    DP = data.get("DP", 0) or 0

    AD = data.get("AD", None)
    if AD is None:
        return DP, 0, 0, 0, 0.0, 0.0, 0.0

    REF_AD = AD[0] if len(AD) > 0 else 0
    ALT1_AD = AD[1] if len(AD) > 1 else 0
    ALT2_AD = AD[2] if len(AD) > 2 else 0

    total = REF_AD + ALT1_AD + ALT2_AD
    if total > 0:
        REF_AF = REF_AD / total
        ALT1_AF = ALT1_AD / total
        ALT2_AF = ALT2_AD / total
    else:
        REF_AF = ALT1_AF = ALT2_AF = 0.0

    return DP, REF_AD, ALT1_AD, ALT2_AD, REF_AF, ALT1_AF, ALT2_AF

# ==========================
# PROCESS EACH GROUP
# ==========================
for idx, group in enumerate(groups, start=1):
```

```
samples = [ancestor] + group
samples = [s for s in samples if s in vcf_samples]

out_tsv = os.path.join(
    OUTPUT_DIR,
    f"Joint_DP_AD_AF_set_{idx}.tsv"
)

with open(out_tsv, "w", newline="") as fh:
    writer = csv.writer(fh, delimiter="\t")

    # Header
    header = ["CHROM", "POS", "REF", "ALT"]
    for s in samples:
        header.extend([
            f"{s}_DP",
            f"{s}_REF_AD",
            f"{s}_ALT1_AD",
            f"{s}_ALT2_AD",
            f"{s}_REF_AF",
            f"{s}_ALT1_AF",
            f"{s}_ALT2_AF"
        ])
    writer.writerow(header)

    # Iterate over variants
    for record in vcf.fetch():
        alt_str = ",".join(record.alts) if record.alts else "."

        row = [
            record.chrom,
            record.pos,
            record.ref,
            alt_str
        ]

        for s in samples:
            DP, REF_AD, ALT1_AD, ALT2_AD, REF_AF, ALT1_AF, ALT2_AF = \
                extract_sample_metrics(record, s)

            row.extend([
                DP,
                REF_AD,
                ALT1_AD,
                ALT2_AD,
                f"{REF_AF:.4f}",
                f"{ALT1_AF:.4f}",
                f"{ALT2_AF:.4f}"
            ])
```

```
            writer.writerow(row)

    print(f"Written: {out_tsv}")

print("All files generated successfully.")
```

3. Removing variants that do not show changes in allele frequency across time

```
#!/usr/bin/env python3

import os
import pandas as pd
import numpy as np
import matplotlib.pyplot as plt

# ==========================
# PATHS
# ==========================
INPUT_DIR = "/14_AD_AF_extracted"
BASE_OUTPUT = "/15_DPless30_AFdiff_heatmap"

DP_FILTER_DIR = os.path.join(BASE_OUTPUT, "01_DP30_filtered")
AF_FILTER_DIR = os.path.join(BASE_OUTPUT, "02_REF_AF_different")
HEATMAP_DIR = os.path.join(BASE_OUTPUT, "03_heatmaps")

os.makedirs(DP_FILTER_DIR, exist_ok=True)
os.makedirs(AF_FILTER_DIR, exist_ok=True)
os.makedirs(HEATMAP_DIR, exist_ok=True)

# ==========================
# PARAMETERS
# ==========================
ANCESTOR = "Ancestor2a"
DP_THRESHOLD = 30
AF_DIFF_THRESHOLD = 0.2

# ==========================
# PROCESS EACH SET
# ==========================
tsv_files = sorted([f for f in os.listdir(INPUT_DIR) if f.endswith(".tsv")])

for tsv in tsv_files:
    print(f"\nProcessing {tsv}")
    df = pd.read_csv(os.path.join(INPUT_DIR, tsv), sep="\t")

    # --------------------------
    # Identify samples (ordered)
    # --------------------------
```

```
dp_cols = [c for c in df.columns if c.endswith("_DP")]
samples = [c.replace("_DP", "") for c in dp_cols]

# Ancestor first, then numeric order
samples = sorted(
    samples,
    key=lambda x: (x != ANCESTOR, int(x[1:]) if x != ANCESTOR else -1)
)

# -------------------------
# STEP 1: DP FILTER
# Keep variant only if ALL samples have DP ≥ 30
# -------------------------
dp_columns = [f"{s}_DP" for s in samples]
df_dp = df[df[dp_columns].min(axis=1) >= DP_THRESHOLD].copy()

dp_out = os.path.join(DP_FILTER_DIR, tsv)
df_dp.to_csv(dp_out, sep="\t", index=False)

# -------------------------
# STEP 2: REF_AF DIFFERENCE FILTER
# Keep variant if at least one evolved sample differs
# from ancestor by > 0.2
# -------------------------
ancestor_af = df_dp[f"{ANCESTOR}_REF_AF"]

keep_mask = np.zeros(len(df_dp), dtype=bool)

for s in samples:
    if s == ANCESTOR:
        continue
    diff = np.abs(df_dp[f"{s}_REF_AF"] - ancestor_af)
    keep_mask |= (diff > AF_DIFF_THRESHOLD)

df_af = df_dp[keep_mask].copy()

af_out = os.path.join(AF_FILTER_DIR, tsv)
df_af.to_csv(af_out, sep="\t", index=False)

# -------------------------
# STEP 3: HEATMAP
# -------------------------
if len(df_af) == 0:
    print("  No variants left after filtering — skipping heatmap")
    continue

af_matrix = df_af[[f"{s}_REF_AF" for s in samples]]

plt.figure(figsize=(1.2 * len(samples), 0.15 * len(af_matrix) + 3))
plt.imshow(af_matrix, aspect="auto", interpolation="nearest")
```

```
    plt.colorbar(label="REF_AF")

    plt.xticks(
        ticks=range(len(samples)),
        labels=samples,
        rotation=45,
        ha="right"
    )
    plt.yticks([])

    plt.title(tsv.replace(".tsv", ""))
    plt.tight_layout()

    heatmap_path = os.path.join(
        HEATMAP_DIR,
        tsv.replace(".tsv", "_REF_AF_heatmap.png")
    )
    plt.savefig(heatmap_path, dpi=300)
    plt.close()

    print(f"  Variants after DP filter: {len(df_dp)}")
    print(f"  Variants after AF filter: {len(df_af)}")
    print("  Heatmap saved")

print("\nAll sets processed successfully.")
```

4. Filter fixed variants

```
#!/usr/bin/env python3

import os
import pandas as pd
import numpy as np
import matplotlib.pyplot as plt

# ==========================
# PATHS
# ==========================
INPUT_DIR = "/15_DPless30_AFdiff_heatmap/02_REF_AF_different"

OUTPUT_BASE = "/16_fixed_variants"
OUTPUT_TSV_DIR = os.path.join(OUTPUT_BASE, "tsv")
OUTPUT_HEATMAP_DIR = os.path.join(OUTPUT_BASE, "heatmaps")

os.makedirs(OUTPUT_TSV_DIR, exist_ok=True)
os.makedirs(OUTPUT_HEATMAP_DIR, exist_ok=True)

# ==========================
```

```
# PARAMETERS
# ===========================
ANCESTOR = "Ancestor2a"

LOW = (0.0, 0.2)
MID = (0.4, 0.6)
HIGH = (0.8, 1.0)

# ===========================
# Helper function
# ===========================
def in_range(x, r):
    return (x >= r[0]) & (x <= r[1])

# ===========================
# PROCESS EACH FILE
# ===========================
tsv_files = sorted([f for f in os.listdir(INPUT_DIR) if f.endswith(".tsv")])

for tsv in tsv_files:
    print(f"\nProcessing {tsv}")
    df = pd.read_csv(os.path.join(INPUT_DIR, tsv), sep="\t")

    # --------------------------
    # Identify samples (ordered)
    # --------------------------
    dp_cols = [c for c in df.columns if c.endswith("_DP")]
    samples = [c.replace("_DP", "") for c in dp_cols]

    # Ancestor first, then numeric order
    samples = sorted(
        samples,
        key=lambda x: (x != ANCESTOR, int(x[1:]) if x != ANCESTOR else -1)
    )

    ancestor_af = df[f"{ANCESTOR}_REF_AF"]

    # --------------------------
    # Categorical REF_AF filter
    # --------------------------
    keep_mask = np.zeros(len(df), dtype=bool)

    for s in samples:
        if s == ANCESTOR:
            continue

        sample_af = df[f"{s}_REF_AF"]

        # Rule 1: Anc low -> sample mid/high
        rule1 = (
```

```
        in_range(ancestor_af, LOW) &
        (in_range(sample_af, MID) | in_range(sample_af, HIGH))
    )

    # Rule 2: Anc high -> sample low/mid
    rule2 = (
        in_range(ancestor_af, HIGH) &
        (in_range(sample_af, LOW) | in_range(sample_af, MID))
    )

    # Rule 3: Anc mid -> sample low/high
    rule3 = (
        in_range(ancestor_af, MID) &
        (in_range(sample_af, LOW) | in_range(sample_af, HIGH))
    )

    keep_mask |= (rule1 | rule2 | rule3)

df_out = df[keep_mask].copy()

# -------------------------
# Write filtered TSV
# -------------------------
out_tsv = os.path.join(OUTPUT_TSV_DIR, tsv)
df_out.to_csv(out_tsv, sep="\t", index=False)

print(f"  Variants kept: {len(df_out)}")

# -------------------------
# Heatmap (REF_AF)
# -------------------------
if len(df_out) == 0:
    print("  No variants left — skipping heatmap")
    continue

af_matrix = df_out[[f"{s}_REF_AF" for s in samples]]

plt.figure(figsize=(1.2 * len(samples), 0.15 * len(af_matrix) + 3))
plt.imshow(af_matrix, aspect="auto", interpolation="nearest")
plt.colorbar(label="REF_AF")

plt.xticks(
    ticks=range(len(samples)),
    labels=samples,
    rotation=45,
    ha="right"
)
plt.yticks([])

plt.title(tsv.replace(".tsv", ""))
```

```
    plt.tight_layout()

    heatmap_path = os.path.join(
        OUTPUT_HEATMAP_DIR,
        tsv.replace(".tsv", "_REF_AF_category_heatmap.png")
    )

    plt.savefig(heatmap_path, dpi=300)
    plt.close()

    print("  Heatmap saved")

print("\nAll categorical filtering and heatmaps complete.")
```

5. Annotate variants

```
#!/usr/bin/env python3

import os
import pandas as pd
import subprocess

# ============================================================
# USER INPUTS
# ============================================================

INPUT_CSV      = "/14_SNP_analysis/3_fixed_variants/tsv/gal6_A.csv"
JOINT_VCF      = "/12_Joint_variantcalling/joint_genotyped_all_samples.vcf"
AF_TSV         = "/14_SNP_analysis/3_fixed_variants/tsv/A12_A24.tsv"

WORK_FOLDER    = "/14_SNP_analysis/4_fixed_annotated"
FINAL_OUTPUT   = "/14_SNP_analysis/5_fixed_annotated_merged"

os.makedirs(WORK_FOLDER, exist_ok=True)
os.makedirs(FINAL_OUTPUT, exist_ok=True)

# intermediate files
FILTERED_VCF   = os.path.join(WORK_FOLDER, "gal6_A_filtered.vcf")
ANNOTATED_VCF  = os.path.join(WORK_FOLDER, "gal6_A_annotated.vcf")
ANNOTATED_TSV  = os.path.join(WORK_FOLDER, "gal6_A_annotated.tsv")
FILTERED_ANN   = os.path.join(WORK_FOLDER, "gal6_A_filtered_annotation.tsv")

FINAL_TSV      = os.path.join(FINAL_OUTPUT, "gal6_A_annotation_AF_merged.tsv")


# ============================================================
# STEP 1: CSV → extract matching variants from joint VCF
# ============================================================
```

```
def csv_to_vcf(csv_file, joint_vcf, output_vcf):

    df = pd.read_csv(csv_file, sep="\t")

    variant_set = set(zip(df["CHROM"], df["POS"], df["REF"], df["ALT"]))

    filtered_lines = []

    with open(joint_vcf) as vcf:
        for line in vcf:

            if line.startswith("#"):
                filtered_lines.append(line)
                continue

            chrom, pos, _, ref, alt = line.strip().split("\t")[:5]

            if (chrom, int(pos), ref, alt) in variant_set:
                filtered_lines.append(line)

    with open(output_vcf, "w") as out:
        out.writelines(filtered_lines)


# ============================================================
# STEP 2: run snpEff
# ============================================================

def run_snpeff(input_vcf, output_vcf):

    print("Running snpEff annotation")

    cmd = f"java -jar snpEff.jar ann SK1_custom {input_vcf} > {output_vcf}"

    subprocess.run(cmd, shell=True, check=True)


# ============================================================
# STEP 3: annotated VCF → TSV
# ============================================================

def vcf_to_tsv(input_vcf, output_tsv):

    awk_cmd = r"""awk 'BEGIN {OFS="\t"; print
"CHROM","POS","REF","ALT","QUAL","Allele","Consequence","Impact","GeneName","GeneID"
,"FeatureType","FeatureID","TranscriptBioType"} {n = split($6, annots, ",");
for(i=1;i<=n;i++){split(annots[i], a, "|"); print
$1,$2,$3,$4,$5,a[1],a[2],a[3],a[4],a[5],a[6],a[7],a[8]}}'"""

    cmd = f"bcftools query -f '%CHROM\t%POS\t%REF\t%ALT\t%QUAL\t%INFO/ANN\n'
```

```
{input_vcf} | {awk_cmd} > {output_tsv}"

    subprocess.run(cmd, shell=True, check=True)


# ==========================================================
# STEP 4: keep all annotations ONLY if first annotation is
# upstream_gene_variant or downstream_gene_variant
# ==========================================================

def keep_first_annotation(input_tsv, output_tsv):

    df = pd.read_csv(input_tsv, sep="\t")

    def filter_annotations(group):

        first_cons = str(group.iloc[0]["Consequence"])

        if ("upstream_gene_variant" in first_cons) or ("downstream_gene_variant" in 
first_cons):
            return group

        # otherwise keep only first annotation
        return group.iloc[[0]]

    filtered_df = (
        df.groupby(["CHROM","POS","REF","ALT"], group_keys=False)
        .apply(filter_annotations)
    )

    filtered_df.to_csv(output_tsv, sep="\t", index=False)

# ==========================================================
# STEP 5: merge with allele frequency table
# ==========================================================

def merge_AF(annotation_tsv, af_tsv, output_tsv):

    ann_df = pd.read_csv(annotation_tsv, sep="\t", dtype=str)
    af_df  = pd.read_csv(af_tsv, sep="\t", dtype=str)

    # detect ALT1_AF columns automatically
    af_cols = [c for c in af_df.columns if c.endswith("_AF_ALT1")]

    merge_cols = ["CHROM","POS","REF","ALT"]

    af_subset = af_df[merge_cols + af_cols]

    merged = ann_df.merge(
        af_subset,
```

```
        on=merge_cols,
        how="left"
    )

    merged.to_csv(output_tsv, sep="\t", index=False)


# ============================================================
# PIPELINE
# ============================================================

print("\nStep 1: Extract variants from joint VCF")
csv_to_vcf(INPUT_CSV, JOINT_VCF, FILTERED_VCF)

print("\nStep 2: Annotate variants")
run_snpeff(FILTERED_VCF, ANNOTATED_VCF)

print("\nStep 3: Convert annotated VCF to TSV")
vcf_to_tsv(ANNOTATED_VCF, ANNOTATED_TSV)

print("\nStep 4: Keep first annotation")
keep_first_annotation(ANNOTATED_TSV, FILTERED_ANN)

print("\nStep 5: Merge allele frequencies")
merge_AF(FILTERED_ANN, AF_TSV, FINAL_TSV)

print("\n✔ Pipeline finished")
print("Final file:", FINAL_TSV)
```

6. CNV analysis - Coverage based pipeline with HMM model

```
import os
import subprocess
import pandas as pd
import numpy as np
from hmmlearn.hmm import GaussianHMM

# === SETTINGS ===
bam_dir = "/8_Sorting_And_Indexing_A"
output_dir = "/25_Final_CNV_analysis/A_Read_depth"
window_size = 500
ancestor_id = "A0_sorted1"

os.makedirs(f"{output_dir}/1_depth", exist_ok=True)
os.makedirs(f"{output_dir}/2_cnv_calls", exist_ok=True)

# === STEP 1: CALCULATE DEPTH WITH SAMTOOLS ===
def generate_depth_file(bam_path, output_path):
    cmd = f"samtools depth {bam_path} > {output_path}"
```

```
    subprocess.run(cmd, shell=True, check=True)

# === STEP 2: BIN DEPTH INTO 500bp WINDOWS ===
def bin_depth_file(depth_file, window_size):
    df = pd.read_csv(depth_file, sep="\t", names=["chrom", "pos", "depth"])
    df["window"] = (df["pos"] // window_size) * window_size
    binned = df.groupby(["chrom", "window"])["depth"].median().reset_index()
    binned["end"] = binned["window"] + window_size
    return binned[["chrom", "window", "end", "depth"]]

# === STEP 3: NORMALIZE DEPTH ===
def compute_relative_depth(df):
    median = df["depth"].median()
    df["relative_depth"] = df["depth"] / median
    return df

# === STEP 4: STANDARDIZE USING ANCESTOR RELATIVE DEPTH ===
def standardize_by_ancestor(sample_df, ancestor_df):
    merged = sample_df.merge(ancestor_df, on=["chrom", "window", "end"],
suffixes=("", "_ancestor"))
    merged = merged[merged["relative_depth_ancestor"] >= 0.25].copy()
    merged["standardized"] = merged["relative_depth"] /
merged["relative_depth_ancestor"]
    return merged

# === STEP 5: HMM CNV DETECTION ===
def run_hmm(df, sample_name, out_path):
    states = [0, 0.5, 1, 1.5, 2, 3, 4]
    X = df["standardized"].values.reshape(-1, 1)
    baseline_var = np.var(X)
    means = np.array([[s] for s in states])
    covars = np.array([[baseline_var * 0.5 if s == 0 else baseline_var * s] for s in
states])

    model = GaussianHMM(n_components=len(states), covariance_type="diag",
init_params="")
    model.means_ = means
    model.covars_ = covars
    model.startprob_ = np.array([0.01] * len(states))
    model.startprob_[states.index(1)] = 0.94
    transmat = np.full((len(states), len(states)), 0.0001)
    np.fill_diagonal(transmat, 0.9994)
    model.transmat_ = transmat

    df["CNV_state"] = [states[s] for s in model.predict(X)]
    df.to_csv(out_path, sep="\t", index=False)

# === MAIN WORKFLOW ===
binned_depths = {}
```

```
# Process ancestor BAM first
bam_files = [f for f in os.listdir(bam_dir) if f.endswith(".bam")]
ancestor_bam = f"{ancestor_id}.bam"
if ancestor_bam not in bam_files:
    raise ValueError(f"Ancestor BAM file {ancestor_bam} not found in {bam_dir}")

bam_files.remove(ancestor_bam)
bam_files.insert(0, ancestor_bam)

for bam in bam_files:
    sample = bam.replace(".bam", "")
    bam_path = os.path.join(bam_dir, bam)
    depth_path = f"{output_dir}/1_depth/{sample}.depth"

    if os.path.exists(depth_path):
        print(f"Depth file already exists for {sample}, skipping samtools depth.")
    else:
        print(f"Running samtools depth for {sample}...")
        generate_depth_file(bam_path, depth_path)

    binned = bin_depth_file(depth_path, window_size)
    binned = compute_relative_depth(binned)
    binned_depths[sample] = binned

# Get ancestor baseline
ancestor_df = binned_depths[ancestor_id].copy()
ancestor_df = ancestor_df[["chrom", "window", "end", "relative_depth"]]
ancestor_df = ancestor_df.rename(columns={"relative_depth":
"relative_depth_ancestor"})

# Process all samples except ancestor
for sample, df in binned_depths.items():
    if sample == ancestor_id:
        continue
    standardized_df = standardize_by_ancestor(df, ancestor_df)
    out_path = f"{output_dir}/2_cnv_calls/{sample}_cnv_calls.tsv"
    run_hmm(standardized_df, sample, out_path)
    print(f"CNV calls saved for {sample} → {out_path}")
```

7. Merge CNV calls

```
#!/usr/bin/env python3

import os
import pandas as pd

# ============================================================
# USER INPUT
# ============================================================
```

```
BASE_INPUT_DIR  = "/25_Final_CNV_analysis/A_Read_depth/2_cnv_calls"
BASE_OUTPUT_DIR = "/25_Final_CNV_analysis/A_Read_depth/3_merged_calls"

os.makedirs(BASE_OUTPUT_DIR, exist_ok=True)

# ============================================================
# SAMPLE GROUP DEFINITIONS
# ============================================================

groups = {
    "glu_300":  range(1, 7),
    "gal_300":  range(7, 13),
    "glu_600":  range(13, 19),
    "gal_600":  range(19, 25),
    "glu_900":  range(25, 31),
    "gal_900":  range(31, 37),
    "glu_1200": range(37, 43),
    "gal_1200": range(43, 49),
}

# ============================================================
# LOOP OVER GROUPS
# ============================================================

for group_name, sample_range in groups.items():

    print(f"\nProcessing {group_name}...")

    merged_df = None

    for i in sample_range:
        sample = f"A{i}"
        filepath = os.path.join(BASE_INPUT_DIR, f"{sample}_sorted1_cnv_calls.tsv")

        if not os.path.exists(filepath):
            print(f"⚠ Missing file: {filepath}")
            continue

        df = pd.read_csv(
            filepath,
            sep="\t",
            usecols=["chrom", "window", "end", "CNV_state"]
        )

        df = df.rename(columns={"CNV_state": sample})

        if merged_df is None:
            merged_df = df
        else:
```

```
            merged_df = pd.merge(
                merged_df,
                df,
                on=["chrom", "window", "end"],
                how="outer"
            )

    if merged_df is not None:
        merged_df = merged_df.sort_values(by=["chrom", "window"])

        output_path = os.path.join(
            BASE_OUTPUT_DIR,
            f"merged_CNV_state_{group_name}.tsv"
        )

        merged_df.to_csv(output_path, sep="\t", index=False)

        print(f" Saved: {output_path}")
    else:
        print(f" No files merged for {group_name}")

print("\n All groups processed.")
```

8. Plot average CNV change across replicates

```
import pandas as pd
import matplotlib.pyplot as plt
import matplotlib.cm as cm
import numpy as np

# === Input file paths ===
file_300 =
"/25_Final_CNV_analysis/A_Read_depth/3_merged_calls/merged_CNV_state_gal_300.tsv"
file_600 =
"/25_Final_CNV_analysis/A_Read_depth/3_merged_calls/merged_CNV_state_gal_600.tsv"
file_900 =
"/25_Final_CNV_analysis/A_Read_depth/3_merged_calls/merged_CNV_state_gal_900.tsv"
file_1200 =
"/25_Final_CNV_analysis/A_Read_depth/3_merged_calls/merged_CNV_state_gal_1200.tsv"

# === Chromosome label mapping (Roman numerals) ===
chrom_label_map = {
    'I': 'I', 'II': 'II', 'III': 'III', 'IV': 'IV',
    'V': 'V', 'VI': 'VI', 'VII': 'VII', 'VIII': 'VIII',
    'IX': 'IX', 'X': 'X', 'XI': 'XI', 'XII': 'XII',
    'XIII': 'XIII', 'XIV': 'XIV', 'XV': 'XV', 'XVI': 'XVI'
}
```

```
# === Preprocess file ===
def process_file(filepath):
    df = pd.read_csv(filepath, sep="\t")

    sample_cols = df.columns[-6:]
    df['avg'] = df[sample_cols].mean(axis=1)

    # FIX: chromosomes are chrI, chrII → strip "chr"
    df['chrom_id'] = df['chrom'].astype(str).str.replace("^chr", "", regex=True)
    df['chrom_label'] = df['chrom_id']

    return df[['chrom_id', 'chrom_label', 'window', 'avg']]

# === Load all datasets ===
df_300 = process_file(file_300)
df_600 = process_file(file_600)
df_900 = process_file(file_900)
df_1200 = process_file(file_1200)

# === Define chromosome order and color mapping ===
chrom_order = list(chrom_label_map.keys())
chrom_to_color = {chrom: cm.tab20(i % 20) for i, chrom in enumerate(chrom_order)}


# === Assign consistent x_pos and chromosome boundaries ===
def assign_global_xpos(df_base):
    xpos_map = {}
    boundaries = {}
    current_offset = 0

    for i, chrom in enumerate(chrom_order):
        chrom_df = df_base[df_base['chrom_id'] == chrom].sort_values('window')
        chrom_len = chrom_df['window'].max()

        boundaries[chrom] = (current_offset, current_offset + chrom_len)

        for _, row in chrom_df.iterrows():
            xpos_map[(chrom, row['window'])] = current_offset + row['window']

        if i < len(chrom_order) - 1:
            current_offset += chrom_len + 50000
        else:
            current_offset += chrom_len

    return xpos_map, boundaries

xpos_map, chrom_boundaries = assign_global_xpos(df_300)

# === Apply consistent x_pos to all datasets ===
def apply_xpos(df, xpos_map):
```

```
    df = df.copy()
    df['x_pos'] = df.apply(lambda row: xpos_map.get((row['chrom_id'],
row['window']), np.nan), axis=1)
    df['color'] = df['chrom_id'].map(chrom_to_color)
    return df

df_300 = apply_xpos(df_300, xpos_map)
df_600 = apply_xpos(df_600, xpos_map)
df_900 = apply_xpos(df_900, xpos_map)
df_1200 = apply_xpos(df_1200, xpos_map)

# === Fixed y-axis limits ===
ymin, ymax = 0.4, 1.8

# === Create subplots ===
fig, axes = plt.subplots(4, 1, figsize=(18, 12), sharex=True)
fig.text(0.04, 0.5, 'CNV state', va='center', rotation='vertical', fontsize=14,
fontname='Arial')

# === Timepoint mapping ===
timepoints = {
    '300 generations': df_300,
    '600 generations': df_600,
    '900 generations': df_900,
    '1200 generations': df_1200
}

# === Plot each panel ===
for ax, (label, df) in zip(axes, timepoints.items()):
    ax.scatter(df['x_pos'], df['avg'], color=df['color'], s=8)
    ax.set_title(label, fontsize=14)
    ax.set_ylim(ymin, ymax)
    ax.tick_params(axis='y', labelsize=12)
    ax.set_yticks(np.arange(ymin, ymax + 0.01, 0.2))

    for chrom in chrom_order[:-1]:
        _, end = chrom_boundaries[chrom]
        ax.axvline(x=end + 25000, color='black', linestyle='-', linewidth=0.8)

# === Set chromosome labels on bottom axis ===
xticks = []
xticklabels = []
for chrom in chrom_order:
    start, end = chrom_boundaries[chrom]
    xticks.append((start + end) // 2)
    xticklabels.append(chrom_label_map[chrom])

axes[-1].set_xticks(xticks)
axes[-1].set_xticklabels(xticklabels, fontsize=12, fontname='Arial')
axes[-1].set_xlabel("Chromosome", fontsize=14, fontname='Arial')
```

```
start_x = chrom_boundaries[chrom_order[0]][0]
end_x = chrom_boundaries[chrom_order[-1]][1]
for ax in axes:
    ax.set_xlim(start_x, end_x)
    for label in ax.get_yticklabels():
        label.set_fontname('Arial')


# === Save and show ===
plt.tight_layout(rect=[0.06, 0.03, 1, 0.97])
plt.savefig(

"/25_Final_CNV_analysis/A_Read_depth/4_avg_CNV_plots/galactose_CNV_state_plot.png",
    dpi=300,
    bbox_inches='tight'
)
plt.show()
```

9. Separate deletions and duplications

```
import pandas as pd
from pathlib import Path

# User configurable paths
input_folder = Path("/25_Final_CNV_analysis/A_Read_depth/2_cnv_calls")
output_folder = Path("/25_Final_CNV_analysis/A_Read_depth/5_CNV_separated")
deletion_folder = output_folder / "deletion"
duplication_folder = output_folder / "duplication"

# Create output subfolders if not exist
deletion_folder.mkdir(parents=True, exist_ok=True)
duplication_folder.mkdir(parents=True, exist_ok=True)

for i in range(1, 49):
    filename = f"A{i}_sorted1_cnv_calls.tsv"
    input_file = input_folder / filename

    if not input_file.exists():
        print(f"Warning: {filename} not found, skipping.")
        continue

    # Read TSV file
    df = pd.read_csv(input_file, sep='\t')

    # Filter rows based on CNV_state
    deletion_df = df[df['CNV_state'] == 0.5]
    duplication_df = df[df['CNV_state'] == 1.5]
```

```
    # Write to respective output files
    deletion_output_file = deletion_folder / filename
    duplication_output_file = duplication_folder / filename

    deletion_df.to_csv(deletion_output_file, sep='\t', index=False)
    duplication_df.to_csv(duplication_output_file, sep='\t', index=False)

    print(f"Processed {filename}: {len(deletion_df)} deletions,
{len(duplication_df)} duplications")

print("All files processed.")
```

10. Annotate deletions and duplications

```
#!/usr/bin/env python3

import os
import subprocess
import pandas as pd
from pathlib import Path

# === USER CONFIG ===
input_folder =
Path("/25_Final_CNV_analysis/A_Read_depth/5_CNV_separated/duplication")
gff_file = Path("/SK1_reference_genome/SK1_lifted_annotation.gff3")
output_base_dir =
Path("/25_Final_CNV_analysis/A_Read_depth/6_CNV_separated_annotated/duplication_anno
tated")

# Create output directories
output_base_dir.mkdir(parents=True, exist_ok=True)
unique_genes_dir = output_base_dir / "unique_genes_dup"
unique_genes_dir.mkdir(parents=True, exist_ok=True)

# ============================================================
# STEP 0: Extract genes.bed (only once)
# ============================================================

genes_bed = output_base_dir / "genes.bed"

if not genes_bed.exists():
    print(" Extracting genes from GFF file...")
    with open(gff_file) as fin, open(genes_bed, "w") as fout:
        for line in fin:
            if line.startswith("#"):
                continue
            parts = line.strip().split('\t')
```

```
            if len(parts) < 9 or parts[2] != "gene":
                continue
            chrom, _, _, start, end, _, strand, _, attr = parts
            attrs = {k: v for k, v in (item.split('=') for item in attr.split(';')
if '=' in item)}
            gene_id = attrs.get('ID', '.')
            gene_name = attrs.get('Name', '.')

fout.write(f"{chrom}\t{int(start)-1}\t{end}\t{gene_id}\t{gene_name}\t{strand}\n")

# ==========================================================
# LOOP OVER A1–A48
# ==========================================================

for i in range(1, 49):

    sample_name = f"A{i}"
    input_filename = f"{sample_name}_sorted1_cnv_calls.tsv"
    window_tsv = input_folder / input_filename

    if not window_tsv.exists():
        print(f" Missing file, skipping: {window_tsv}")
        continue

    print(f"\n Processing {sample_name} ...")

    # Define output files
    windows_bed = output_base_dir / f"{sample_name}_windows.bed"
    merged_bed = output_base_dir / f"{sample_name}_merged_blocks.bed"
    intersect_file = output_base_dir / f"{sample_name}_full_gene_overlap.tsv"
    parsed_output = output_base_dir / f"{sample_name}_parsed_annotation.tsv"
    filtered_output = output_base_dir /
f"{sample_name}_parsed_annotation_filtered.tsv"
    unique_genes_txt = unique_genes_dir / f"{sample_name}_unique_gene_names_dup.txt"

    # ==========================================================
    # STEP 1: Convert TSV → BED
    # ==========================================================

    with open(window_tsv) as fin, open(windows_bed, "w") as fout:
        next(fin)  # skip header
        for line in fin:
            chrom, start, end = line.strip().split('\t')[:3]
            if chrom == "BK006947.2":
                chrom = "BK006947.3"
            start_bed = max(0, int(start) - 1)
            fout.write(f"{chrom}\t{start_bed}\t{end}\n")

    # ==========================================================
    # STEP 2: Merge adjacent windows
```

```
# ===========================================================

subprocess.run([
    "bedtools", "merge",
    "-i", str(windows_bed)
], stdout=open(merged_bed, "w"), check=True)

# ===========================================================
# STEP 3: Intersect genes fully contained in CNV blocks
# ===========================================================

subprocess.run([
    "bedtools", "intersect",
    "-a", str(genes_bed),
    "-b", str(merged_bed),
    "-wa", "-u",
    "-f", "1.0"
], stdout=open(intersect_file, "w"), check=True)

# ===========================================================
# STEP 4: Parse intersect results
# ===========================================================

if os.path.getsize(intersect_file) == 0:
    print(f"⚠ No fully duplicated genes found for {sample_name}")
    continue

cols = [
    "gene_chrom", "gene_start", "gene_end",
    "gene_id", "gene_name", "strand"
]

df = pd.read_csv(intersect_file, sep='\t', header=None, names=cols)
df.drop_duplicates(inplace=True)
df.to_csv(parsed_output, sep='\t', index=False)

# ===========================================================
# STEP 5: Filter gene_id != gene_name
# ===========================================================

df["core_gene_id"] = df["gene_id"].str.replace(
    r"^(gene-|transcript-|.*?:)?", "", regex=True
)

filtered_df = df[df["core_gene_id"] != df["gene_name"]].copy()
filtered_df.drop(columns=["core_gene_id"], inplace=True)
filtered_df.to_csv(filtered_output, sep='\t', index=False)

# ===========================================================
# STEP 6: Extract unique gene names
```

```
    # ==========================================================

    unique_genes = sorted(filtered_df["gene_name"].unique())

    with open(unique_genes_txt, "w") as fout:
        fout.write("\n".join(unique_genes))

    print(f" Completed {sample_name}")
    print(f"  Fully duplicated genes: {len(unique_genes)}")

print("\n All samples processed successfully.")
```

11. Extract gene count, gene names in each replicate

```
#!/usr/bin/env python3

import os
import pandas as pd

folder_del =
"/25_Final_CNV_analysis/A_Read_depth/6_CNV_separated_annotated/deletion_annotated/un
ique_genes_del"
folder_dup =
"/25_Final_CNV_analysis/A_Read_depth/6_CNV_separated_annotated/duplication_annotated
/unique_genes_dup"

total_genes = 6281

samples = []
count_del = []
count_dup = []

missing_del = []
missing_dup = []

for i in range(1, 49):

    sample = f"A{i}"

    del_file = os.path.join(folder_del, f"{sample}_unique_gene_names_del.txt")
    dup_file = os.path.join(folder_dup, f"{sample}_unique_gene_names_dup.txt")

    # deletion count
    if os.path.exists(del_file):
        with open(del_file) as f:
            del_count = sum(1 for _ in f)
    else:
        del_count = 0
```

```
        missing_del.append(sample)

    # duplication count
    if os.path.exists(dup_file):
        with open(dup_file) as f:
            dup_count = sum(1 for _ in f)
    else:
        dup_count = 0
        missing_dup.append(sample)

    samples.append(sample)
    count_del.append(del_count)
    count_dup.append(dup_count)

# Create dataframe
df = pd.DataFrame({
    "sample": samples,
    "count_del": count_del,
    "count_dup": count_dup
})

# Calculate percentages
df["del_ratio"] = (df["count_del"] / total_genes) * 100
df["dup_ratio"] = (df["count_dup"] / total_genes) * 100

# Save TSV
output_file =
"/25_Final_CNV_analysis/A_Read_depth/6_CNV_separated_annotated/CNV_gene_counts_summa
ry.tsv"
df.to_csv(output_file, sep="\t", index=False)

print(f"Summary file saved as {output_file}")

# Only print missing files to terminal
if missing_del:
    print("\nDeletion file missing for:", ", ".join(missing_del))

if missing_dup:
    print("\nDuplication file missing for:", ", ".join(missing_dup))


#!/usr/bin/env python3

import os
import pandas as pd

# === Input folders ===
folder_del =
"/25_Final_CNV_analysis/A_Read_depth/6_CNV_separated_annotated/deletion_annotated/un
ique_genes_del"
```

```
folder_dup =
"/25_Final_CNV_analysis/A_Read_depth/6_CNV_separated_annotated/duplication_annotated
/unique_genes_dup"

# === Output files ===
output_del =
"/25_Final_CNV_analysis/A_Read_depth/6_CNV_separated_annotated/RD_deletion_gene_list
.tsv"
output_dup =
"/25_Final_CNV_analysis/A_Read_depth/6_CNV_separated_annotated/RD_duplication_gene_l
ist.tsv"


def compile_gene_lists(folder, suffix=""):

    sample_dict = {}
    max_len = 0
    missing_samples = []

    for i in range(1, 49):

        sample_name = f"A{i}"
        file_path = os.path.join(folder,
f"{sample_name}_unique_gene_names{suffix}.txt")

        if os.path.exists(file_path):
            with open(file_path) as f:
                genes = [line.strip() for line in f if line.strip()]
        else:
            genes = []   # empty column if file missing
            missing_samples.append(sample_name)

        sample_dict[sample_name] = genes
        max_len = max(max_len, len(genes))

    # Pad lists so all columns have equal length
    for sample in sample_dict:
        sample_dict[sample] += [""] * (max_len - len(sample_dict[sample]))

    df = pd.DataFrame(sample_dict)

    if missing_samples:
        print(f"Missing files for: {', '.join(missing_samples)}")

    return df


# === Compile deletion and duplication gene tables ===
df_del = compile_gene_lists(folder_del, suffix="_del")
df_dup = compile_gene_lists(folder_dup, suffix="_dup")
```

```
# === Save files ===
df_del.to_csv(output_del, sep="\t", index=False)
df_dup.to_csv(output_dup, sep="\t", index=False)

print(f"Deletion gene list saved: {output_del}")
print(f"Duplication gene list saved: {output_dup}")
```

12. Extract list of genes undergoing deleion / duplication commonly across replicates

```
#!/usr/bin/env Rscript

library(readxl)

# =========================
# INPUT FILES
# =========================
del_file <- "/NGF/ms_figures/RD_deletion_gene_list.xlsx"
dup_file <- "/NGF/ms_figures/RD_duplication_gene_list.xlsx"

# =========================
# OUTPUT DIRECTORY
# =========================
out_dir <- /NGF/ms_figures/CNV_unique_gene_lists"

if (!dir.exists(out_dir)) {
  dir.create(out_dir, recursive = TRUE)
}

# =========================
# FUNCTION
# =========================
process_file <- function(file, prefix, out_dir) {

  cat("\nProcessing:", file, "\n")

  raw <- read_excel(file, col_names = FALSE)

  # Assign A1–A48
  colnames(raw) <- as.character(unlist(raw[2, ]))

  # Remove header rows
  df <- raw[-c(1,2,3), ]

  # Define groups
  groups <- list(
    "300_glu"  = paste0("A", 1:6),
    "300_gal"  = paste0("A", 7:12),
```

```
    "600_glu"  = paste0("A", 13:18),
    "600_gal"  = paste0("A", 19:24),

    "900_glu"  = paste0("A", 25:30),
    "900_gal"  = paste0("A", 31:36),

    "1200_glu" = paste0("A", 37:42),
    "1200_gal" = paste0("A", 43:48)
  )

  for (name in names(groups)) {

    cols <- groups[[name]]

    sub_df <- df[, cols]

    # Count gene frequency across 6 replicates
    gene_counts <- table(unlist(sub_df))

    # Remove NA if present
    gene_counts <- gene_counts[names(gene_counts) != "NA"]

    # Keep genes present in ≥4 replicates
    genes <- names(gene_counts[gene_counts >= 4])

    genes <- sort(genes)

    # Output file
    out_file <- file.path(out_dir, paste0(prefix, "_", name, ".txt"))
    writeLines(genes, out_file)

    cat("Written:", out_file, "|", length(genes), "genes\n")
  }
}

# ========================
# RUN
# ========================
process_file(del_file, "Deletion", out_dir)
process_file(dup_file, "Duplication", out_dir)

cat("\nAll files generated successfully in:", out_dir, "\n")
```

13. Pathway Enrichment Analysis

```
#!/usr/bin/env Rscript

# ============================================
```

```
# KEGG Enrichment with Bootstrapped Binning
# ============================================

library(clusterProfiler)
library(org.Sc.sgd.db)
library(enrichplot)
library(ggplot2)
library(dplyr)
library(stringr)

# ---- USER INPUT ----
input_file  <- "/NGF/ms_figures/CNV_unique_gene_lists/Deletion_600_glu.txt"
output_dir  <- "/NGF/ms_figures/FigS5/glu_600_del"
bin_size    <- 100   # genes per bin
n_repeats   <- 3     # number of bootstrap replicates

# ---- SETUP ----
dir.create(output_dir, showWarnings = FALSE)

# ---- 1. Load gene list ----
gene_list <- read.table(input_file, header = FALSE, stringsAsFactors = FALSE)[[1]]
gene_list <- unique(gene_list[gene_list != ""])  # remove blanks & duplicates
total_genes <- length(gene_list)
cat("Total genes:", total_genes, "\n")

# ---- 2. Bootstrap repeats ----
for (rep in 1:n_repeats) {
    cat("\n--- Bootstrap replicate", rep, "---\n")

    rep_dir <- file.path(output_dir, paste0("rep", rep))
    dir.create(rep_dir, showWarnings = FALSE)

    # Shuffle genes
    set.seed(100 + rep)  # different seed each time
    shuffled <- sample(gene_list)

    # Split into bins of bin_size
    num_bins <- ceiling(total_genes / bin_size)
    gene_bins <- split(shuffled, ceiling(seq_along(shuffled) / bin_size))
    cat("Bin sizes for replicate", rep, ":", sapply(gene_bins, length), "\n")

    # ---- 3. Process each bin ----
    for (i in seq_along(gene_bins)) {
        bin_genes <- gene_bins[[i]]
        cat("Processing replicate", rep, "bin", i, "with", length(bin_genes),
"genes...\n")

        # ---- Map GENENAME -> ENTREZ ID ----
        mapped <- bitr(bin_genes,
                       fromType = "GENENAME",
```

```
}

                toType   = "ENTREZID",
                OrgDb    = org.Sc.sgd.db)
        entrez_genes <- unique(mapped$ENTREZID)

        if (length(entrez_genes) == 0) {
            cat("No valid ENTREZ IDs for bin", i, "in replicate", rep, "\n")
            next
        }

        # ---- Run KEGG enrichment ----
        kegg_res <- enrichKEGG(
            gene         = entrez_genes,
            organism     = "sce",
            keyType      = "ncbi-geneid",
            pvalueCutoff = 1
        )

        if (is.null(kegg_res) || nrow(as.data.frame(kegg_res)) == 0) {
            cat("No enrichment found for bin", i, "in replicate", rep, "\n")
            next
        }

        # ---- Shorten descriptions ----
        kegg_res@result$Description <- str_replace(
            kegg_res@result$Description,
            " - Saccharomyces cerevisiae \\(budding yeast\\)", ""
        )

        # ---- Map Entrez IDs in results to GENENAME ----
        all_entrez <- unique(unlist(strsplit(kegg_res@result$geneID, "/")))
        entrez2gene <- AnnotationDbi::select(
            org.Sc.sgd.db,
            keys = all_entrez,
            columns = c("GENENAME"),
            keytype = "ENTREZID"
        )
        lookup <- setNames(entrez2gene$GENENAME, entrez2gene$ENTREZID)
        kegg_res@result$geneID <- sapply(kegg_res@result$geneID, function(ids) {
            genes <- unlist(strsplit(ids, "/"))
            paste(lookup[genes], collapse = "/")
        })

        # ---- Save CSV ----
        csv_file <- file.path(rep_dir, paste0("KEGG_bin_", i, ".csv"))
        write.csv(as.data.frame(kegg_res), csv_file, row.names = FALSE)

        cat("Results saved:", csv_file, "\n")
    }
}
```

```
cat("\nAll bootstrap replicates complete.\n")


#!/usr/bin/env Rscript

# =========================================
# Combine KEGG Enrichment Results across Bins (Multiple Reps)
# =========================================

# ---- Load libraries ----
library(dplyr)
library(purrr)
library(metap)   # for Fisher’s method
library(stringr)

# ---- User Input ----
# Parent folder containing rep1, rep2, ..., repN
input_parent <- "/NGF/ms_figures/FigS5/glu_600_del"
output_parent <- file.path(input_parent, "g_del_600_combined")

# Create output folder if missing
if (!dir.exists(output_parent)) dir.create(output_parent, recursive = TRUE)

# ---- 1. Fisher’s method ----
combine_fisher <- function(pvals) {
    pvals <- as.numeric(pvals)
    pvals <- pvals[!is.na(pvals)]
    if (length(pvals) == 0) {
        return(NA)
    } else if (length(pvals) == 1) {
        return(pvals)   # keep the single value if only one
    } else {
        return(sumlog(pvals)$p)
    }
}

# ---- 2. Column name mapping ----
colmap <- c(
  "ID"         = "ID",
  "Description"= "Description",
  "GeneRatio"  = "GeneRatio",
  "BgRatio"    = "BgRatio",
  "pvalue"     = "pvalue",
  "p.adjust"   = "p.adjust",
  "qvalue"     = "qvalue",
  "geneID"     = "geneID",
  "Count"      = "Count"
)

# ---- 3. Loop over all reps ----
```

```
rep_dirs <- list.dirs(input_parent, recursive = FALSE, full.names = TRUE)
rep_dirs <- rep_dirs[grepl("rep[0-9]+$", rep_dirs)]  # keep only rep1, rep2, etc.

cat("Found", length(rep_dirs), "replicate folders.\n")

for (rep_dir in rep_dirs) {
    rep_name <- basename(rep_dir)
    output_file <- file.path(output_parent, paste0(rep_name, "_combined.csv"))

    # --- Read all CSVs inside this rep ---
    csv_files <- list.files(rep_dir, pattern = "KEGG_bin_.*\\.csv$", full.names =
TRUE)
    if (length(csv_files) == 0) {
        cat("Skipping", rep_name, "-> No CSV files found.\n")
        next
    }

    all_results <- lapply(csv_files, function(f) {
        df <- read.csv(f, stringsAsFactors = FALSE, check.names = FALSE)
        colnames(df) <- trimws(colnames(df))

        # Create aligned output with only mapped columns
        out <- data.frame(matrix(ncol = length(colmap), nrow = nrow(df)))
        colnames(out) <- names(colmap)

        for (nm in names(colmap)) {
            src <- colmap[[nm]]
            if (src %in% colnames(df)) {
                out[[nm]] <- df[[src]]
            } else {
                out[[nm]] <- NA
            }
        }
        return(out)
    }) %>% bind_rows(.id = "bin_id")

    cat(rep_name, "-> Normalized input:", nrow(all_results),
        "rows across", length(csv_files), "files.\n")

    # --- Aggregate across bins ---
    combined <- all_results %>%
      group_by(ID, Description) %>%
      summarise(
        bins_present     = n(),
        combined_p       = combine_fisher(pvalue),
        combined_padjust = combine_fisher(p.adjust),
        combined_q       = combine_fisher(qvalue),
        geneID           = paste(unique(na.omit(geneID)), collapse = "; "),
        total_count      = sum(as.numeric(Count), na.rm = TRUE),
        GeneRatio        = paste(unique(na.omit(GeneRatio)), collapse = "; "),
```

```
        BgRatio          = paste(unique(na.omit(BgRatio)), collapse = "; "),
        .groups = "drop"
      ) %>%
      arrange(combined_p)

    # --- Save ---
    write.csv(combined, output_file, row.names = FALSE)
    cat("Saved:", output_file, "\n")
}


#!/usr/bin/env Rscript

# ==========================================
# Combine KEGG Enrichment Results across Replicates
# ==========================================

# ---- Load libraries ----
library(dplyr)
library(purrr)
library(metap)   # for Fisher’s method
library(stringr)

# ---- User Input ----
input_dir <- "/NGF/ms_figures/FigS5/glu_300_del/g_del_300_combined"
output_file <- file.path(input_dir,
"g_del_300_combined_final_combined_replicates.csv")

# ---- 1. Read all replicate CSVs ----
csv_files <- list.files(input_dir, pattern = "^rep[0-9]+_combined\\.csv$",
full.names = TRUE)

if (length(csv_files) == 0) {
  stop("No replicate combined CSV files found in: ", input_dir)
}

all_data <- lapply(csv_files, function(f) {
  df <- read.csv(f, stringsAsFactors = FALSE, check.names = FALSE)
  df$source_file <- basename(f)   # track which replicate
  return(df)
}) %>% bind_rows()

cat("Loaded", nrow(all_data), "rows from", length(csv_files), "replicate CSVs.\n")

# ---- 2. Fisher’s method helper ----
combine_fisher <- function(pvals) {
  pvals <- as.numeric(pvals)
  pvals <- pvals[!is.na(pvals)]
  if (length(pvals) == 0) {
    return(NA)
```

```
  } else if (length(pvals) == 1) {
    return(pvals)   # keep the single value
  } else {
    return(sumlog(pvals)$p)
  }
}

# ---- 3. Aggregate across replicates ----
final_combined <- all_data %>%
  group_by(ID, Description) %>%
  summarise(
    bins_present     = sum(bins_present, na.rm = TRUE),
    combined_p       = combine_fisher(combined_p),
    combined_padjust = combine_fisher(combined_padjust),
    combined_q       = combine_fisher(combined_q),
    geneID           = paste(unique(unlist(strsplit(paste(geneID, collapse = "; "),
"; "))), collapse = "; "),
    total_count      = sum(as.numeric(total_count), na.rm = TRUE),
    GeneRatio        = paste(unique(na.omit(GeneRatio)), collapse = "; "),
    BgRatio          = paste(unique(na.omit(BgRatio)), collapse = "; "),
    .groups = "drop"
  ) %>%
  arrange(combined_p)

# ---- 4. Save results ----
write.csv(final_combined, output_file, row.names = FALSE)
cat("Final combined results saved to:", output_file, "\n")


#!/usr/bin/env Rscript

# ==========================================
# KEGG Enrichment Plots (Dotplot & Cnetplot)
# ==========================================

# ---- Load libraries ----
library(ggplot2)
library(dplyr)
library(stringr)
library(igraph)
library(ggraph)
library(tidyr)

# ---- User Input ----
input_file  <-
"/NGF/ms_figures/FigS5/glu_300_del/g_del_300_combined/g_del_300_combined_final_combi
ned_replicates.csv"
output_dir  <- "/NGF/ms_figures/FigS5/glu_300_del/g_del_300_combined/"

if (!dir.exists(output_dir)) dir.create(output_dir, recursive = TRUE)
```

```r
# ---- 1. Load data ----
df <- read.csv(input_file, stringsAsFactors = FALSE, check.names = FALSE)

# ---- 1a. Filter rows with bins_present >= 2 ----
df <- df %>% filter(bins_present >= 2)

# ---- 2. Bar plot (Top 10 by combined_padjust, cutoff < 0.05) ----
df_bar <- df %>%
  filter(combined_padjust < 0.05) %>%
  arrange(combined_padjust) %>%
  head(10) %>%
  # Calculate unique gene count from geneID column
  rowwise() %>%
  mutate(gene_count = length(unique(unlist(str_split(geneID, "/|;\\s*"))))) %>%
  ungroup()

# Reorder factor for decreasing order (most significant on top)
df_bar$Description <- factor(df_bar$Description, levels =
rev(df_bar$Description[order(df_bar$combined_padjust)]))

# ---- 3. Plot ----
# ---- 3. Plot ----
p <- ggplot(df_bar, aes(x = Description, y = gene_count, fill = combined_padjust)) +
  geom_bar(stat = "identity") +
  scale_fill_gradientn(
    colours = c("#FF6666", "#FFCC66", "#66CC99", "#6699CC"),
    values = scales::rescale(c(0, 0.01, 0.03, 0.05)),
    limits = c(0, 0.05),
    breaks = c(0, 0.01, 0.02, 0.05),
    labels = c("0", "0.01", "0.02", "0.05"),
    name = "Adjusted p-value"
  ) +
  coord_flip() +
  labs(title = "300 generations",x = "Pathway", y = "Unique Gene Count") +
  theme_minimal(base_size = 12, base_family = "sans") +
  theme(
    plot.title = element_text(size = 14, face = "bold", hjust = 0.5),
    axis.title.x = element_text(size = 16, face = "bold", color = "black"),
    axis.title.y = element_text(size = 18, face = "bold", color = "black"),
    axis.text.x  = element_text(size = 14, color = "black"),
    axis.text.y  = element_text(size = 14, color = "black")
  )

# Save PNG
png(file.path(output_dir, "g_del_300_kegg_barplot.png"), width = 10, height = 6,
units = "in", res = 300)
print(p)
dev.off()
```

14. Plot heatmap for opposing CNVs

```
#!/usr/bin/env Rscript

library(readxl)
library(dplyr)
library(tidyr)
library(ggplot2)
library(grid)

#=========================
# INPUT FILES
#=========================

del_file <- "/NGF/ms_figures/RD_deletion_gene_list.xlsx"
dup_file <- "/NGF/ms_figures/RD_duplication_gene_list.xlsx"

#=========================
# OUTPUT DIRECTORY
#=========================

output_dir <- "/NGF/ms_figures/Fig4"
dir.create(output_dir, showWarnings = FALSE, recursive = TRUE)

#=========================
# READ DATA
#=========================

del_raw <- read_excel(del_file, col_names = FALSE)
dup_raw <- read_excel(dup_file, col_names = FALSE)

sample_ids <- as.character(del_raw[2, ])
env_ids    <- as.character(del_raw[3, ])
timepoints <- rep(c("300","600","900","1200"), each = 12)

meta <- data.frame(
  sample = sample_ids,
  env = env_ids,
  time = timepoints,
  stringsAsFactors = FALSE
)

# Remove header rows
del <- del_raw[-c(1:3), ]
dup <- dup_raw[-c(1:3), ]

colnames(del) <- sample_ids
colnames(dup) <- sample_ids
```

```
#=========================
# LONG FORMAT
#=========================

del_long <- del %>%
  pivot_longer(cols = everything(), names_to="sample", values_to="gene") %>%
  filter(!is.na(gene) & gene != "") %>%
  distinct(sample, gene) %>%
  left_join(meta, by="sample")

dup_long <- dup %>%
  pivot_longer(cols = everything(), names_to="sample", values_to="gene") %>%
  filter(!is.na(gene) & gene != "") %>%
  distinct(sample, gene) %>%
  left_join(meta, by="sample")

#=========================
# DEFINE GENE SETS
#=========================

# Required sets
glu_del_genes <- del_long %>%
  filter(grepl("glu", env)) %>%
  pull(gene) %>% unique()

gal_dup_genes <- dup_long %>%
  filter(grepl("gal", env)) %>%
  pull(gene) %>% unique()

# Forbidden sets
glu_dup_genes <- dup_long %>%
  filter(grepl("glu", env)) %>%
  pull(gene) %>% unique()

gal_del_genes <- del_long %>%
  filter(grepl("gal", env)) %>%
  pull(gene) %>% unique()

#=========================
# STRICT FILTERING
#=========================

strict_genes <- intersect(glu_del_genes, gal_dup_genes)

strict_genes <- setdiff(strict_genes, glu_dup_genes)
strict_genes <- setdiff(strict_genes, gal_del_genes)

cat("Strict genes retained:", length(strict_genes), "\n")

#=========================
```

```
# COUNT MATRICES
#=========================

del_counts <- del_long %>%
  filter(gene %in% strict_genes, grepl("glu", env)) %>%
  group_by(gene, time) %>%
  summarise(count=n(), .groups="drop") %>%
  pivot_wider(names_from=time, values_from=count, values_fill=0)

dup_counts <- dup_long %>%
  filter(gene %in% strict_genes, grepl("gal", env)) %>%
  group_by(gene, time) %>%
  summarise(count=n(), .groups="drop") %>%
  pivot_wider(names_from=time, values_from=count, values_fill=0)

colnames(del_counts)[-1] <- paste0("Glu_",colnames(del_counts)[-1])
colnames(dup_counts)[-1] <- paste0("Gal_",colnames(dup_counts)[-1])

#=========================
# MERGE MATRICES
#=========================

matrix <- full_join(del_counts, dup_counts, by="gene")
matrix[is.na(matrix)] <- 0

# Make gal duplications negative
matrix[,grep("Gal_", colnames(matrix))] <- -matrix[,grep("Gal_", colnames(matrix))]

#=========================
# SAVE MATRIX
#=========================

write.table(matrix,
            file.path(output_dir,"gene_CNV_matrix.tsv"),
            sep="\t",
            row.names=FALSE,
            quote=FALSE)

#=========================
# HEATMAP MATRIX
#=========================

mat <- as.matrix(matrix[,-1])
rownames(mat) <- matrix$gene

mat <- mat[,c("Glu_300","Glu_600","Glu_900","Glu_1200",
              "Gal_300","Gal_600","Gal_900","Gal_1200")]

#=========================
# CLUSTER
```

```
#=========================

d <- dist(mat)
hc <- hclust(d, method="ward.D2")
row_order <- hc$order

#=========================
# LONG FORMAT
#=========================

df_long <- as.data.frame(mat) %>%
  mutate(Row=factor(rownames(mat), levels=rownames(mat)[row_order])) %>%
  pivot_longer(-Row, names_to="Condition", values_to="Value") %>%
  mutate(
    Generation=factor(sub(".*_(\\d+)", "\\1", Condition),
                      levels=c("300","600","900","1200")),

    Environment=factor(ifelse(grepl("^Glu",Condition),
                              "Glucose deletions",
                              "Galactose duplications"),
                       levels=c("Glucose deletions","Galactose duplications")),

    Gen_Env=factor(paste(Environment,Generation,sep="_"),
                   levels=c("Glucose deletions_300","Glucose deletions_600",
                            "Glucose deletions_900","Glucose deletions_1200",
                            "Galactose duplications_300","Galactose
duplications_600",
                            "Galactose duplications_900","Galactose
duplications_1200"))
  )

#=========================
# HEATMAP
#=========================

p <- ggplot(df_long, aes(x=Gen_Env, y=Row, fill=Value)) +
  geom_tile(color="black", size=0.2) +

  scale_fill_gradient2(
    low="blue",
    mid="white",
    high="red",
    midpoint=0,
    limits=c(-6,6),
    breaks=c(-6,0,6),
    labels=c("Duplication","No change","Deletion"),
    name=""
  ) +

  scale_x_discrete(labels=rep(c("300","600","900","1200"),2)) +
```

```
  labs(x="Time (generations)", y="Genes") +

  theme_minimal(base_size=14) +
  theme(
    axis.ticks.y=element_blank(),
    panel.grid=element_blank(),
    axis.text.x=element_text(size=18,face="bold"),
    axis.text.y=element_text(size=6, face="italic"),
    axis.title=element_text(face="bold"),
    legend.text=element_text(size=11),
    plot.margin=margin(10,10,30,10)
  ) +

  annotate("text",
           x=c(2.5,6.5),
           y=-Inf,
           label=c("Glucose","Galactose"),
           fontface="bold", size=5, vjust=4) +

  coord_cartesian(clip="off")

#==========================
# SAVE HEATMAP
#==========================

ggsave(file.path(output_dir,"CNV_heatmap_grouped.png"),
       p,
       width=10,
       height=40,
       dpi=600)

#==========================
# SAVE GENE LIST
#==========================

gene_list <- rownames(mat)[row_order]

write.table(gene_list,
            file.path(output_dir,"heatmap_genes.txt"),
            quote=FALSE,
            row.names=FALSE,
            col.names=FALSE)

#==========================
# SUMMARY
#==========================

cat("Total strict genes:", length(gene_list), "\n")
cat("Results saved in:", output_dir, "\n")
```

```
---
#Linear model/
import pandas as pd
import statsmodels.api as sm
import statsmodels.formula.api as smf
================================
1. Load data
================================
file_path = r"C:\Users\supreet\OneDrive\Desktop\fitness_template.xlsx"  # update as
needed
df = pd.read_excel(file_path)
================================
2. Create ancestor lookup
================================
Filter ancestor rows
ancestor_df = df[df['lineage'] == 'ancestor'].copy()
Build a lookup dictionary based on precond1_env, recovery_gens, and assay_env
ancestor_lookup = {}
for _, row in ancestor_df.iterrows():
key = (row['precond1_env'], row['recovery_gens'], row['assay_env'])
ancestor_lookup[key] = row['fitness_composite']
Map ancestor fitness to all rows
def get_ancestor(row):
key = (row['precond1_env'], row['recovery_gens'], row['assay_env'])
if key not in ancestor_lookup:
raise KeyError(f"No ancestor found for key: {key}")
return ancestor_lookup[key]
df['ancestor'] = df.apply(get_ancestor, axis=1)
================================
3. Fit linear model
================================
Include ancestor fitness as covariate, plus all main effects and interactions
formula = 'fitness_composite ~ ancestor + C(precond1_env) * C(recovery_gens) *
evolution_gen'
model = smf.ols(formula, data=df).fit()
================================
4. ANOVA table
================================
anova_table = sm.stats.anova_lm(model, typ=2)
print("=== ANOVA Table ===\n")
print(anova_table)
================================
5. Effect sizes (η²)
================================
anova_table['eta_sq'] = anova_table['sum_sq'] / anova_table['sum_sq'].sum()
print("\n=== Effect sizes (η²) ===\n")
print(anova_table[['sum_sq', 'eta_sq', 'PR(>F)']])
================================
6. Linear regression summary
```

```
================================
print("\n=== Linear Regression Summary ===\n")
print(model.summary())
================================
7. Save predicted fitness and coefficients
================================
df['predicted_fitness'] = model.predict(df)
df[['precond1_env', 'recovery_gens', 'evolution_gen', 'assay_env',
'ancestor', 'fitness_composite', 'predicted_fitness']].to_excel(
"predicted_fitness_and_ancestor.xlsx", index=False
)
model.params.to_frame(name='coefficient').to_excel("model_coefficients.xlsx")
print("\n Predicted fitness and coefficients saved successfully.")
```
